# FGFR1 is critical for *Rbl2* loss-driven tumor development and requires PLCG1 activation for continued growth of small cell lung cancer

**DOI:** 10.1101/796607

**Authors:** Kee-Beom Kim, Youngchul Kim, Dong-Wook Kim, Kwon-Sik Park

## Abstract

Small cell lung cancer (SCLC) remains a recalcitrant disease; limited therapeutic options have not improved overall survival and approved targeted therapies are lacking. Amplification of the tyrosine kinase receptor FGFR1 (fibroblast growth factor receptor 1) is one of the few actionable alterations found in SCLC genome. However, efforts to develop targeted therapies for *FGFR1*-amplified SCLC are hindered by critical gaps in knowledge around the molecular origins and mediator of FGFR1-driven signalling and the physiological impact of targeting FGFR1. Here we demonstrate the oncogenic impact of increased FGFR1 on the malignant progression of precancerous cells and the necessity of nonamplified *FGFR1* for SCLC development and homeostasis. Unexpected dependency of *Rbl2* loss-driven tumor development on FGFR1 in autochthonous mouse models reveals a novel function of p130 (encoded by *Rbl2*), a member of the RB family of tumor suppressors, as a regulator of FGFR1 expression. Additionally, FGFR1-dependent SCLC cells require activation of PLCG1 for continued growth. Together, this study uncovers mechanisms of FGFR1-driven SCLC that involves p130 upstream and PLCG1 downstream and provides potential biomarkers for anti-FGFR1 therapeutic strategy.

## Introduction

SCLC accounts for approximately 13% of all lung cancer and remains a recalcitrant disease. The standard chemotherapy of cisplatin and etoposide has not significantly improved overall survival in SCLC patients and the development of targeted therapies has been challenging due to the paucity of clinically viable targets (1). SCLC is mostly driven by loss of tumor suppressors, including near universal inactivation of RB and p53 and frequent loss of other tumor suppressors such as the RB family members p107 and p130 (coded by the *RBL1* and *RBL2*, respectively) (2, 3). Unfortunately, these loss-of-function alterations are not directly actionable. Although oncogenic alterations are less frequent in SCLC, gene amplifications of *FGFR1* are detected in approximately 6% of SCLC tumors and high FGFR1 expression is also detected in a subset of tumors without the gene amplification (2-6). Importantly, an FGFR-selective tyrosine kinase inhibitor (TKI) can inhibit the growth of SCLC cell lines with high FGFR1 expression (7-9). However, *FGFR1* amplification and high FGFR1 levels alone do not guarantee sensitivity to FGFR inhibitors (7-9). Intracellular mediators of FGFR1 signaling as well as other genetic alterations may modulate sensitivity to treatment, but these mechanisms remain poorly defined. For example, while the MEK-ERK pathway mediates oncogenic FGFR1 signaling in some SCLC cell lines, a hyperactive RAF-MEK-ERK pathway is tumor suppressive in others (10, 11). In addition, while AKT, STAT, and PLCG1 act as transducers downstream of FGFR1 during organ development, little is known of their roles in SCLC (12). Importantly, physiological significance of FGFR1, either amplified or nonamplified, in SCLC has not formally been determined using autochthonous model. The oncogenic role has also recently been challenged by a study in which a constitutively active form of FGFR1 suppressed SCLC development in *Rb1/Trp53*-mutant genetically engineered mouse model (*Rb1/Trp53*-GEMM) (13). Thus, there are critical gaps in knowledge that need to be addressed not only to target FGFR1 in SCLC, but also to identify SCLC patients in ongoing clinical trials who would benefit from anti-FGFR1 therapies.

Here, we characterize FGFR1 in SCLC using *Rb1/Trp53*-GEMM and its variants in which adenoviral Cre (Ad-Cre)-mediated deletion of *Rb1* and *Trp53* results in lung tumors recapitulating major histopathological features of human SCLC (14,15). We also introduced genetic and chemical perturbations in primary tumor cells (mSCLC) as well as precancerous neuroendocrine cells (preSC) derived from the *Rb1/Trp53*-GEMM and that can transform into SCLC upon activation of oncogenic drivers (16). Our data show that FGFR1 is required for the development and growth of *Rbl2* loss-driven SCLC, and that this dependency is due to the induction of FGFR1 expression. Furthermore, FGFR1 promotes tumor development and growth via activation of PLCG1 and alteration in neural differentiation. Taken together, our findings provide insight into the mechanism of SCLC progression and potential biomarkers for identifying patients appropriate for anti-FGFR1 therapies.

## Materials and Methods

### Plasmids and chemicals

All plasmids used in this study are listed in Supplementary Table 1. The sequences of shRNAs and gRNAs are listed in Supplementary Table 2 and 3. *Fgfr1* cDNA was cloned using total RNAs from murine SCLC cells and verified by Sanger sequencing. U73122 and U73343 were obtained from Cayman and PD173074 from Sigma.

### Cell culture

Murine SCLC cells (mSCLC) and precancerous cells (preSC) were derived from lung tumors and early-stage neuroendocrine lesions, respectively, developed in the *Rb1/Trp53-*GEMM (16). Human SCLC cell lines (NCI-H82, NCI-H209, NCI-H524, also designated simply as H82, H209, H524) were obtained from ATCC. 293T cells were cultured in DMEM media and human and mouse cells were cultured in RPMI1640 media, both supplemented with 10% bovine growth serum (GE Healthcare) and 1% penicillin-streptomycin (Invitrogen). Puromycin (Thermo Fisher) was used to select stably transduced cells following lentiviral infection. For lentivirus production we transfected lentiviral plasmids with packaging plasmids in 293T cells using polyethylenimine (Sigma), harvested supernatants containing viral particles 48 and 72 hours later, filtered through 0.45μm PVDF filter before adding to culture of target cells in the presence of 5μg/ml polybrene (Sigma). To induce FGFR1 expression in *Fgfr1-preSC*, cells were treated with doxycycline (0.2ug/mL). CRISPR-mediated knockout of *Rbl2* was achieved by transient transfection of Cas9 and gRNA in a single vector using Lipofectamine 2000 (Invitrogen). Forty-eight hours later, transfected cells were enriched by FACS sorting for red fluorescent protein expressed from the same vector. Sanger sequencing verified mutation in the target sequence in *Rbl2* gene and immunoblot validated loss of p130 coded by *Rbl2*. MTT assay was performed to measure cell viability using thiazolyl blue tetrazolium bromide (Sigma). Cells were seeded at 1 × 10^4^ per well in 96-well plates at day 0, and MTT reagents were added on day 4. O.D values were determined at a wavelength of 590 nm using an ELISA reader (Thermo Scientific) The percentage survival was determined as the ratio of treated cells versus vehicle control after background subtraction. Soft-agar assay was performed by seeding 1×10^4^ cells per well in 12-well plates as previously described (16). Images of wells are acquired using Olympus MVX10 scope, and colonies from the whole field of image were counted using the imaging software NIS-Elements Basic Research (Nikon). All of the cell culture experiments were performed in triplicates and repeated for a minimum of two biological replicates.

### Tumor induction and allograft

*Rb1*^*lox/lox*^; *Trp53*^*lox/lox*^ (*Rb1/Trp53*-GEMM) and *Rb1*^*lox/lox*^; *Trp53*^*lox/lox*^; *Rbl2*^*lox/lox*^ (*Rb1/Trp53/Rbl2-*GEMM) have been previously described (14, 16). Mouse strains with *Fgfr1*^*lox*^ and *Fgfr2*^*lox*^ alleles have been previously described (17, 18). For tumor induction, lungs of 10-week-old mice were infected with adenoviral Cre via intratracheal instillation as previously described and mice were aged 6 months (19). For allograft experiment, we inject 5.0×10^5^ control or *Fgfr1*-preSC in the flanks of nude mice (*Foxn1*^*nu/nu*^; Envigo) and 1.0×10^6^ *Fgfr1*^*lox/lox*^ *Rb1/Trp53/Rbl2* murine cells infected with Ad-Cre in the flanks of B6/129S F1 hybrid mice (Jackson Laboratory). To induce FGFR1 expression in *Fgfr1*-preSC following implantation, mice were fed doxycycline diet (625mg/kg, Envigo). Mice were maintained according to guidelines from the National Institutes of Health. Animal procedures were approved by the Animal Care and Use Committee at the University of Virginia.

### Histology, immunohistochemistry, and immunoblot

Mouse tissues were fixed in 4% paraformaldehyde in phosphate-buffered saline before paraffin embedding. Five micrometer-thick tissue sections were stained with hematoxylin and eosin staining and immunostaining as previously described (16). For quantification of pHH3-positive cells, tumors of similar size and area were included. For immunoblot analysis, protein was extracted from mouse tumor tissues and human and murine cell lines in RIPA buffer. All primary and secondary antibodies used are listed in Supplementary Table 4. Macroscopic images of lung sections were acquired using Olympus MVX10. Images of H&E and immunostained tissues were acquired using Nikon Eclipse Ni-U microscope. Image analysis and automated quantification were performed using NIS-Elements Basic Research (Nikon)

### RNA sequencing and analysis

Sequencing libraries were generated by the Genome Analysis and Technology Core at University of Virginia using oligo dT-purified mRNA from 500ng of total RNA and the NEB Next Ultra RNA library preparation kit (New England Biolabs), and 50-bp single-end sequencing was performed on Illumina NextSeq 500 platform (Illumina). The sequencing data have been deposited to the SRA Database (https://www.ncbi.nlm.nih.gov/sra, identifier: PRJNA564798). Reads were mapped to the Mus musculus genome assembly GRCm38 (mm10) using TopHat and counts of reads map to each gene were obtained using HTSeq (20, 21). Differentially expressed genes regulated by FGFR1 were identified using DESeq2 package (22). A regularized log-transformation was applied to the read count data of all sequenced samples and resultant data were used for gene ontology (GO) term analysis at DAVID Bioinformatics Resources (https://david.ncifcrf.gov/home.jsp).

### Statistical analysis

Using GraphPad Prism, the results were presented as the mean ± standard deviation. and evaluated using an unpaired two-tailed Student’s t-test. p<0.05 was considered statistically significant.

## Results

### Ectopic FGFR1 promotes progression of precancerous cells to SCLC

The lung tumors in *Rb1/Trp53*-GEMM, unlike a subset of human SCLC tumors, have not shown *Fgfr1* amplification (3, 23). Instead, cell derivatives of these lung tumors (murine SCLC, mSCLC) showed markedly higher levels of *Fgfr1* mRNA and FGFR1 protein than precancerous cells (preSC) (Fig. 1A). To begin determining the functional impact of these alterations on tumor development, we expressed FGFR1 in preSC using a lentiviral vector and tested the transduced cells for tumorigenic potential. Compared with control preSC infected with empty vector, *Fgfr1*-preSC gave rise to more colonies in soft agar and more rapid development of subcutaneous tumors in immunocompromised nude mice (Fig. 1B, C). These subcutaneous tumors displayed histological features of SCLC including distinct small-cell morphology with scanty cytoplasm and positive staining for the neuroendocrine marker calcitonin gene-related peptide (CGRP) (Fig. 1D). These data suggest that FGFR1 overexpression is sufficient to promote tumorigenic transformation of precancerous precursors.

**Fig. 1.**
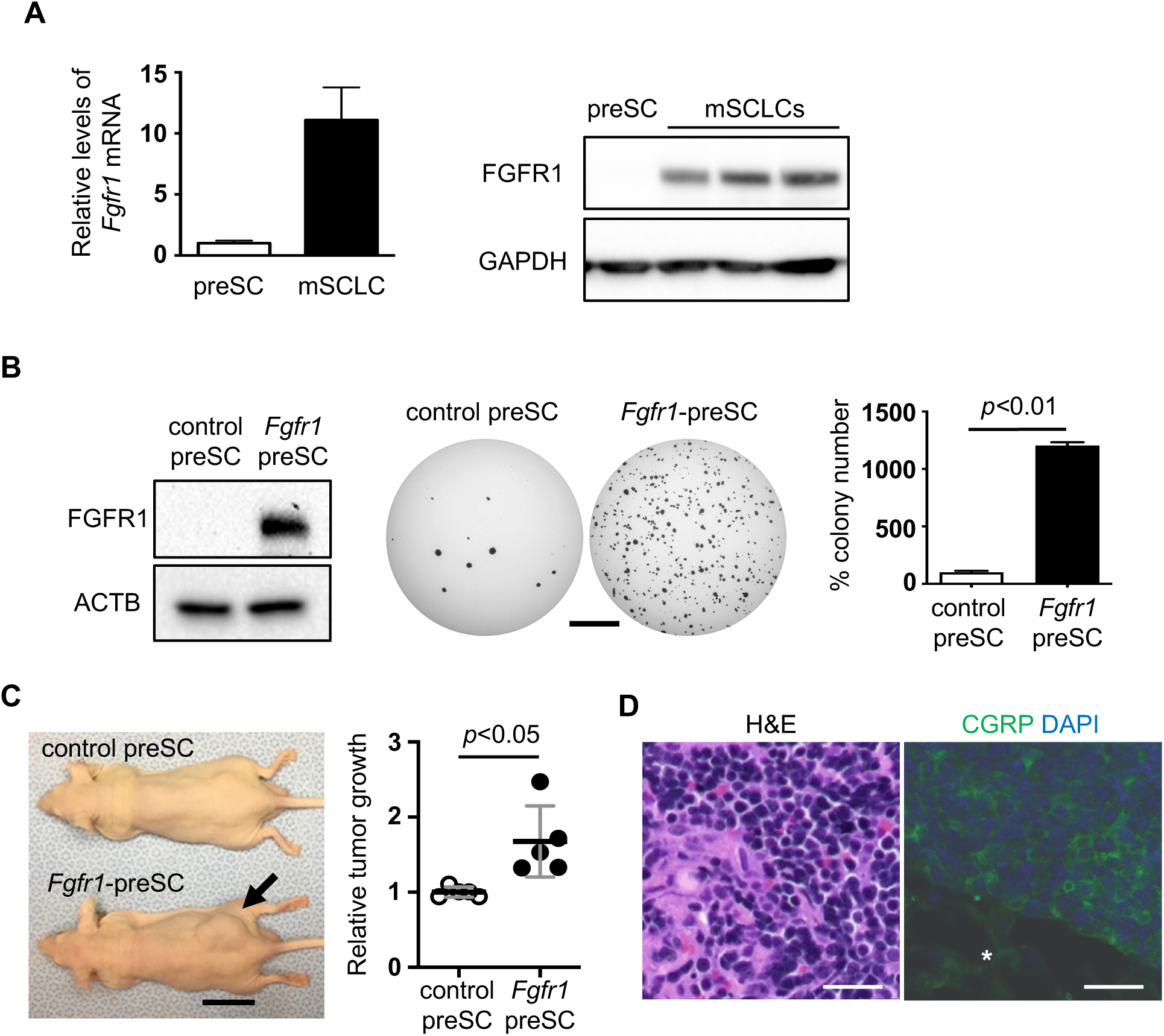
Increased FGFR1 promotes SCLC development. **(A)** RT-qPCR and immunoblot for *Fgfr1* mRNA normalized to *ARBPP0* and FGFR1 protein in preSC and mSCLC (n=3 replicates per cell type). GAPDH blot verifies equal loading of total proteins. (**B)** Left, expression of FGFR1 in preSC infected with empty vector (control preSC) or an FGFR1 expression construct (*Fgfr1*-preSC); ACTB blot verifies equal loading of total proteins. Middle, representative images of soft agar colonies formed by control and *Fgfr1-*preSC. Right, quantification of colonies >0.2mm in diameter (unpaired t-test n=3 replicates per cell type). (**C)** Left, representative image of subcutaneous tumors (arrow) in athymic nude mice (n=5 mice per cell type). Right, quantification of tumor development of subcutaneous tumors >1.5cm in diameter (unpaired t-test n=5 mice per cell type); Relative tumor growth represents tumor weight (g) divided by latency (days after allograft). (**D)** Representative image of hematoxylin-eosin (H&E) stained section of *Fgfr1*-preSC-derived subcutaneous tumors and, right, of immunostaining for calcitonin gene-related peptide (CGRP, green). DAPI stains for nuclei (blue). Asterisk indicates non-tumor area. Statistical tests performed using unpaired t-test (ns: not significant). Scale bars: B, 5mm; D, 100μm. Error bars represent standard deviation.

### Deletion of FGFR1 suppresses Rbl2-deficient SCLC

The physiological significance of FGFR1 in SCLC has not been formally validated in autochthonous model. To begin testing whether FGFR1 is required for SCLC development *in vivo*, we deleted *Fgfr1* in *Rb1/Trp53*-GEMM carrying *Fgfr1*^*lox/lox*^ alleles using intratracheal instillation of Ad-Cre. We found that there was no significant difference in tumor burden (tumor area/lung area) between *Fgfr1*^*Δ/Δ*^ and *Fgfr1*^*+/+*^ *Rb1*/*Trp53* controls (Fig. 2A). This indicates that in *Rb1*/*Trp53-*mutant mSCLC, FGFR1 is likely dispensable for tumor development even though its overexpression was sufficient for tumorigenic progression of *Rb1*/*Trp53-*mutant preSC (Fig. 1). To validate this finding, we performed a similar experiment using *Rb1*/*Trp53*/*Rbl2*-GEMM in which tumor development is initiated by the loss of *Rbl2* (encoding p130, an RB family member) in addition to *Rb1* and *Trp53* (24). This model represents a subset of human SCLC with loss of all three genes and develops more than a dozen tumors with latency of 6 months as opposed to 1-2 tumors with latency of 9-12 months in *Rb1*/*Trp53-*GEMM, thus providing a robust experimental system for measuring the effects of genetic or pharmacological manipulation (25, 26). Intriguingly, *Fgfr1*^*Δ/Δ*^ *Rb1*/*Trp53*/*Rbl2* mice had significantly less tumor burden than *Fgfr1*^*+/+*^ *Rb1*/*Trp53*/*Rbl2* mice (Fig. 2B). Further, lung tumors in the in the *Fgfr1*^*Δ/Δ*^ *Rb1*/*Trp53*/*Rbl2* mice showed significantly less staining for phosphorylated histone H3 (pHH3), a marker for mitotic cell, than tumors in *Fgfr1*^*+/+*^ *Rb1*/*Trp53*/*Rbl2* mice (Fig. 2C). These findings suggest that loss of FGFR1 suppresses tumor development through reduced proliferation.

**Fig. 2.**
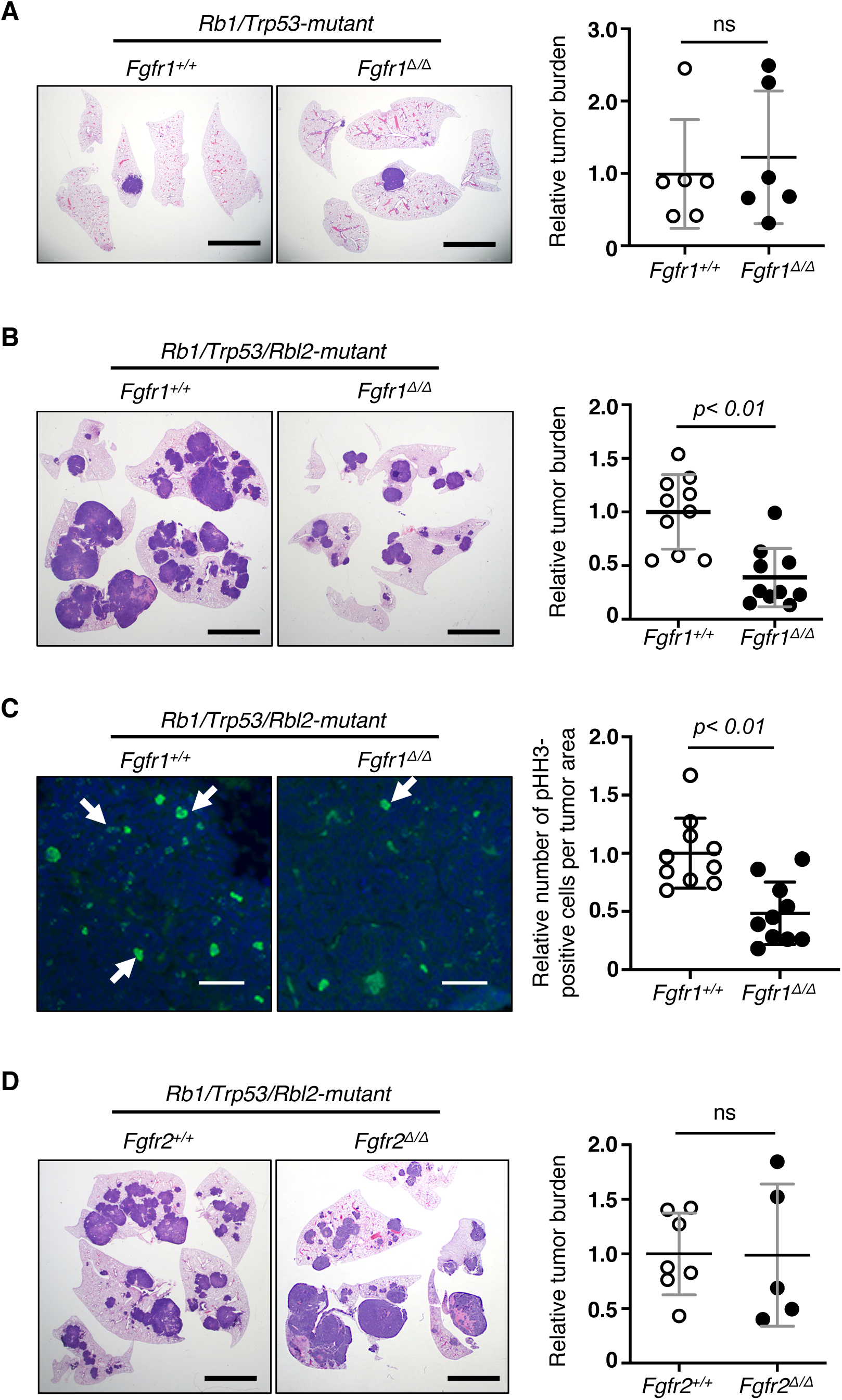
Deletion of *Fgfr1* suppresses SCLC development in vivo. Left, representative images of H&E stained or immunostained sections of the lungs from different SCLC GEMMs and right, quantification of tumor burden (tumor area/lung area) or phosphorylated histone H3 (pHH3)-positive cells per tumor area in **(A)** *Fgfr1*^*+/+*^ *vs. Fgfr1*^*Δ/Δ*^ *Rb1/Trp53* (n=6 and n=6, respectively), **(B**, **C)** *Fgfr1*^*+/+*^ *vs. Fgfr1*^*Δ/Δ*^ *Rb1/Trp53/Rbl2* (n=10 and n=10, respectively); in panel **C**, pHH3 stain in green and DAPI in blue. Arrows indicate pHH3-positive nuclei, **(D)** *Fgfr2*^*+/+*^ *vs. Fgfr2*^*Δ/Δ*^ *Rb1/Trp53/Rbl2* n=7 and n=5, respectively). Statistical tests performed using unpaired t-test (ns: not significant). Error bars represent standard deviation. Scale bars: A, B, D, 5mm; C, 50μm.

To determine whether the decrease in tumor burden is truly due to loss of FGFR1, we validated the recombination of all floxed alleles using genotyping PCR and immunoblot of primary tumor cells from *Fgfr1*^*Δ/Δ*^ *Rb1*/*Trp53*/*Rbl2* mice. Primary cells from one of four randomly selected *Fgfr1*^*Δ/Δ*^ tumors expressed FGFR1 at a level comparable to *Fgfr1*^*+/+*^ tumor cells (Fig. 3A; Supplementary Fig. S1), indicating incomplete recombination and retention of one *Fgfr1*^*lox*^ allele.

**Fig. 3.**
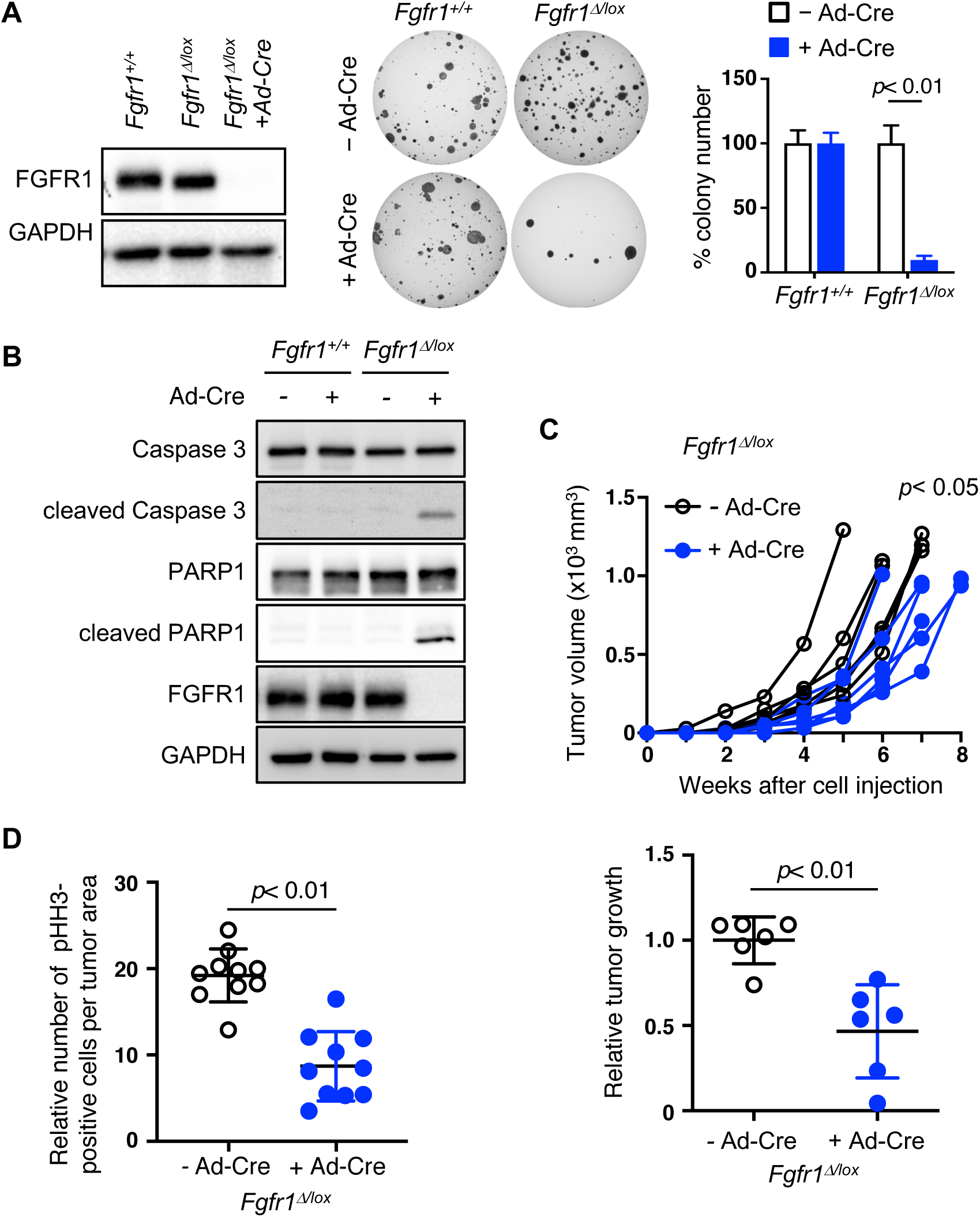
FGFR1 is critical for continued growth of tumor cells. **(A)** Left, immunoblot for FGFR1 in primary cells derived from the lung tumors in *Fgfr1*^*+/+*^ and *Fgfr1*^*lox/lox*^ *Rb1/Trp53/Rbl2-*GEMM infected with Ad-Cre. The cells from one of the *Fgfr1*^*lox/lox*^ mice show little reduction of FGFR1, indicating incomplete recombination of floxed *Fgfr1* alleles (*Fgfr1*^*Δ/lox*^). Following Ad-Cre infection, these cells completely lose FGFR1 expression. Middle, representative images of soft agar colonies formed by *Fgfr1*^*+/+*^ and *Fgfr1*^*Δ/lox*^ *Rb1/Trp53/Rbl2* cells in soft agar 3 weeks following Ad-Cre infection. Right, quantification of colonies >0.2mm in diameter (n=3 replicates per cell type). **(B)** Immunoblot for cleaved and total Caspase 3 and PARP1 in *Fgfr1*^*+/+*^ and *Fgfr1*^*Δ/lox*^ *Rb1/Trp53/Rbl2* cells with or without Ad-Cre infection. Cleaved proteins are a marker for apoptosis and FGFR1 blot verifies loss of FGFR1 expression in Ad-Cre infected *Fgfr1*^*Δ/lox*^ cells. GADPH blot verifies equal loading of total proteins. **(C)** Top, volumes of tumors (n=6) generated from subcutaneous injection of *Fgfr1*^*Δ/lox*^ *Rb1/Trp53/Rbl2* cells with or without Ad-Cre infection. Bottom, quantification of tumor development of subcutaneous tumors >1.5cm in diameter; relative tumor growth represent tumor weight (g) divided by latency (days after allograft). **(D)** Quantification of pHH3 staining in allograft tumors derived from *Fgfr1*^*Δ/lox*^ *Rb1/Trp53/Rbl2* cells with or without Ad-Cre infection. Statistical tests performed using unpaired t-test. Error bars represent standard deviation.

Subsequently, these *Fgfr1*^*Δ/lox*^ cells were infected with Ad-Cre in culture, completely lost FGFR1, and gave rise to fewer colonies in soft agar than uninfected *Fgfr1*^*Δ/lox*^ cells or Ad-Cre-infected *Fgfr1*^*+/+*^ cells (Fig. 3A). Immunoblot showed that levels of cleaved Caspase 3 and PARP1 indicative for apoptosis markedly increased in Ad-Cre-infected *Fgfr1*^*Δ/lox*^ cells relative to uninfected *Fgfr1*^*Δ/lox*^ cells and Ad-Cre-infected *Fgfr1*^*+/+*^ cells (Fig. 3B). Ad-Cre-infected *Fgfr1*^*Δ/lox*^ cells smaller subcutaneous tumors than uninfected *Fgfr1*^*Δ/lox*^ cells (Fig. 3C). Similar to the primary tumors with the same genotypes, allograft tumors derived from Ad-Cre-infected *Fgfr1*^*Δ/lox*^ cells had significantly less pHH3-positive cells per tumor area than those from uninfected cells (Fig. 3D; Supplementary Fig. S2). These findings suggest that acute deletion of FGFR1 in tumor cells suppressed continued growth of tumor cells through increased cell death and decreased proliferation.

Additionally, to determine whether this impact is driven specifically by FGFR1, we used similar methods to examine the impact of deleting *Fgfr2* in the *Rb1*/*Trp53*/*Rbl2*-GEMM. FGFR2 is expressed in the lung epithelium, and the growth and survival of numerous tumor types appear to depend on *FGFR2* amplification or deregulation (27-29). However, loss of *Fgfr2* did not result in decreased tumor burden (Fig. 2C). Taken together, these findings suggest that *Rbl2* loss-driven tumor development depends specifically on FGFR1 signaling.

### Loss of p130 induces FGFR1 and the receptor dependency in SCLC

To our knowledge, functional relationships between p130 and FGFR1 have never been documented in cancer. However, p130 was found to repress *Fgfr1* expression during muscle cell differentiation by co-binding with E2F4 at the *Fgfr1* promoter (30). We thus surmised that *Rbl2* loss may amplify FGFR1 dependency in SCLC by increasing FGFR1 expression levels. We first determined whether p130 loss induces FGFR1 expression in preSC. We found that *Rbl2*-*KO* preSC generated using CRIPSR showed a near-complete loss of p130 but did not induce FGFR1 expression (Fig. 4A). Notably, however, subcutaneous tumors derived from *Rbl2*-*KO* preSC showed a significant induction of FGFR1 compared to those derived from *Rbl2-WT* preSC (Fig. 4B).

**Fig. 4.**
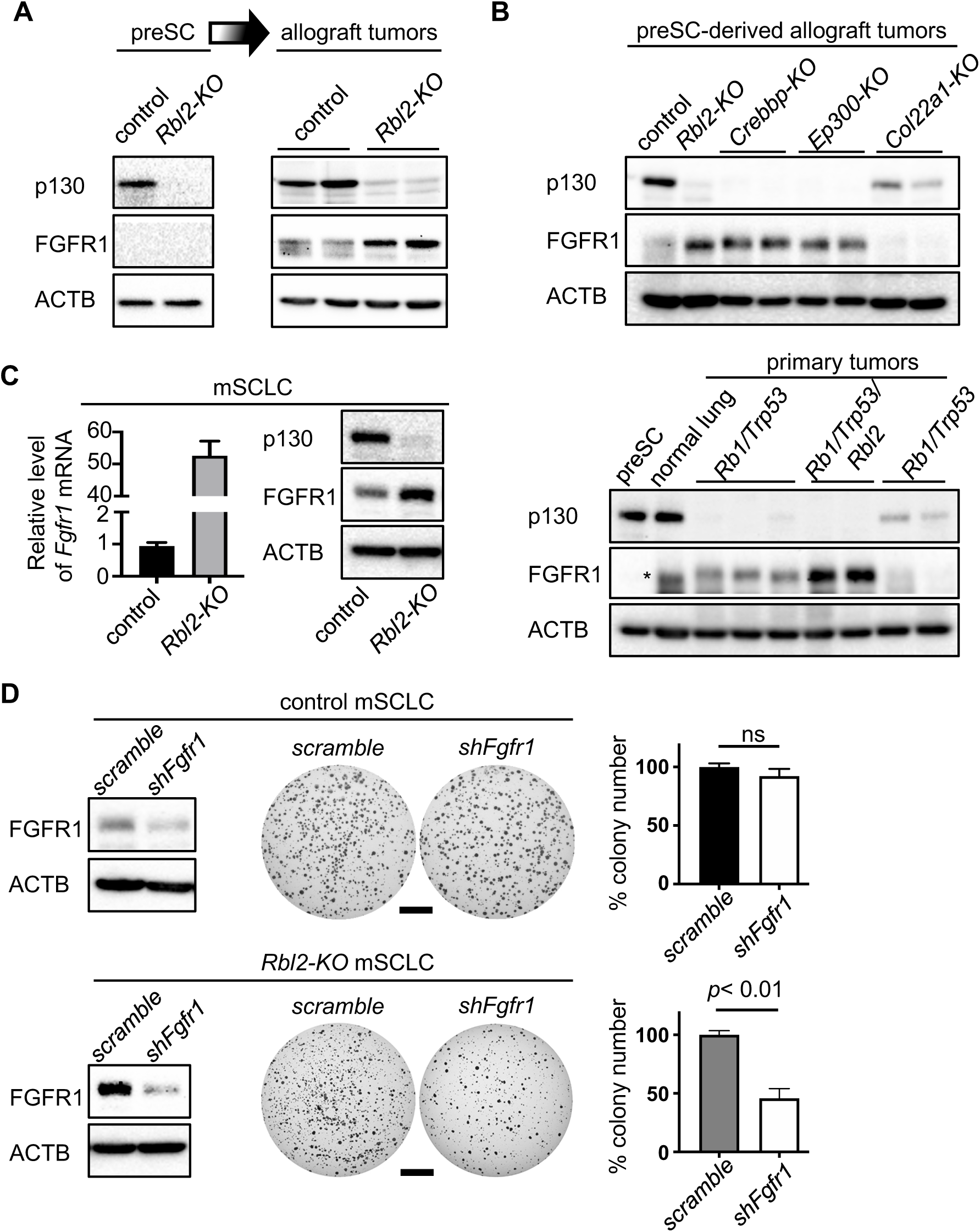
p130 loss induces FGFR1 expression and dependency. **(A)** Immunoblots for FGFR1 and p130 in control and *Rbl2*-*KO* preSC (left) and in preSC-derived subcutaneous tumors (right). (**B)** Immunoblots for FGFR1 and p130 in the subcutaneous tumors generated from control and various targeted preSC (top) and in primary lung tumors developed in *Rb1/Trp53*-mutant and *Rb1/Trp53/Rbl2-*mutant GEMMs (bottom). Asterisk indicates a non-specific band. (**C**) Left, RT-qPCR for *Fgfr1* mRNA normalized to *ARBPP0*. Right, immunoblots for FGFR1 and p130 in control and *Rbl2*-*KO* preSC. (**D**) Left, immunoblots for FGFR1 in control and *Rbl2*-*KO* preSC after shRNA-mediated knockdown of *Fgfr1*. Middle, representative images of soft agar colonies formed by control and *Fgfr1*-knockdown cells. Right, quantification of colonies >0.2mm in diameter (n=3 per cell type). ACTB blot verifies equal loading of total proteins. Statistical tests performed using unpaired t-test (ns: not significant). Error bar represents standard deviation. Scale bar: 5mm.

To determine whether this inverse correlation between p130 and FGFR1 levels exists across molecular subtypes of SCLC cells, we examined the subcutaneous tumors derived from variants of mutant preSC in which *Crebbp, Ep300*, or *Col22a1* was mutated using CRISPR. Loss-of-function mutations in these genes are among the most frequent alterations found in the SCLC genome and have been shown to promote malignant progression of preSC (3, 31). Similar to *Rbl2*-*KO* tumors, *Crebbp-KO* or *Ep300-KO* tumors expressed FGFR1 but lacked p130, whereas *Col22a1-KO* tumor showed the opposite expression patterns (Fig. 4B). Additionally, a survey of multiple primary tumors developed in *Rb1*/*Trp53*-GEMM and *Rb1*/*Trp53/Rbl2*-GEMM showed a similar inverse correlation between p130 and FGFR1 expression (Fig. 4B).

We also tested whether p130 regulates FGFR1 expression in malignant SCLC cells. CRISPR-mediated knockout of *Rbl2* increased *Fgfr1* mRNA and FGFR1 protein levels in mSCLC (Fig. 4C). Notably, subsequent shRNA-mediated knockdown of *Fgfr1* in *Rbl2-*KO mSCLC cells resulted in these cells giving rise to fewer colonies in soft agar than *Rbl2-*KO cells treated with a scrambled shRNA control, whereas *Fgfr1* knockdown in *Rbl2*-*WT* cells did not impact colony-forming ability (Fig. 4D). Taken together, these findings suggest that loss of p130 induces FGFR1 expression and results in the dependency of tumor development on FGFR1 signaling.

### FGFR1-dependent SCLC cells require activation of PLCG1

To begin to identify intracellular mediators of FGFR1-driven SCLC development, we examined the phosphorylation status of ERK1/2, AKT1, STAT1, and PLCG1, major transducers downstream of FGFR1 during organ development (12). Phosphorylation of AKT1 and PLCG1 increased in *Fgfr1*-preSC relative to control preSC while that of ERK1/2 and STAT did not change, suggesting lack of activation of ERK1/2 and STAT by FGFR1 (Fig. 5A). This finding was intriguing as FGFR1 signaling in SCLC has so far been linked to MEK-ERK pathway (32, 33). We performed similar analysis on human SCLC lines with varying levels of FGFR1, including H82, H209, and H524. Lentiviral shRNA-mediated knockdown of FGFR1 significantly reduced growth of H82 and H209 cells in soft agar and reduced phosphorylation of PLCG1, but did not impact the growth of H524 colonies and PLCG1 phosphorylation despite comparable levels of knockdown (Fig. 5B, C). In all cell lines, phosphorylation levels of ERK1/2, AKT1, and STAT1 did not change regardless of FGFR1 expression, further demonstrating a specific functional relationship between FGFR1 and PLCG1 in SCLC (Fig. 5C).

**Fig. 5.**
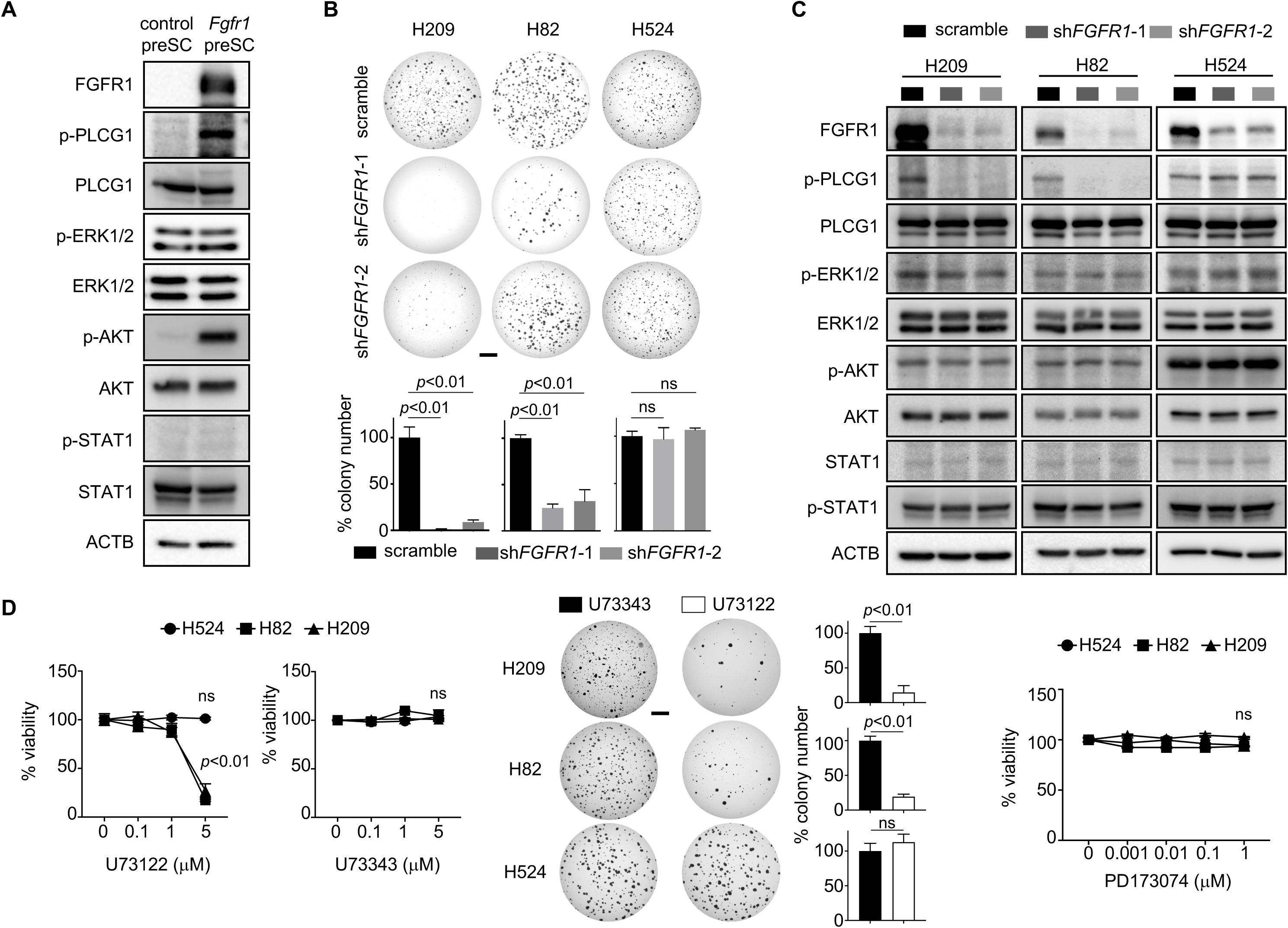
FGFR1-dependent SCLC cells require activation of PLCG1. **(A)** Immunoblot for phosphorylated forms and respective total protein in control and *Fgfr1*-preSC. ACTB blot verifies equal loading of total proteins. (**B)** Representative images of soft agar colonies formed from human SCLC lines with FGFR1 knockdown (top) and quantification of colonies >0.2mm in diameter (bottom) (n=3 replicates per cell type). (**C)** Immunoblot for phosphorylated and respective total protein in human SCLC lines with FGFR1 knockdown. (**D)** Left, viability of human SCLC lines treated with U73122 (PLCG1 inhibitor) and U73343 (innocuous analog of U73122) once for 4 days, as measured by MTT assay. Middle, representative images of soft agar culture of human SCLC lines treated with 1μM U73122 and U73343 every three days for 14 days (H82 and H524) or 21 days (H209) and quantification of colonies (>0.2mm) in diameter (n=3 replicates per cell line). Right, viability of human SCLC lines treated with PD173074 (pan-FGFR inhibitor) once for 4 days, as measured by MTT assay. Statistical tests performed using unpaired t-test (ns: not significant). Error bar represents standard deviation. Scale bars: 5mm.

To determine the significance of PLCG1 activation in SCLC cells, we tested effects of U73122, a chemical inhibitor of PLCG1, and its structural analog U73344 as an innocuous control (34). Compared with control, U73122 reduced viability of H82 and H209 cells in a dose-dependent manner (Fig. 5D) and suppressed their ability to give rise to colonies in soft agar (Fig. 5D). However, these effects of U73122 were not seen for H524 (Fig. 5D), consistent with the observation that this cell line does not respond to FGFR1 for growth. Importantly, treatment of a pan-FGFR inhibitor PD173074 did not affect viability in the tested SCLC lines (Fig. 5D). Taken together, these data suggest that PLCG1 mediates FGFR1 signaling to promote tumorigenic progression of preSC and continued growth of SCLC cells independently of ERK1/2, AKT1, and STAT1.

To further explore the mechanism by which FGFR1 drives tumor growth, we performed RNA sequencing and analyzed a total of 1823 genes that were differentially expressed (DE) in *Fgfr1*-preSC (Fig. 6A; Supplementary Data 1). Gene ontology (GO) analysis of DE genes indicated enrichment of GO terms for cell cycle/mitosis and neuron differentiation and development (Fig. 6B; Supplementary Data 2). The GO terms related to neuron differentiation were further enriched in top 228 DE genes (with 1.5-fold or higher changes in expression) (Fig. 6C; Supplementary Data 3). Enrichment of these GO terms reflected the upregulation of *Rhoa, Gli3, Etv4, Arx*, and *Lif* and downregulation of *Nefl, Chl1, Dcx, Dcc, Slitrk3, Nr2e1*, and *Bdnf* (Fig. 6A). These results suggest that, in addition to promoting cell cycle, FGFR1 signaling regulates neural differentiation in cells progressing from precancerous neuroendocrine cells to neuroendocrine tumor. This observation is consistent with established roles of FGFR1 in proliferation of neural progenitor cells and neural development (35), and with recent studies showing that dysregulation in neural differentiation is a major event associated with SCLC progression (36, 37).

**Fig. 6.**
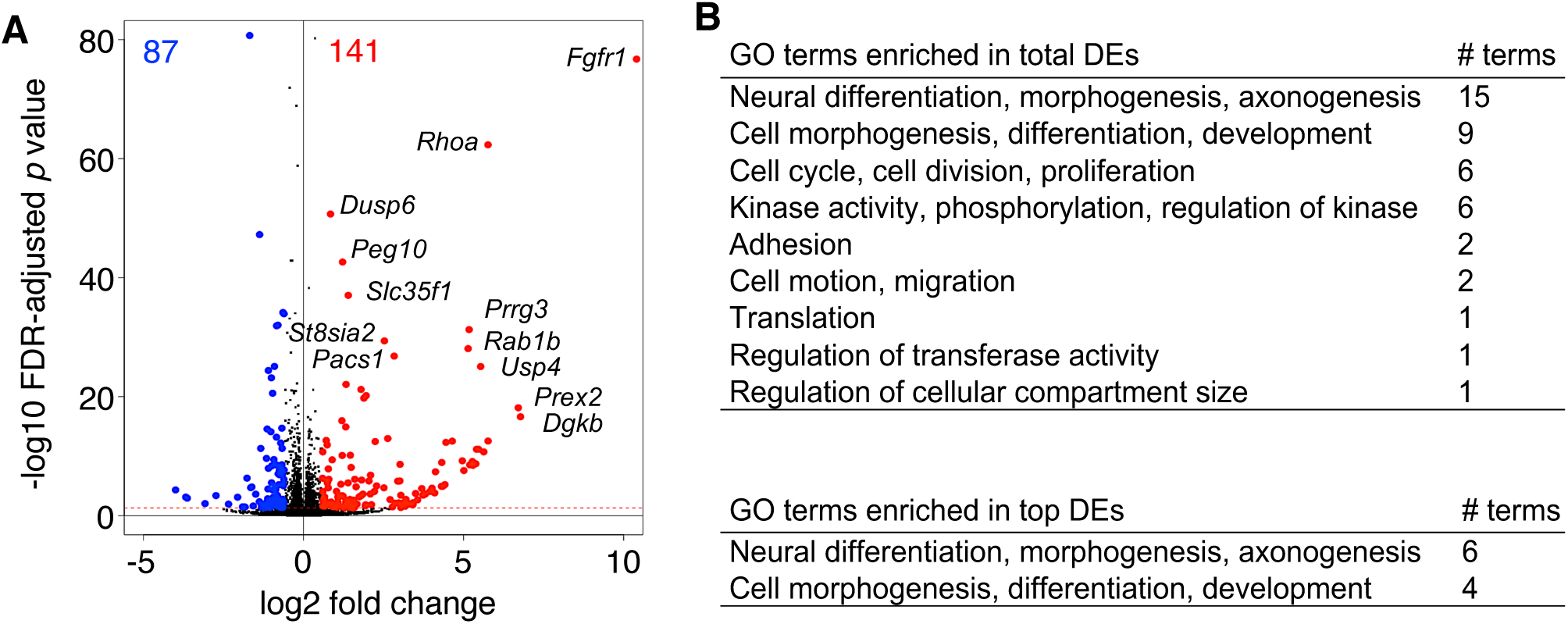
FGFR1 induces transcriptome-wide change in the expression of genes involved in proliferation and differentiation. **(A)** Volcano plot showing differential gene expression between control and *Fgfr1*-preSC; changes in gene expression are significant above the red horizontal line, with an FDR-adjusted *p*-value < 0.05, and red and blue dots indicate genes with a an 1.5-fold or higher increase or decrease in expression. FDR: false discovery rate. **(B)**, Gene ontology (GO) terms enriched in genes differentially expressed (DE) in *Fgfr1*-preSC relative to control preSC (FDR-adjusted *p*-value < 0.05). Bottom, summary of GO terms enriched in top DE genes with an 1.5-fold or higher increase or decrease in expression.

## Discussion

Here we report for the first time a direct role for nonamplified FGFR1 in SCLC development. *Fgfr1* knockout significantly inhibited tumor growth in various SCLC models, notably in human SCLC lines that have previously shown little sensitivity to existing pharmacological inhibitors (8). While previous studies utilizing FGFR1 inhibition in human cell lines have focused on *FGFR1* amplification as a biomarker for drug sensitivity, our findings suggest that FGFR1 expression may confer sensitivity as well, including in subtypes that lack *FGFR1* amplification.

Little is known about regulators of FGFR1 expression in cancer, and further study of these regulators would provide insight into the heterogeneity in FGFR1 dependency among SCLC patient tumors. Our study shows that p130 loss, found in a subset of SCLC patient tumors, induces FGFR1 expression and is sufficient to render p130*-*deficient mouse tumor dependent on the receptor. The role of p130 loss in FGFR1 induction is permissive as it appears to require additional unknown factor present in transforming or malignant cells, which may not be present in preSC. During myogenesis, *Fgfr1* expression is induced in myoblast by complexes containing the Sp1 transcription factor, RB family member p107, and E2F4, and is inhibited in myogenic cells by p130-E2F4 complexes (30). While Sp1, RB family member p107, and E2F4 are variably expressed in SCLC (11, 38, 39), their involvement in the regulation of FGFR1 is not known, and it will be interesting to assess whether these factors also impact FGFR1 dependency.

Despite the importance of FGFR1 as a prognostic and therapeutic biomarker, the intracellular pathways downstream of FGFR1 have been poorly defined. Our findings indicate that PLCG1 is a critical mediator of FGFR1 signaling in SCLC whereas other pathways, including the ERK pathway, do not appear consistently engaged. PLCG1 is a member of the phospholipase C (PLC) family of enzymes. When activated by growth factor receptors, PLCG1 hydrolyzes phosphatidylinositol 4,5-bisphsphate (PIP2) to generate inositol 1,4,5-triphosphate (IP3) and diacylglycerol (DAG). IP3 then trigger Ca^2+^ release and signaling while DAG activates protein kinase C (PKC), giving rise to various intracellular signaling cascades that control proliferation, differentiation, morphology, and migration in multiple cell types (40). The functional impact of the interaction between FGFR1 and PLCG1 has remained poorly understood. Almost two decades ago, the first reports establishing their interaction suggested that it is not critical for mitogenesis (41, 42). More recently, PLCG1 has been shown to mediate FGFR1 signaling in maintaining the differentiation capacity of adult neural stem cells (43). Our data show that PLCG1 activation cooccurs with transcriptomic change related to neural differentiation during FGFR1-driven tumorigenesis, and thus suggest a role for PLCG1 in regulating neuroendocrine differentiation in SCLC. Our findings help add SCLC to the growing list of malignancies in which the activity of PLCG1 is implicated and indicate PLCG1 as a potential therapeutic target in SCLC (44-48).

Our findings are in line with those of previous studies in which the activation of ERK1/2 and AKT varied among SCLC primary tumors and did not strongly correlate with disease-free or overall survival (49, 50). The role of the ERK pathway in FGFR signaling in SCLC remains unclear. A hyperactive RAF-MEK-ERK pathway has been shown to be tumor-suppressive in SCLC cell lines, in contrast to the oncogenic role of this pathway in non-small cell lung cancer cells (10, 11). Further, in a recent study in *Rb1/Trp53-GEMM*, constitutive FGFR1 activation suppressed the development of SCLC originating from CGRP-positive neuroendocrine cells while promoting the tumor development from keratin 14 (K14)-positive airway epithelial cells (13). These studies suggest that whether FGFR1 and the ERK pathway are oncogenic in SCLC may depend on cell of origin. Importantly, our findings demonstrate an oncogenic role for FGFR1 in SCLC tumors with neuroendocrine origin. We surmised that differences in FGFR1 dependency may be due to on the extent of signaling from constitutively active versus wild-type of FGFR1 and the interactions between FGFR1 and its signal transducers.

Taken together, our identification of two novel players in in FGFR1 signaling, PLCG1 and p130, shed light on the mechanism of FGFR1-driven SCLC development and homeostasis, and may facilitate the development of biomarkers for SCLC subtypes appropriate for FGFR1-targeted therapies, as well as targeted therapeutic strategies for SCLC.

## Acknowledgements

We thank Drs. Anton Berns, Tyler Jacks, Julien Sage, Juha Partanen, and David Ornitz for sharing *Trp53* ^*lox*^, *Rb1*^*lox*^, *Rbl2*^*lox*^, *Fgfr1*^*lox*^ and *Fgfr2*^*lox*^ mice, respectively. We thank J. Hsu, J. Sage, and C. Dunn for comments on the manuscript. This work was supported by National Institutes of Health (NCI R01CA194461; U01CA224293; R03CA215777) and American Cancer Society (RSG-15-066-01-TBG) to K-S. P. We also thank S. Vanhoose and her team at the Research Histology Core and the experts of Genome Analysis and Technology Core at the University of Virginia Cancer Center and Biostatistics and Bioinformatics Shared Resources at the H. Lee Moffitt Cancer Center & Research Institute, which are supported by the National Cancer Institute grants (P30CA044579 and P30CA076292).

## References

1. Byers LA, Rudin CM. Small cell lung cancer: where do we go from here? Cancer 2015;121:664–72

2. Peifer M, Fernandez-Cuesta L, Sos ML, George J, Seidel D, Kasper LH, et al. Integrative genome analyses identify key somatic driver mutations of small-cell lung cancer. Nat Genet 2012;44:1104–10

3. George J, Lim JS, Jang SJ, Cun Y, Ozretic L, Kong G, et al. Comprehensive genomic profiles of small cell lung cancer. Nature 2015;524:47–53

4. Voortman J, Lee JH, Killian JK, Suuriniemi M, Wang Y, Lucchi M, et al. Array comparative genomic hybridization-based characterization of genetic alterations in pulmonary neuroendocrine tumors. Proc Natl Acad Sci U S A 2010;107:13040–5

5. Zhang L, Yu H, Badzio A, Boyle TA, Schildhaus HU, Lu X, et al. Fibroblast Growth Factor Receptor 1 and Related Ligands in Small-Cell Lung Cancer. J Thorac Oncol 2015; 10:1083–90.

6. Desai A, Adjei AA. FGFR Signaling as a Target for Lung Cancer Therapy. J Thorac Oncol 2016;11:9–20

7. Pardo OE, Latigo J, Jeffery RE, Nye E, Poulsom R, Spencer-Dene B, et al. The fibroblast growth factor receptor inhibitor PD173074 blocks small cell lung cancer growth in vitro and in vivo. Cancer Res 2009;69:8645–51

8. Sos ML, Dietlein F, Peifer M, Schottle J, Balke-Want H, Muller C, et al. A framework for identification of actionable cancer genome dependencies in small cell lung cancer. Proc Natl Acad Sci U S A 2012;109:17034–9

9. Thomas A, Lee JH, Abdullaev Z, Park KS, Pineda M, Saidkhodjaeva L, et al. Characterization of fibroblast growth factor receptor 1 in small-cell lung cancer. J Thorac Oncol 2014;9:567–71

10. Ravi RK, Weber E, McMahon M, Williams JR, Baylin S, Mal A, et al. Activated Raf-1 causes growth arrest in human small cell lung cancer cells. J Clin Invest 1998;101:153–9

11. Ravi RK, Thiagalingam A, Weber E, McMahon M, Nelkin BD, Mabry M. Raf-1 causes growth suppression and alteration of neuroendocrine markers in DMS53 human small-cell lung cancer cells. Am J Respir Cell Mol Biol 1999;20:543–9

12. Ornitz DM, Itoh N. The Fibroblast Growth Factor signaling pathway. Wiley Interdiscip Rev Dev Biol 2015;4:215–66

13. Ferone G, Song JY, Krijgsman O, van der Vliet J, Cozijnsen M, Semenova EA, et al. FGFR1 Oncogenic Activation Reveals an Alternative Cell of Origin of SCLC in Rb1/p53 Mice. Cell Rep 2020;30:3837–50 e3

14. Meuwissen R, Linn SC, Linnoila RI, Zevenhoven J, Mooi WJ, Berns A. Induction of small cell lung cancer by somatic inactivation of both Trp53 and Rb1 in a conditional mouse model. Cancer Cell 2003;4:181–9

15. Gazdar AF, Savage TK, Johnson JE, Berns A, Sage J, Linnila RI, et al. The comparative pathology of genetically engineed mouse models for neuroendocrine carcinomas of the lung. J Thorac Oncol 2015;10:553–64

16. Kim DW, Wu N, Kim YC, Cheng PF, Basom R, Kim D, et al. Genetic requirement for Mycl and efficacy of RNA Pol I inhibition in mouse models of small cell lung cancer. Genes Dev 2016;30:1289–99

17. Trokovic R, Trokovic N, Hernesniemi S, Pirvola U, Vogt Weisenhorn DM, Rossant J, et al. FGFR1 is independently required in both developing mid- and hindbrain for sustained response to isthmic signals. EMBO J 2003;22:1811–23

18. Yu K, Xu J, Liu Z, Sosic D, Shao J, Olson EN, et al. Conditional inactivation of FGF receptor 2 reveals an essential role for FGF signaling in the regulation of osteoblast function and bone growth. Development 2003;130:3063–74

19. DuPage M, Dooley AL, Jacks T. Conditional mouse lung cancer models using adenoviral or lentiviral delivery of Cre recombinase. Nat Protoc 2009;4:1064–72

20. Trapnell C, Pachter L, Salzberg SL. TopHat: discovering splice junctions with RNA-Seq. Bioinformatics 2009;25:1105–1111.

21. Anders S, Pyl PT, Huber W. HTSeq--a Python framework to work with high-throughput sequencing data. Bioinformatics 2015;31:166–169.

22. Love MI, Huber W, Anders S. Moderated estimation of fold change and dispersion for RNA-seq data with DESeq2. Genome Biol 2014;15:550.

23. McFadden DG, Papagiannakopoulos T, Taylor-Weiner A, Stewart C, Carter SL, Cibulskis K, et al. Genetic and clonal dissection of murine small cell lung carcinoma progression by genome sequencing. Cell 2014;156:1298–311

24. Schaffer BE, Park KS, Yiu G, Conklin JF, Lin C, Burkhart DL, et al. Loss of p130 accelerates tumor development in a mouse model for human small-cell lung carcinoma. Cancer Res 2010;70:3877–83

25. Park KS, Martelotto LG, Peifer M, Sos ML, Karnezis AN, Mahjoub MR, et al. A crucial requirement for Hedgehog signaling in small cell lung cancer. Nat Med 2011;17:1504–8

26. Sen T, Tong P, Stewart CA, Cristea S, Valliani A, Shames DS, et al. CHK1 Inhibition in Small-Cell Lung Cancer Produces Single-Agent Activity in Biomarker-Defined Disease Subsets and Combination Activity with Cisplatin or Olaparib. Cancer Res 2017;77:3870–84

27. Arman E, Haffner-Krausz R, Gorivodsky M, Lonai P. Fgfr2 is required for limb outgrowth and lung-branching morphogenesis. Proc Natl Acad Sci U S A 1999;96:11895–9

28. Yuan T, Klinkhammer K, Liu H, Gao S, Yuan J, Hopkins S, et al. Temporale Expression of Fgfr1 and 2 During Lung Development, Homeostasis, and Regeneration. Front Pharmacol 2020;11:120

29. Pearson A, Smyth E, Babina IS, Herrera-Abreu MT, Tarazona N, Peckitt C, et al. High-Level Clonal FGFR Amplification and Response to FGFR Inhibition in a Translational Clinical Trial. Cancer Discov 2016;6:838–51

30. Parakati R, DiMario JX. Dynamic transcriptional regulatory complexes, including E2F4, p107, p130, and Sp1, control fibroblast growth factor receptor 1 gene expression during myogenesis. J Biol Chem 2005;280:21284–94

31. Jia D, Augert A, Kim DW, Eastwood E, Wu N, Ibrahim AH, et al. Crebbp Loss Drives Small Cell Lung Cancer and Increases Sensitivity to HDAC Inhibition. Cancer Discovery 2018;8:1422–1437.

32. Pardo OE, Acaro A, Salerno G, Tetley TD, Valovka T, Gout I, et al. Novel cross talk between MEK and S6K2 in FGF-2 induced proliferation of SCLC cells. Oncogene 2001; 20:7658–67

33. Wang K, Ji W, Yu Y, Li Z, Niu X, Xia W, et al. FGFR1-ERK1/2-SOX2 axis promotes cell proliferation, epithelial-mesenchymal transition, and metastasis in FGFR1-amplified lung cancer. Oncogene 2018;37:5340–54

34. Bleasdale JE, Bundy GL, Bunting S, Fitzpatrick FA, Huff RM, Sun FF, et al. Inhibition of phospholipase C dependent processes by U-73, 122. Adv Prostaglandin Thromboxane Leukot Res 1989;19:590–3

35. Ohkubo Y, Uchida AO, Shin D, Partanen J, Vaccarino FM. Fibroblast growth factor receptor 1 is required for the proliferation of hippocampal progenitor cells and for hippocampal growth in mouse. J Neurosci 2004;24:6057–69

36. Denny SK, Yang D, Chuang CH, Brady JJ, Lim JS, Gruner BM et al. Nfib Promotes Metastasis through a Widespread Increase in Chromatin Accessibility. Cell 2016;66:328–342.

37. Yang D, Qu F, Cai H, Chuang CH, Lim JS, Jahchan N et al. Axon-like protrusions promote small cell lung cancer migration and metastasis. Elife 2019;8.

38. Kim WY, Jang JY, Jeon YK, Chung DH, Kim YG, Kim CW. Syntenin increases the invasiveness of small cell lung cancer cells by activating p38, AKT, focal adhesion kinase and SP1. Exp Mol Med 2014;46:e90.

39. Brown KC, Witte TR, Hardman WE, Luo H, Chen YC, Carpenter AB, et al. Capsaicin displays anti-proliferative activity against human small cell lung cancer in cell culture and nude mice models via E2F pathway. PLoS One 2010;5:e10243.

40. Koss H, Bunney TD, Behjati S, Katan M. Dysfunction of phospholipase Cg in immune disorders and cancer. Trends Biochem Sci 2014;39:603–11.

41. Peters KG, Marie J, Wilson E, Ives HE, Esobedo J, Del Rosario M, et al. Point mutation of an FGF recentor abolishes phosphatidylinositol turnover and Ca^2+^flux but nor mitogenesis. Nature 1992;358:678–681

42. Mohammadi M, Dionne CA, Li W, Li N, Spival T, Honegger M, et al. Point mutation in FGF receptor eliminates phosphatidylinositol hydrolysis without affecting mitogenesis. Nature 1992;358:681–684.

43. Vaque JP, Gomex-Lopez G, Monsalvez V, Valera I, Martinez N, Perez C, et al. PLCG1 mutations in cutaneous T-cell lymphomas. Blood 2014;123:2034–43.

44. Kunze K, Spieker T, Gamerdinger U, Nau K, Berger J, Dreyer T, et al. Arecurrent activating PLCG1 mutation in cardiac angiosarcomas increases apoptosis resistance and invasiveness of endothelial cells. Cancer Res 2014;74:6173–83.

45. Gouaze-Andersson V, Delmas C, Taurance M, Martinex-Gala J, Evrard S, Mazoyer S, et al. FGFR1 Induces Glioblastoma Resistance through the PLC*γ*1/Hif1*α* Pathway. Cancer Res 2016;76:3036–44.

46. Walker K, Boyd NH, Anderson JC, Willey CD, Hjelmeland AB. Kinome profiling of glioblastoma cells revelas PLCG1 as a target in restricted glucose. Biomarker Res 2018;6:22.

47. Tang W, Zhou Y, Sun D, Dong L, Xia J, Yang B. Oncogenic role of phospholipase C-*γ*1 in progression of hepatocellular carcinoma. Hepatol Res 2019;49:559–569.

48. Ma DK, Ponnusamy K, Song MR, Ming GI, Song H. Molecualr genetic analysis of FGFR1 signalling reveals distinct roles of MAPK and PLAgamma1 activation for self-renewal of adult neural stem cells. Mol Brain 2009;2:16.

49. Blackhall FH, Pintillie M, Michael M, Leighl N, Feld R, Tsao MS, et al. Expression and prognostic significance of kit, protein kinase B, and mitogen-activated protein kinase in patients with small cell lung cancer. Clin Cancer Res 2003;9:2241–7.

50. Schmid K, Bago-Horvath Z, Berger W, Haitel A, Cejka D, Werzowa J, et al. Dual inhibition of EGFR and mTOR pathways in small cell lung cancer. Br J Cancer 2010;103:622–8.

