## Supplementary Tables for "FGFR1 is critical for *Rbl2* loss-driven tumor development and requires PLCG1 activation for continued growth of small cell lung cancer": Supplementary Tables .pdf

**Supplementary Table 1.**

| **Plasmids** | **Source** |
|---|---|
| pCW57.1 | Gift from David Root, Addgene #41393 |
| Tet-pLKO-puro | Gift from Dmitri Wiederschain, Addgene #21915 |
| pL-CRISPR.EFS.tRFP | Gift from Benjamin Ebert, Addgene #57819 |
| psPAX2 and pMD2.G | Gift from Didier Trono, Addgene #12259, #12260 |

**Supplementary Table 2.**

Sequences of guide RNAs for mouse genome

| Target | Sequence (5'-3') |
|---|---|
| *Rbl2* exon1 | TCAGATCCAGCAGCGGTTCG |
| *Rbl2* exon3 | GTACGTTCTCGGAAATGTGG |

**Supplementary Table 3.**

Sequences of shRNAs for human and mouse genes

| Target | Sequence (5'-3') |
|---|---|
| *FGFR1* (human) | GAGATGGAGGTGCTTCACTTA |
| *FGFR1* (human) | TCTTGAAGACTGCTGGAGTTA |
| *Fgfr1* (mouse) | CGAGGATAACGTAATGAAGAT |
| *Fgfr1* (mouse) | CCTGGAGCATCATAATGGATT |

**Supplementary Table 4.**

| **Antibodies** | **Source** |
|---|---|

| | |
|---|---|
| FGFR1 | Cell Signaling Technology, 9740 |
| phospho-FGFR1 | Cell Signaling Technology, 52928 |
| ERK1/2 | Cell Signaling Technology, 4695 |
| phospho-ERK1/2 | Cell Signaling Technology, 4376 |
| AKT | Cell Signaling Technology, 9272 |
| phospho-AKT | Cell Signaling Technology, 4060 |
| STAT1 | Cell Signaling Technology, 9172 |
| phopho-STAT1 | Cell Signaling Technology, 7649 |
| PLCG1 | Cell Signaling Technology, 2822 |
| phosphor-PLCG1 | Cell Signaling Technology, 2821 |
| p130 | Santa Cruz Biotechnology, sc-317 |
| GAPDH | Santa Cruz Biotechnology, sc-47724 |
| ACTB | Santa Cruz Biotechnology, sc-47778 |
| Calca (CGRP) | Sigma, C8198 |
| Secondary Ab-488 | Thermo Fisher Scientific, A11034 |
| Secondary Ab-HRP (Rabbit) | Jackson Immuno Research. 111-035-003 |
| Secondary Ab-HRP (mouse) | Jackson Immuno Research, 115-035-003 |
