## Supplementary Fig 1 for "FGFR1 is critical for *Rbl2* loss-driven tumor development and requires PLCG1 activation for continued growth of small cell lung cancer": Supplementaty Fig 1.pdf

**Supplementary Fig. S1**

**Genotyping PCR for tail DNAs**

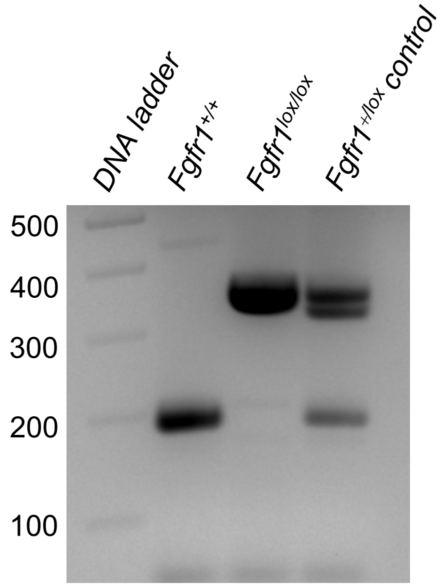


**Immunoblot for primary cells derived from lung tumors**

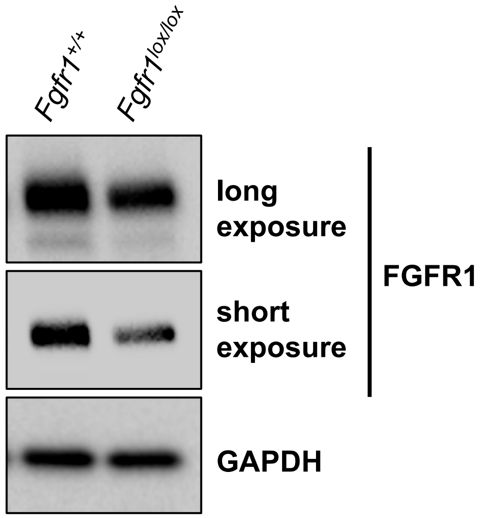


**Supplementary Fig. S1.** *Fgfr1*$^{+/+}$ *Rb1/Trp53/Rbl2*-GEMM and *Fgfr1*$^{lox/lox}$ *Rb1/Trp53/Rbl2*-GEMM that were infected with Ad-Cre and aged for 6 months and primary cells were derived from lung tumors. FGFR1 expression in the primary cells from *Fgfr1*$^{lox/lox}$ *Rb1/Trp53/Rbl2*-GEMM indicate incomplete recombination of *Fgfr1*$^{lox}$ alleles.
