## Supplementary Fig 2 for "FGFR1 is critical for *Rbl2* loss-driven tumor development and requires PLCG1 activation for continued growth of small cell lung cancer": Supplementaty Fig 2.pdf

**Supplementary Fig. S2**

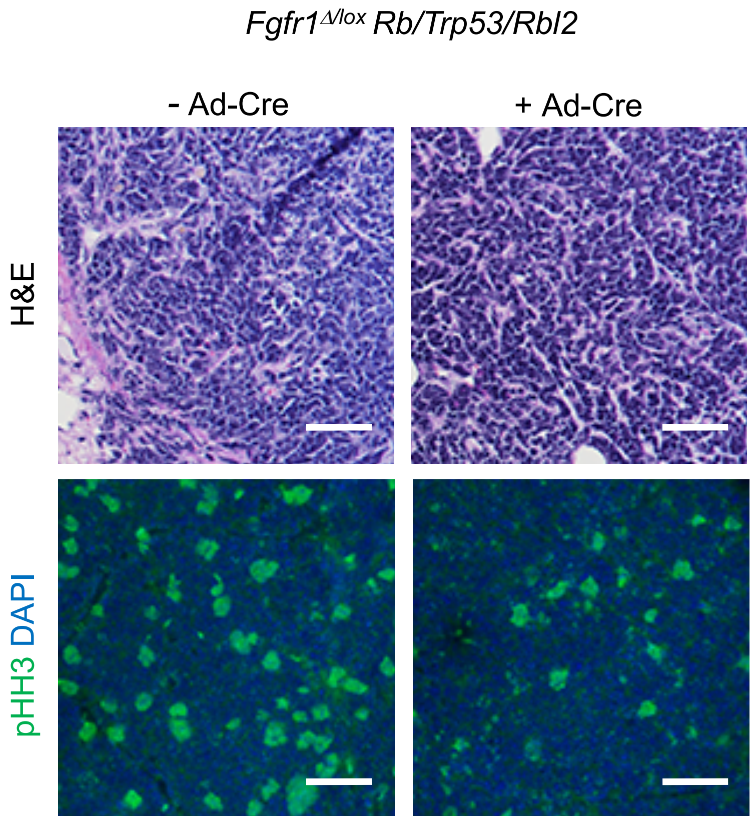


**Supplementary Fig. S2.** H&E staining for section of subcutaneous tumors generated from control or Ad-Cre-infected *Fgfr1*$^{\Delta/lox}$ *Rb1/Trp53/Rbl2* SCLC cells. Histology is typical of SCLC. Scale bars: 50 μm
