## Supplementary Data 1 for "FGFR1 is critical for *Rbl2* loss-driven tumor development and requires PLCG1 activation for continued growth of small cell lung cancer": Supplementaty Data 1.pdf

| gene | m.experimei | m.control_pi | FC.tradition | log2FC.tradit | baseMean | log2FoldCha | lfcSE | stat | pvalue | padj |
|---|---|---|---|---|---|---|---|---|---|---|
| Chl1 | 379.011954 | 994.339555 | 0.38116954 | -1.3914952 | 686.675754 | -1.3852152 | 0.04073412 | -34.006264 | 1.8E-253 | 2.668E-249 |
| Dpp10 | 232.716366 | 574.056145 | 0.40538956 | -1.3026192 | 403.386255 | -1.2963161 | 0.04930678 | -26.290826 | 2.442E-152 | 1.809E-148 |
| Rn45s | 20670.9963 | 16415.2398 | 1.25925643 | 0.3325721 | 18543.1181 | 0.3298368 | 0.01479608 | 22.2921705 | 4.401E-110 | 2.174E-106 |
| Clca3b | 5761.44139 | 4504.90692 | 1.27892573 | 0.35493249 | 5133.17415 | 0.35383662 | 0.01731982 | 20.4295823 | 9.1279E-93 | 3.3821E-89 |
| Scara3 | 66.0691481 | 211.795067 | 0.31194848 | -1.6806203 | 138.932107 | -1.6813761 | 0.08611841 | -19.524004 | 6.8643E-85 | 2.0347E-81 |
| Car8 | 3549.44936 | 2766.86402 | 1.282842 | 0.35934349 | 3158.15669 | 0.35927003 | 0.0184625 | 19.4594499 | 2.4237E-84 | 5.987E-81 |
| Fgfr1 | 1062.17511 | 0.79741706 | 1332.01954 | 10.3793995 | 531.486262 | 10.4039689 | 0.54656737 | 19.0351079 | 8.7312E-81 | 1.8487E-77 |
| Alcam | 1902.77415 | 2558.46184 | 0.74371801 | -0.4271724 | 2230.61799 | -0.4262399 | 0.02312302 | -18.433571 | 7.0661E-76 | 1.3091E-72 |
| Krt8 | 8271.77341 | 9680.76552 | 0.85445447 | -0.2269245 | 8976.26946 | -0.2267366 | 0.01256485 | -18.045299 | 8.5898E-73 | 1.4145E-69 |
| Rhoa | 124.687708 | 2.32739839 | 53.5738568 | 5.74345726 | 63.5075532 | 5.76208565 | 0.33520693 | 17.1896377 | 3.1751E-66 | 4.7058E-63 |
| Nnat | 16773.9187 | 19024.9996 | 0.88167774 | -0.1816767 | 17899.4592 | -0.1827529 | 0.01094117 | -16.703229 | 1.2416E-62 | 1.6729E-59 |
| Dusp6 | 368.036744 | 205.279692 | 1.7928551 | 0.8422589 | 286.658218 | 0.84253764 | 0.05419087 | 15.5475935 | 1.652E-54 | 2.0403E-51 |
| AW551984 | 64.0886484 | 166.07072 | 0.38591179 | -1.373657 | 115.079684 | -1.3721122 | 0.09132753 | -15.02408 | 5.1066E-51 | 5.8219E-48 |
| Scnn1a | 1777.98171 | 2276.06924 | 0.78116328 | -0.356304 | 2027.02548 | -0.355244 | 0.02478973 | -14.330291 | 1.4153E-46 | 1.4216E-43 |
| Clstn2 | 1329.26099 | 1739.7134 | 0.76406895 | -0.3882253 | 1534.4872 | -0.3908801 | 0.02727868 | -14.329145 | 1.4388E-46 | 1.4216E-43 |
| Peg10 | 205.636899 | 89.0359781 | 2.3095933 | 1.20763883 | 147.336438 | 1.21979673 | 0.08536033 | 14.2899714 | 2.5271E-46 | 2.3409E-43 |
| H3f3b | 10299.1693 | 9149.91375 | 1.12560288 | 0.17069793 | 9724.54152 | 0.17045853 | 0.01255956 | 13.5720113 | 5.8689E-42 | 5.1167E-39 |
| Slc35f1 | 128.476347 | 49.2270881 | 2.60987095 | 1.38397847 | 88.8517175 | 1.39793526 | 0.10468435 | 13.3538138 | 1.1254E-40 | 9.2661E-38 |
| Necab1 | 252.663606 | 394.271552 | 0.64083651 | -0.6419717 | 323.467579 | -0.6424787 | 0.05001966 | -12.844524 | 9.2318E-38 | 7.2013E-35 |
| Homer2 | 2500.55577 | 3009.57928 | 0.83086556 | -0.267313 | 2755.06753 | -0.2665646 | 0.02079574 | -12.818232 | 1.2962E-37 | 9.6056E-35 |
| Frem2 | 292.066543 | 446.479534 | 0.65415438 | -0.6122969 | 369.273039 | -0.6096278 | 0.04763353 | -12.798293 | 1.676E-37 | 1.1828E-34 |
| Astn2 | 1485.9335 | 2022.32053 | 0.73476656 | -0.4446421 | 1754.12702 | -0.4335367 | 0.03426736 | -12.651593 | 1.0963E-36 | 7.3856E-34 |
| Dcn | 226.066663 | 400.405402 | 0.56459444 | -0.8247132 | 313.236033 | -0.8000975 | 0.0642843 | -12.446234 | 1.466E-35 | 9.4469E-33 |
| Pcdh8 | 159.077671 | 287.399736 | 0.55350667 | -0.8533274 | 223.238703 | -0.8408363 | 0.06768622 | -12.422562 | 1.9715E-35 | 1.2175E-32 |
| Prrg3 | 53.7425408 | 1.51491534 | 35.4756067 | 5.14875545 | 27.628728 | 5.173638 | 0.42060478 | 12.3004738 | 9.0044E-35 | 5.3382E-32 |
| Rragb | 936.433892 | 1327.42291 | 0.70545256 | -0.503379 | 1131.9284 | -0.5082123 | 0.04165448 | -12.200666 | 3.0829E-34 | 1.7574E-31 |
| St8sia2 | 50.2568155 | 8.58271007 | 5.85558816 | 2.54981409 | 29.4197627 | 2.52718136 | 0.21171841 | 11.9365216 | 7.635E-33 | 4.191E-30 |
| Rab1b | 48.2443367 | 1.40465493 | 34.3460416 | 5.10207193 | 24.8244957 | 5.14247516 | 0.44022354 | 11.6815087 | 1.5846E-31 | 8.3875E-29 |
| Insrr | 747.917154 | 985.994616 | 0.75854081 | -0.3987013 | 866.955885 | -0.3976081 | 0.03441002 | -11.555008 | 6.9641E-31 | 3.5591E-28 |
| Pacs1 | 49.7387453 | 6.46466501 | 7.69394009 | 2.9437226 | 28.101705 | 2.8336655 | 0.24800666 | 11.4257637 | 3.1091E-30 | 1.536E-27 |
| Dclk1 | 88.680428 | 166.731054 | 0.53187709 | -0.9108352 | 127.705741 | -0.9076983 | 0.08198146 | -11.071995 | 1.7154E-28 | 8.2012E-26 |
| Usp4 | 47.7366369 | 1.05154673 | 45.3965911 | 5.50451206 | 24.3940917 | 5.53197973 | 0.49994357 | 11.0652082 | 1.8503E-28 | 8.5699E-26 |
| Kcnip4 | 59.2051984 | 126.392925 | 0.46842177 | -1.09412 | 92.7990618 | -1.0964445 | 0.10036019 | -10.925094 | 8.745E-28 | 3.9275E-25 |
| Col6a5 | 81.7890795 | 165.938631 | 0.49288752 | -1.0206697 | 123.863855 | -1.0046122 | 0.09423526 | -10.660683 | 1.5545E-26 | 6.7763E-24 |
| Ptpru | 2666.66762 | 3065.39123 | 0.86992733 | -0.2010332 | 2866.02943 | -0.2003721 | 0.01915977 | -10.45796 | 1.3473E-25 | 5.705E-23 |
| Dag1 | 112.244792 | 42.8039138 | 2.62230208 | 1.39083389 | 77.524353 | 1.32844685 | 0.12754975 | 10.4151273 | 2.1151E-25 | 8.7079E-23 |
| Smarca2 | 56.9459723 | 16.2646092 | 3.50121983 | 1.80785764 | 36.6052907 | 1.79587213 | 0.17562211 | 10.225775 | 1.5201E-24 | 6.0889E-22 |
| Rps3a1 | 2756.85747 | 2236.98574 | 1.23239832 | 0.30146862 | 2496.9216 | 0.30264276 | 0.02965259 | 10.2062855 | 1.8584E-24 | 7.2484E-22 |
| Notch2 | 235.416418 | 347.302038 | 0.67784347 | -0.5609759 | 291.359228 | -0.5617222 | 0.05505898 | -10.202189 | 1.9385E-24 | 7.3669E-22 |
| Mageh1 | 664.008124 | 858.857578 | 0.77312949 | -0.371218 | 761.432851 | -0.3743112 | 0.03675603 | -10.183669 | 2.3455E-24 | 8.6906E-22 |
| Vwf | 2008.5787 | 2365.46389 | 0.84912677 | -0.2359481 | 2187.02129 | -0.232198 | 0.02284514 | -10.163998 | 2.8706E-24 | 1.0377E-21 |
| Por | 1034.54611 | 1270.20691 | 0.81447054 | -0.2960656 | 1152.37651 | -0.2947354 | 0.02903396 | -10.151403 | 3.2663E-24 | 1.1526E-21 |
| Slc17a6 | 64.4288323 | 125.441412 | 0.51361693 | -0.9612353 | 94.9351219 | -0.9641177 | 0.09576554 | -10.06748 | 7.6923E-24 | 2.6514E-21 |
| Chml | 1164.67983 | 1436.90599 | 0.81054699 | -0.3030323 | 1300.79291 | -0.3018848 | 0.03001748 | -10.056968 | 8.5594E-24 | 2.8831E-21 |
| Mmp15 | 41.827245 | 10.637473 | 3.93206592 | 1.97528751 | 26.2323589 | 1.95747703 | 0.19626822 | 9.97347945 | 1.9913E-23 | 6.5584E-21 |
| Med12l | 43.6615354 | 11.5952112 | 3.76547996 | 1.91283377 | 27.6283732 | 1.88562364 | 0.19102292 | 9.87119078 | 5.5502E-23 | 1.7882E-20 |
| Cd109 | 422.906041 | 564.830143 | 0.74873136 | -0.4174799 | 493.868092 | -0.4156191 | 0.042415 | -9.7988698 | 1.1385E-22 | 3.5902E-20 |
| Inhbb | 397.440415 | 581.59052 | 0.68336811 | -0.5492652 | 489.515467 | -0.5485294 | 0.0567194 | -9.6709302 | 4.0072E-22 | 1.2373E-19 |
| Dner | 3550.11327 | 3965.19912 | 0.89531778 | -0.1595283 | 3757.65619 | -0.159508 | 0.01664755 | -9.5814714 | 9.5672E-22 | 2.8938E-19 |
| Atf5 | 1866.29807 | 2213.56042 | 0.84312046 | -0.2461893 | 2039.92924 | -0.2500108 | 0.02631085 | -9.5021944 | 2.0551E-21 | 6.0918E-19 |
| Prex2 | 40.5429233 | 0.34527906 | 117.420743 | 6.87554348 | 20.4441011 | 6.70934664 | 0.7076392 | 9.48131005 | 2.5111E-21 | 7.2975E-19 |
| Ddx3y | 595.29542 | 461.860913 | 1.28890626 | 0.36614734 | 528.578166 | 0.36604306 | 0.03925773 | 9.32410141 | 1.1193E-20 | 3.1902E-18 |
| Map2 | 3005.98145 | 3376.83959 | 0.89017597 | -0.1678375 | 3191.41052 | -0.1675133 | 0.01816188 | -9.223349 | 2.8796E-20 | 8.0526E-18 |
| Dgkb | 24.7737261 | 0.11719444 | 211.389952 | 7.72376299 | 12.4454602 | 6.77721903 | 0.74437141 | 9.10462024 | 8.6569E-20 | 2.376E-17 |
| Rab3c | 4546.11685 | 4117.21952 | 1.1041716 | 0.1429644 | 4331.66818 | 0.1428983 | 0.01571102 | 9.09541941 | 9.4222E-20 | 2.539E-17 |
| Rpl23a | 2115.72948 | 1820.02003 | 1.16247593 | 0.21720085 | 1967.87476 | 0.21746981 | 0.02417605 | 8.99525893 | 2.3568E-19 | 6.2374E-17 |
| Mdga1 | 282.593078 | 387.087632 | 0.73004936 | -0.4539341 | 334.840355 | -0.4538637 | 0.05058796 | -8.971772 | 2.9178E-19 | 7.5868E-17 |
| Sdk1 | 166.050482 | 246.102472 | 0.6747209 | -0.5676372 | 206.076477 | -0.5713363 | 0.06393847 | -8.9357207 | 4.0453E-19 | 1.0337E-16 |
| Gria4 | 65.713468 | 28.7250717 | 2.28766941 | 1.19387859 | 47.2192698 | 1.19935969 | 0.13434343 | 8.92756459 | 4.3549E-19 | 1.094E-16 |
| Kif1a | 4177.75801 | 4639.43997 | 0.90048757 | -0.1512217 | 4408.59899 | -0.1506385 | 0.01692366 | -8.9010572 | 5.5317E-19 | 1.3664E-16 |
| Cd55 | 239.436991 | 332.585151 | 0.71992688 | -0.4740777 | 286.011071 | -0.4727465 | 0.05360665 | -8.8188046 | 1.1569E-18 | 2.8108E-16 |
| Gpc1 | 3531.65431 | 4061.31705 | 0.8695835 | -0.2016035 | 3796.48568 | -0.1992365 | 0.0227363 | -8.7629241 | 1.9025E-18 | 4.5479E-16 |
| Etv5 | 57.157874 | 23.1546278 | 2.46852917 | 1.30365169 | 40.1562508 | 1.31890644 | 0.15245897 | 8.6508945 | 5.1097E-18 | 1.2021E-15 |
| Btbd11 | 467.370217 | 599.434663 | 0.779685 | -0.3590367 | 533.40244 | -0.3567234 | 0.04137486 | -8.6217427 | 6.5943E-18 | 1.5271E-15 |
| Fbp1 | 107.934951 | 172.082253 | 0.62722884 | -0.6729362 | 140.008602 | -0.6744014 | 0.07849696 | -8.5914334 | 8.5891E-18 | 1.9584E-15 |
| Hes6 | 2063.55703 | 2348.57967 | 0.87864042 | -0.1866552 | 2206.06835 | -0.1868436 | 0.02178367 | -8.5772318 | 9.7184E-18 | 2.1824E-15 |

| | | | | | | | | | | |
|---|---|---|---|---|---|---|---|---|---|---|
| Tmem163 | 1573.23343 | 1895.96501 | 0.82977978 | -0.2691996 | 1734.59922 | -0.2703531 | 0.03157953 | -8.5610246 | 1.1187E-17 | 2.4746E-15 |
| Tll1 | 34.0421574 | 75.2499975 | 0.45238749 | -1.1443691 | 54.6460774 | -1.1345189 | 0.13273572 | -8.547201 | 1.2611E-17 | 2.7486E-15 |
| Cyfip2 | 2066.81188 | 1813.80455 | 1.13948985 | 0.18838807 | 1940.30822 | 0.18707091 | 0.02195517 | 8.52058413 | 1.5875E-17 | 3.4099E-15 |
| Aldh1a2 | 269.134223 | 364.347926 | 0.73867368 | -0.4369909 | 316.741074 | -0.432729 | 0.05109518 | -8.469078 | 2.4734E-17 | 5.237E-15 |
| Aldh1a7 | 3525.49001 | 3169.60236 | 1.11228148 | 0.15352193 | 3347.54619 | 0.15358235 | 0.01820863 | 8.43458971 | 3.3237E-17 | 6.938E-15 |
| Rpl23 | 1596.21435 | 1360.70754 | 1.17307673 | 0.23029738 | 1478.46095 | 0.23038032 | 0.02732615 | 8.43076311 | 3.4342E-17 | 7.0692E-15 |
| Tmem196 | 41.4783485 | 83.7509135 | 0.49525846 | -1.0137465 | 62.6146309 | -1.0184226 | 0.12102072 | -8.4152745 | 3.9198E-17 | 7.9582E-15 |
| Dpp6 | 600.387429 | 786.989918 | 0.76289088 | -0.3904514 | 693.688674 | -0.374243 | 0.04448806 | -8.4122133 | 4.0235E-17 | 8.0583E-15 |
| Nfasc | 1947.41849 | 2231.00079 | 0.8728901 | -0.1961281 | 2089.20964 | -0.1966692 | 0.02355213 | -8.3503803 | 6.8044E-17 | 1.3446E-14 |
| Ccdc85a | 370.047088 | 488.777184 | 0.75708748 | -0.4014681 | 429.412136 | -0.4044993 | 0.04848187 | -8.343312 | 7.2238E-17 | 1.4087E-14 |
| Igfbp5 | 31621.3546 | 34677.8333 | 0.91186073 | -0.1331146 | 33149.594 | -0.1309932 | 0.01570841 | -8.339048 | 7.4891E-17 | 1.4415E-14 |
| Rpl19 | 2915.4665 | 2570.11741 | 1.13437094 | 0.18189248 | 2742.79196 | 0.18344112 | 0.02210229 | 8.29964216 | 1.0443E-16 | 1.9842E-14 |
| Rps8 | 2082.08359 | 1835.77429 | 1.13417188 | 0.1816393 | 1958.92894 | 0.18164692 | 0.02216399 | 8.19558705 | 2.4937E-16 | 4.6784E-14 |
| Dpysl2 | 1769.86962 | 2012.26475 | 0.87954113 | -0.185177 | 1891.06719 | -0.1865065 | 0.02280121 | -8.1796767 | 2.8461E-16 | 5.2727E-14 |
| Astn1 | 731.318121 | 900.633088 | 0.8120045 | -0.3004404 | 815.975604 | -0.2996155 | 0.03665811 | -8.17324 | 3.0022E-16 | 5.4932E-14 |
| Zeb2 | 55.9044855 | 100.401833 | 0.55680742 | -0.8447497 | 78.1531593 | -0.8435137 | 0.10346646 | -8.1525326 | 3.5638E-16 | 6.4414E-14 |
| Trib2 | 422.193622 | 316.12174 | 1.33554124 | 0.41742453 | 369.157681 | 0.41408827 | 0.05103701 | 8.11349046 | 4.9186E-16 | 8.7159E-14 |
| Psap | 6872.67157 | 7457.10707 | 0.92162705 | -0.117745 | 7164.88932 | -0.1174279 | 0.0144741 | -8.1129666 | 4.9399E-16 | 8.7159E-14 |
| Sv2b | 301.834063 | 396.970308 | 0.76034418 | -0.3952755 | 349.402185 | -0.3944824 | 0.04865273 | -8.1081259 | 5.1406E-16 | 8.9635E-14 |
| Pygb | 1854.99055 | 2120.20401 | 0.87491135 | -0.1927913 | 1987.59728 | -0.193113 | 0.02383351 | -8.1025817 | 5.3805E-16 | 9.2726E-14 |
| Actg1 | 11108.016 | 9989.14252 | 1.11200896 | 0.15316841 | 10548.5792 | 0.15469903 | 0.01910684 | 8.09652501 | 5.6551E-16 | 9.6339E-14 |
| Sox21 | 23.2396843 | 3.6720576 | 6.3287908 | 2.66192988 | 13.4558709 | 2.63180535 | 0.32563496 | 8.08207253 | 6.3675E-16 | 1.0724E-13 |
| Psat1 | 442.823211 | 324.381769 | 1.36512978 | 0.44903811 | 383.60249 | 0.45699497 | 0.05668608 | 8.06185535 | 7.5145E-16 | 1.2514E-13 |
| Dll4 | 261.981522 | 365.883287 | 0.71602484 | -0.4819185 | 313.932404 | -0.4793933 | 0.0595339 | -8.0524415 | 8.1159E-16 | 1.3365E-13 |
| Copa | 4393.01622 | 4837.02558 | 0.90820612 | -0.1389083 | 4615.0209 | -0.1383574 | 0.01719954 | -8.0442523 | 8.6774E-16 | 1.4133E-13 |
| Rps12 | 1055.53399 | 888.379554 | 1.18815655 | 0.24872494 | 971.956771 | 0.24757923 | 0.0307839 | 8.0424917 | 8.803E-16 | 1.4181E-13 |
| Kctd12 | 2382.90488 | 2674.72886 | 0.89089586 | -0.1666713 | 2528.81687 | -0.1653853 | 0.02056942 | -8.0403476 | 8.9584E-16 | 1.4277E-13 |
| Akt3 | 3048.47643 | 3390.76481 | 0.89905275 | -0.1535223 | 3219.62062 | -0.1526592 | 0.01909857 | -7.9932274 | 1.3145E-15 | 2.0726E-13 |
| Grb10 | 148.86291 | 89.7179039 | 1.65923304 | 0.73051653 | 119.290407 | 0.70972184 | 0.08890245 | 7.98315314 | 1.4264E-15 | 2.2254E-13 |
| Ctsc | 175.617726 | 250.129305 | 0.70210776 | -0.5102356 | 212.873515 | -0.5073431 | 0.06356925 | -7.980951 | 1.4521E-15 | 2.2418E-13 |
| Gm16294 | 18.1594839 | 0.22670627 | 80.1013741 | 6.32375509 | 9.193095 | 5.7645484 | 0.72512831 | 7.94969429 | 1.8697E-15 | 2.8277E-13 |
| Cntn2 | 128.122362 | 190.907104 | 0.67112412 | -0.5753485 | 159.514733 | -0.5706051 | 0.07177504 | -7.9499099 | 1.8665E-15 | 2.8277E-13 |
| Sgpp2 | 20.1832866 | 0.80798957 | 24.9796376 | 4.64268064 | 10.495638 | 4.64627377 | 0.5850441 | 7.94174968 | 1.9935E-15 | 2.9844E-13 |
| Sgip1 | 22.9944752 | 4.86790548 | 4.72368974 | 2.23991421 | 13.9311902 | 2.23959287 | 0.28270206 | 7.92209593 | 2.3354E-15 | 3.4613E-13 |
| Gabra4 | 18.2785973 | 0.80828395 | 22.6140793 | 4.49914936 | 9.54344055 | 4.45032984 | 0.56486388 | 7.87858806 | 3.311E-15 | 4.8304E-13 |
| Tubb3 | 666.691576 | 807.539125 | 0.82558424 | -0.2765127 | 737.11535 | -0.2805033 | 0.03560552 | -7.8780848 | 3.3244E-15 | 4.8304E-13 |
| Unc80 | 3272.49775 | 3601.90762 | 0.90854572 | -0.138369 | 3437.20268 | -0.1388709 | 0.01768225 | -7.8536906 | 4.0397E-15 | 5.8129E-13 |
| Tmem132e | 81.7435717 | 132.766796 | 0.61569289 | -0.6997172 | 107.255184 | -0.7095723 | 0.09044986 | -7.844924 | 4.3322E-15 | 6.1737E-13 |
| Rps21 | 1729.40828 | 1508.15369 | 1.1467056 | 0.19749505 | 1618.78098 | 0.19976704 | 0.02575507 | 7.75641637 | 8.7363E-15 | 1.2241E-12 |
| Dnm3 | 662.821162 | 839.530303 | 0.78951428 | -0.3409627 | 751.175732 | -0.3283423 | 0.04233318 | -7.7561448 | 8.755E-15 | 1.2241E-12 |
| St6galnac5 | 108.348158 | 64.704544 | 1.67450617 | 0.74373569 | 86.5263509 | 0.74031726 | 0.09554295 | 7.74852818 | 9.2964E-15 | 1.2758E-12 |
| Id1 | 1152.0793 | 970.72671 | 1.18682147 | 0.24710293 | 1061.403 | 0.24729302 | 0.03191486 | 7.74852368 | 9.2967E-15 | 1.2758E-12 |
| Fxyd6 | 1673.48202 | 2013.67579 | 0.83105832 | -0.2669784 | 1843.57891 | -0.259295 | 0.03378443 | -7.6749841 | 1.6544E-14 | 2.2495E-12 |
| Dach1 | 406.775488 | 508.57231 | 0.79983806 | -0.3222202 | 457.673899 | -0.3214544 | 0.04191759 | -7.6687225 | 1.7372E-14 | 2.3406E-12 |
| Irf6 | 2920.992 | 3235.36844 | 0.90283133 | -0.1474716 | 3078.18022 | -0.1469365 | 0.0193043 | -7.6115944 | 2.7074E-14 | 3.6149E-12 |
| Nme2 | 906.332403 | 720.312678 | 1.25824858 | 0.33141697 | 813.322541 | 0.34215959 | 0.04521573 | 7.56726853 | 3.8115E-14 | 4.9553E-12 |
| Bdnf | 82.0477182 | 130.666276 | 0.62791809 | -0.6713517 | 106.356997 | -0.6680324 | 0.08826883 | -7.5681568 | 3.7856E-14 | 4.9553E-12 |
| Cabp2 | 17.4357275 | 43.9937792 | 0.39632257 | -1.335253 | 30.7147533 | -1.336098 | 0.17652773 | -7.5687714 | 3.7677E-14 | 4.9553E-12 |
| Ankrd33b | 258.548338 | 179.646095 | 1.43920934 | 0.52527645 | 219.097216 | 0.53053365 | 0.07013363 | 7.56461088 | 3.8903E-14 | 5.0137E-12 |
| Flt4 | 166.163274 | 111.114799 | 1.49541983 | 0.58055057 | 138.639036 | 0.58155433 | 0.07693015 | 7.55951128 | 4.0459E-14 | 5.1693E-12 |
| Actn1 | 3282.65106 | 3617.92543 | 0.90732966 | -0.1403013 | 3450.28824 | -0.1394349 | 0.01850918 | -7.5332814 | 4.9481E-14 | 6.268E-12 |
| Gpx1 | 17.1114235 | 0.35317379 | 48.4504344 | 5.5984377 | 8.73229857 | 5.46777003 | 0.72725398 | 7.51837756 | 5.546E-14 | 6.9659E-12 |
| Negr1 | 17.1652487 | 0.33523743 | 51.2032585 | 5.67816372 | 8.75024298 | 5.4062328 | 0.71932405 | 7.51571254 | 5.6602E-14 | 7.0495E-12 |
| Col8a1 | 12389.0088 | 13419.7972 | 0.92318898 | -0.1153021 | 12904.403 | -0.1142019 | 0.01539464 | -7.4182878 | 1.1864E-13 | 1.4654E-11 |
| Tenm1 | 395.00075 | 496.765788 | 0.79514483 | -0.3307104 | 445.883269 | -0.3342835 | 0.04511705 | -7.4092489 | 1.2702E-13 | 1.5558E-11 |
| Kcnn3 | 318.942238 | 409.913103 | 0.77807281 | -0.3620229 | 364.42767 | -0.3622932 | 0.04901756 | -7.3910906 | 1.4563E-13 | 1.7575E-11 |
| Lrrn2 | 312.532191 | 423.258866 | 0.73839491 | -0.4375355 | 367.895529 | -0.4300377 | 0.0581849 | -7.3908829 | 1.4586E-13 | 1.7575E-11 |
| Csrp1 | 3284.03331 | 3593.2216 | 0.91395235 | -0.1298091 | 3438.62745 | -0.1298951 | 0.0175837 | -7.3872411 | 1.4991E-13 | 1.7917E-11 |
| Yif1a | 14.5911446 | 0.23510138 | 62.0632023 | 5.95566623 | 7.41312288 | 5.63136474 | 0.76309055 | 7.37968092 | 1.5867E-13 | 1.8664E-11 |
| Neat1 | 455.711508 | 302.103829 | 1.50845989 | 0.59307633 | 378.907668 | 0.60206844 | 0.08157692 | 7.3803768 | 1.5784E-13 | 1.8664E-11 |
| Plk5 | 708.907422 | 835.319659 | 0.84866603 | -0.2367312 | 772.11354 | -0.2367563 | 0.03243928 | -7.2984448 | 2.9111E-13 | 3.3973E-11 |
| Rpl12 | 1053.41512 | 906.393516 | 1.16220505 | 0.21686462 | 979.904317 | 0.21870401 | 0.0300831 | 7.26999688 | 3.595E-13 | 4.1626E-11 |
| Rps24 | 2444.39484 | 2198.83701 | 1.11167623 | 0.15273667 | 2321.61593 | 0.15229024 | 0.02100922 | 7.24873183 | 4.2069E-13 | 4.8334E-11 |
| Rassf4 | 351.197729 | 461.233795 | 0.76143104 | -0.3932147 | 406.215762 | -0.390803 | 0.0542296 | -7.2064521 | 5.7429E-13 | 6.5473E-11 |
| Tro | 32.3732392 | 11.6976369 | 2.76750248 | 1.46858461 | 22.035438 | 1.46217514 | 0.20317757 | 7.1965383 | 6.176E-13 | 6.9874E-11 |
| Tfcp2l1 | 979.552266 | 1139.26865 | 0.85980797 | -0.2179136 | 1059.41046 | -0.2179632 | 0.03030716 | -7.1918049 | 6.394E-13 | 7.1792E-11 |
| Gpr88 | 40.5872434 | 17.5457875 | 2.31321868 | 1.20990166 | 29.0665153 | 1.20970923 | 0.16833626 | 7.18626651 | 6.6587E-13 | 7.4202E-11 |

| | | | | | | | | | | |
|---|---|---|---|---|---|---|---|---|---|---|
| Svil | 2513.70635 | 2807.94368 | 0.89521253 | -0.1596979 | 2660.82502 | -0.161262 | 0.02256674 | -7.1460045 | 8.934E-13 | 9.8814E-11 |
| Kit | 1011.44887 | 1176.48432 | 0.8597215 | -0.2180587 | 1093.96659 | -0.2201789 | 0.03098732 | -7.1054527 | 1.1993E-12 | 1.3166E-10 |
| Tuba1a | 6528.25668 | 7000.15249 | 0.93258778 | -0.1006886 | 6764.20458 | -0.1006095 | 0.01419883 | -7.0857588 | 1.3828E-12 | 1.507E-10 |
| Kdm2a | 3163.61873 | 3458.45715 | 0.91474857 | -0.1285528 | 3311.03794 | -0.1285322 | 0.01819423 | -7.0644523 | 1.6125E-12 | 1.7445E-10 |
| Ptprz1 | 2653.52181 | 2382.72964 | 1.11364788 | 0.15529314 | 2518.12573 | 0.15313874 | 0.0217102 | 7.05377 | 1.7413E-12 | 1.8702E-10 |
| Cadm2 | 852.397103 | 994.109694 | 0.85744773 | -0.2218794 | 923.253398 | -0.2214133 | 0.03146198 | -7.037488 | 1.9574E-12 | 2.0871E-10 |
| Sema5a | 268.16339 | 353.706762 | 0.75815172 | -0.3994415 | 310.935075 | -0.3871649 | 0.05507942 | -7.0292115 | 2.077E-12 | 2.1988E-10 |
| Pld5 | 21.0923797 | 46.8275549 | 0.45042667 | -1.1506358 | 33.9599672 | -1.1564098 | 0.16471041 | -7.0208664 | 2.205E-12 | 2.3177E-10 |
| Rpl7 | 2858.81942 | 2548.71694 | 1.12167004 | 0.16564834 | 2703.76818 | 0.16650763 | 0.02374989 | 7.01088073 | 2.3682E-12 | 2.4718E-10 |
| Nsg2 | 102.532523 | 199.895067 | 0.51293173 | -0.9631613 | 151.213795 | -0.9257577 | 0.13310848 | -6.9549115 | 3.5278E-12 | 3.6564E-10 |
| Srebf2 | 2197.51378 | 2456.47926 | 0.8945786 | -0.1607198 | 2326.99652 | -0.1594513 | 0.02293464 | -6.9524224 | 3.5907E-12 | 3.6956E-10 |
| Rpl5 | 3082.92805 | 2802.76316 | 1.09996024 | 0.13745138 | 2942.84561 | 0.1376266 | 0.01982296 | 6.94278695 | 3.8444E-12 | 3.9295E-10 |
| Rapgef4 | 716.820495 | 847.15843 | 0.84614692 | -0.2410199 | 781.989463 | -0.2407463 | 0.03469877 | -6.9381806 | 3.9718E-12 | 4.0319E-10 |
| Dusp4 | 60.0521937 | 32.2838787 | 1.86012946 | 0.89540303 | 46.1680361 | 0.89520226 | 0.12908039 | 6.93523058 | 4.0556E-12 | 4.089E-10 |
| Hmgb2 | 4470.1921 | 4034.30356 | 1.10804555 | 0.14801719 | 4252.24783 | 0.15211716 | 0.02199381 | 6.9163631 | 4.6339E-12 | 4.6404E-10 |
| Cacna2d1 | 4339.31855 | 4701.866 | 0.92289286 | -0.1157649 | 4520.59227 | -0.1163021 | 0.01684552 | -6.904036 | 5.0545E-12 | 5.0277E-10 |
| Ppp3ca | 5105.47295 | 5506.85684 | 0.92711198 | -0.1091845 | 5306.1649 | -0.1091749 | 0.01582329 | -6.8996309 | 5.2138E-12 | 5.1516E-10 |
| Tacc2 | 367.290073 | 454.256722 | 0.80855176 | -0.306588 | 410.773397 | -0.3067874 | 0.04448592 | -6.8962801 | 5.3382E-12 | 5.2396E-10 |
| Upf2 | 424.173408 | 519.864146 | 0.81593126 | -0.2934805 | 472.018776 | -0.2936017 | 0.04262962 | -6.8872697 | 5.6873E-12 | 5.5455E-10 |
| Klc2 | 12.4866396 | 0.35412494 | 35.2605481 | 5.139983 | 6.4203822 | 4.95992875 | 0.72152005 | 6.87427705 | 6.2305E-12 | 5.9981E-10 |
| Ptprf | 3916.89006 | 4249.86698 | 0.92165004 | -0.117709 | 4083.37852 | -0.1187648 | 0.01727681 | -6.8742341 | 6.2324E-12 | 5.9981E-10 |
| Ccdc71 | 11.7370628 | 0.23438877 | 50.0751925 | 5.64602416 | 5.98572571 | 5.27710417 | 0.77203664 | 6.8353027 | 8.1832E-12 | 7.8247E-10 |
| Pcdhb20 | 109.783148 | 159.494152 | 0.68832084 | -0.5388469 | 134.63865 | -0.5389859 | 0.07888833 | -6.8322636 | 8.3585E-12 | 7.9411E-10 |
| Fam210b | 433.530102 | 544.240421 | 0.79657829 | -0.3281119 | 488.885261 | -0.3236173 | 0.04752728 | -6.8090851 | 9.8221E-12 | 9.2722E-10 |
| Neto2 | 3827.32634 | 3447.14488 | 1.1102888 | 0.15093498 | 3637.23561 | 0.15158585 | 0.02229415 | 6.79935657 | 1.0509E-11 | 9.8576E-10 |
| Slc38a11 | 145.669601 | 202.664893 | 0.71877077 | -0.4763964 | 174.167247 | -0.4772954 | 0.07026225 | -6.7930557 | 1.0978E-11 | 1.0233E-09 |
| Eda2r | 14.1258045 | 0.67073105 | 21.060311 | 4.39645484 | 7.39826766 | 4.31983557 | 0.63731852 | 6.77814227 | 1.2173E-11 | 1.1276E-09 |
| Mid1 | 408.063128 | 326.144618 | 1.25117235 | 0.32328054 | 367.103873 | 0.32322227 | 0.04805828 | 6.72563059 | 1.7483E-11 | 1.6094E-09 |
| F5 | 3663.70494 | 3978.88408 | 0.92078705 | -0.1190605 | 3821.29451 | -0.119026 | 0.01774641 | -6.7070509 | 1.986E-11 | 1.8156E-09 |
| Foxp2 | 125.463742 | 177.298841 | 0.70763995 | -0.4989126 | 151.381292 | -0.4974019 | 0.07416985 | -6.7062545 | 1.9968E-11 | 1.8156E-09 |
| Brms1 | 8.30387265 | 0.0000001 | 83038726.5 | 26.307281 | 4.15193628 | 5.37944113 | 0.80270868 | 6.70161073 | 2.0613E-11 | 1.8629E-09 |
| Tmcc2 | 1124.77367 | 1325.39136 | 0.84863513 | -0.2367837 | 1225.08251 | -0.2300458 | 0.03436658 | -6.693882 | 2.1733E-11 | 1.9521E-09 |
| Ece1 | 2458.8225 | 2721.88857 | 0.90335164 | -0.1466404 | 2590.35554 | -0.1450541 | 0.02169256 | -6.6868137 | 2.2808E-11 | 2.0364E-09 |
| Rpl27 | 1350.00666 | 1182.11526 | 1.14202624 | 0.1915958 | 1266.06096 | 0.19483843 | 0.02919934 | 6.67269905 | 2.5114E-11 | 2.2288E-09 |
| Plscr4 | 318.36309 | 399.464485 | 0.7969747 | -0.3273942 | 358.913788 | -0.3266919 | 0.04899658 | -6.6676465 | 2.5994E-11 | 2.2932E-09 |
| Strc | 13.3157148 | 1.58819375 | 8.38418785 | 3.06767104 | 7.45195415 | 3.01422937 | 0.45236806 | 6.66322331 | 2.6789E-11 | 2.3493E-09 |
| Chrdl1 | 8.8947697 | 0.11790705 | 75.4388313 | 6.23723542 | 4.50633827 | 5.24333813 | 0.78776567 | 6.65596167 | 2.8145E-11 | 2.4538E-09 |
| Pogk | 1162.49828 | 1011.77879 | 1.14896486 | 0.20033468 | 1087.13854 | 0.20048384 | 0.03017308 | 6.64445994 | 3.0433E-11 | 2.6377E-09 |
| Wnt5a | 65.0439244 | 104.308036 | 0.62357539 | -0.6813641 | 84.6759799 | -0.6758765 | 0.10176506 | -6.6415377 | 3.1043E-11 | 2.6749E-09 |
| Rgs16 | 794.858822 | 970.117182 | 0.8193431 | -0.2874604 | 882.488002 | -0.286502 | 0.04321267 | -6.6300466 | 3.3558E-11 | 2.8749E-09 |
| B4gat1 | 8.49396523 | 0.11719444 | 72.4775471 | 6.17946222 | 4.30557973 | 5.19480703 | 0.78369667 | 6.62859394 | 3.389E-11 | 2.8867E-09 |
| Pitx2 | 42.2762822 | 74.6134946 | 0.5666037 | -0.8195881 | 58.4448883 | -0.8370053 | 0.12647972 | -6.617704 | 3.6482E-11 | 3.0897E-09 |
| Gm16159 | 6.55128884 | 0.0000001 | 65512888.4 | 25.9652754 | 3.27564437 | 5.29318939 | 0.80174113 | 6.60211781 | 4.0533E-11 | 3.4133E-09 |
| Jag1 | 387.977631 | 486.718787 | 0.79712894 | -0.327115 | 437.348209 | -0.3244118 | 0.04929359 | -6.5812179 | 4.6661E-11 | 3.9071E-09 |
| Ildr2 | 29.4385373 | 58.5833222 | 0.50250713 | -0.992784 | 44.0109297 | -0.9887904 | 0.15049056 | -6.5704477 | 5.0164E-11 | 4.1769E-09 |
| Ppp1r3a | 552.834392 | 687.08597 | 0.80460731 | -0.3136433 | 619.960181 | -0.3015733 | 0.04592734 | -6.5663128 | 5.1576E-11 | 4.2705E-09 |
| Rplp1 | 2137.71084 | 1940.82995 | 1.1014416 | 0.13939301 | 2039.2704 | 0.13950748 | 0.02128616 | 6.55390413 | 5.6052E-11 | 4.6153E-09 |
| Clca3a1 | 192.463589 | 138.454818 | 1.39008228 | 0.47517028 | 165.459203 | 0.46973546 | 0.07202643 | 6.52170954 | 6.9511E-11 | 5.6918E-09 |
| Rps7 | 2159.46703 | 1947.58739 | 1.10879082 | 0.14898722 | 2053.52721 | 0.15078973 | 0.02318931 | 6.50255482 | 7.8967E-11 | 6.4306E-09 |
| Acsl5 | 862.07865 | 737.140706 | 1.16948995 | 0.22587947 | 799.609678 | 0.22511493 | 0.03464496 | 6.49776829 | 8.152E-11 | 6.6022E-09 |
| Etv4 | 27.7096233 | 9.84237859 | 2.81533809 | 1.49330818 | 18.7760009 | 1.4884153 | 0.22993791 | 6.47311842 | 9.6001E-11 | 7.7327E-09 |
| Mbip | 295.341908 | 229.416614 | 1.28736059 | 0.36441621 | 262.379261 | 0.36422173 | 0.05628783 | 6.47070142 | 9.7549E-11 | 7.815E-09 |
| Smarca1 | 681.956366 | 804.455694 | 0.84772396 | -0.2383335 | 743.20603 | -0.2362028 | 0.03663493 | -6.4474734 | 1.1373E-10 | 9.0623E-09 |
| Eef1a1 | 27173.5328 | 25630.7182 | 1.06019396 | 0.08432823 | 26402.1255 | 0.08474588 | 0.01316025 | 6.43953496 | 1.1984E-10 | 9.4981E-09 |
| Nme1 | 1047.04315 | 904.940413 | 1.15702994 | 0.2104262 | 975.991782 | 0.21411482 | 0.03329136 | 6.43154367 | 1.2631E-10 | 9.958E-09 |
| Scn3b | 18.9179183 | 40.3982507 | 0.46828558 | -1.0945395 | 29.6580844 | -1.0949456 | 0.1706212 | -6.4174064 | 1.3862E-10 | 1.087E-08 |
| Gm3002 | 1694.36777 | 1344.97486 | 1.25977653 | 0.33316784 | 1519.67132 | 0.34129919 | 0.05321805 | 6.41322284 | 1.4248E-10 | 1.1114E-08 |
| Thbs4 | 1179.42832 | 1338.71546 | 0.88101494 | -0.1827616 | 1259.07189 | -0.1830431 | 0.0286105 | -6.3977593 | 1.5767E-10 | 1.2235E-08 |
| Ppp1ca | 7488.75235 | 7929.42313 | 0.94442587 | -0.0824905 | 7709.08774 | -0.08265 | 0.01292229 | -6.3959254 | 1.5958E-10 | 1.2318E-08 |
| Chrna3 | 80.7819041 | 47.3264865 | 1.70690685 | 0.77138433 | 64.0541952 | 0.77664819 | 0.12170537 | 6.38137967 | 1.755E-10 | 1.3477E-08 |
| Nrxn1 | 2124.12174 | 2342.46863 | 0.9067877 | -0.1411633 | 2233.29519 | -0.1401926 | 0.02198285 | -6.3773645 | 1.8016E-10 | 1.3764E-08 |
| Rps15a | 1218.12803 | 1082.79809 | 1.12498169 | 0.16990153 | 1150.46306 | 0.16968588 | 0.02666632 | 6.36330325 | 1.9746E-10 | 1.5008E-08 |
| Egfl6 | 380.743672 | 463.666351 | 0.82115873 | -0.284267 | 422.205012 | -0.2849972 | 0.04481417 | -6.3595331 | 2.0237E-10 | 1.5303E-08 |
| Plcb1 | 467.673879 | 383.538335 | 1.21936671 | 0.28613207 | 425.606107 | 0.28613586 | 0.04520874 | 6.32921602 | 2.4641E-10 | 1.8538E-08 |
| Tbc1d9 | 386.083469 | 467.744559 | 0.8254152 | -0.2768081 | 426.914014 | -0.2756776 | 0.04364806 | -6.3159184 | 2.6856E-10 | 2.0103E-08 |
| Sox5 | 68.2000373 | 104.015881 | 0.65566947 | -0.6089594 | 86.107959 | -0.608324 | 0.09640586 | -6.3100309 | 2.7898E-10 | 2.0778E-08 |
| Med22 | 1780.04835 | 1965.40409 | 0.90569077 | -0.1429095 | 1872.72622 | -0.1434005 | 0.02273983 | -6.3061368 | 2.8609E-10 | 2.12E-08 |

| | | | | | | | | | | |
|---|---|---|---|---|---|---|---|---|---|---|
| Pcsk2 | 2439.68182 | 2709.44241 | 0.90043686 | -0.151303 | 2574.56212 | -0.1522262 | 0.0242452 | -6.2786122 | 3.4161E-10 | 2.5189E-08 |
| Klhdc8b | 7.29817253 | 0.11798978 | 61.8542785 | 5.95080148 | 3.70808105 | 5.00939795 | 0.79837515 | 6.27449133 | 3.5078E-10 | 2.5737E-08 |
| Mcm6 | 4223.92893 | 4617.02676 | 0.9148591 | -0.1283785 | 4420.47784 | -0.1293879 | 0.02062881 | -6.2721935 | 3.56E-10 | 2.5991E-08 |
| Adgrf5 | 43.9954705 | 74.3245615 | 0.59193717 | -0.756484 | 59.1600159 | -0.7537442 | 0.12043612 | -6.2584559 | 3.8881E-10 | 2.8248E-08 |
| Antxr1 | 248.784466 | 313.698065 | 0.79306981 | -0.3344802 | 281.241266 | -0.3394685 | 0.05425816 | -6.2565417 | 3.9361E-10 | 2.8457E-08 |
| Capn2 | 2307.27881 | 2524.72885 | 0.91387192 | -0.1299361 | 2416.00383 | -0.1300076 | 0.02083233 | -6.2406638 | 4.3572E-10 | 3.1348E-08 |
| Prom1 | 72.8509931 | 108.875782 | 0.66912027 | -0.5796625 | 90.8633876 | -0.5810404 | 0.0932808 | -6.2289391 | 4.696E-10 | 3.3623E-08 |
| Dhrs3 | 170.503945 | 121.647117 | 1.4016275 | 0.48710298 | 146.075531 | 0.48462778 | 0.07783994 | 6.22595261 | 4.7864E-10 | 3.4105E-08 |
| Hmgcs1 | 1357.76554 | 1555.91301 | 0.87264874 | -0.196527 | 1456.83928 | -0.1920581 | 0.03086342 | -6.2228384 | 4.8824E-10 | 3.4623E-08 |
| Cnot6 | 2882.50276 | 2583.16262 | 1.11588126 | 0.15818352 | 2732.83269 | 0.15955392 | 0.02575292 | 6.1955673 | 5.8075E-10 | 4.0793E-08 |
| Rpsa | 5241.30006 | 4900.86769 | 1.06946369 | 0.0968875 | 5071.08387 | 0.0969253 | 0.0156427 | 6.19619878 | 5.7843E-10 | 4.0793E-08 |
| Nicn1 | 10.2281268 | 0.58056305 | 17.6175988 | 4.1389454 | 5.40434483 | 4.12244365 | 0.66555102 | 6.19403097 | 5.8645E-10 | 4.0999E-08 |
| Rpl11 | 2947.11816 | 2684.00014 | 1.09803204 | 0.13492015 | 2815.55915 | 0.13710175 | 0.02217772 | 6.18195737 | 6.3312E-10 | 4.4009E-08 |
| Syn2 | 503.37706 | 606.520323 | 0.82994261 | -0.2689165 | 554.948691 | -0.2662157 | 0.04306737 | -6.1813783 | 6.3544E-10 | 4.4009E-08 |
| Dpp3 | 1360.94058 | 1519.13653 | 0.89586456 | -0.1586475 | 1440.03855 | -0.1586023 | 0.02575436 | -6.1582691 | 7.3544E-10 | 5.0698E-08 |
| Rpl6 | 2490.95973 | 2275.5488 | 1.09466329 | 0.13048717 | 2383.25427 | 0.13100567 | 0.02130661 | 6.14859244 | 7.8174E-10 | 5.3639E-08 |
| Eif3a | 1810.3297 | 1608.20447 | 1.12568379 | 0.17080162 | 1709.26709 | 0.16976504 | 0.0276575 | 6.13811874 | 8.3504E-10 | 5.7033E-08 |
| Rpl3 | 3204.45735 | 2953.96615 | 1.08479826 | 0.11742677 | 3079.21175 | 0.11715891 | 0.01910975 | 6.13084358 | 8.7414E-10 | 5.943E-08 |
| Kirrel3 | 121.910278 | 171.471252 | 0.71096628 | -0.492147 | 146.690765 | -0.4774541 | 0.077973 | -6.1233257 | 9.1642E-10 | 6.202E-08 |
| Dnajc12 | 818.157779 | 951.5145 | 0.85984794 | -0.2178466 | 884.83614 | -0.2194103 | 0.03587423 | -6.1160966 | 9.5895E-10 | 6.4603E-08 |
| Tmtc1 | 96.1806453 | 141.797965 | 0.67829355 | -0.5600183 | 118.989305 | -0.5563859 | 0.09112891 | -6.105482 | 1.0249E-09 | 6.8734E-08 |
| Col18a1 | 454.971844 | 351.382314 | 1.29480576 | 0.37273569 | 403.177079 | 0.37502299 | 0.06146905 | 6.10100471 | 1.054E-09 | 7.0369E-08 |
| Vps13d | 1366.28714 | 1536.52071 | 0.88920841 | -0.1694065 | 1451.40393 | -0.1746752 | 0.02867081 | -6.092439 | 1.112E-09 | 7.3908E-08 |
| Plekha6 | 893.509891 | 1013.86362 | 0.88129199 | -0.182308 | 953.686757 | -0.1823801 | 0.02996836 | -6.0857556 | 1.1594E-09 | 7.6714E-08 |
| Decr1 | 97.9094574 | 138.181038 | 0.70855929 | -0.4970395 | 118.045247 | -0.4972382 | 0.08177866 | -6.0802929 | 1.1996E-09 | 7.9021E-08 |
| Ube2c | 2450.3591 | 2208.40537 | 1.10956038 | 0.14998818 | 2329.38224 | 0.1547443 | 0.02547243 | 6.07497202 | 1.2401E-09 | 8.1325E-08 |
| Rps27a | 906.952841 | 792.609007 | 1.1442626 | 0.19441818 | 849.780924 | 0.19515585 | 0.03214643 | 6.07084106 | 1.2724E-09 | 8.3077E-08 |
| Dpysl3 | 38.083392 | 65.1803863 | 0.58427687 | -0.7752759 | 51.6318891 | -0.7889111 | 0.13026758 | -6.0560814 | 1.3948E-09 | 9.0666E-08 |
| Rps27l | 518.974656 | 433.34642 | 1.19759765 | 0.2601433 | 476.160538 | 0.25758823 | 0.0425985 | 6.04688449 | 1.4767E-09 | 9.5575E-08 |
| Sez6l2 | 1602.38717 | 1770.57014 | 0.90501197 | -0.1439912 | 1686.47865 | -0.1430119 | 0.02366368 | -6.0435186 | 1.5079E-09 | 9.7167E-08 |
| Pkm | 7099.2304 | 7515.92535 | 0.9445584 | -0.0822881 | 7307.57788 | -0.0823735 | 0.01366631 | -6.0274841 | 1.6653E-09 | 1.0685E-07 |
| Kitl | 542.905159 | 658.46214 | 0.82450475 | -0.2784003 | 600.68365 | -0.2762325 | 0.04584484 | -6.025379 | 1.6871E-09 | 1.0778E-07 |
| Rplp0 | 4790.6075 | 4474.28138 | 1.07069875 | 0.09855263 | 4632.44444 | 0.09909584 | 0.01646048 | 6.02022677 | 1.7417E-09 | 1.1079E-07 |
| Pnisr | 760.501214 | 657.649026 | 1.15639374 | 0.2096327 | 709.07512 | 0.21158484 | 0.03520064 | 6.0108245 | 1.8458E-09 | 1.1691E-07 |
| Mapk10 | 40.4790883 | 67.8230188 | 0.59683407 | -0.7445982 | 54.1510534 | -0.7556684 | 0.12601983 | -5.9964248 | 2.0171E-09 | 1.2721E-07 |
| Ulbp1 | 50.1284957 | 79.0061725 | 0.63448835 | -0.6563344 | 64.567334 | -0.664489 | 0.11083894 | -5.9950861 | 2.0338E-09 | 1.2772E-07 |
| Fnbp1l | 627.666927 | 733.399926 | 0.85583173 | -0.2246009 | 680.533427 | -0.22495 | 0.03754578 | -5.9913526 | 2.081E-09 | 1.3014E-07 |
| Oat | 14.326091 | 3.34498266 | 4.28285958 | 2.09857438 | 8.83553674 | 2.09483341 | 0.35136502 | 5.96198628 | 2.4919E-09 | 1.5518E-07 |
| Bcas1 | 230.715719 | 290.353557 | 0.79460269 | -0.3316944 | 260.534638 | -0.330913 | 0.05557841 | -5.9539856 | 2.6169E-09 | 1.6228E-07 |
| St6gal1 | 365.182189 | 294.089429 | 1.24173857 | 0.31236147 | 329.635809 | 0.31077041 | 0.05232959 | 5.93871307 | 2.8727E-09 | 1.774E-07 |
| Nrp1 | 582.058198 | 467.318598 | 1.24552757 | 0.31675696 | 524.688398 | 0.30997059 | 0.05228859 | 5.92807337 | 3.0651E-09 | 1.885E-07 |
| Arhgap24 | 265.443126 | 335.705967 | 0.79070125 | -0.3387954 | 300.574546 | -0.338193 | 0.05713261 | -5.9194394 | 3.2304E-09 | 1.9784E-07 |
| Gch1 | 1926.74118 | 2114.67029 | 0.91113077 | -0.13427 | 2020.70574 | -0.1349392 | 0.02283878 | -5.908336 | 3.4558E-09 | 2.1078E-07 |
| Rpl4 | 4154.92645 | 3856.98563 | 1.07724707 | 0.10734917 | 4005.95604 | 0.10795066 | 0.01827696 | 5.90637974 | 3.4971E-09 | 2.1242E-07 |
| Krt7 | 7718.25847 | 8156.27606 | 0.94629687 | -0.0796352 | 7937.26726 | -0.0794398 | 0.0134731 | -5.896178 | 3.7202E-09 | 2.2436E-07 |
| Thsd7a | 275.183038 | 339.508971 | 0.81053245 | -0.3030581 | 307.346005 | -0.3049323 | 0.05171844 | -5.8960075 | 3.724E-09 | 2.2436E-07 |
| Nov | 25048.1092 | 27689.7421 | 0.90459886 | -0.1446499 | 26368.9257 | -0.1446471 | 0.02454818 | -5.892377 | 3.8068E-09 | 2.2842E-07 |
| P4hb | 1671.28031 | 1496.74613 | 1.11660907 | 0.15912418 | 1584.01322 | 0.16083935 | 0.02734435 | 5.88199559 | 4.0535E-09 | 2.4225E-07 |
| Kif20b | 408.904405 | 330.728345 | 1.23637544 | 0.30611691 | 369.816375 | 0.30991263 | 0.05273882 | 5.87636596 | 4.1937E-09 | 2.4962E-07 |
| Clca3a2 | 103.069224 | 69.8077271 | 1.476473 | 0.56215497 | 86.4384756 | 0.55774921 | 0.09500669 | 5.87063075 | 4.3414E-09 | 2.5635E-07 |
| Pip4k2c | 1608.26918 | 1776.42639 | 0.90533961 | -0.143469 | 1692.34779 | -0.1434934 | 0.02444136 | -5.8709239 | 4.3337E-09 | 2.5635E-07 |
| Kalrn | 1317.4896 | 1470.6888 | 0.89583167 | -0.1587004 | 1394.0892 | -0.1623163 | 0.02766299 | -5.8676325 | 4.4206E-09 | 2.5999E-07 |
| Kcnh2 | 939.829003 | 1077.22197 | 0.87245621 | -0.1968454 | 1008.52549 | -0.1958665 | 0.03339369 | -5.8653741 | 4.4812E-09 | 2.6251E-07 |
| Slc7a5 | 472.126506 | 391.2197 | 1.20680658 | 0.27119446 | 431.673103 | 0.27843067 | 0.04747747 | 5.86447995 | 4.5054E-09 | 2.6289E-07 |
| Pum3 | 229.664114 | 179.208574 | 1.28154646 | 0.35788578 | 204.436344 | 0.35752157 | 0.06102568 | 5.8585426 | 4.6695E-09 | 2.714E-07 |
| Rps17 | 1388.46757 | 1247.15313 | 1.11330961 | 0.15485486 | 1317.81035 | 0.15584579 | 0.02670502 | 5.83582336 | 5.3526E-09 | 3.0988E-07 |
| Cenpa | 823.929409 | 718.714151 | 1.14639375 | 0.19710265 | 771.32178 | 0.20115779 | 0.03452025 | 5.82724103 | 5.6351E-09 | 3.2497E-07 |
| Rps2 | 3490.41168 | 3229.36846 | 1.08083414 | 0.11214515 | 3359.89007 | 0.11304911 | 0.01947294 | 5.80544529 | 6.4195E-09 | 3.6877E-07 |
| Rpl10 | 2624.20431 | 2417.05121 | 1.08570489 | 0.11863201 | 2520.62776 | 0.11846337 | 0.02044349 | 5.79467509 | 6.8454E-09 | 3.9172E-07 |
| Fdps | 1071.60641 | 1203.33247 | 0.89053228 | -0.1672602 | 1137.46944 | -0.1652343 | 0.02856153 | -5.7852037 | 7.2425E-09 | 4.1285E-07 |
| Tnks2 | 862.56034 | 742.515308 | 1.16167348 | 0.21620461 | 802.537824 | 0.21389322 | 0.03700421 | 5.78023987 | 7.4594E-09 | 4.2359E-07 |
| Basp1 | 2208.3043 | 1975.13888 | 1.11805014 | 0.16098489 | 2091.72159 | 0.1611193 | 0.02792304 | 5.77012026 | 7.9215E-09 | 4.4811E-07 |
| Ftsj3 | 450.828449 | 377.297435 | 1.19488872 | 0.25687626 | 414.062942 | 0.25744009 | 0.04466314 | 5.76403903 | 8.2124E-09 | 4.628E-07 |
| Adgrv1 | 427.796283 | 518.403248 | 0.82521914 | -0.2771508 | 473.099765 | -0.2904011 | 0.050421 | -5.7595265 | 8.435E-09 | 4.7354E-07 |
| Nmu | 5.52954779 | 20.0340358 | 0.27600768 | -1.8572197 | 12.7817917 | -1.7648649 | 0.30662845 | -5.7557114 | 8.6278E-09 | 4.8254E-07 |
| Ppia | 6816.15745 | 6386.94576 | 1.0672014 | 0.09383246 | 6601.55161 | 0.09703803 | 0.01687893 | 5.74906372 | 8.9739E-09 | 5.0001E-07 |
| Trp73 | 122.707511 | 81.4598603 | 1.50635553 | 0.59106231 | 102.083685 | 0.5924109 | 0.10306644 | 5.74785439 | 9.0383E-09 | 5.0171E-07 |

| | | | | | | | | | | |
|---|---|---|---|---|---|---|---|---|---|---|
| App | 6438.57164 | 6792.06303 | 0.94795523 | -0.0771092 | 6615.31733 | -0.0771264 | 0.0134392 | -5.7389113 | 9.5287E-09 | 5.2696E-07 |
| Itm2c | 2587.78853 | 2903.27892 | 0.89133308 | -0.1659635 | 2745.53373 | -0.1600738 | 0.02789851 | -5.7377181 | 9.5961E-09 | 5.2871E-07 |
| Amotl1 | 186.67206 | 141.131584 | 1.32268097 | 0.40346513 | 163.901822 | 0.4040807 | 0.07070115 | 5.71533437 | 1.0949E-08 | 6.0101E-07 |
| Cdc14b | 1586.2964 | 1772.63064 | 0.89488265 | -0.1602296 | 1679.46352 | -0.1551527 | 0.02717276 | -5.7098596 | 1.1307E-08 | 6.1838E-07 |
| Vgf | 174.63052 | 128.954486 | 1.35420275 | 0.43744376 | 151.792503 | 0.43770173 | 0.07668302 | 5.70793566 | 1.1435E-08 | 6.2311E-07 |
| Impdh2 | 506.236336 | 420.039076 | 1.20521248 | 0.26928752 | 463.137706 | 0.26772153 | 0.04695649 | 5.70148082 | 1.1877E-08 | 6.448E-07 |
| Nsg1 | 662.650948 | 759.181402 | 0.87284929 | -0.1961955 | 710.916175 | -0.1970255 | 0.0345729 | -5.6988427 | 1.2062E-08 | 6.5247E-07 |
| Luc7l3 | 1145.22246 | 1006.4503 | 1.13788278 | 0.18635194 | 1075.83638 | 0.19080064 | 0.03352796 | 5.69079249 | 1.2645E-08 | 6.815E-07 |
| Cfl1 | 2548.88065 | 2296.85614 | 1.10972586 | 0.15020332 | 2422.8684 | 0.15222319 | 0.02678033 | 5.68414134 | 1.3147E-08 | 7.0599E-07 |
| Ddr1 | 1360.25859 | 1506.86431 | 0.90270808 | -0.1476686 | 1433.56145 | -0.1473732 | 0.02593208 | -5.6830479 | 1.3232E-08 | 7.0796E-07 |
| Shmt1 | 170.677464 | 98.8150346 | 1.72724186 | 0.78847011 | 134.746249 | 0.79012655 | 0.13925602 | 5.67391291 | 1.3957E-08 | 7.441E-07 |
| Adamts4 | 124.220755 | 165.507511 | 0.75054451 | -0.4139905 | 144.864133 | -0.4139636 | 0.07297555 | -5.6726337 | 1.4062E-08 | 7.4699E-07 |
| Dchs1 | 18.8535682 | 6.07444021 | 3.10375401 | 1.63401422 | 12.4640041 | 1.60223645 | 0.28253486 | 5.67093365 | 1.4202E-08 | 7.5175E-07 |
| Stx16 | 1144.47059 | 996.175957 | 1.1488639 | 0.2002079 | 1070.32328 | 0.20060542 | 0.03540771 | 5.66558582 | 1.4652E-08 | 7.7282E-07 |
| Dcc | 54.1135242 | 82.8195554 | 0.65339066 | -0.6139823 | 68.4665397 | -0.6129719 | 0.10829683 | -5.6601094 | 1.5128E-08 | 7.9506E-07 |
| Rpl22l1 | 370.456442 | 306.353965 | 1.20924318 | 0.2741044 | 338.405203 | 0.27457624 | 0.0486324 | 5.64595264 | 1.6427E-08 | 8.5726E-07 |
| Kpna2 | 1874.7633 | 1561.05079 | 1.2009624 | 0.26419099 | 1717.90705 | 0.27037309 | 0.04788652 | 5.64612103 | 1.6411E-08 | 8.5726E-07 |
| Dmpk | 213.851965 | 163.816286 | 1.30543775 | 0.38453367 | 188.834126 | 0.38964916 | 0.06917243 | 5.63301256 | 1.7709E-08 | 9.1519E-07 |
| Ctnna2 | 639.654569 | 742.070396 | 0.86198637 | -0.214263 | 690.862482 | -0.2108797 | 0.03743631 | -5.633026 | 1.7707E-08 | 9.1519E-07 |
| Cds1 | 271.059382 | 332.728095 | 0.81465733 | -0.2957347 | 301.893739 | -0.2945294 | 0.0522875 | -5.6328828 | 1.7722E-08 | 9.1519E-07 |
| Itpr1 | 748.94116 | 853.455748 | 0.87753953 | -0.188464 | 801.198454 | -0.1923693 | 0.03420002 | -5.6248308 | 1.8569E-08 | 9.5559E-07 |
| Brinp2 | 316.728552 | 386.826928 | 0.81878621 | -0.2884413 | 351.77774 | -0.2871323 | 0.05106814 | -5.622533 | 1.8818E-08 | 9.6505E-07 |
| Eif4a1 | 5642.88715 | 5145.90188 | 1.09657885 | 0.13300955 | 5394.39452 | 0.13466285 | 0.02398333 | 5.61485228 | 1.9673E-08 | 1.0054E-06 |
| Gli3 | 20.0355848 | 5.30900121 | 3.77388967 | 1.91605225 | 12.6722929 | 1.81442216 | 0.32346955 | 5.60925187 | 2.032E-08 | 1.0349E-06 |
| Rps13 | 1536.15698 | 1392.91472 | 1.10283635 | 0.14121873 | 1464.53585 | 0.14174851 | 0.02537121 | 5.58698223 | 2.3105E-08 | 1.1727E-06 |
| Dzip1 | 149.428983 | 107.975931 | 1.38391012 | 0.46875024 | 128.702457 | 0.46540037 | 0.0833396 | 5.58438413 | 2.3453E-08 | 1.1863E-06 |
| Tmem45b | 403.292868 | 489.322619 | 0.82418603 | -0.2789581 | 446.307743 | -0.2831484 | 0.05074753 | -5.5795501 | 2.4114E-08 | 1.2156E-06 |
| Coro1b | 1351.56468 | 1489.29153 | 0.9075219 | -0.1399956 | 1420.42811 | -0.140076 | 0.02518022 | -5.5629388 | 2.6527E-08 | 1.3327E-06 |
| Rack1 | 3223.58409 | 2942.92629 | 1.09536692 | 0.13141421 | 3083.25519 | 0.132094 | 0.02376308 | 5.55879181 | 2.7165E-08 | 1.3602E-06 |
| Igfbp2 | 9.23510148 | 1.12510422 | 8.20821871 | 3.03706917 | 5.18010275 | 3.00912774 | 0.54241247 | 5.54767434 | 2.8949E-08 | 1.4446E-06 |
| Usp13 | 13.4072607 | 3.06574565 | 4.37324627 | 2.12870459 | 8.23650308 | 2.04550344 | 0.3688018 | 5.54634882 | 2.917E-08 | 1.4508E-06 |
| Pnmal2 | 1999.22305 | 2233.43914 | 0.89513209 | -0.1598275 | 2116.3311 | -0.1553122 | 0.0280089 | -5.5451 | 2.9379E-08 | 1.4563E-06 |
| Anxa11 | 1868.51893 | 2063.16864 | 0.90565497 | -0.1429666 | 1965.84378 | -0.1433655 | 0.02587497 | -5.5407027 | 3.0126E-08 | 1.4883E-06 |
| Litaf | 1021.51149 | 1162.96368 | 0.87836921 | -0.1871006 | 1092.23758 | -0.183659 | 0.03315873 | -5.5387817 | 3.0458E-08 | 1.4997E-06 |
| Fmn2 | 807.747535 | 912.532068 | 0.88517167 | -0.1759708 | 860.139801 | -0.1763411 | 0.03186031 | -5.5348221 | 3.1154E-08 | 1.5289E-06 |
| Slitrk3 | 10.8075826 | 1.3259448 | 8.15085408 | 3.02695124 | 6.0667636 | 2.94417631 | 0.53213066 | 5.53280714 | 3.1515E-08 | 1.5415E-06 |
| Osbpl3 | 435.638601 | 514.075844 | 0.84742088 | -0.2388494 | 474.857222 | -0.235806 | 0.04269317 | -5.5232728 | 3.3274E-08 | 1.6222E-06 |
| Plec | 976.584288 | 1095.05365 | 0.8918141 | -0.1651851 | 1035.81897 | -0.1662647 | 0.03015632 | -5.5134283 | 3.5191E-08 | 1.7101E-06 |
| 2700081O15 | 80.2465909 | 48.7749776 | 1.64524096 | 0.71829889 | 64.5107841 | 0.74477908 | 0.13543492 | 5.49916599 | 3.8159E-08 | 1.8482E-06 |
| Mgst3 | 162.685216 | 208.643603 | 0.77972779 | -0.3589575 | 185.664409 | -0.3602697 | 0.06554176 | -5.4967969 | 3.8675E-08 | 1.8671E-06 |
| Tm9sf3 | 1403.96353 | 1260.20213 | 1.11407804 | 0.1558503 | 1332.08283 | 0.15558956 | 0.02836018 | 5.48619746 | 4.1068E-08 | 1.9706E-06 |
| Lrfn4 | 1136.78302 | 1286.89571 | 0.88335287 | -0.1789382 | 1211.83936 | -0.1802552 | 0.03285657 | -5.4861231 | 4.1085E-08 | 1.9706E-06 |
| Rpl29 | 1529.71499 | 1394.45741 | 1.09699657 | 0.13355902 | 1462.0862 | 0.13430487 | 0.02464214 | 5.45021188 | 5.031E-08 | 2.4012E-06 |
| Bbs1 | 551.283541 | 638.95746 | 0.86278598 | -0.2129254 | 595.1205 | -0.2122786 | 0.03895059 | -5.4499445 | 5.0386E-08 | 2.4012E-06 |
| Abcc8 | 80.2361794 | 113.247883 | 0.70850048 | -0.4971593 | 96.7420311 | -0.4982192 | 0.0914567 | -5.4475966 | 5.1055E-08 | 2.4253E-06 |
| Lamc1 | 802.103427 | 903.780012 | 0.88749852 | -0.1721834 | 852.941719 | -0.1726897 | 0.03174792 | -5.4394022 | 5.346E-08 | 2.5314E-06 |
| Rabep1 | 1416.34952 | 1282.31952 | 1.10452153 | 0.14342154 | 1349.33452 | 0.14292437 | 0.02628473 | 5.43754386 | 5.402E-08 | 2.5498E-06 |
| Cox6a2 | 108.670473 | 76.8310985 | 1.41440738 | 0.50019771 | 92.7507856 | 0.50543879 | 0.0930387 | 5.43256498 | 5.555E-08 | 2.6137E-06 |
| Cacna1h | 2737.66184 | 3091.56461 | 0.88552632 | -0.1753929 | 2914.61322 | -0.1700827 | 0.03131803 | -5.4308235 | 5.6095E-08 | 2.6309E-06 |
| Rufy3 | 1725.94384 | 1920.53429 | 0.898679 | -0.1541222 | 1823.23906 | -0.1541397 | 0.02838871 | -5.4296147 | 5.6476E-08 | 2.6405E-06 |
| Arhgap19 | 371.838041 | 303.411079 | 1.22552559 | 0.29340061 | 337.62456 | 0.29434092 | 0.05422432 | 5.42820822 | 5.6923E-08 | 2.653E-06 |
| Oprd1 | 170.038497 | 220.739493 | 0.77031298 | -0.3764834 | 195.388995 | -0.3911516 | 0.07223306 | -5.4151329 | 6.1243E-08 | 2.8454E-06 |
| Dgkz | 1816.83247 | 1983.54979 | 0.91595002 | -0.1266592 | 1900.19113 | -0.1266276 | 0.02339893 | -5.4116837 | 6.2435E-08 | 2.8917E-06 |
| Slc18a3 | 17.3026541 | 34.862809 | 0.49630694 | -1.0106955 | 26.0827314 | -0.9962952 | 0.1842114 | -5.4084343 | 6.3578E-08 | 2.9355E-06 |
| Knstrn | 1270.94222 | 1132.48582 | 1.12225884 | 0.16640546 | 1201.71402 | 0.16940475 | 0.03135007 | 5.4036481 | 6.5299E-08 | 3.0056E-06 |
| Ralyl | 19.3488 | 7.30591027 | 2.64837636 | 1.40510816 | 13.327355 | 1.39228409 | 0.25773385 | 5.4020227 | 6.5894E-08 | 3.0142E-06 |
| Rps4x | 3032.40454 | 2812.77851 | 1.07808152 | 0.10846628 | 2922.59153 | 0.10852271 | 0.02008786 | 5.40240224 | 6.5754E-08 | 3.0142E-06 |
| Serpinb12 | 183.799767 | 108.672507 | 1.69131799 | 0.75814793 | 146.236137 | 0.74430273 | 0.13780373 | 5.40117997 | 6.6204E-08 | 3.0191E-06 |
| H2afz | 4804.30542 | 4466.16865 | 1.0757107 | 0.10529014 | 4635.23704 | 0.10712486 | 0.01984113 | 5.39913135 | 6.6964E-08 | 3.0444E-06 |
| Srsf5 | 1691.16095 | 1546.37795 | 1.09362718 | 0.129121 | 1618.76945 | 0.1292597 | 0.02397616 | 5.39117525 | 6.9998E-08 | 3.1726E-06 |
| Tmem56 | 615.523113 | 530.983125 | 1.15921408 | 0.21314702 | 573.253119 | 0.21418214 | 0.0397577 | 5.38718704 | 7.1569E-08 | 3.2339E-06 |
| Arhgef40 | 120.795935 | 159.640028 | 0.75667699 | -0.4022505 | 140.217982 | -0.401131 | 0.07466784 | -5.3722058 | 7.7779E-08 | 3.5038E-06 |
| Racgap1 | 3155.93108 | 2883.42972 | 1.09450598 | 0.13027984 | 3019.6804 | 0.13317733 | 0.02479555 | 5.37101625 | 7.8294E-08 | 3.5164E-06 |
| Robo2 | 254.214061 | 321.530472 | 0.79063754 | -0.3389116 | 287.872266 | -0.321919 | 0.05994378 | -5.3703493 | 7.8584E-08 | 3.5187E-06 |
| Rpl9 | 2223.85168 | 2046.90531 | 1.0864458 | 0.11961621 | 2135.3785 | 0.12106012 | 0.02262176 | 5.35149088 | 8.7233E-08 | 3.8825E-06 |
| Igsf9 | 2057.96671 | 2248.51199 | 0.91525717 | -0.1277509 | 2153.23935 | -0.1297776 | 0.02425028 | -5.3515927 | 8.7183E-08 | 3.8825E-06 |
| Ifitm3 | 121.927429 | 87.5804666 | 1.39217606 | 0.47734168 | 104.753948 | 0.47543921 | 0.08897353 | 5.34360297 | 9.1117E-08 | 4.0433E-06 |

| | | | | | | | | | | |
|---|---|---|---|---|---|---|---|---|---|---|
| Nkx2-1 | 2539.93943 | 2347.68072 | 1.08189304 | 0.11355788 | 2443.81007 | 0.11355032 | 0.02127876 | 5.33632125 | 9.4851E-08 | 4.1964E-06 |
| Cep170 | 1416.87904 | 1558.06095 | 0.90938615 | -0.1370351 | 1487.47 | -0.1370797 | 0.02569754 | -5.3343496 | 9.5888E-08 | 4.2296E-06 |
| Itga4 | 522.264825 | 435.03168 | 1.20052136 | 0.26366107 | 478.648252 | 0.25847739 | 0.04858387 | 5.32023036 | 1.0364E-07 | 4.5578E-06 |
| Pcdhb11 | 40.1993417 | 63.1663932 | 0.63640394 | -0.6519853 | 51.6828674 | -0.6529506 | 0.12292675 | -5.311705 | 1.086E-07 | 4.7622E-06 |
| Sowahc | 600.9127 | 689.35591 | 0.87170167 | -0.1980936 | 645.134305 | -0.195206 | 0.036762 | -5.3099949 | 1.0963E-07 | 4.7929E-06 |
| Map4 | 2392.26581 | 2641.24711 | 0.90573343 | -0.1428416 | 2516.75646 | -0.1421412 | 0.02677484 | -5.3087591 | 1.1037E-07 | 4.8113E-06 |
| Abca7 | 943.812118 | 1057.25347 | 0.89270184 | -0.1637497 | 1000.53279 | -0.164554 | 0.0310187 | -5.3049928 | 1.1268E-07 | 4.8973E-06 |
| Fev | 75.6194643 | 108.162774 | 0.69912652 | -0.5163745 | 91.8911192 | -0.5162768 | 0.09737671 | -5.301851 | 1.1463E-07 | 4.9678E-06 |
| Kcnk2 | 889.879349 | 998.966603 | 0.8907999 | -0.1668267 | 944.422976 | -0.1670793 | 0.03152407 | -5.3000545 | 1.1577E-07 | 5.0023E-06 |
| Hnrnpa2b1 | 7490.02829 | 7074.88573 | 1.05867834 | 0.08226432 | 7282.45701 | 0.08467845 | 0.01600458 | 5.29088935 | 1.2172E-07 | 5.2291E-06 |
| Grxcr2 | 410.509905 | 485.833436 | 0.84496017 | -0.2430448 | 448.171671 | -0.2455694 | 0.04641239 | -5.2910324 | 1.2163E-07 | 5.2291E-06 |
| Hectd2 | 185.145431 | 142.66841 | 1.29773249 | 0.37599302 | 163.90692 | 0.37501634 | 0.07088957 | 5.29014849 | 1.2222E-07 | 5.2352E-06 |
| Sprr1a | 29.1373878 | 13.1423588 | 2.21705922 | 1.1486473 | 21.1398732 | 1.16845628 | 0.22096089 | 5.28806827 | 1.2361E-07 | 5.2798E-06 |
| Birc5 | 484.024361 | 405.76454 | 1.19287003 | 0.25443687 | 444.89445 | 0.26869066 | 0.05087487 | 5.28140291 | 1.282E-07 | 5.4599E-06 |
| Pnn | 1142.06349 | 1021.44898 | 1.11808177 | 0.1610257 | 1081.75623 | 0.16474073 | 0.0312398 | 5.27342435 | 1.339E-07 | 5.6864E-06 |
| Grk2 | 3767.9106 | 4016.80664 | 0.93803634 | -0.0922843 | 3892.35862 | -0.092644 | 0.01759063 | -5.2666647 | 1.3892E-07 | 5.8829E-06 |
| Sema6d | 5.48648707 | 0.11790705 | 46.532309 | 5.54016087 | 2.80219696 | 4.4178131 | 0.83912343 | 5.26479529 | 1.4035E-07 | 5.9261E-06 |
| Hnrnph1 | 4891.96133 | 4481.41294 | 1.09161137 | 0.12645933 | 4686.68713 | 0.13277269 | 0.0252348 | 5.26149259 | 1.4289E-07 | 6.0164E-06 |
| Atp2b4 | 131.614698 | 172.882096 | 0.76129744 | -0.3934679 | 152.248397 | -0.3908795 | 0.07430496 | -5.260477 | 1.4368E-07 | 6.0326E-06 |
| Arhgap11a | 1615.39092 | 1435.1802 | 1.12556662 | 0.17065144 | 1525.28556 | 0.17446107 | 0.03318599 | 5.25707024 | 1.4637E-07 | 6.128E-06 |
| Foxq1 | 29.148075 | 49.6627881 | 0.58691983 | -0.7687647 | 39.4054315 | -0.7757425 | 0.147608 | -5.2554233 | 1.4768E-07 | 6.1657E-06 |
| Rpl13 | 2233.25063 | 2066.3788 | 1.08075568 | 0.11204042 | 2149.81472 | 0.11159047 | 0.0212826 | 5.2432726 | 1.5775E-07 | 6.5676E-06 |
| Map1b | 2526.26603 | 2820.62791 | 0.89563959 | -0.1590098 | 2673.44697 | -0.1653332 | 0.03155236 | -5.2399627 | 1.6061E-07 | 6.6678E-06 |
| 5330417C22 | 1801.04028 | 1976.66142 | 0.91115264 | -0.1342353 | 1888.85085 | -0.1320885 | 0.02529041 | -5.2228669 | 1.7617E-07 | 7.2935E-06 |
| Gstm5 | 530.785959 | 610.274248 | 0.86974989 | -0.2013275 | 570.530104 | -0.2012341 | 0.03853717 | -5.2218192 | 1.7717E-07 | 7.3145E-06 |
| Txndc16 | 1578.0058 | 1418.54145 | 1.1124143 | 0.1536942 | 1498.27362 | 0.16004643 | 0.0306702 | 5.21830429 | 1.8057E-07 | 7.4339E-06 |
| Srebf1 | 1779.62907 | 1950.77902 | 0.91226584 | -0.1324738 | 1865.20405 | -0.1287255 | 0.02467561 | -5.2167118 | 1.8213E-07 | 7.4773E-06 |
| Rpl38 | 638.277341 | 554.233096 | 1.15164061 | 0.20369057 | 596.255218 | 0.20490669 | 0.03928561 | 5.21582031 | 1.8301E-07 | 7.4926E-06 |
| Map4k4 | 3169.31559 | 3386.79932 | 0.93578488 | -0.0957512 | 3278.05745 | -0.095822 | 0.01839599 | -5.2088553 | 1.9001E-07 | 7.7579E-06 |
| Atf6 | 1591.34751 | 1747.58079 | 0.91060025 | -0.1351102 | 1669.46415 | -0.1351899 | 0.02599451 | -5.2007089 | 1.9853E-07 | 8.0835E-06 |
| Ldlr | 1097.06992 | 1218.05913 | 0.90067049 | -0.1509287 | 1157.56452 | -0.1560586 | 0.03004124 | -5.1948144 | 2.0492E-07 | 8.321E-06 |
| Aip | 1684.81475 | 1835.21716 | 0.91804653 | -0.1233608 | 1760.01596 | -0.1235004 | 0.02379413 | -5.1903729 | 2.0987E-07 | 8.4987E-06 |
| Cdh9 | 185.212092 | 236.983064 | 0.78154147 | -0.3556057 | 211.097578 | -0.3495966 | 0.06740225 | -5.1867203 | 2.1403E-07 | 8.6434E-06 |
| Inppl1 | 3481.74019 | 3713.75523 | 0.93752549 | -0.0930702 | 3597.74771 | -0.093173 | 0.0179685 | -5.1853498 | 2.1561E-07 | 8.6836E-06 |
| Rps28 | 966.944345 | 865.696022 | 1.11695598 | 0.15957232 | 916.320184 | 0.15955486 | 0.03080216 | 5.17998858 | 2.219E-07 | 8.9127E-06 |
| Lynx1 | 40.3444929 | 62.4851368 | 0.64566543 | -0.6311413 | 51.4148147 | -0.6309834 | 0.12195144 | -5.1740541 | 2.2907E-07 | 9.1757E-06 |
| Tmem132a | 274.937482 | 221.047994 | 1.24379089 | 0.31474396 | 247.992738 | 0.31603328 | 0.0611666 | 5.16676229 | 2.3818E-07 | 9.5152E-06 |
| Tmcc3 | 9.494637 | 1.91454369 | 4.95921668 | 2.31011226 | 5.70459025 | 2.2832652 | 0.44235626 | 5.16159807 | 2.4485E-07 | 9.7552E-06 |
| Cpne5 | 364.118504 | 428.774549 | 0.84920736 | -0.2358112 | 396.446526 | -0.2367899 | 0.04591135 | -5.1575462 | 2.5021E-07 | 9.9419E-06 |
| Rps18 | 1759.5146 | 1606.50998 | 1.09524038 | 0.13124755 | 1683.01229 | 0.13133399 | 0.02549242 | 5.1518834 | 2.5788E-07 | 1.0203E-05 |
| Kcnq5 | 320.698619 | 380.220372 | 0.84345459 | -0.2456177 | 350.459495 | -0.246877 | 0.04792151 | -5.1516934 | 2.5814E-07 | 1.0203E-05 |
| Arhgef15 | 234.127258 | 286.369052 | 0.81757179 | -0.2905827 | 260.248155 | -0.2875425 | 0.05583301 | -5.150044 | 2.6043E-07 | 1.0265E-05 |
| Trim66 | 146.358487 | 189.133557 | 0.7738367 | -0.3698989 | 167.746022 | -0.3676365 | 0.07140872 | -5.148342 | 2.628E-07 | 1.0331E-05 |
| Tspan6 | 4.37725437 | 0.11822832 | 37.0237377 | 5.21037864 | 2.24774125 | 4.35303435 | 0.84822866 | 5.13191139 | 2.8681E-07 | 1.1198E-05 |
| Psmc6 | 1172.00514 | 1059.61695 | 1.10606493 | 0.14543608 | 1115.81105 | 0.14646068 | 0.02854026 | 5.13172269 | 2.871E-07 | 1.1198E-05 |
| Kif5a | 852.666568 | 950.858931 | 0.89673299 | -0.1572496 | 901.762749 | -0.1572251 | 0.03063699 | -5.1318705 | 2.8688E-07 | 1.1198E-05 |
| Frmd3 | 4.7086716 | 0.11822832 | 39.8269342 | 5.31567252 | 2.41344986 | 4.30921908 | 0.84080225 | 5.12512793 | 2.9734E-07 | 1.1566E-05 |
| Zbtb20 | 94.8675433 | 127.71004 | 0.74283544 | -0.4288855 | 111.288791 | -0.4302207 | 0.08411936 | -5.114408 | 3.1473E-07 | 1.2211E-05 |
| Aff3 | 666.292879 | 766.654143 | 0.86909187 | -0.2024194 | 716.473511 | -0.2001259 | 0.03913364 | -5.1139083 | 3.1556E-07 | 1.2211E-05 |
| Maged1 | 2482.44297 | 2688.91979 | 0.92321198 | -0.1152661 | 2585.68138 | -0.1150522 | 0.02252591 | -5.1075477 | 3.2637E-07 | 1.2597E-05 |
| Syp | 1262.5553 | 1386.25631 | 0.91076614 | -0.1348474 | 1324.4058 | -0.1341019 | 0.02629086 | -5.1007044 | 3.3839E-07 | 1.3027E-05 |
| Casr | 1034.25572 | 1204.5587 | 0.85861795 | -0.2199118 | 1119.40721 | -0.2023079 | 0.0397174 | -5.0936852 | 3.5117E-07 | 1.3449E-05 |
| Enho | 451.783301 | 523.958021 | 0.86225095 | -0.2138203 | 487.870661 | -0.2139774 | 0.04200837 | -5.0936847 | 3.5117E-07 | 1.3449E-05 |
| St8sia1 | 255.41441 | 314.012388 | 0.8133896 | -0.2979816 | 284.713399 | -0.2878449 | 0.05653578 | -5.091375 | 3.5548E-07 | 1.3579E-05 |
| Nes | 14.8691236 | 4.79037573 | 3.1039577 | 1.6341089 | 9.82974958 | 1.64190755 | 0.3228532 | 5.0856165 | 3.6643E-07 | 1.3961E-05 |
| Slc12a2 | 1045.32949 | 1161.61965 | 0.89988965 | -0.15218 | 1103.47457 | -0.152342 | 0.02997626 | -5.0820889 | 3.7331E-07 | 1.4187E-05 |
| Rtn1 | 704.028018 | 830.103667 | 0.8481206 | -0.2376587 | 767.065842 | -0.238565 | 0.04695195 | -5.0810449 | 3.7536E-07 | 1.4228E-05 |
| Adipor1 | 2308.96859 | 2517.52435 | 0.91715839 | -0.1247572 | 2413.24647 | -0.1243268 | 0.02449819 | -5.0749398 | 3.8762E-07 | 1.4655E-05 |
| Esr1 | 4.78559775 | 14.6813392 | 0.32596466 | -1.6172125 | 9.73346835 | -1.6145716 | 0.31819104 | -5.0742209 | 3.8909E-07 | 1.4673E-05 |
| Usmg5 | 338.168772 | 281.937702 | 1.19944502 | 0.26236703 | 310.053237 | 0.26698921 | 0.05273494 | 5.06285265 | 4.1303E-07 | 1.5537E-05 |
| Rpl35a | 1260.01523 | 1133.47923 | 1.11163504 | 0.15268321 | 1196.74723 | 0.15591194 | 0.03082958 | 5.05721859 | 4.2542E-07 | 1.5962E-05 |
| Fau | 650.279801 | 563.62914 | 1.15373701 | 0.20631441 | 606.954471 | 0.206726 | 0.04089406 | 5.05515915 | 4.3003E-07 | 1.6095E-05 |
| Btg2 | 1179.24045 | 1297.1594 | 0.90909448 | -0.1374979 | 1238.19992 | -0.1368755 | 0.02707971 | -5.0545409 | 4.3143E-07 | 1.6106E-05 |
| Dsg2 | 1353.41436 | 1482.38907 | 0.91299537 | -0.1313206 | 1417.90171 | -0.1315248 | 0.02607262 | -5.0445548 | 4.5458E-07 | 1.6928E-05 |
| Kif11 | 1562.01355 | 1288.10556 | 1.21264406 | 0.27815614 | 1425.05955 | 0.28310739 | 0.05615761 | 5.04130018 | 4.6238E-07 | 1.7175E-05 |
| Ptms | 3880.0783 | 4135.8774 | 0.93815119 | -0.0921077 | 4007.97785 | -0.0920404 | 0.01827576 | -5.0362027 | 4.7486E-07 | 1.7595E-05 |
| Dsp | 3760.0439 | 4069.19407 | 0.92402668 | -0.1139936 | 3914.61899 | -0.1217245 | 0.0242129 | -5.02726 | 4.9754E-07 | 1.8389E-05 |

| | | | | | | | | | | |
|---|---|---|---|---|---|---|---|---|---|---|
| Fhdc1 | 526.130369 | 455.149097 | 1.15595169 | 0.20908111 | 490.639733 | 0.20850249 | 0.0414892 | 5.02546469 | 5.0221E-07 | 1.8516E-05 |
| Pam | 10307.0503 | 11193.1494 | 0.92083559 | -0.1189845 | 10750.0999 | -0.1238468 | 0.02464823 | -5.024572 | 5.0456E-07 | 1.8525E-05 |
| Tox2 | 108.484341 | 146.630402 | 0.73984889 | -0.4346975 | 127.557371 | -0.4441727 | 0.08840279 | -5.0244197 | 5.0496E-07 | 1.8525E-05 |
| Ptp4a3 | 1022.07953 | 1131.3191 | 0.90344054 | -0.1464984 | 1076.69932 | -0.145953 | 0.02905183 | -5.0238829 | 5.0637E-07 | 1.8531E-05 |
| Plekha1 | 1507.23327 | 1639.15409 | 0.91951896 | -0.1210488 | 1573.19368 | -0.1209115 | 0.02409516 | -5.018082 | 5.219E-07 | 1.9052E-05 |
| Grm1 | 4.57885902 | 14.1852234 | 0.32279076 | -1.6313288 | 9.38204111 | -1.6469113 | 0.32853703 | -5.0128637 | 5.3626E-07 | 1.9528E-05 |
| Plcl1 | 9.31234195 | 1.47983202 | 6.29283716 | 2.65371061 | 5.39608689 | 2.51219332 | 0.50226569 | 5.00172195 | 5.6821E-07 | 2.0641E-05 |
| Mcam | 90.6644461 | 62.7050883 | 1.44588659 | 0.53195439 | 76.6847671 | 0.53332347 | 0.10671927 | 4.99744326 | 5.8095E-07 | 2.1052E-05 |
| Galnt18 | 47.5773353 | 28.0485059 | 1.69625204 | 0.76235055 | 37.8129205 | 0.77105955 | 0.15438051 | 4.9945393 | 5.8976E-07 | 2.1216E-05 |
| Kremen1 | 972.08417 | 1115.43509 | 0.8714843 | -0.1984534 | 1043.75963 | -0.1923311 | 0.03850381 | -4.995118 | 5.88E-07 | 2.1216E-05 |
| 1810034E14I | 139.252132 | 178.924259 | 0.77827418 | -0.3616496 | 159.088195 | -0.3574117 | 0.07155808 | -4.9947083 | 5.8925E-07 | 2.1216E-05 |
| Cavin2 | 121.011876 | 165.313922 | 0.73201261 | -0.4500596 | 143.162899 | -0.4674426 | 0.09360755 | -4.9936426 | 5.9251E-07 | 2.1263E-05 |
| Calm1 | 9247.84511 | 8805.60049 | 1.05022311 | 0.07069584 | 9026.7228 | 0.07192221 | 0.01442921 | 4.98448672 | 6.2127E-07 | 2.2241E-05 |
| Prc1 | 2719.1667 | 2524.30761 | 1.07719308 | 0.10727687 | 2621.73716 | 0.10842696 | 0.02176553 | 4.98159081 | 6.3064E-07 | 2.2522E-05 |
| Bok | 403.765646 | 468.748527 | 0.86136942 | -0.215296 | 436.257086 | -0.2171861 | 0.04362175 | -4.9788495 | 6.3963E-07 | 2.2788E-05 |
| Meis1 | 6.19222152 | 0.33496936 | 18.4859342 | 4.20835605 | 3.26359534 | 4.01353857 | 0.80688265 | 4.97412921 | 6.5542E-07 | 2.3295E-05 |
| Spats2l | 20.3068086 | 36.3314829 | 0.55893145 | -0.8392567 | 28.3191457 | -0.8724128 | 0.17560706 | -4.9679822 | 6.7653E-07 | 2.3988E-05 |
| Snca | 241.686089 | 291.548819 | 0.82897297 | -0.270603 | 266.617454 | -0.2691895 | 0.05422359 | -4.9644354 | 6.8901E-07 | 2.4372E-05 |
| Prps1 | 524.666366 | 451.376831 | 1.16236885 | 0.21706794 | 488.021598 | 0.21835618 | 0.04402397 | 4.95993882 | 7.0515E-07 | 2.4884E-05 |
| Slf2 | 509.829661 | 437.120031 | 1.1663379 | 0.22198582 | 473.474846 | 0.21961628 | 0.04435024 | 4.95186209 | 7.3507E-07 | 2.5877E-05 |
| Tcta | 6.090867 | 0.47179335 | 12.9100316 | 3.69042063 | 3.28133007 | 3.6947204 | 0.74779986 | 4.94078773 | 7.7808E-07 | 2.7327E-05 |
| Efr3b | 1024.56494 | 1132.83385 | 0.90442649 | -0.1449248 | 1078.6994 | -0.1448076 | 0.02932735 | -4.9376303 | 7.9078E-07 | 2.7707E-05 |
| Pim1 | 414.557552 | 481.84675 | 0.86035145 | -0.217002 | 448.202151 | -0.2193215 | 0.04452621 | -4.9256716 | 8.4071E-07 | 2.9387E-05 |
| Skap2 | 117.883808 | 86.6853659 | 1.35990437 | 0.4435052 | 102.284587 | 0.44696236 | 0.09087606 | 4.91837322 | 8.7266E-07 | 3.0432E-05 |
| Elf4 | 295.873484 | 352.622428 | 0.83906598 | -0.2531438 | 324.247956 | -0.2526908 | 0.05140557 | -4.9156305 | 8.8497E-07 | 3.0789E-05 |
| Rps25 | 1435.47147 | 1303.88282 | 1.10092061 | 0.13871043 | 1369.67714 | 0.13760186 | 0.02801133 | 4.91236508 | 8.9984E-07 | 3.1233E-05 |
| Clvs1 | 223.068706 | 278.378055 | 0.8013157 | -0.3195574 | 250.72338 | -0.3086322 | 0.06284679 | -4.9108659 | 9.0675E-07 | 3.1399E-05 |
| Pcp4 | 11.1529848 | 23.7961934 | 0.46868777 | -1.093301 | 17.474589 | -1.1086931 | 0.22607052 | -4.9041912 | 9.3813E-07 | 3.241E-05 |
| Pex5l | 1079.22839 | 1191.10284 | 0.9060749 | -0.1422978 | 1135.16561 | -0.1429018 | 0.02915879 | -4.9008134 | 9.5441E-07 | 3.2896E-05 |
| Atp1a3 | 344.000555 | 406.493909 | 0.84626251 | -0.2408228 | 375.247232 | -0.2394494 | 0.04889836 | -4.8968801 | 9.737E-07 | 3.3483E-05 |
| Setd7 | 1625.43652 | 1494.92033 | 1.08730645 | 0.12075861 | 1560.17843 | 0.12080795 | 0.02468774 | 4.89343905 | 9.9089E-07 | 3.3995E-05 |
| Atp6v0b | 2905.6098 | 3095.95527 | 0.93851802 | -0.0915437 | 3000.78254 | -0.0913003 | 0.01866434 | -4.8917004 | 9.9969E-07 | 3.4218E-05 |
| Rpl15 | 2887.6777 | 2702.81852 | 1.06839497 | 0.09544509 | 2795.24811 | 0.09596351 | 0.01964848 | 4.88401605 | 1.0395E-06 | 3.5498E-05 |
| Eif2s3y | 482.65145 | 417.408521 | 1.15630474 | 0.20952166 | 450.029985 | 0.21141329 | 0.04331172 | 4.8812024 | 1.0544E-06 | 3.5925E-05 |
| Glul | 1003.43327 | 1118.49479 | 0.89712825 | -0.1566139 | 1060.96403 | -0.1568985 | 0.03220901 | -4.8712607 | 1.1089E-06 | 3.7694E-05 |
| Esyt3 | 1011.31169 | 1119.0601 | 0.90371526 | -0.1460598 | 1065.18589 | -0.1456616 | 0.02991359 | -4.8694121 | 1.1193E-06 | 3.7962E-05 |
| Ascl1 | 20585.5988 | 21286.5746 | 0.96706958 | -0.0483084 | 20936.0867 | -0.0482379 | 0.00991447 | -4.8654029 | 1.1422E-06 | 3.8651E-05 |
| Tanc2 | 3095.86702 | 2791.29835 | 1.10911362 | 0.14940717 | 2943.58269 | 0.13491142 | 0.02778425 | 4.85567908 | 1.1997E-06 | 4.0504E-05 |
| Vwa5a | 113.448299 | 150.453324 | 0.75404316 | -0.407281 | 131.950811 | -0.3962925 | 0.08163216 | -4.8546124 | 1.2062E-06 | 4.0631E-05 |
| Ppm1e | 587.569539 | 664.437781 | 0.88431085 | -0.1773745 | 626.00366 | -0.1781627 | 0.036726 | -4.8511326 | 1.2276E-06 | 4.1256E-05 |
| Rps23 | 913.045762 | 820.495409 | 1.11279814 | 0.15419191 | 866.770585 | 0.15511085 | 0.03198696 | 4.84919084 | 1.2397E-06 | 4.1568E-05 |
| Hpse | 105.797318 | 140.676946 | 0.75205868 | -0.4110829 | 123.237131 | -0.408833 | 0.08432928 | -4.8480559 | 1.2468E-06 | 4.1712E-05 |
| Neo1 | 1290.0398 | 1408.15767 | 0.91611886 | -0.1263933 | 1349.09874 | -0.1259046 | 0.02602444 | -4.8379356 | 1.3119E-06 | 4.3794E-05 |
| Cenpe | 1358.10514 | 1201.21809 | 1.13060663 | 0.17709706 | 1279.66161 | 0.17581164 | 0.0363443 | 4.83739245 | 1.3155E-06 | 4.3815E-05 |
| Ctnna1 | 3194.08078 | 3404.64777 | 0.93815308 | -0.0921048 | 3299.36427 | -0.0920214 | 0.01903602 | -4.8340664 | 1.3377E-06 | 4.4399E-05 |
| Dcx | 18.5792321 | 33.7633864 | 0.55027751 | -0.8617687 | 26.1713091 | -0.8722399 | 0.18044352 | -4.8338664 | 1.3391E-06 | 4.4399E-05 |
| Dis3l2 | 311.937523 | 375.430941 | 0.83087857 | -0.2672904 | 343.684231 | -0.2645407 | 0.0547849 | -4.8287153 | 1.3742E-06 | 4.5461E-05 |
| Acot12 | 0.13075673 | 4.25339671 | 0.03074172 | -5.0236583 | 2.19207662 | -3.9966038 | 0.82847553 | -4.8240456 | 1.4068E-06 | 4.6435E-05 |
| Has3 | 80.2961738 | 55.4760291 | 1.44740305 | 0.53346672 | 67.8861014 | 0.54258392 | 0.11248916 | 4.82343301 | 1.4111E-06 | 4.6475E-05 |
| Rpl17 | 1593.23405 | 1463.32664 | 1.0887754 | 0.12270638 | 1528.28034 | 0.1235581 | 0.02564217 | 4.81854987 | 1.4461E-06 | 4.7441E-05 |
| Cacnb2 | 48.7054539 | 72.3652768 | 0.67305006 | -0.5712143 | 60.5353652 | -0.5706823 | 0.11843703 | -4.8184449 | 1.4468E-06 | 4.7441E-05 |
| Prkd3 | 15.55802 | 31.1045823 | 0.50018418 | -0.9994687 | 23.3313011 | -0.9552145 | 0.19844191 | -4.8135725 | 1.4826E-06 | 4.8505E-05 |
| Igsf3 | 419.443079 | 490.049854 | 0.8559192 | -0.2244535 | 454.746466 | -0.2222632 | 0.04618355 | -4.8126058 | 1.4897E-06 | 4.8633E-05 |
| Atp5h | 1144.24935 | 1037.48252 | 1.10290953 | 0.14131445 | 1090.86594 | 0.14192854 | 0.02950923 | 4.8096328 | 1.5121E-06 | 4.9254E-05 |
| Sept4 | 83.0589096 | 113.674193 | 0.73067516 | -0.4526979 | 98.3665512 | -0.4434238 | 0.09221578 | -4.8085453 | 1.5203E-06 | 4.9414E-05 |
| Tsc22d1 | 2449.16661 | 2620.23608 | 0.93471219 | -0.0974059 | 2534.70134 | -0.0974833 | 0.02036691 | -4.786358 | 1.6984E-06 | 5.5079E-05 |
| Rpl14 | 1824.28281 | 1690.57366 | 1.079091 | 0.10981653 | 1757.42824 | 0.1098892 | 0.02298669 | 4.78055825 | 1.7481E-06 | 5.6569E-05 |
| Cdc42ep3 | 77.2287358 | 106.056663 | 0.72818373 | -0.4576256 | 91.642699 | -0.4586521 | 0.09600681 | -4.7772867 | 1.7768E-06 | 5.7371E-05 |
| Rec8 | 275.874586 | 225.973639 | 1.2208264 | 0.28785807 | 250.924112 | 0.28593198 | 0.0598718 | 4.77573731 | 1.7905E-06 | 5.7689E-05 |
| Hint1 | 1477.51158 | 1357.864 | 1.08811456 | 0.12183045 | 1417.68779 | 0.12306242 | 0.02577536 | 4.77441999 | 1.8023E-06 | 5.7942E-05 |
| Rab3gap2 | 1694.88434 | 1846.066 | 0.91810604 | -0.1232673 | 1770.47517 | -0.1223279 | 0.02566551 | -4.7662362 | 1.877E-06 | 6.0214E-05 |
| Igsf8 | 1261.93981 | 1383.46238 | 0.91216055 | -0.1326403 | 1322.7011 | -0.1319464 | 0.02772196 | -4.7596334 | 1.9394E-06 | 6.2083E-05 |
| Nfe2l2 | 322.190704 | 271.386077 | 1.18720425 | 0.24756816 | 296.788391 | 0.24783632 | 0.0520786 | 4.75888946 | 1.9466E-06 | 6.2178E-05 |
| Kcnh3 | 399.38562 | 465.038473 | 0.85882275 | -0.2195677 | 432.212047 | -0.2199465 | 0.04634678 | -4.7456696 | 2.0782E-06 | 6.6238E-05 |
| Arrdc1 | 151.768349 | 190.96492 | 0.79474465 | -0.3314367 | 171.366635 | -0.3311441 | 0.06978697 | -4.7450705 | 2.0843E-06 | 6.6292E-05 |
| Rpl13a | 2488.32738 | 2310.0878 | 1.07715706 | 0.10722862 | 2399.20759 | 0.10920564 | 0.0230168 | 4.74460519 | 2.0891E-06 | 6.6302E-05 |
| Nkd1 | 307.353157 | 361.43343 | 0.8503728 | -0.2338326 | 334.393293 | -0.2338631 | 0.04929562 | -4.744095 | 2.0944E-06 | 6.6327E-05 |

| | | | | | | | | | | |
|---|---|---|---|---|---|---|---|---|---|---|
| Gpc6 | 237.077953 | 194.721279 | 1.21752463 | 0.28395096 | 215.899616 | 0.28391908 | 0.05988042 | 4.74143414 | 2.1221E-06 | 6.7061E-05 |
| Arhgef19 | 452.823137 | 518.998909 | 0.87249343 | -0.1967838 | 485.911023 | -0.1973214 | 0.04162782 | -4.7401336 | 2.1358E-06 | 6.735E-05 |
| Efhd2 | 518.427396 | 601.296725 | 0.8621823 | -0.2139351 | 559.86206 | -0.2126585 | 0.04488249 | -4.7381176 | 2.1571E-06 | 6.7878E-05 |
| Aldoc | 85.3536221 | 121.397383 | 0.70309277 | -0.508213 | 103.375502 | -0.464956 | 0.09826494 | -4.7316574 | 2.2269E-06 | 6.9927E-05 |
| Mprip | 1782.00781 | 1645.82796 | 1.08274246 | 0.11469012 | 1713.91789 | 0.11483474 | 0.02429023 | 4.72761094 | 2.2718E-06 | 7.1184E-05 |
| Hp1bp3 | 2242.01902 | 2400.61638 | 0.93393474 | -0.0986064 | 2321.3177 | -0.0987967 | 0.02090789 | -4.7253316 | 2.2974E-06 | 7.1835E-05 |
| Smarce1 | 1134.14251 | 1034.00141 | 1.09684813 | 0.13336379 | 1084.07196 | 0.13384005 | 0.02835407 | 4.72031194 | 2.3548E-06 | 7.3476E-05 |
| Nlgn2 | 526.816801 | 598.697206 | 0.87993863 | -0.1845252 | 562.757004 | -0.1850044 | 0.0392046 | -4.718946 | 2.3707E-06 | 7.3815E-05 |
| Rap1gap | 1939.03752 | 2100.62584 | 0.92307611 | -0.1154785 | 2019.83168 | -0.1153458 | 0.02447013 | -4.7137385 | 2.4321E-06 | 7.5569E-05 |
| Arhgap32 | 1524.39772 | 1669.47831 | 0.91309825 | -0.131158 | 1596.93802 | -0.1329468 | 0.0282126 | -4.7123209 | 2.4491E-06 | 7.5938E-05 |
| Zfp703 | 193.456206 | 154.849649 | 1.2493164 | 0.3211389 | 174.152927 | 0.32588849 | 0.06920348 | 4.70913414 | 2.4877E-06 | 7.6974E-05 |
| Adcy9 | 1457.24988 | 1581.98783 | 0.92115113 | -0.1184902 | 1519.61885 | -0.1174636 | 0.02495609 | -4.7068101 | 2.5162E-06 | 7.7694E-05 |
| Rps26 | 1225.36842 | 1115.12378 | 1.09886315 | 0.13601172 | 1170.2461 | 0.13534365 | 0.02876854 | 4.70457114 | 2.544E-06 | 7.8245E-05 |
| Pim3 | 441.328847 | 505.341691 | 0.8733276 | -0.1954052 | 473.335269 | -0.1951731 | 0.04148629 | -4.7045201 | 2.5446E-06 | 7.8245E-05 |
| Rps5 | 1854.06466 | 1721.24719 | 1.0771635 | 0.10723726 | 1787.65593 | 0.1072926 | 0.0228191 | 4.70187601 | 2.5778E-06 | 7.9101E-05 |
| Tlr4 | 280.574025 | 338.123241 | 0.82979811 | -0.2691677 | 309.348633 | -0.2648957 | 0.05635641 | -4.7003651 | 2.597E-06 | 7.9524E-05 |
| Ssh3 | 839.673562 | 930.485612 | 0.9024036 | -0.1481553 | 885.079587 | -0.149291 | 0.03177074 | -4.6990098 | 2.6143E-06 | 7.9889E-05 |
| Sgce | 209.959158 | 169.752019 | 1.23685809 | 0.30667999 | 189.855588 | 0.30686947 | 0.06532174 | 4.69781502 | 2.6296E-06 | 8.0192E-05 |
| Ddx3x | 4164.9023 | 3916.50367 | 1.06342357 | 0.08871634 | 4040.70299 | 0.09002189 | 0.019169 | 4.69622343 | 2.6502E-06 | 8.0653E-05 |
| Epha2 | 26.8886369 | 13.2458197 | 2.02997154 | 1.0214595 | 20.0672282 | 1.03112409 | 0.21965101 | 4.69437454 | 2.6742E-06 | 8.1053E-05 |
| Clcn6 | 350.417765 | 407.232818 | 0.86048508 | -0.2167779 | 378.825292 | -0.2167099 | 0.04615991 | -4.694765 | 2.6691E-06 | 8.1053E-05 |
| Rplp2 | 1550.00371 | 1428.1402 | 1.08533022 | 0.11813406 | 1489.07196 | 0.11820029 | 0.0252045 | 4.68965008 | 2.7367E-06 | 8.2778E-05 |
| Phactr1 | 794.384934 | 882.548171 | 0.90010377 | -0.1518368 | 838.466552 | -0.1518316 | 0.03238893 | -4.6877604 | 2.7621E-06 | 8.33E-05 |
| Krt80 | 65.8912078 | 91.6872722 | 0.71865163 | -0.4766355 | 78.7892399 | -0.4734004 | 0.10099144 | -4.68753 | 2.7652E-06 | 8.33E-05 |
| St6gal2 | 4.24509126 | 0.21806577 | 19.4670222 | 4.28296031 | 2.23157841 | 3.95314098 | 0.84346316 | 4.68679744 | 2.7751E-06 | 8.3429E-05 |
| Eef1g | 978.540942 | 875.626391 | 1.11753249 | 0.16031678 | 927.083667 | 0.15982454 | 0.03410673 | 4.68601176 | 2.7858E-06 | 8.358E-05 |
| Snap25 | 1533.03628 | 1678.49416 | 0.91334026 | -0.1307757 | 1605.76522 | -0.1291944 | 0.0275766 | -4.6849292 | 2.8006E-06 | 8.3853E-05 |
| Slc22a17 | 371.830111 | 432.428111 | 0.85986572 | -0.2178167 | 402.129111 | -0.2158363 | 0.04607841 | -4.6841085 | 2.8118E-06 | 8.402E-05 |
| Usp31 | 1246.80496 | 1137.85436 | 1.09575092 | 0.13191989 | 1192.32966 | 0.13073136 | 0.02792467 | 4.68157207 | 2.8468E-06 | 8.4895E-05 |
| Slc9a2 | 9.31629823 | 1.91233152 | 4.87169621 | 2.28442417 | 5.61431477 | 2.21478163 | 0.4733287 | 4.67916195 | 2.8805E-06 | 8.5727E-05 |
| Arhgef28 | 492.08933 | 568.001426 | 0.86635228 | -0.2069743 | 530.045378 | -0.201406 | 0.04304977 | -4.6784448 | 2.8906E-06 | 8.5855E-05 |
| Pfkfb3 | 62.415637 | 41.1090425 | 1.5182946 | 0.60245175 | 51.7623397 | 0.60257833 | 0.12881368 | 4.67790634 | 2.8982E-06 | 8.5908E-05 |
| Plxnb2 | 1936.60839 | 2087.23494 | 0.9278344 | -0.1080608 | 2011.92166 | -0.1088 | 0.02326667 | -4.6762171 | 2.9222E-06 | 8.6446E-05 |
| Hhatl | 167.76782 | 210.053935 | 0.79868925 | -0.3242938 | 188.910878 | -0.3253988 | 0.0695921 | -4.6758007 | 2.9281E-06 | 8.6449E-05 |
| Cdc20 | 2251.01697 | 2097.09132 | 1.07339959 | 0.10218724 | 2174.05414 | 0.1028671 | 0.02201268 | 4.67308401 | 2.9671E-06 | 8.7426E-05 |
| Kcnq1ot1 | 260.534955 | 165.220631 | 1.57689117 | 0.6570831 | 212.877793 | 0.56628548 | 0.12128107 | 4.66919942 | 3.0238E-06 | 8.8919E-05 |
| Rpl35 | 1180.34879 | 1081.08736 | 1.09181628 | 0.12673012 | 1130.71807 | 0.12820197 | 0.02746391 | 4.66801575 | 3.0412E-06 | 8.9079E-05 |
| Ephb3 | 1258.55998 | 1370.03181 | 0.91863559 | -0.1224354 | 1314.29589 | -0.1225007 | 0.02624154 | -4.6682004 | 3.0385E-06 | 8.9079E-05 |
| Elovl6 | 452.803529 | 390.149919 | 1.16058855 | 0.2148566 | 421.476724 | 0.21487551 | 0.04605738 | 4.66538734 | 3.0804E-06 | 9.0047E-05 |
| Hnrnpab | 6156.17868 | 5770.2639 | 1.06687992 | 0.0933978 | 5963.22129 | 0.09525943 | 0.02042481 | 4.66390697 | 3.1026E-06 | 9.0519E-05 |
| Hnrnpa3 | 5530.68417 | 5264.65468 | 1.05053123 | 0.07111905 | 5397.66942 | 0.0710489 | 0.01523514 | 4.66348989 | 3.1089E-06 | 9.0525E-05 |
| Etv1 | 762.35032 | 886.84646 | 0.85961929 | -0.2182302 | 824.59839 | -0.2220034 | 0.04761313 | -4.6626506 | 3.1216E-06 | 9.0717E-05 |
| Smc3 | 1154.98211 | 1050.37685 | 1.09958831 | 0.13696347 | 1102.67948 | 0.13634091 | 0.02924889 | 4.66140498 | 3.1406E-06 | 9.1089E-05 |
| Slc12a7 | 365.566074 | 426.935798 | 0.85625538 | -0.2238869 | 396.250936 | -0.223632 | 0.04798484 | -4.660472 | 3.1549E-06 | 9.1324E-05 |
| Pbx1 | 3387.72654 | 3196.7836 | 1.0597297 | 0.08369634 | 3292.25507 | 0.08351418 | 0.01793445 | 4.65663427 | 3.2142E-06 | 9.2861E-05 |
| Qsox1 | 4443.51448 | 4741.91507 | 0.93707171 | -0.0937686 | 4592.71478 | -0.0931124 | 0.02004693 | -4.6447221 | 3.4053E-06 | 9.8001E-05 |
| Sh3glb2 | 765.412518 | 852.187532 | 0.8981738 | -0.1549334 | 808.800025 | -0.1550629 | 0.03338302 | -4.6449621 | 3.4014E-06 | 9.8001E-05 |
| Cdh8 | 64.3382653 | 42.9650594 | 1.49745552 | 0.58251315 | 53.6516623 | 0.57172172 | 0.12314807 | 4.64255513 | 3.4413E-06 | 9.8843E-05 |
| Tgfb3 | 375.965406 | 314.494256 | 1.19546033 | 0.25756625 | 345.229831 | 0.25892017 | 0.05579174 | 4.64083348 | 3.4701E-06 | 9.9477E-05 |
| Adamts5 | 871.476682 | 784.348467 | 1.11108355 | 0.15196731 | 827.912575 | 0.15145914 | 0.03264673 | 4.63933551 | 3.4953E-06 | 0.00010001 |
| Sf3b2 | 1757.91868 | 1626.64675 | 1.08070094 | 0.11196735 | 1692.28272 | 0.11139151 | 0.02402043 | 4.63736597 | 3.5288E-06 | 0.00010077 |
| Fam20b | 2340.11389 | 2521.7741 | 0.92796333 | -0.1078603 | 2430.944 | -0.1077713 | 0.02327884 | -4.6295799 | 3.6641E-06 | 0.00010443 |
| Tmem158 | 1492.1833 | 1662.2919 | 0.89766622 | -0.155749 | 1577.2376 | -0.1523785 | 0.03294192 | -4.6256706 | 3.7339E-06 | 0.00010622 |
| Tvp23b | 467.950104 | 407.476475 | 1.14841011 | 0.19963794 | 437.713289 | 0.19979505 | 0.04324682 | 4.61987779 | 3.8397E-06 | 0.0001089 |
| Rpl32 | 2058.42927 | 1902.91219 | 1.08172583 | 0.11333488 | 1980.67073 | 0.11497467 | 0.0248879 | 4.61970207 | 3.8429E-06 | 0.0001089 |
| Eif2s2 | 1496.67771 | 1372.72361 | 1.09029793 | 0.12472242 | 1434.70066 | 0.12495559 | 0.02706672 | 4.61657602 | 3.9012E-06 | 0.00011034 |
| Sart1 | 394.748484 | 340.650498 | 1.15880789 | 0.21264141 | 367.699491 | 0.21294555 | 0.04613569 | 4.61563638 | 3.9189E-06 | 0.00011063 |
| Car10 | 220.545598 | 267.095499 | 0.82571814 | -0.2762787 | 243.820548 | -0.2786589 | 0.06043066 | -4.6112166 | 4.0032E-06 | 0.0001128 |
| Tmem169 | 136.400225 | 171.247135 | 0.79651099 | -0.3282338 | 153.82368 | -0.3292472 | 0.07147596 | -4.6064043 | 4.0969E-06 | 0.00011522 |
| Erlin1 | 289.064186 | 241.09701 | 1.19895384 | 0.26177612 | 265.080598 | 0.26306279 | 0.05725939 | 4.5942295 | 4.3435E-06 | 0.00012192 |
| Cep55 | 213.065151 | 171.214301 | 1.24443548 | 0.31549144 | 192.139726 | 0.31683241 | 0.06898675 | 4.59265585 | 4.3764E-06 | 0.00012261 |
| Pkp4 | 2839.60818 | 3022.16769 | 0.93959319 | -0.0898918 | 2930.88794 | -0.0894603 | 0.01949361 | -4.5892113 | 4.4492E-06 | 0.00012418 |
| Clstn1 | 768.895735 | 854.846898 | 0.89945432 | -0.1528781 | 811.871316 | -0.1526856 | 0.03326836 | -4.5895134 | 4.4428E-06 | 0.00012418 |
| Tsc22d3 | 390.777401 | 335.977394 | 1.16310623 | 0.21798287 | 363.377397 | 0.21907724 | 0.04774167 | 4.58880539 | 4.4579E-06 | 0.00012419 |
| Mkrn1 | 2457.26236 | 2224.838 | 1.10446799 | 0.14335161 | 2341.05018 | 0.14522225 | 0.03167681 | 4.58449679 | 4.5508E-06 | 0.00012654 |
| Aurkb | 679.23743 | 598.388322 | 1.13511144 | 0.18283394 | 638.812876 | 0.18514096 | 0.04041188 | 4.58135007 | 4.6198E-06 | 0.00012822 |
| Larp1 | 1885.24382 | 1745.32189 | 1.0801697 | 0.11125798 | 1815.28286 | 0.11085301 | 0.02420526 | 4.57970718 | 4.6563E-06 | 0.00012899 |

| | | | | | | | | | | |
|---|---|---|---|---|---|---|---|---|---|---|
| Pls3 | 1143.87791 | 1298.03505 | 0.88123807 | -0.1823963 | 1220.95648 | -0.1731016 | 0.03783873 | -4.5747183 | 4.7686E-06 | 0.00013186 |
| Rab11fip4 | 503.890479 | 570.213242 | 0.88368779 | -0.1783913 | 537.051861 | -0.178363 | 0.0390066 | -4.5726358 | 4.8163E-06 | 0.00013284 |
| Ptpn14 | 219.057659 | 267.977526 | 0.81744788 | -0.2908014 | 243.517592 | -0.2796339 | 0.06115722 | -4.5723777 | 4.8222E-06 | 0.00013284 |
| Ddx5 | 12821.8934 | 12371.8817 | 1.03637375 | 0.05154439 | 12596.8876 | 0.05160967 | 0.01129188 | 4.57051089 | 4.8654E-06 | 0.00013378 |
| Rps6kb1 | 688.710216 | 611.770675 | 1.12576533 | 0.17090613 | 650.240445 | 0.17163087 | 0.03765223 | 4.55831946 | 5.1565E-06 | 0.00014153 |
| Sptbn1 | 1948.89177 | 2127.17801 | 0.9161865 | -0.1262868 | 2038.03489 | -0.1306769 | 0.02867042 | -4.557901 | 5.1667E-06 | 0.00014155 |
| Cyp51 | 620.640236 | 695.723594 | 0.89207875 | -0.164757 | 658.181915 | -0.1643824 | 0.03611029 | -4.5522307 | 5.308E-06 | 0.00014515 |
| Vill | 321.55338 | 388.859703 | 0.82691361 | -0.2741915 | 355.206541 | -0.2568942 | 0.05644761 | -4.5510209 | 5.3386E-06 | 0.00014572 |
| Vstm2a | 476.178359 | 548.063959 | 0.86883721 | -0.2028422 | 512.121159 | -0.2038692 | 0.04482518 | -4.5480954 | 5.4134E-06 | 0.00014721 |
| Irs2 | 241.002151 | 288.458358 | 0.83548333 | -0.259317 | 264.730254 | -0.2539154 | 0.05582759 | -4.5482065 | 5.4105E-06 | 0.00014721 |
| Hepacam2 | 169.079379 | 207.936896 | 0.81312832 | -0.298445 | 188.508138 | -0.3045722 | 0.06700184 | -4.5457285 | 5.4746E-06 | 0.00014861 |
| Arhgef4 | 138.282801 | 107.496148 | 1.28639773 | 0.36333677 | 122.889474 | 0.36227886 | 0.07973286 | 4.54365804 | 5.5286E-06 | 0.00014953 |
| Clint1 | 430.619438 | 358.395129 | 1.20152146 | 0.26486242 | 394.507284 | 0.26236734 | 0.05774053 | 4.54390276 | 5.5222E-06 | 0.00014953 |
| Slc29a2 | 2.97327778 | 0.0000001 | 29732777.8 | 24.8255509 | 1.48663884 | 4.09689456 | 0.90247024 | 4.53964504 | 5.6349E-06 | 0.00015212 |
| Soat1 | 311.909075 | 363.83747 | 0.85727585 | -0.2221686 | 337.873272 | -0.2220373 | 0.04896168 | -4.5349195 | 5.7625E-06 | 0.00015528 |
| Id4 | 2578.51714 | 2809.65241 | 0.91773528 | -0.12385 | 2694.08478 | -0.1163557 | 0.02568007 | -4.5309721 | 5.8713E-06 | 0.00015751 |
| Dnm2 | 1046.26105 | 1145.63842 | 0.9132559 | -0.1309089 | 1095.94973 | -0.1308604 | 0.02887951 | -4.5312538 | 5.8635E-06 | 0.00015751 |
| Clock | 1637.50148 | 1992.85553 | 0.821686 | -0.2833409 | 1815.17851 | -0.2769102 | 0.0611178 | -4.5307623 | 5.8771E-06 | 0.00015751 |
| Chd5 | 219.70442 | 262.761424 | 0.83613651 | -0.2581896 | 241.232922 | -0.2579581 | 0.05695849 | -4.5288787 | 5.9298E-06 | 0.00015864 |
| Rab3a | 470.993022 | 534.491448 | 0.88119842 | -0.1824612 | 502.742235 | -0.182411 | 0.04030637 | -4.5256136 | 6.0221E-06 | 0.00016082 |
| Ptprm | 350.108495 | 410.954197 | 0.85194043 | -0.2311755 | 380.531346 | -0.2291938 | 0.05064889 | -4.5251501 | 6.0353E-06 | 0.00016084 |
| Rnasel | 104.488274 | 135.443095 | 0.77145516 | -0.3743458 | 119.965684 | -0.3701726 | 0.08180927 | -4.5248252 | 6.0445E-06 | 0.00016084 |
| St6galnac2 | 220.784537 | 182.020537 | 1.21296498 | 0.2785379 | 201.402537 | 0.27893385 | 0.06165322 | 4.52423816 | 6.0613E-06 | 0.00016099 |
| Kcnj11 | 47.8253069 | 69.6525312 | 0.68662698 | -0.5424016 | 58.738919 | -0.5430074 | 0.12008499 | -4.5218591 | 6.1299E-06 | 0.00016252 |
| Rnf187 | 2751.86275 | 2592.09466 | 1.06163667 | 0.08629011 | 2671.9787 | 0.08623437 | 0.01907722 | 4.52027896 | 6.1758E-06 | 0.00016345 |
| Top1 | 1457.85154 | 1349.22493 | 1.08051038 | 0.11171293 | 1403.53823 | 0.11189825 | 0.02476316 | 4.51873933 | 6.2209E-06 | 0.00016435 |
| Abcb10 | 692.91372 | 769.958539 | 0.89993641 | -0.152105 | 731.436129 | -0.1523112 | 0.03372742 | -4.5159465 | 6.3035E-06 | 0.00016623 |
| Rpl27a | 2218.52492 | 2076.93329 | 1.06817341 | 0.09514588 | 2147.72911 | 0.09561102 | 0.02117955 | 4.51430833 | 6.3524E-06 | 0.00016723 |
| Rpl8 | 3238.4521 | 3046.53538 | 1.06299507 | 0.08813491 | 3142.49374 | 0.08902454 | 0.0197287 | 4.51243887 | 6.4086E-06 | 0.00016841 |
| Tmem2 | 88.9039142 | 64.9212287 | 1.36941207 | 0.45355663 | 76.9125714 | 0.45017824 | 0.09977492 | 4.51193795 | 6.4238E-06 | 0.00016851 |
| Lyst | 959.216904 | 1054.07549 | 0.91000779 | -0.1360492 | 1006.6462 | -0.140524 | 0.03116852 | -4.5085226 | 6.5281E-06 | 0.00017094 |
| Npm1 | 1540.96049 | 1401.83041 | 1.09924887 | 0.13651805 | 1471.39545 | 0.140097 | 0.03108419 | 4.50701835 | 6.5745E-06 | 0.00017185 |
| Tubb6 | 875.440102 | 791.309724 | 1.10631789 | 0.14576599 | 833.374913 | 0.14917146 | 0.03316921 | 4.49728734 | 6.8826E-06 | 0.00017959 |
| Ipo9 | 3590.55483 | 3803.68423 | 0.94396764 | -0.0831907 | 3697.11953 | -0.0831678 | 0.01850727 | -4.4937894 | 6.9967E-06 | 0.00018225 |
| Foxd2 | 137.176537 | 171.780111 | 0.7985589 | -0.3245293 | 154.478324 | -0.3226231 | 0.07180781 | -4.4928699 | 7.027E-06 | 0.00018271 |
| Syce2 | 170.506363 | 136.446523 | 1.24962043 | 0.32148994 | 153.476443 | 0.32368195 | 0.07205962 | 4.49186293 | 7.0603E-06 | 0.00018326 |
| Cetn3 | 581.236241 | 508.002514 | 1.14416017 | 0.19428902 | 544.619377 | 0.19298898 | 0.0429739 | 4.49084215 | 7.0942E-06 | 0.00018382 |
| Olfml2b | 119.347999 | 152.479444 | 0.78271534 | -0.3534404 | 135.913722 | -0.357738 | 0.07966943 | -4.4902797 | 7.113E-06 | 0.00018398 |
| Fam19a5 | 5.27551003 | 0.4531739 | 11.6412486 | 3.5411739 | 2.86434187 | 3.50271714 | 0.78034999 | 4.48864893 | 7.1676E-06 | 0.00018507 |
| Mbnl2 | 1383.20459 | 1276.31935 | 1.08374491 | 0.11602522 | 1329.76197 | 0.11610197 | 0.02588567 | 4.48518326 | 7.2851E-06 | 0.00018778 |
| Homer1 | 78.0813682 | 55.1469575 | 1.41587808 | 0.50169704 | 66.6141627 | 0.50069676 | 0.11166148 | 4.48405972 | 7.3236E-06 | 0.00018779 |
| Syngr2 | 286.39088 | 241.502438 | 1.18587159 | 0.2459478 | 263.946659 | 0.24818539 | 0.05534438 | 4.48438316 | 7.3125E-06 | 0.00018779 |
| Tpt1 | 5557.57975 | 5238.68221 | 1.06087362 | 0.0852528 | 5398.13098 | 0.0854904 | 0.0190641 | 4.48436621 | 7.3131E-06 | 0.00018779 |
| D10Wsu102e | 607.747443 | 535.772592 | 1.13433843 | 0.18185114 | 571.760018 | 0.18456497 | 0.04132913 | 4.4657361 | 7.9794E-06 | 0.00020425 |
| Zfp704 | 2010.0463 | 2151.17039 | 0.9343966 | -0.0978931 | 2080.60834 | -0.100233 | 0.0224674 | -4.4612624 | 8.1478E-06 | 0.00020821 |
| Ddc | 8249.94106 | 8617.67967 | 0.95732742 | -0.0629157 | 8433.81036 | -0.0631122 | 0.01415694 | -4.4580368 | 8.2714E-06 | 0.000211 |
| 6820431F20I | 2380.18935 | 2234.43658 | 1.06523021 | 0.09116525 | 2307.31296 | 0.09233012 | 0.02074554 | 4.45060097 | 8.563E-06 | 0.00021806 |
| Hmmr | 1478.4075 | 1346.7324 | 1.09777377 | 0.13458077 | 1412.56995 | 0.1367369 | 0.03074966 | 4.44677769 | 8.7168E-06 | 0.0002216 |
| Rpl41 | 2788.33989 | 2627.9927 | 1.06101508 | 0.08544516 | 2708.1663 | 0.08688148 | 0.01956777 | 4.44002999 | 8.9946E-06 | 0.00022827 |
| Rapgef5 | 9.57254615 | 2.14985914 | 4.45263877 | 2.15466057 | 5.86120254 | 2.08537228 | 0.46990939 | 4.43781782 | 9.0875E-06 | 0.00022984 |
| Mgat5 | 707.066228 | 783.176119 | 0.90281893 | -0.1474914 | 745.121173 | -0.1469318 | 0.03310689 | -4.4381024 | 9.0755E-06 | 0.00022984 |
| Cap2 | 4.18313458 | 0.23542266 | 17.7686151 | 4.15125933 | 2.20927852 | 3.80396914 | 0.85742301 | 4.43651393 | 9.1427E-06 | 0.00023084 |
| Slc4a7 | 563.826228 | 634.52313 | 0.88858262 | -0.1704222 | 599.174679 | -0.1699633 | 0.03832439 | -4.4348601 | 9.2132E-06 | 0.00023223 |
| Abcg1 | 917.370556 | 1011.34148 | 0.9070829 | -0.1406937 | 964.356016 | -0.1375869 | 0.03102984 | -4.4340197 | 9.2492E-06 | 0.00023274 |
| Gfra1 | 258.386332 | 304.606444 | 0.84826286 | -0.2374167 | 281.496388 | -0.2363881 | 0.05335169 | -4.4307524 | 9.3905E-06 | 0.00023589 |
| Spry4 | 8.96199606 | 2.47747261 | 3.61739461 | 1.85495099 | 5.71973424 | 1.85613126 | 0.41907237 | 4.42914255 | 9.4608E-06 | 0.00023726 |
| Smc4 | 3493.56605 | 3312.07509 | 1.05479676 | 0.07696504 | 3402.82057 | 0.07687811 | 0.01736409 | 4.42742066 | 9.5367E-06 | 0.00023875 |
| Rps11 | 1591.75443 | 1473.49292 | 1.0802593 | 0.11137765 | 1532.62368 | 0.11336799 | 0.02562147 | 4.42472551 | 9.6565E-06 | 0.00024135 |
| Aldh3b1 | 262.838502 | 310.91982 | 0.84535782 | -0.242366 | 286.879161 | -0.2433574 | 0.05502304 | -4.4228268 | 9.7418E-06 | 0.00024307 |
| Nr2e1 | 4.45592443 | 13.0415168 | 0.34167225 | -1.549315 | 8.74872053 | -1.4920441 | 0.33740749 | -4.4220836 | 9.7754E-06 | 0.0002435 |
| Dll3 | 521.420826 | 590.91843 | 0.88239053 | -0.1805108 | 556.169628 | -0.1844728 | 0.04188674 | -4.4040863 | 1.0623E-05 | 0.00026329 |
| Nyap1 | 380.847027 | 440.39251 | 0.86478997 | -0.2095783 | 410.619769 | -0.2076549 | 0.04715009 | -4.4041251 | 1.0621E-05 | 0.00026329 |
| Tspan8 | 571.570663 | 668.470483 | 0.85504249 | -0.225932 | 620.020573 | -0.2092773 | 0.04751475 | -4.4044714 | 1.0604E-05 | 0.00026329 |
| Pi4ka | 1370.32147 | 1482.21764 | 0.92450759 | -0.1132429 | 1426.26955 | -0.1135575 | 0.02580286 | -4.4009666 | 1.0777E-05 | 0.00026622 |
| Crim1 | 559.452155 | 632.244061 | 0.8848674 | -0.1764668 | 595.848108 | -0.1803425 | 0.04097802 | -4.400958 | 1.0777E-05 | 0.00026622 |
| Cdkn1a | 1143.1211 | 1244.77022 | 0.91833905 | -0.1229012 | 1193.94566 | -0.1228463 | 0.02792385 | -4.399333 | 1.0858E-05 | 0.00026777 |
| Aldh1a3 | 1141.52806 | 1243.2272 | 0.91819746 | -0.1231237 | 1192.37763 | -0.1231596 | 0.02800013 | -4.3985373 | 1.0898E-05 | 0.00026831 |

| | | | | | | | | | | |
|---|---|---|---|---|---|---|---|---|---|---|
| Srsf10 | 2125.70075 | 1982.33049 | 1.07232409 | 0.100741 | 2054.01562 | 0.10268189 | 0.02336674 | 4.39436037 | 1.111E-05 | 0.00027307 |
| Nusap1 | 1632.0945 | 1512.0448 | 1.07939559 | 0.1102237 | 1572.06965 | 0.11133792 | 0.02534732 | 4.39249334 | 1.1206E-05 | 0.00027497 |
| Cyb5rl | 88.7264394 | 64.9698238 | 1.36565615 | 0.44959428 | 76.8481315 | 0.44704304 | 0.1018674 | 4.38847977 | 1.1415E-05 | 0.00027963 |
| Gnai1 | 3.9177161 | 0.22726116 | 17.2388283 | 4.10758982 | 2.07248853 | 3.76309574 | 0.85942408 | 4.37862499 | 1.1943E-05 | 0.00029171 |
| Lmo4 | 108.696162 | 76.7244848 | 1.41670761 | 0.50254204 | 92.7103231 | 0.49205806 | 0.11237912 | 4.37855422 | 1.1947E-05 | 0.00029171 |
| Pgap1 | 258.96516 | 305.117272 | 0.84873976 | -0.2366058 | 282.041216 | -0.2361978 | 0.05396623 | -4.3767707 | 1.2045E-05 | 0.00029362 |
| Pgam1 | 1067.1096 | 970.520879 | 1.09952256 | 0.13687721 | 1018.81524 | 0.13665225 | 0.0312306 | 4.37558815 | 1.2111E-05 | 0.00029473 |
| Atp7b | 104.147217 | 134.013971 | 0.77713702 | -0.3637591 | 119.080594 | -0.3584763 | 0.08198121 | -4.3726645 | 1.2274E-05 | 0.00029822 |
| Kat2a | 490.373555 | 432.04463 | 1.13500671 | 0.18270083 | 461.209092 | 0.18155393 | 0.04157596 | 4.36680038 | 1.2608E-05 | 0.00030583 |
| Sdad1 | 241.337391 | 201.374629 | 1.19844983 | 0.26116952 | 221.35601 | 0.25981829 | 0.05951117 | 4.36587444 | 1.2662E-05 | 0.00030663 |
| Irs1 | 222.981975 | 265.345444 | 0.84034597 | -0.2509447 | 244.163709 | -0.2512281 | 0.05757779 | -4.3632809 | 1.2813E-05 | 0.00030978 |
| Psmd12 | 792.428597 | 713.037269 | 1.11134247 | 0.15230346 | 752.732933 | 0.15380855 | 0.03526484 | 4.36152738 | 1.2916E-05 | 0.00031177 |
| Parp1 | 2098.89255 | 2237.12225 | 0.93821093 | -0.0920158 | 2168.0074 | -0.0919918 | 0.02113232 | -4.3531308 | 1.3421E-05 | 0.0003231 |
| St3gal6 | 155.775659 | 190.78614 | 0.81649358 | -0.2924865 | 173.280899 | -0.2933483 | 0.06739546 | -4.3526413 | 1.3451E-05 | 0.0003231 |
| Plekha5 | 104.569784 | 134.131382 | 0.77960714 | -0.3591808 | 119.350583 | -0.355038 | 0.08156498 | -4.3528239 | 1.344E-05 | 0.0003231 |
| Leng8 | 990.281807 | 874.728195 | 1.13210231 | 0.17900434 | 932.505001 | 0.18789462 | 0.04319107 | 4.35031132 | 1.3594E-05 | 0.0003255 |
| Pcyt1b | 380.169709 | 451.286367 | 0.84241346 | -0.2473996 | 415.728038 | -0.2471515 | 0.05680802 | -4.3506439 | 1.3574E-05 | 0.0003255 |
| Ldha | 1866.17867 | 1732.92209 | 1.07689704 | 0.10688032 | 1799.55038 | 0.10554252 | 0.02429457 | 4.34428376 | 1.3973E-05 | 0.00033402 |
| Mettl2 | 136.214766 | 105.932969 | 1.2858581 | 0.36273144 | 121.073867 | 0.36292469 | 0.08367862 | 4.33712575 | 1.4436E-05 | 0.00034453 |
| Sema3c | 3.79743909 | 0.23581399 | 16.103536 | 4.0093056 | 2.01662644 | 3.7306595 | 0.86116628 | 4.33210125 | 1.4769E-05 | 0.00035192 |
| Myh14 | 1598.80549 | 1716.83344 | 0.93125254 | -0.1027556 | 1657.81946 | -0.1033727 | 0.02388757 | -4.3274665 | 1.5083E-05 | 0.00035883 |
| Igsf10 | 5.20600496 | 0.5702855 | 9.12876958 | 3.19042042 | 2.88814513 | 3.04154928 | 0.7042728 | 4.31870903 | 1.5694E-05 | 0.00037271 |
| Dnajc7 | 1207.52618 | 1109.87523 | 1.08798372 | 0.12165697 | 1158.70071 | 0.12319937 | 0.02852899 | 4.31839212 | 1.5717E-05 | 0.00037271 |
| Hpgd | 234.032077 | 276.481572 | 0.84646537 | -0.240477 | 255.256824 | -0.2389272 | 0.05533313 | -4.3179775 | 1.5747E-05 | 0.00037281 |
| Plekhb2 | 1993.37769 | 2182.72794 | 0.91325064 | -0.1309172 | 2088.05281 | -0.127399 | 0.02953017 | -4.3141984 | 1.6018E-05 | 0.00037812 |
| Armcx2 | 678.36033 | 751.075824 | 0.90318488 | -0.1469068 | 714.718077 | -0.1470702 | 0.03409018 | -4.3141499 | 1.6022E-05 | 0.00037812 |
| Abca1 | 16.6013491 | 29.4460497 | 0.56378867 | -0.8267736 | 23.0236993 | -0.817613 | 0.18955458 | -4.3133379 | 1.6081E-05 | 0.00037891 |
| Arpp21 | 48.2474708 | 68.4416184 | 0.70494345 | -0.5044206 | 58.3445445 | -0.504584 | 0.11704983 | -4.3108478 | 1.6263E-05 | 0.00038259 |
| Gdi1 | 2772.5755 | 2942.84083 | 0.94214253 | -0.0859828 | 2857.70817 | -0.0859365 | 0.01996367 | -4.3046444 | 1.6725E-05 | 0.00039285 |
| Unc93b1 | 180.657488 | 217.682245 | 0.82991375 | -0.2689667 | 199.169866 | -0.2692038 | 0.06255306 | -4.3036081 | 1.6804E-05 | 0.00039407 |
| Atp2c1 | 1461.81229 | 1589.63681 | 0.91958885 | -0.1209391 | 1525.72455 | -0.1197836 | 0.02784299 | -4.302109 | 1.6918E-05 | 0.00039612 |
| Mtor | 842.517291 | 935.457262 | 0.90064755 | -0.1509654 | 888.987277 | -0.1565726 | 0.03640583 | -4.300756 | 1.7022E-05 | 0.00039791 |
| Thbs1 | 880.518649 | 609.077262 | 1.44566002 | 0.53172831 | 744.797955 | 0.54213966 | 0.12612211 | 4.29853 | 1.7193E-05 | 0.0004013 |
| Rps19 | 878.195531 | 799.661169 | 1.09820955 | 0.13515336 | 838.92835 | 0.1351455 | 0.03150306 | 4.28991629 | 1.7874E-05 | 0.00041653 |
| Wdfy4 | 0.92087503 | 6.20229056 | 0.14847338 | -2.7517238 | 3.5615827 | -2.7331265 | 0.63725477 | -4.2889071 | 1.7955E-05 | 0.00041777 |
| Dock5 | 639.528988 | 708.534833 | 0.90260769 | -0.147829 | 674.03191 | -0.1476578 | 0.03444763 | -4.2864428 | 1.8156E-05 | 0.00042176 |
| Ccnd1 | 17.6978123 | 7.6339666 | 2.31829837 | 1.21306626 | 12.6658894 | 1.20470272 | 0.2811616 | 4.2847342 | 1.8296E-05 | 0.00042435 |
| Ica1l | 659.676137 | 736.850553 | 0.89526449 | -0.1596141 | 698.263345 | -0.1581333 | 0.03695076 | -4.2795686 | 1.8726E-05 | 0.00043364 |
| Aplp1 | 1954.96627 | 2171.7815 | 0.90016711 | -0.1517352 | 2063.37388 | -0.1480725 | 0.03463239 | -4.2755515 | 1.9066E-05 | 0.00044085 |
| Gnaq | 676.063195 | 606.734787 | 1.11426477 | 0.15609208 | 641.398991 | 0.15465337 | 0.03618246 | 4.27426375 | 1.9177E-05 | 0.00044271 |
| Sept5 | 848.401099 | 936.573601 | 0.9058563 | -0.1426459 | 892.48735 | -0.1423717 | 0.03331461 | -4.2735497 | 1.9239E-05 | 0.00044344 |
| Satb2 | 9.3616097 | 2.4946767 | 3.75263444 | 1.90790376 | 5.9281431 | 1.7942166 | 0.41999139 | 4.27203183 | 1.937E-05 | 0.00044578 |
| Wdr43 | 461.633008 | 405.326504 | 1.13891641 | 0.18766187 | 433.479756 | 0.18940824 | 0.04439503 | 4.26642849 | 1.9863E-05 | 0.00045641 |
| Capzb | 1197.23538 | 1297.13226 | 0.92298636 | -0.1156188 | 1247.18382 | -0.1161136 | 0.02722436 | -4.2650637 | 1.9985E-05 | 0.0004585 |
| Galnt6 | 16.5328712 | 29.4053243 | 0.56224074 | -0.8307401 | 22.9690977 | -0.8193485 | 0.19221873 | -4.2625841 | 2.0208E-05 | 0.0004629 |
| Incenp | 574.229096 | 469.768733 | 1.22236551 | 0.28967574 | 521.998914 | 0.29808335 | 0.069974 | 4.2599159 | 2.045E-05 | 0.00046643 |
| Kif1b | 1128.12576 | 1224.05796 | 0.92162773 | -0.117744 | 1176.09186 | -0.1183614 | 0.02778533 | -4.2598532 | 2.0456E-05 | 0.00046643 |
| Enpp2 | 144.300609 | 178.628823 | 0.80782377 | -0.3078875 | 161.464716 | -0.3106054 | 0.07290865 | -4.2601988 | 2.0425E-05 | 0.00046643 |
| Rcor2 | 120.703654 | 93.3158983 | 1.29349506 | 0.37127455 | 107.009776 | 0.37116304 | 0.08715746 | 4.25853461 | 2.0577E-05 | 0.00046847 |
| Chuk | 340.78268 | 293.701404 | 1.1603032 | 0.21450185 | 317.242042 | 0.21463407 | 0.05042167 | 4.25678247 | 2.0739E-05 | 0.00047143 |
| Sec61g | 570.112838 | 508.47232 | 1.12122689 | 0.16507825 | 539.292579 | 0.16953473 | 0.03985038 | 4.25428134 | 2.0972E-05 | 0.000476 |
| Gna13 | 917.320634 | 835.204746 | 1.09831827 | 0.13529618 | 876.26269 | 0.13515739 | 0.0318051 | 4.2495509 | 2.142E-05 | 0.00048542 |
| Fgf13 | 1188.61071 | 1096.37737 | 1.08412554 | 0.11653183 | 1142.49404 | 0.1161705 | 0.02735052 | 4.24746966 | 2.162E-05 | 0.0004892 |
| Ryr3 | 270.335466 | 223.059007 | 1.21194598 | 0.27732539 | 246.697237 | 0.2730826 | 0.06433119 | 4.24494888 | 2.1864E-05 | 0.00049398 |
| Uty | 237.14612 | 198.887983 | 1.19236023 | 0.25382016 | 218.017051 | 0.25422559 | 0.05996709 | 4.23941875 | 2.241E-05 | 0.00050554 |
| Rpl36a | 482.420253 | 423.495195 | 1.13913985 | 0.18794487 | 452.957724 | 0.18826164 | 0.04442706 | 4.23754451 | 2.2598E-05 | 0.000509 |
| Ywhaq | 5643.34745 | 5391.97108 | 1.0466205 | 0.06573842 | 5517.65926 | 0.06565122 | 0.0154983 | 4.23602746 | 2.2751E-05 | 0.00051167 |
| Pcdhb21 | 39.4277736 | 57.3202468 | 0.68785073 | -0.5398326 | 48.3740101 | -0.5399596 | 0.12753238 | -4.2339022 | 2.2967E-05 | 0.00051575 |
| Tmem178b | 447.347824 | 508.505841 | 0.87972996 | -0.1848673 | 477.926832 | -0.1820454 | 0.0430038 | -4.2332394 | 2.3035E-05 | 0.00051649 |
| Prr16 | 130.931351 | 101.01901 | 1.29610606 | 0.37418378 | 115.97518 | 0.3787467 | 0.08948156 | 4.23267895 | 2.3092E-05 | 0.000517 |
| Kif5b | 5441.25891 | 5204.95014 | 1.04540077 | 0.06405613 | 5323.10452 | 0.0641305 | 0.015164 | 4.22912751 | 2.346E-05 | 0.00052443 |
| Cdkl5 | 14.4549845 | 6.16511039 | 2.34464326 | 1.22936843 | 10.3100473 | 1.23781561 | 0.29323308 | 4.22126861 | 2.4293E-05 | 0.00054224 |
| Rpl36 | 1032.81466 | 948.401387 | 1.08900585 | 0.12301171 | 990.608024 | 0.12337606 | 0.02926887 | 4.21526589 | 2.4948E-05 | 0.00055517 |
| Csnk1d | 1697.26048 | 1578.97857 | 1.07491039 | 0.1042164 | 1638.11953 | 0.10435939 | 0.02475574 | 4.21556336 | 2.4916E-05 | 0.00055517 |
| Mfsd6 | 1617.02984 | 1737.05731 | 0.93090184 | -0.103299 | 1677.04357 | -0.1033819 | 0.02452751 | -4.2149381 | 2.4985E-05 | 0.00055517 |
| Trim32 | 414.932782 | 362.673585 | 1.1440943 | 0.19420597 | 388.803184 | 0.19422818 | 0.04610852 | 4.2124139 | 2.5266E-05 | 0.00056057 |
| Alyref | 666.114829 | 586.385789 | 1.13596687 | 0.18392076 | 626.250309 | 0.18497884 | 0.04392124 | 4.21160364 | 2.5356E-05 | 0.00056175 |

| | | | | | | | | | | |
|---|---|---|---|---|---|---|---|---|---|---|
| Rxra | 267.374671 | 326.730935 | 0.81833289 | -0.2892403 | 297.052803 | -0.2704694 | 0.06425279 | -4.2094572 | 2.5598E-05 | 0.00056626 |
| Spry1 | 308.182586 | 262.659357 | 1.17331661 | 0.23059236 | 285.420971 | 0.2318525 | 0.05515704 | 4.20349771 | 2.6282E-05 | 0.00058052 |
| Hmgn5 | 571.494294 | 510.949801 | 1.11849401 | 0.16155753 | 541.222047 | 0.16285296 | 0.03875319 | 4.20231116 | 2.642E-05 | 0.0005827 |
| Gm14322 | 19.7700823 | 9.35206525 | 2.11398037 | 1.07996198 | 14.5610737 | 1.08232714 | 0.25761259 | 4.20137511 | 2.653E-05 | 0.00058327 |
| Cdc14a | 1049.07804 | 961.349187 | 1.09125597 | 0.12598955 | 1005.21361 | 0.1268397 | 0.03019214 | 4.20108397 | 2.6564E-05 | 0.00058327 |
| Cul7 | 502.737075 | 566.193921 | 0.88792383 | -0.1714922 | 534.465498 | -0.1708598 | 0.04066807 | -4.2013249 | 2.6536E-05 | 0.00058327 |
| 2510039O18 | 800.171709 | 883.952272 | 0.90522049 | -0.1436589 | 842.06199 | -0.1450684 | 0.0345507 | -4.1987108 | 2.6844E-05 | 0.00058854 |
| Samd4b | 838.953095 | 921.555454 | 0.91036637 | -0.1354808 | 880.254275 | -0.1355655 | 0.03231539 | -4.1950745 | 2.7278E-05 | 0.00059718 |
| Celf6 | 269.241736 | 315.210675 | 0.8541644 | -0.2274143 | 292.226205 | -0.2267554 | 0.05412699 | -4.1893223 | 2.7979E-05 | 0.00061162 |
| Tra2a | 1010.19283 | 913.390884 | 1.10598085 | 0.14532641 | 961.791855 | 0.15163084 | 0.03621068 | 4.18746184 | 2.8209E-05 | 0.00061559 |
| Hcn1 | 976.221166 | 1060.68938 | 0.92036479 | -0.1197223 | 1018.45528 | -0.1197078 | 0.02858911 | -4.1871834 | 2.8244E-05 | 0.00061559 |
| Casp7 | 160.895869 | 129.890578 | 1.23870315 | 0.3088305 | 145.393223 | 0.3084802 | 0.0736821 | 4.18663694 | 2.8312E-05 | 0.00061617 |
| Ddost | 1924.59471 | 2054.86839 | 0.93660242 | -0.0944913 | 1989.73155 | -0.0962544 | 0.02303494 | -4.1786246 | 2.9328E-05 | 0.00063561 |
| Ncam1 | 1270.89112 | 1378.45895 | 0.92196515 | -0.1172159 | 1324.67503 | -0.1180493 | 0.02824883 | -4.1789082 | 2.9291E-05 | 0.00063561 |
| Mdga2 | 141.883951 | 174.954633 | 0.81097567 | -0.3022695 | 158.419292 | -0.3013058 | 0.07210729 | -4.178576 | 2.9334E-05 | 0.00063561 |
| Pitpnm1 | 1625.07233 | 1754.89144 | 0.92602442 | -0.1108779 | 1689.98188 | -0.1120421 | 0.02682196 | -4.1772505 | 2.9505E-05 | 0.00063839 |
| Fam163a | 606.482874 | 677.346357 | 0.89538073 | -0.1594268 | 641.914615 | -0.1592744 | 0.03813901 | -4.1761557 | 2.9648E-05 | 0.00064054 |
| Ccdc47 | 1021.23623 | 935.602494 | 1.09152791 | 0.12634902 | 978.419364 | 0.12639143 | 0.03028116 | 4.17392901 | 2.9939E-05 | 0.00064589 |
| Olfm2 | 503.777594 | 578.164329 | 0.8713398 | -0.1986926 | 540.970961 | -0.1898345 | 0.04548735 | -4.1733466 | 3.0016E-05 | 0.0006466 |
| Padi2 | 317.948043 | 368.183203 | 0.86355934 | -0.2116328 | 343.065623 | -0.2103865 | 0.05042856 | -4.1719716 | 3.0198E-05 | 0.00064958 |
| Ckap2 | 619.704578 | 543.616253 | 1.13996698 | 0.18899204 | 581.660415 | 0.18950586 | 0.04545445 | 4.16913796 | 3.0575E-05 | 0.00065675 |
| Tlk2 | 644.585125 | 579.814116 | 1.11170995 | 0.15278044 | 612.19962 | 0.15280131 | 0.03667061 | 4.16686081 | 3.0882E-05 | 0.00066238 |
| Camk2n1 | 1415.9595 | 1547.1744 | 0.91519062 | -0.1278558 | 1481.56695 | -0.1272334 | 0.03054443 | -4.1655179 | 3.1065E-05 | 0.00066533 |
| Cmklr1 | 8.59119121 | 2.5750259 | 3.33635137 | 1.73827124 | 5.58310846 | 1.73889598 | 0.41769259 | 4.16309986 | 3.1396E-05 | 0.00067145 |
| Rarb | 222.32687 | 143.624283 | 1.54797549 | 0.63038263 | 182.975576 | 0.56471269 | 0.13590568 | 4.15518108 | 3.2503E-05 | 0.00069316 |
| Rrp12 | 131.479084 | 103.142726 | 1.27472958 | 0.35019123 | 117.310905 | 0.34505465 | 0.0830488 | 4.15484193 | 3.2551E-05 | 0.00069316 |
| Ulk1 | 1120.72878 | 1213.1338 | 0.9238295 | -0.1143015 | 1166.93129 | -0.1142193 | 0.02748898 | -4.1550928 | 3.2516E-05 | 0.00069316 |
| Actn4 | 2851.89019 | 3027.21095 | 0.94208505 | -0.0860708 | 2939.55057 | -0.0865152 | 0.02082972 | -4.1534515 | 3.275E-05 | 0.00069639 |
| Obsl1 | 3221.63603 | 3000.5448 | 1.0736837 | 0.10256904 | 3111.09041 | 0.09969 | 0.02401052 | 4.1519298 | 3.2968E-05 | 0.00070003 |
| Pea15a | 895.36882 | 979.803513 | 0.91382487 | -0.1300104 | 937.586167 | -0.1307046 | 0.03148909 | -4.1507911 | 3.3133E-05 | 0.00070252 |
| Apba2 | 44.3131295 | 27.9993153 | 1.58265047 | 0.66234267 | 36.1562223 | 0.6585891 | 0.15873152 | 4.1490757 | 3.3382E-05 | 0.00070679 |
| Rorc | 105.270335 | 80.8397132 | 1.30221065 | 0.38096284 | 93.0550241 | 0.37829899 | 0.09119745 | 4.14813103 | 3.352E-05 | 0.0007087 |
| Phtf2 | 435.705319 | 384.093858 | 1.134372 | 0.18189383 | 409.899589 | 0.18192405 | 0.04390042 | 4.14401645 | 3.4128E-05 | 0.00072052 |
| Idh1 | 380.436636 | 432.9791 | 0.87864896 | -0.1866412 | 406.707868 | -0.1878522 | 0.04534856 | -4.1424065 | 3.4368E-05 | 0.00072456 |
| Clip4 | 63.1290547 | 44.7123085 | 1.41189432 | 0.49763211 | 53.9206815 | 0.5004785 | 0.12089911 | 4.1396375 | 3.4786E-05 | 0.00073232 |
| Psmc3ip | 149.323417 | 119.137614 | 1.25336921 | 0.32581146 | 134.230516 | 0.33063952 | 0.0799623 | 4.13494271 | 3.5504E-05 | 0.0007464 |
| Nefl | 0.0000001 | 2.59320444 | 3.8562E-08 | -24.628233 | 1.29660217 | -3.677755 | 0.88988912 | -4.1328238 | 3.5833E-05 | 0.00075225 |
| Zfp536 | 11.0066783 | 21.1898682 | 0.51943118 | -0.9449955 | 16.0982731 | -0.942033 | 0.22819965 | -4.1281088 | 3.6576E-05 | 0.00076675 |
| Srp19 | 247.058851 | 208.876987 | 1.18279594 | 0.24220119 | 227.967919 | 0.24093461 | 0.05837073 | 4.12766173 | 3.6647E-05 | 0.00076716 |
| Rcl1 | 75.5307634 | 54.3717408 | 1.38915478 | 0.47420735 | 64.951252 | 0.4739671 | 0.11492591 | 4.12410991 | 3.7217E-05 | 0.00077799 |
| Clu | 356.660814 | 478.438851 | 0.74546792 | -0.4237818 | 417.549832 | -0.4049544 | 0.09821649 | -4.1230794 | 3.7384E-05 | 0.00078038 |
| Zfp423 | 1.7573416 | 7.34075068 | 0.23939535 | -2.0625329 | 4.54904604 | -2.0578437 | 0.49918379 | -4.122417 | 3.7492E-05 | 0.00078153 |
| Aldh1a1 | 679.389193 | 750.038109 | 0.90580623 | -0.1427256 | 714.713651 | -0.14195 | 0.03444197 | -4.1214263 | 3.7653E-05 | 0.00078379 |
| Slc6a20a | 139.235241 | 172.811181 | 0.80570736 | -0.3116722 | 156.023211 | -0.3081161 | 0.07478015 | -4.1202933 | 3.7839E-05 | 0.00078655 |
| Nufip2 | 1040.88823 | 946.377753 | 1.09986549 | 0.1373271 | 993.632992 | 0.13607809 | 0.03303681 | 4.11898338 | 3.8055E-05 | 0.00078993 |
| Cic | 1507.54336 | 1616.30061 | 0.93271224 | -0.100496 | 1561.92198 | -0.1004661 | 0.02439817 | -4.1177707 | 3.8255E-05 | 0.00079299 |
| Id3 | 622.391407 | 556.720251 | 1.11796078 | 0.16086958 | 589.555829 | 0.1611852 | 0.03915063 | 4.11705213 | 3.8375E-05 | 0.00079435 |
| Rpl21 | 2538.18295 | 2393.52224 | 1.06043842 | 0.08466085 | 2465.8526 | 0.08478537 | 0.02060002 | 4.11579078 | 3.8585E-05 | 0.00079759 |
| Fgfr1op | 240.947297 | 202.824415 | 1.18796002 | 0.24848629 | 221.885856 | 0.24789809 | 0.06026957 | 4.11315541 | 3.9029E-05 | 0.00080563 |
| Agtrap | 326.565676 | 374.345068 | 0.87236537 | -0.1969956 | 350.455372 | -0.1974678 | 0.04802642 | -4.1116487 | 3.9284E-05 | 0.00080978 |
| Gpx2 | 327.83559 | 282.283749 | 1.16136898 | 0.21582641 | 305.059669 | 0.21368973 | 0.05200412 | 4.10909247 | 3.9722E-05 | 0.00081766 |
| Mafb | 264.205275 | 310.702395 | 0.85034837 | -0.2338741 | 287.453835 | -0.234377 | 0.05709721 | -4.1048764 | 4.0453E-05 | 0.00083156 |
| Pcdh7 | 3112.66836 | 3310.13553 | 0.94034469 | -0.0887384 | 3211.40195 | -0.0856367 | 0.02087834 | -4.1017 | 4.1013E-05 | 0.00084189 |
| Cdc27 | 931.390055 | 841.815536 | 1.10640635 | 0.14588134 | 886.602796 | 0.14579547 | 0.03554821 | 4.10134439 | 4.1076E-05 | 0.00084202 |
| Bcam | 948.530543 | 1044.66789 | 0.90797329 | -0.1392782 | 996.599218 | -0.1393005 | 0.03397934 | -4.0995657 | 4.1393E-05 | 0.00084618 |
| Jhy | 333.073086 | 382.072931 | 0.87175264 | -0.1980093 | 357.573008 | -0.1953724 | 0.04765459 | -4.0997595 | 4.1358E-05 | 0.00084618 |
| Lrrc59 | 1464.50559 | 1359.44049 | 1.07728555 | 0.10740071 | 1411.97304 | 0.10770653 | 0.02632501 | 4.0914148 | 4.2875E-05 | 0.00087527 |
| Syt13 | 593.149277 | 659.105438 | 0.89993079 | -0.152114 | 626.127358 | -0.1524021 | 0.03726597 | -4.0895784 | 4.3216E-05 | 0.00088102 |
| Dlg5 | 666.92235 | 736.227047 | 0.90586505 | -0.142632 | 701.574698 | -0.1425638 | 0.03487715 | -4.0875975 | 4.3586E-05 | 0.00088735 |
| Txnip | 1203.9556 | 1089.67836 | 1.10487245 | 0.14387983 | 1146.81698 | 0.12964315 | 0.03174122 | 4.08437762 | 4.4195E-05 | 0.00089851 |
| Ppp2r1a | 1918.27152 | 2051.75617 | 0.93494127 | -0.0970524 | 1985.01385 | -0.0973118 | 0.02389031 | -4.0732738 | 4.6357E-05 | 0.00094117 |
| Crk | 1535.89003 | 1421.12944 | 1.08075308 | 0.11203695 | 1478.50974 | 0.11313092 | 0.02777903 | 4.07252946 | 4.6505E-05 | 0.00094289 |
| Bub1b | 1441.1388 | 1337.14387 | 1.07777393 | 0.10805459 | 1389.14133 | 0.109695 | 0.02694327 | 4.07133228 | 4.6745E-05 | 0.00094646 |
| Minpp1 | 185.987039 | 153.48422 | 1.21176652 | 0.27711176 | 169.735629 | 0.2768556 | 0.06803685 | 4.06920071 | 4.7175E-05 | 0.00095386 |
| Bach2 | 16.1135438 | 7.64448551 | 2.10786505 | 1.0757825 | 11.8790146 | 1.0824246 | 0.26605337 | 4.06844904 | 4.7327E-05 | 0.00095551 |
| Fhod3 | 102.326651 | 77.880048 | 1.31390072 | 0.39385627 | 90.1033494 | 0.39513321 | 0.09712817 | 4.06816263 | 4.7385E-05 | 0.00095551 |
| Ablim1 | 222.522417 | 266.523891 | 0.83490608 | -0.2603142 | 244.523154 | -0.2574532 | 0.06337998 | -4.0620593 | 4.8642E-05 | 0.00097951 |

| | | | | | | | | | | |
|---|---|---|---|---|---|---|---|---|---|---|
| Clpx | 385.135801 | 336.822463 | 1.14343859 | 0.19337888 | 360.979132 | 0.19382087 | 0.04777856 | 4.05664969 | 4.9782E-05 | 0.0010011 |
| Cog6 | 689.882382 | 758.647669 | 0.90935807 | -0.1370796 | 724.265025 | -0.1371105 | 0.03380817 | -4.0555449 | 5.0018E-05 | 0.00100449 |
| Thra | 533.468786 | 612.697537 | 0.87068864 | -0.1997712 | 573.083161 | -0.1942211 | 0.04793817 | -4.0514907 | 5.0892E-05 | 0.00102067 |
| Gpa33 | 326.788097 | 380.109361 | 0.85972126 | -0.2180591 | 353.448729 | -0.2147233 | 0.05316761 | -4.038611 | 5.3769E-05 | 0.0010769 |
| Casp9 | 367.306689 | 420.127516 | 0.8742743 | -0.1938421 | 393.717102 | -0.1937225 | 0.04797594 | -4.0379108 | 5.3929E-05 | 0.00107866 |
| Asic1 | 129.814179 | 103.514515 | 1.2540674 | 0.32661489 | 116.664347 | 0.32764376 | 0.0811738 | 4.03632414 | 5.4295E-05 | 0.00108451 |
| Homer3 | 69.0453448 | 49.337364 | 1.39945346 | 0.48486351 | 59.1913543 | 0.4930622 | 0.12219194 | 4.03514514 | 5.4569E-05 | 0.00108851 |
| Lnx1 | 143.572989 | 176.208892 | 0.81478856 | -0.2955024 | 159.890941 | -0.2956695 | 0.07328565 | -4.0344805 | 5.4723E-05 | 0.00109013 |
| Cdca8 | 1393.77548 | 1280.96322 | 1.0880683 | 0.12176913 | 1337.36935 | 0.12440316 | 0.03084182 | 4.03358735 | 5.4932E-05 | 0.00109281 |
| Rhbdd1 | 445.256116 | 500.200722 | 0.89015488 | -0.1678717 | 472.728419 | -0.1676615 | 0.0416652 | -4.0240185 | 5.7213E-05 | 0.00113668 |
| Mfn2 | 667.093831 | 738.274592 | 0.90358498 | -0.1462678 | 702.684211 | -0.1461714 | 0.03633826 | -4.0225206 | 5.7579E-05 | 0.0011424 |
| Tspan7 | 1522.59399 | 1652.98814 | 0.9211161 | -0.1185451 | 1587.79107 | -0.1172421 | 0.02915294 | -4.0216223 | 5.7799E-05 | 0.00114523 |
| Dbp | 600.748541 | 689.633397 | 0.87111289 | -0.1990684 | 645.190969 | -0.2009492 | 0.04997656 | -4.0208691 | 5.7984E-05 | 0.00114737 |
| Pcdha2 | 0.0000001 | 2.7735258 | 3.6055E-08 | -24.725218 | 1.38676285 | -3.6312491 | 0.90322752 | -4.0203039 | 5.8123E-05 | 0.00114859 |
| Cdk5r2 | 655.124683 | 723.427521 | 0.90558441 | -0.143079 | 689.276102 | -0.143015 | 0.03561734 | -4.0153202 | 5.9365E-05 | 0.00117157 |
| Jup | 2871.05506 | 3034.78434 | 0.94604912 | -0.080013 | 2952.9197 | -0.0800618 | 0.01994215 | -4.0147016 | 5.9521E-05 | 0.00117309 |
| Cuedc2 | 202.527701 | 168.9887 | 1.1984689 | 0.26119247 | 185.7582 | 0.26201107 | 0.06527041 | 4.01423951 | 5.9638E-05 | 0.00117383 |
| Fgf14 | 975.115939 | 886.455123 | 1.10001726 | 0.13752617 | 930.785531 | 0.13783568 | 0.03435589 | 4.01199599 | 6.0208E-05 | 0.00118347 |
| Gatsl2 | 1079.68885 | 1178.10441 | 0.91646279 | -0.1258518 | 1128.89663 | -0.1245371 | 0.03104523 | -4.0114713 | 6.0342E-05 | 0.00118453 |
| Rbm3 | 3162.73772 | 2967.49152 | 1.06579503 | 0.09193002 | 3065.11462 | 0.09855714 | 0.02459138 | 4.00779201 | 6.1289E-05 | 0.00120154 |
| Crip2 | 341.996082 | 391.487185 | 0.87358181 | -0.1949853 | 366.741634 | -0.1934927 | 0.04829302 | -4.0066383 | 6.1589E-05 | 0.00120583 |
| Nras | 977.35776 | 899.0853 | 1.08705788 | 0.12042876 | 938.22153 | 0.12127527 | 0.03029179 | 4.00356894 | 6.2394E-05 | 0.00121998 |
| Tmem258 | 187.443761 | 154.747254 | 1.21128974 | 0.276544 | 171.095507 | 0.27771041 | 0.06938897 | 4.00222682 | 6.2749E-05 | 0.0012253 |
| G2e3 | 964.689166 | 888.701814 | 1.08550377 | 0.11836473 | 926.69549 | 0.11837212 | 0.02958858 | 4.00060222 | 6.3181E-05 | 0.00123212 |
| Kcnk16 | 135.648407 | 166.743548 | 0.81351518 | -0.2977588 | 151.195977 | -0.3009037 | 0.07528851 | -3.9966748 | 6.4238E-05 | 0.00125109 |
| Pde1a | 372.129346 | 325.128621 | 1.1445604 | 0.1947936 | 348.628984 | 0.19359929 | 0.04844646 | 3.99614916 | 6.4381E-05 | 0.00125222 |
| Hspa4 | 2666.53572 | 2503.51599 | 1.06511631 | 0.09101098 | 2585.02586 | 0.09091053 | 0.02275564 | 3.99507717 | 6.4673E-05 | 0.00125625 |
| Npnt | 482.559012 | 429.693414 | 1.12303097 | 0.16739772 | 456.126213 | 0.16931487 | 0.04238433 | 3.9947516 | 6.4762E-05 | 0.00125633 |
| St14 | 1338.24123 | 1438.29086 | 0.93043853 | -0.1040173 | 1388.26604 | -0.1036391 | 0.02597025 | -3.9906861 | 6.5882E-05 | 0.0012764 |
| Atp1a1 | 5464.38348 | 5745.35107 | 0.95109653 | -0.0723363 | 5604.86727 | -0.0736814 | 0.01847067 | -3.9891044 | 6.6323E-05 | 0.00128326 |
| Amfr | 1667.70075 | 1779.69038 | 0.93707353 | -0.0937658 | 1723.69557 | -0.093835 | 0.02352527 | -3.988691 | 6.6439E-05 | 0.00128382 |
| Akap13 | 637.806894 | 706.190059 | 0.90316606 | -0.1469368 | 671.998477 | -0.1537848 | 0.0385598 | -3.9882154 | 6.6572E-05 | 0.00128472 |
| Rps6 | 2630.04264 | 2477.90311 | 1.0613985 | 0.08596641 | 2553.97288 | 0.08640095 | 0.021676 | 3.98601857 | 6.7191E-05 | 0.00129498 |
| Sytl1 | 180.935221 | 216.643413 | 0.83517527 | -0.2598491 | 198.789317 | -0.2632 | 0.06605222 | -3.9847255 | 6.7558E-05 | 0.00130036 |
| Faim2 | 200.88774 | 165.35469 | 1.21488988 | 0.28082555 | 183.121215 | 0.28982273 | 0.07281008 | 3.98053028 | 6.8762E-05 | 0.00132049 |
| Met | 279.674814 | 324.44482 | 0.86201041 | -0.2142228 | 302.059817 | -0.209353 | 0.05259814 | -3.980236 | 6.8847E-05 | 0.00132049 |
| S1pr1 | 185.524111 | 224.563962 | 0.82615264 | -0.2755197 | 205.044036 | -0.2642545 | 0.06639308 | -3.9801521 | 6.8871E-05 | 0.00132049 |
| Tmem255a | 641.364498 | 714.234585 | 0.89797458 | -0.1552535 | 677.799542 | -0.1509437 | 0.03793229 | -3.9792942 | 6.912E-05 | 0.00132185 |
| Ahdc1 | 555.159237 | 616.697575 | 0.9002131 | -0.1516615 | 585.928406 | -0.1515509 | 0.03808354 | -3.9794343 | 6.9079E-05 | 0.00132185 |
| Pttg1 | 481.51808 | 418.842061 | 1.14964118 | 0.20118364 | 450.18007 | 0.20750105 | 0.05214923 | 3.97898578 | 6.921E-05 | 0.00132186 |
| Mapk3 | 1171.02614 | 1263.96213 | 0.92647249 | -0.11018 | 1217.49413 | -0.1098592 | 0.02761612 | -3.9780828 | 6.9473E-05 | 0.00132518 |
| Magoh | 467.521163 | 415.530564 | 1.12511859 | 0.17007708 | 441.525864 | 0.1717309 | 0.04320024 | 3.97522982 | 7.0311E-05 | 0.00133944 |
| Lrrk2 | 494.633761 | 553.17775 | 0.89416785 | -0.1613824 | 523.905755 | -0.1603641 | 0.04035825 | -3.9735154 | 7.082E-05 | 0.00134739 |
| Slc29a3 | 116.974508 | 144.896699 | 0.80729588 | -0.3088306 | 130.935603 | -0.310839 | 0.07829598 | -3.9700502 | 7.1857E-05 | 0.00136538 |
| Naca | 1984.49306 | 1860.47479 | 1.0666589 | 0.0930989 | 1922.48443 | 0.09339666 | 0.02354405 | 3.96689015 | 7.2817E-05 | 0.00138184 |
| Psme3 | 1349.13092 | 1251.62856 | 1.07790039 | 0.10822387 | 1300.37974 | 0.10865134 | 0.02740094 | 3.96524118 | 7.3322E-05 | 0.00138964 |
| S100a11 | 6.78031108 | 14.6249028 | 0.4636141 | -1.1090037 | 10.7026068 | -1.1188521 | 0.28225823 | -3.9639307 | 7.3726E-05 | 0.00139551 |
| Foxa1 | 3176.38538 | 3363.16055 | 0.94446439 | -0.0824317 | 3269.77296 | -0.0818258 | 0.02064526 | -3.9634188 | 7.3884E-05 | 0.00139673 |
| Gpr101 | 38.6721931 | 55.9075589 | 0.69171672 | -0.5317468 | 47.2898759 | -0.5221958 | 0.13177317 | -3.9628385 | 7.4064E-05 | 0.00139834 |
| Rab37 | 77.2958888 | 102.554544 | 0.75370516 | -0.4079278 | 89.9252164 | -0.4070233 | 0.10273061 | -3.9620454 | 7.431E-05 | 0.00140121 |
| Derl3 | 39.0913221 | 25.1699583 | 1.55309443 | 0.63514555 | 32.1306401 | 0.6433038 | 0.16253263 | 3.95799792 | 7.5581E-05 | 0.00142335 |
| Nalcn | 734.60593 | 809.287875 | 0.90771894 | -0.1396824 | 771.946902 | -0.1417597 | 0.03583507 | -3.9558938 | 7.6249E-05 | 0.00143412 |
| Cnnm3 | 421.129508 | 482.121581 | 0.87349234 | -0.195133 | 451.625544 | -0.1948276 | 0.0492803 | -3.953457 | 7.703E-05 | 0.00144698 |
| Sptbn2 | 1633.30803 | 1763.00293 | 0.92643524 | -0.110238 | 1698.15548 | -0.1059361 | 0.0268164 | -3.9504213 | 7.8014E-05 | 0.0014636 |
| Gadd45b | 295.050685 | 339.257412 | 0.86969562 | -0.2014175 | 317.154048 | -0.2038056 | 0.051603 | -3.9494908 | 7.8318E-05 | 0.00146744 |
| Tjp1 | 1233.61421 | 1329.09278 | 0.92816261 | -0.1075505 | 1281.3535 | -0.1086343 | 0.02750832 | -3.9491448 | 7.8431E-05 | 0.00146771 |
| Rpl39 | 661.526622 | 594.083888 | 1.11352392 | 0.15513255 | 627.805255 | 0.1554081 | 0.03938703 | 3.94566675 | 7.9578E-05 | 0.0014873 |
| Snhg4 | 40.0881302 | 26.1690514 | 1.53189084 | 0.6153135 | 33.1285907 | 0.63760706 | 0.16197485 | 3.93645711 | 8.2693E-05 | 0.00154206 |
| Trip12 | 3158.39847 | 3324.74919 | 0.94996593 | -0.0740523 | 3241.57383 | -0.075064 | 0.01906925 | -3.9363911 | 8.2716E-05 | 0.00154206 |
| Mgea5 | 1301.1042 | 1208.29731 | 1.07680799 | 0.10676102 | 1254.70075 | 0.10686396 | 0.0271676 | 3.93350781 | 8.3715E-05 | 0.00155872 |
| Ctnnbip1 | 205.791528 | 241.609784 | 0.85175163 | -0.2314953 | 223.700656 | -0.2327783 | 0.0592187 | -3.9308247 | 8.4655E-05 | 0.00157424 |
| Cdc25b | 1594.28417 | 1491.62449 | 1.06882408 | 0.09602441 | 1542.95433 | 0.09602062 | 0.02444173 | 3.92855342 | 8.5458E-05 | 0.00158719 |
| Prr11 | 765.609429 | 690.546731 | 1.10870039 | 0.14886955 | 728.07808 | 0.15610191 | 0.03974902 | 3.92718896 | 8.5944E-05 | 0.00159422 |
| Pxylp1 | 634.683976 | 698.578311 | 0.90853662 | -0.1383834 | 666.631143 | -0.1382438 | 0.03521863 | -3.9253022 | 8.6621E-05 | 0.00160476 |
| Celf2 | 2026.9891 | 2149.36331 | 0.9430649 | -0.084571 | 2088.17621 | -0.084528 | 0.02153853 | -3.9245007 | 8.691E-05 | 0.0016081 |
| Tmem63a | 408.209708 | 460.193949 | 0.88703841 | -0.1729315 | 434.201828 | -0.1731789 | 0.04414932 | -3.9225722 | 8.7609E-05 | 0.00161872 |
| Cryz | 160.227203 | 198.563439 | 0.80693205 | -0.3094809 | 179.395321 | -0.3082836 | 0.07859733 | -3.9223159 | 8.7702E-05 | 0.00161872 |

| | | | | | | | | | | |
|---|---|---|---|---|---|---|---|---|---|---|
| Podxl | 707.651969 | 777.551527 | 0.91010299 | -0.1358983 | 742.601748 | -0.1364026 | 0.03478336 | -3.9214893 | 8.8003E-05 | 0.00162226 |
| Anxa2 | 284.073089 | 326.795097 | 0.86926974 | -0.2021242 | 305.434093 | -0.2028724 | 0.05175343 | -3.9199796 | 8.8556E-05 | 0.00163043 |
| Akap12 | 200.440079 | 235.396809 | 0.85149871 | -0.2319238 | 217.918444 | -0.2315063 | 0.05906586 | -3.9194612 | 8.8747E-05 | 0.00163191 |
| Actr3 | 3042.06312 | 3194.63481 | 0.95224127 | -0.0706009 | 3118.34897 | -0.0705242 | 0.01800685 | -3.9165231 | 8.9835E-05 | 0.00164987 |
| Rpl10a | 1334.5846 | 1239.65067 | 1.0765812 | 0.10645713 | 1287.11763 | 0.10725076 | 0.02739707 | 3.91467935 | 9.0524E-05 | 0.00166047 |
| Gm2a | 238.518754 | 278.976907 | 0.8549767 | -0.226043 | 258.747831 | -0.2270184 | 0.05803503 | -3.9117483 | 9.163E-05 | 0.00167868 |
| Plekhb1 | 24.9790525 | 39.433341 | 0.63345007 | -0.6586972 | 32.2061966 | -0.6518655 | 0.16666102 | -3.9113257 | 9.1791E-05 | 0.00167955 |
| Disp3 | 34.8906836 | 50.9133113 | 0.6852959 | -0.545201 | 42.9019973 | -0.5475634 | 0.14001392 | -3.9107781 | 9.1999E-05 | 0.00168128 |
| Rpl24 | 1452.16116 | 1352.85616 | 1.07340395 | 0.10219311 | 1402.50866 | 0.10352446 | 0.02648664 | 3.90855331 | 9.285E-05 | 0.00169475 |
| Gpam | 192.015654 | 158.673656 | 1.21012938 | 0.2751613 | 175.344655 | 0.27185579 | 0.06967502 | 3.90176824 | 9.5493E-05 | 0.00174083 |
| Tns3 | 1675.38555 | 1784.93306 | 0.93862654 | -0.0913768 | 1730.1593 | -0.0920192 | 0.02358603 | -3.901427 | 9.5627E-05 | 0.00174115 |
| Nedd4l | 161.187884 | 193.557936 | 0.83276298 | -0.2640222 | 177.37291 | -0.2643804 | 0.06778653 | -3.9001914 | 9.6117E-05 | 0.00174791 |
| Trpc4 | 12.3764544 | 22.6498792 | 0.54642474 | -0.8719053 | 17.5131667 | -0.8677184 | 0.22262247 | -3.8977125 | 9.7106E-05 | 0.00176373 |
| Snrpg | 336.380878 | 294.388827 | 1.14264145 | 0.19237278 | 315.384852 | 0.19233367 | 0.04937706 | 3.8952032 | 9.8116E-05 | 0.00177991 |
| Hnrnpul2 | 1663.89606 | 1536.77346 | 1.08272045 | 0.1146608 | 1600.33476 | 0.11509362 | 0.02957622 | 3.89142424 | 9.9658E-05 | 0.00180345 |
| Rps16 | 1830.53691 | 1711.45407 | 1.06957993 | 0.09704429 | 1770.99549 | 0.09699748 | 0.0249251 | 3.89155797 | 9.9603E-05 | 0.00180345 |
| Srsf11 | 1187.49029 | 1097.62965 | 1.08186791 | 0.11352436 | 1142.55997 | 0.11554997 | 0.02969769 | 3.89087456 | 9.9884E-05 | 0.00180533 |
| Frem1 | 22.2514962 | 35.7264086 | 0.62283048 | -0.6830885 | 28.9889523 | -0.6843918 | 0.17597608 | -3.8891183 | 0.00010061 | 0.00181623 |
| Msmo1 | 319.225682 | 364.930718 | 0.87475695 | -0.1930459 | 342.0782 | -0.1920711 | 0.04941487 | -3.8869087 | 0.00010153 | 0.00183061 |
| Ttc14 | 580.086601 | 513.732159 | 1.12916155 | 0.17525191 | 546.90938 | 0.17815985 | 0.04584966 | 3.88574013 | 0.00010202 | 0.0018372 |
| Stx1b | 9.09618017 | 3.29181873 | 2.76326886 | 1.46637594 | 6.19399935 | 1.4517356 | 0.37376056 | 3.88413266 | 0.0001027 | 0.00184644 |
| Nudcd2 | 296.355136 | 256.01582 | 1.15756571 | 0.2110941 | 276.185478 | 0.21558914 | 0.05550797 | 3.8839314 | 0.00010278 | 0.00184644 |
| Tmem151a | 3.6320125 | 0.35412494 | 10.2563024 | 3.35843879 | 1.99306862 | 3.20109632 | 0.82442817 | 3.88280805 | 0.00010326 | 0.00185275 |
| R3hdm2 | 1111.96018 | 1196.48207 | 0.92935799 | -0.1056937 | 1154.22112 | -0.1049674 | 0.02704184 | -3.8816656 | 0.00010374 | 0.00185923 |
| Macrod2 | 279.922946 | 321.1309 | 0.87167864 | -0.1981317 | 300.526923 | -0.1982576 | 0.05110803 | -3.8791863 | 0.00010481 | 0.00187601 |
| Insm1 | 4147.026 | 4353.53405 | 0.95256542 | -0.0701099 | 4250.28003 | -0.0703891 | 0.01814689 | -3.8788536 | 0.00010495 | 0.00187631 |
| Pclaf | 1756.32154 | 1650.09252 | 1.06437761 | 0.09001007 | 1703.20703 | 0.0924179 | 0.02384233 | 3.87621147 | 0.0001061 | 0.00189421 |
| Fdft1 | 788.146979 | 861.764873 | 0.91457311 | -0.1288296 | 824.955926 | -0.1283469 | 0.0331136 | -3.8759569 | 0.00010621 | 0.00189421 |
| Malat1 | 1900.8322 | 1193.39991 | 1.59278728 | 0.67155361 | 1547.11605 | 0.53762283 | 0.13884597 | 3.87208086 | 0.00010791 | 0.00192228 |
| Cbln4 | 5.18382412 | 0.78872889 | 6.5723776 | 2.71641537 | 2.98627641 | 2.70472356 | 0.69910539 | 3.8688352 | 0.00010936 | 0.00194571 |
| Bag6 | 2389.08298 | 2529.93515 | 0.94432578 | -0.0826434 | 2459.50907 | -0.0828178 | 0.02141449 | -3.8673721 | 0.00011001 | 0.00195507 |
| Pcdh15 | 51.0805252 | 35.5286851 | 1.43772631 | 0.52378906 | 43.304605 | 0.51834532 | 0.1340835 | 3.86583965 | 0.00011071 | 0.00196503 |
| Fth1 | 1105.91537 | 1022.44065 | 1.0816426 | 0.11322388 | 1064.17801 | 0.11319865 | 0.02932472 | 3.86017791 | 0.0001133 | 0.00200872 |
| Gstm7 | 200.373285 | 236.836722 | 0.84603977 | -0.2412026 | 218.605004 | -0.2369429 | 0.06139284 | -3.8594544 | 0.00011364 | 0.00201226 |
| Igf1r | 739.452414 | 811.089487 | 0.91167797 | -0.1334038 | 775.27095 | -0.1353645 | 0.03508556 | -3.8581261 | 0.00011426 | 0.00202082 |
| Emc1 | 506.944982 | 563.758984 | 0.89922289 | -0.1532493 | 535.351983 | -0.1526698 | 0.03958919 | -3.8563521 | 0.00011509 | 0.0020331 |
| Snrpb2 | 401.391726 | 354.899083 | 1.13100243 | 0.17760203 | 378.145404 | 0.17746213 | 0.0460327 | 3.85513221 | 0.00011567 | 0.00204084 |
| Robo1 | 7502.87962 | 7780.17351 | 0.96435891 | -0.0523579 | 7641.52656 | -0.0523616 | 0.01358957 | -3.8530743 | 0.00011664 | 0.00205352 |
| Tnrc18 | 1214.30474 | 1311.53492 | 0.92586536 | -0.1111257 | 1262.91983 | -0.112097 | 0.02909317 | -3.8530345 | 0.00011666 | 0.00205352 |
| Mphosph10 | 241.090878 | 205.743373 | 1.17180385 | 0.2287311 | 223.417125 | 0.22873454 | 0.05939405 | 3.8511356 | 0.00011757 | 0.0020646 |
| Gng2 | 926.429505 | 1028.89789 | 0.90040957 | -0.1513467 | 977.663698 | -0.1501022 | 0.03897465 | -3.8512763 | 0.0001175 | 0.0020646 |
| G3bp1 | 1852.38666 | 1727.11552 | 1.072532 | 0.10102069 | 1789.75109 | 0.10037967 | 0.02607597 | 3.84950802 | 0.00011836 | 0.00207591 |
| Cpne7 | 119.567834 | 94.7883199 | 1.26141949 | 0.33504813 | 107.178077 | 0.33398585 | 0.08684238 | 3.84588564 | 0.00012012 | 0.00210185 |
| Rph3al | 510.590347 | 566.585973 | 0.90117012 | -0.1501286 | 538.58816 | -0.1495387 | 0.03888188 | -3.8459739 | 0.00012007 | 0.00210185 |
| Ncam2 | 391.576897 | 344.770836 | 1.13575992 | 0.1836579 | 368.173866 | 0.18292859 | 0.04759184 | 3.84369684 | 0.00012119 | 0.00211819 |
| Slc35g1 | 195.227104 | 163.98652 | 1.19050702 | 0.25157613 | 179.606812 | 0.2541577 | 0.06614356 | 3.84251607 | 0.00012178 | 0.00212548 |
| Phldb1 | 312.190746 | 356.574302 | 0.87552789 | -0.191775 | 334.382524 | -0.1912849 | 0.04978427 | -3.8422766 | 0.0001219 | 0.00212548 |
| Parva | 1102.84634 | 1196.43587 | 0.9217764 | -0.1175113 | 1149.6411 | -0.117153 | 0.03052615 | -3.8377898 | 0.00012415 | 0.00216214 |
| Nrd1 | 1833.18054 | 1726.18694 | 1.06198263 | 0.08676017 | 1779.68374 | 0.08661962 | 0.02258791 | 3.83477695 | 0.00012568 | 0.00218624 |
| Cks2 | 611.294549 | 547.808806 | 1.11589033 | 0.15819524 | 579.551677 | 0.16289122 | 0.0425149 | 3.83139135 | 0.00012742 | 0.00221395 |
| Tex101 | 2.64547203 | 0.10938243 | 24.1855291 | 4.5960722 | 1.37742713 | 3.56219945 | 0.92986017 | 3.83089799 | 0.00012768 | 0.0022158 |
| Add1 | 2109.91227 | 2251.15844 | 0.93725623 | -0.0934846 | 2180.53536 | -0.0920866 | 0.02404538 | -3.8297015 | 0.0001283 | 0.002224 |
| Fam184a | 73.8114994 | 95.2623952 | 0.77482305 | -0.3680612 | 84.5369472 | -0.3652098 | 0.09539197 | -3.8285176 | 0.00012892 | 0.00223211 |
| Glis3 | 218.147614 | 254.49007 | 0.85719499 | -0.2223047 | 236.318842 | -0.2223981 | 0.0582046 | -3.8209716 | 0.00013293 | 0.00229885 |
| Senp5 | 612.225367 | 549.364535 | 1.11442463 | 0.15629904 | 580.794951 | 0.15651277 | 0.04100539 | 3.81688264 | 0.00013515 | 0.00233454 |
| Mad2l1 | 736.735684 | 669.952404 | 1.09968362 | 0.13708852 | 703.344044 | 0.14169205 | 0.03713828 | 3.81525623 | 0.00013604 | 0.00234723 |
| Selenot | 2181.79904 | 2055.39189 | 1.06150027 | 0.08610473 | 2118.59547 | 0.08584375 | 0.02253075 | 3.81007019 | 0.00013893 | 0.00239424 |
| Klhl29 | 223.507115 | 261.334793 | 0.85525204 | -0.2255785 | 242.420954 | -0.2242647 | 0.05886698 | -3.8096867 | 0.00013914 | 0.00239517 |
| Nell2 | 8.81633336 | 2.84876709 | 3.09478911 | 1.6298411 | 5.83255013 | 1.56203657 | 0.4101152 | 3.80877517 | 0.00013966 | 0.00240122 |
| Tpx2 | 3352.08575 | 3127.79192 | 1.07170996 | 0.09991452 | 3239.93884 | 0.10451001 | 0.0274569 | 3.80633012 | 0.00014104 | 0.00242226 |
| Ubr4 | 3065.76907 | 3390.56705 | 0.90420541 | -0.1452775 | 3228.16806 | -0.1688673 | 0.04439671 | -3.8035999 | 0.00014261 | 0.0024463 |
| Anxa4 | 1129.81227 | 1213.47762 | 0.93105324 | -0.1030644 | 1171.64494 | -0.1032367 | 0.02715204 | -3.8021713 | 0.00014343 | 0.0024576 |
| Kcnq4 | 146.133174 | 175.680164 | 0.83181374 | -0.2656676 | 160.906669 | -0.264708 | 0.06962662 | -3.8018215 | 0.00014364 | 0.00245824 |
| Ctnnd2 | 686.274938 | 766.476012 | 0.89536388 | -0.159454 | 726.375475 | -0.152627 | 0.04015921 | -3.800547 | 0.00014438 | 0.00246806 |
| Vat1 | 762.345728 | 836.826175 | 0.91099651 | -0.1344826 | 799.585952 | -0.1339289 | 0.03528039 | -3.7961301 | 0.00014697 | 0.00250954 |
| Spen | 1125.07084 | 1214.80908 | 0.92612976 | -0.1107138 | 1169.93996 | -0.1106068 | 0.02917164 | -3.7915882 | 0.00014969 | 0.00255295 |
| Megf8 | 1129.31226 | 1215.56456 | 0.92904343 | -0.1061821 | 1172.43841 | -0.1074055 | 0.02833534 | -3.7905135 | 0.00015034 | 0.00256107 |

| | | | | | | | | | | |
|---|---|---|---|---|---|---|---|---|---|---|
| Cldn6 | 468.930794 | 523.540884 | 0.89569088 | -0.1589272 | 496.235839 | -0.1586041 | 0.04184821 | -3.7899862 | 0.00015066 | 0.00256357 |
| Phlda1 | 35.7240185 | 23.4285618 | 1.52480629 | 0.60862597 | 29.57629 | 0.60884898 | 0.1607251 | 3.78813863 | 0.00015178 | 0.00257974 |
| Tmem117 | 768.839779 | 847.303746 | 0.9073957 | -0.1401963 | 808.071763 | -0.1402181 | 0.03702411 | -3.787211 | 0.00015235 | 0.00258622 |
| Spata13 | 41.1819052 | 57.6718672 | 0.71407269 | -0.4858572 | 49.4268861 | -0.4841986 | 0.1278599 | -3.7869462 | 0.00015251 | 0.00258622 |
| Ints1 | 1009.95295 | 1109.73229 | 0.91008702 | -0.1359236 | 1059.84262 | -0.1377596 | 0.03638733 | -3.7859217 | 0.00015314 | 0.00259393 |
| Cldn10 | 9.33293259 | 3.48834761 | 2.67545945 | 1.41978666 | 6.41064 | 1.3837398 | 0.36605943 | 3.78009606 | 0.00015677 | 0.00265235 |
| Vat1l | 1052.74139 | 1135.23445 | 0.9273339 | -0.1088392 | 1093.98792 | -0.108943 | 0.02885 | -3.7761851 | 0.00015925 | 0.00269125 |
| Manba | 563.013396 | 626.05898 | 0.89929769 | -0.1531293 | 594.536188 | -0.1482725 | 0.0392744 | -3.7752958 | 0.00015982 | 0.00269472 |
| Pcsk1n | 989.859476 | 1154.04328 | 0.85773168 | -0.2214017 | 1071.95138 | -0.2191913 | 0.05805647 | -3.7754838 | 0.0001597 | 0.00269472 |
| Cdc42bpa | 1957.75158 | 2087.6184 | 0.93779187 | -0.0926603 | 2022.68499 | -0.0933505 | 0.02473122 | -3.7746026 | 0.00016026 | 0.00269916 |
| Eif4h | 2975.02761 | 2832.5823 | 1.05028815 | 0.07078518 | 2903.80495 | 0.07090075 | 0.01878762 | 3.77380092 | 0.00016078 | 0.00270477 |
| Cbarp | 570.389648 | 632.349103 | 0.90201701 | -0.1487735 | 601.369375 | -0.1469026 | 0.03895673 | -3.7709157 | 0.00016265 | 0.00273314 |
| Ttc9c | 279.311933 | 240.58217 | 1.16098351 | 0.21534749 | 259.947052 | 0.215198 | 0.05707717 | 3.77029943 | 0.00016305 | 0.0027368 |
| Eno2 | 267.284091 | 307.879003 | 0.86814654 | -0.2039895 | 287.581547 | -0.2023435 | 0.05367613 | -3.7697115 | 0.00016344 | 0.00274015 |
| Akap1 | 539.199662 | 480.73725 | 1.12160991 | 0.165571 | 509.968456 | 0.16670458 | 0.04422862 | 3.76915654 | 0.0001638 | 0.00274314 |
| Zwint | 870.132388 | 801.252874 | 1.08596476 | 0.11897729 | 835.692631 | 0.11911231 | 0.03161737 | 3.76730652 | 0.00016502 | 0.00276043 |
| Pik3cb | 334.858465 | 380.939679 | 0.87903278 | -0.1860111 | 357.899072 | -0.1860551 | 0.049406 | -3.765841 | 0.00016599 | 0.00277354 |
| Arpc5 | 1648.03162 | 1756.69263 | 0.93814455 | -0.0921179 | 1702.36213 | -0.0926434 | 0.02463917 | -3.7600051 | 0.00016991 | 0.00283585 |
| Cacnb4 | 126.590031 | 102.766536 | 1.23182153 | 0.30079325 | 114.678283 | 0.30155412 | 0.08026573 | 3.75694709 | 0.000172 | 0.00286749 |
| Sema3d | 203.866988 | 244.409161 | 0.83412171 | -0.2616702 | 224.138074 | -0.2516258 | 0.06705994 | -3.7522521 | 0.00017525 | 0.00291846 |
| Osbp | 573.591687 | 511.29214 | 1.12184726 | 0.16587627 | 542.441913 | 0.16413195 | 0.04377817 | 3.74917349 | 0.00017742 | 0.00294795 |
| Ehmt2 | 1254.09787 | 1344.65819 | 0.93265179 | -0.1005895 | 1299.37803 | -0.1007039 | 0.02686034 | -3.7491656 | 0.00017742 | 0.00294795 |
| Colgalt2 | 81.7394299 | 104.056571 | 0.78552877 | -0.348264 | 92.8980004 | -0.3582308 | 0.09555658 | -3.7488874 | 0.00017762 | 0.00294795 |
| Zfp91 | 814.716095 | 730.732752 | 1.11493031 | 0.15695354 | 772.724423 | 0.15343893 | 0.0409359 | 3.74827266 | 0.00017806 | 0.00295188 |
| Jpt1 | 2019.51542 | 1876.43458 | 1.07625144 | 0.10601517 | 1947.975 | 0.11155719 | 0.02979142 | 3.74460793 | 0.00018068 | 0.00299082 |
| Kpnb1 | 4019.96779 | 3829.73958 | 1.04967132 | 0.06993765 | 3924.85369 | 0.06948637 | 0.0185573 | 3.74442261 | 0.00018081 | 0.00299082 |
| Trp53i11 | 1491.40742 | 1396.53294 | 1.06793573 | 0.09482482 | 1443.97018 | 0.09532702 | 0.02547695 | 3.74169637 | 0.00018278 | 0.00302008 |
| Gnb2 | 3707.72547 | 3878.82032 | 0.95588998 | -0.0650835 | 3793.2729 | -0.0659511 | 0.01763163 | -3.7405005 | 0.00018365 | 0.00303111 |
| Foxo6 | 221.895171 | 257.719189 | 0.86099592 | -0.2159217 | 239.80718 | -0.2159404 | 0.05779558 | -3.7362796 | 0.00018676 | 0.00307899 |
| Esf1 | 319.385074 | 279.817042 | 1.1414068 | 0.19081306 | 299.601058 | 0.19162232 | 0.05133084 | 3.73308351 | 0.00018915 | 0.00311488 |
| Rangrf | 101.683002 | 80.420931 | 1.26438479 | 0.33843558 | 91.0519663 | 0.33851756 | 0.0906879 | 3.73277518 | 0.00018938 | 0.00311523 |
| Gprc5b | 459.927894 | 512.391065 | 0.89761107 | -0.1558376 | 486.15948 | -0.1552997 | 0.0416162 | -3.7317134 | 0.00019018 | 0.00312493 |
| Morf4l2 | 2726.2851 | 2586.76462 | 1.05393629 | 0.07578766 | 2656.52486 | 0.07570245 | 0.02029325 | 3.73042513 | 0.00019116 | 0.00313217 |
| Smg1 | 2247.7144 | 2376.95328 | 0.94562835 | -0.0806548 | 2312.33384 | -0.0870807 | 0.02334593 | -3.7300155 | 0.00019147 | 0.00313217 |
| 1190002N15 | 578.777886 | 642.77865 | 0.9004311 | -0.1513122 | 610.778268 | -0.1507189 | 0.04040072 | -3.7305997 | 0.00019102 | 0.00313217 |
| Klhl5 | 245.035899 | 287.05485 | 0.85362048 | -0.2283333 | 266.045375 | -0.2244845 | 0.06018175 | -3.730109 | 0.0001914 | 0.00313217 |
| Scn7a | 31.9306667 | 20.2700832 | 1.57526076 | 0.65559067 | 26.1003748 | 0.67729839 | 0.18163437 | 3.72891105 | 0.00019231 | 0.00314246 |
| Ctsf | 796.419369 | 899.265914 | 0.88563278 | -0.1752195 | 847.842641 | -0.1663257 | 0.04462197 | -3.7274399 | 0.00019343 | 0.00315737 |
| Psma5 | 631.988229 | 571.932831 | 1.10500429 | 0.14405196 | 601.96053 | 0.14546913 | 0.03904062 | 3.72609717 | 0.00019447 | 0.00316726 |
| Cnn3 | 578.768712 | 524.20483 | 1.10408886 | 0.14285629 | 551.486771 | 0.14295574 | 0.03836475 | 3.72622626 | 0.00019437 | 0.00316726 |
| Sstr2 | 80.0660364 | 102.25284 | 0.78302017 | -0.3528786 | 91.159438 | -0.3475463 | 0.09335901 | -3.722686 | 0.00019711 | 0.00320684 |
| Rps6kb2 | 374.6003 | 423.642535 | 0.88423675 | -0.1774954 | 399.121417 | -0.1758442 | 0.04724843 | -3.7216937 | 0.00019789 | 0.00321595 |
| Ckmt1 | 586.994722 | 651.375829 | 0.90116135 | -0.1501427 | 619.185275 | -0.1490031 | 0.04007524 | -3.7180843 | 0.00020074 | 0.00325866 |
| Capza1 | 1592.01061 | 1496.91273 | 1.06352935 | 0.08885984 | 1544.46167 | 0.08942398 | 0.02408064 | 3.71352123 | 0.0002044 | 0.00331438 |
| Mbtd1 | 438.233653 | 380.892119 | 1.15054534 | 0.20231784 | 409.562886 | 0.20062197 | 0.05405927 | 3.71114837 | 0.00020632 | 0.00334195 |
| Senp2 | 797.474667 | 728.414651 | 1.09480866 | 0.13067875 | 762.944659 | 0.13075508 | 0.0352506 | 3.70930082 | 0.00020783 | 0.00336273 |
| Ptprt | 2184.87668 | 2322.95231 | 0.94056028 | -0.0884077 | 2253.9145 | -0.0842609 | 0.02271778 | -3.7090267 | 0.00020806 | 0.00336273 |
| Tmem167 | 753.113183 | 687.656258 | 1.09518844 | 0.13117912 | 720.38472 | 0.13195437 | 0.03560473 | 3.70609083 | 0.00021048 | 0.00339822 |
| Bace2 | 388.300605 | 443.385384 | 0.8757632 | -0.1913873 | 415.842994 | -0.1799074 | 0.04855303 | -3.7053794 | 0.00021107 | 0.00340407 |
| L2hgdh | 233.548495 | 199.485213 | 1.17075592 | 0.22744034 | 216.516854 | 0.22764274 | 0.06152531 | 3.69998529 | 0.00021561 | 0.00346969 |
| Ccdc148 | 147.462148 | 177.470479 | 0.83091086 | -0.2672344 | 162.466314 | -0.2662753 | 0.07196649 | -3.6999903 | 0.00021561 | 0.00346969 |
| Atp5j | 889.611338 | 822.292567 | 1.08186718 | 0.11352339 | 855.951952 | 0.1142195 | 0.03087643 | 3.69924523 | 0.00021624 | 0.00347228 |
| Nxph3 | 78.8908115 | 101.356583 | 0.77834916 | -0.3615106 | 90.123697 | -0.3558771 | 0.09619869 | -3.699397 | 0.00021611 | 0.00347228 |
| Ncstn | 1485.68439 | 1592.98137 | 0.93264392 | -0.1006017 | 1539.33288 | -0.1012874 | 0.02740643 | -3.6957525 | 0.00021924 | 0.00351656 |
| Cachd1 | 19.5986366 | 32.1544583 | 0.60951537 | -0.7142655 | 25.8765474 | -0.6840371 | 0.18511609 | -3.6951791 | 0.00021973 | 0.0035207 |
| Yes1 | 308.296251 | 269.809788 | 1.14264294 | 0.19237466 | 289.053019 | 0.19261326 | 0.05215669 | 3.69297355 | 0.00022165 | 0.00354755 |
| Cbx6 | 2899.7651 | 3046.15863 | 0.95194159 | -0.071055 | 2972.96186 | -0.0710746 | 0.01925019 | -3.6921486 | 0.00022237 | 0.00355524 |
| Chrnb4 | 72.8109026 | 54.5538177 | 1.33466191 | 0.41647433 | 63.6823601 | 0.42319417 | 0.11463159 | 3.69177608 | 0.00022269 | 0.00355662 |
| Atp1b2 | 268.113575 | 318.331625 | 0.84224612 | -0.2476862 | 293.2226 | -0.233027 | 0.0631811 | -3.6882395 | 0.00022581 | 0.00360252 |
| Opn3 | 100.875169 | 126.763293 | 0.79577587 | -0.3295659 | 113.819231 | -0.3267566 | 0.0886282 | -3.6868238 | 0.00022707 | 0.00361872 |
| Hmgb3 | 1212.96296 | 1117.86273 | 1.08507326 | 0.11779246 | 1165.41284 | 0.12242642 | 0.03320907 | 3.68653562 | 0.00022733 | 0.00361893 |
| Gna14 | 99.5971949 | 78.8419939 | 1.26325058 | 0.33714085 | 89.2195943 | 0.36262675 | 0.09838251 | 3.6858865 | 0.00022791 | 0.00362427 |
| Plgrkt | 77.9808577 | 58.7556609 | 1.32720586 | 0.40839217 | 68.3682592 | 0.40518521 | 0.11002102 | 3.68279803 | 0.00023069 | 0.00366455 |
| Qars | 351.605141 | 306.401742 | 1.14752984 | 0.19853167 | 329.003442 | 0.19778868 | 0.05373892 | 3.68054777 | 0.00023273 | 0.00369309 |
| Becn1 | 1279.1523 | 1193.62435 | 1.07165399 | 0.09983917 | 1236.38833 | 0.09987415 | 0.02715048 | 3.67854026 | 0.00023457 | 0.00371829 |
| Polr2g | 381.100942 | 336.438893 | 1.13274936 | 0.17982868 | 358.769917 | 0.17975538 | 0.04888923 | 3.67678914 | 0.00023619 | 0.0037399 |
| Arl4c | 401.859647 | 449.035258 | 0.89494007 | -0.160137 | 425.447452 | -0.1599161 | 0.04349812 | -3.6763908 | 0.00023656 | 0.00374174 |

| | | | | | | | | | | |
|---|---|---|---|---|---|---|---|---|---|---|
| Dnm1 | 410.099245 | 463.657353 | 0.88448774 | -0.1770859 | 436.878299 | -0.1700941 | 0.04627955 | -3.6753629 | 0.00023751 | 0.00375283 |
| D430019H16 | 1083.2987 | 1165.0134 | 0.92985944 | -0.1049155 | 1124.15605 | -0.1053947 | 0.02872155 | -3.6695318 | 0.000243 | 0.00383539 |
| Stk11ip | 741.338876 | 805.641641 | 0.92018441 | -0.1200051 | 773.490258 | -0.120078 | 0.03274562 | -3.6669949 | 0.00024542 | 0.00386951 |
| Spred2 | 257.487995 | 221.119194 | 1.164476 | 0.21968091 | 239.303595 | 0.2210797 | 0.0604399 | 3.65784334 | 0.00025435 | 0.00400177 |
| Grid2 | 155.461774 | 186.055823 | 0.83556521 | -0.2591757 | 170.758798 | -0.2544566 | 0.06956176 | -3.6579949 | 0.0002542 | 0.00400177 |
| Slc35b1 | 385.778024 | 334.622469 | 1.15287543 | 0.20523664 | 360.200247 | 0.20551859 | 0.05619062 | 3.65752448 | 0.00025466 | 0.0040025 |
| Ntn1 | 56.4484104 | 41.1577451 | 1.37151368 | 0.45576901 | 48.8030777 | 0.45520497 | 0.12448662 | 3.65665784 | 0.00025553 | 0.00400755 |
| Arhgef3 | 706.052911 | 769.304665 | 0.91778062 | -0.1237788 | 737.678788 | -0.12374 | 0.03383889 | -3.656739 | 0.00025544 | 0.00400755 |
| Epas1 | 350.774796 | 402.362163 | 0.87178872 | -0.1979496 | 376.568479 | -0.1957975 | 0.05363546 | -3.6505223 | 0.00026171 | 0.00410018 |
| Ptchd1 | 324.229672 | 367.176968 | 0.8830338 | -0.1794594 | 345.70332 | -0.1792006 | 0.0491096 | -3.6489923 | 0.00026327 | 0.00412032 |
| Sf3b1 | 3826.95258 | 4010.90408 | 0.95413715 | -0.0677314 | 3918.92833 | -0.0671949 | 0.01841606 | -3.6487096 | 0.00026356 | 0.0041205 |
| Etl4 | 361.347373 | 405.748737 | 0.89056931 | -0.1672002 | 383.548055 | -0.1659363 | 0.04549217 | -3.6475797 | 0.00026472 | 0.0041343 |
| Ndufv1 | 2095.43925 | 2224.78747 | 0.94186042 | -0.0864148 | 2160.11336 | -0.0862489 | 0.023651 | -3.6467352 | 0.00026559 | 0.00414354 |
| Cacnb3 | 589.692644 | 649.254136 | 0.90826167 | -0.1388201 | 619.47339 | -0.1404147 | 0.03850962 | -3.6462248 | 0.00026612 | 0.00414741 |
| Carmil1 | 1400.60007 | 1495.71198 | 0.93641028 | -0.0947873 | 1448.15603 | -0.0944725 | 0.0259195 | -3.6448406 | 0.00026756 | 0.00416541 |
| Kctd15 | 40.5232017 | 26.9881272 | 1.50151959 | 0.5864233 | 33.7556644 | 0.6239191 | 0.17126959 | 3.64290643 | 0.00026958 | 0.00418366 |
| B3gat1 | 224.077626 | 190.180374 | 1.17823738 | 0.23663023 | 207.129 | 0.2340336 | 0.06424027 | 3.64309825 | 0.00026938 | 0.00418366 |
| Gpr137b-ps | 103.546599 | 129.1546 | 0.80172598 | -0.3188189 | 116.350599 | -0.3186154 | 0.0874536 | -3.6432509 | 0.00026922 | 0.00418366 |
| Plcb4 | 1998.13969 | 1885.13667 | 1.05994421 | 0.08398833 | 1941.63818 | 0.08431052 | 0.02317273 | 3.63835112 | 0.00027439 | 0.00425389 |
| Vps13b | 889.242976 | 959.549645 | 0.92672951 | -0.1097798 | 924.39631 | -0.1114376 | 0.03063204 | -3.6379429 | 0.00027482 | 0.00425619 |
| Tmem131 | 1796.05241 | 1906.40149 | 0.94211656 | -0.0860225 | 1851.22695 | -0.0871535 | 0.02397121 | -3.6357559 | 0.00027717 | 0.00428183 |
| Arhgef17 | 1381.65335 | 1481.61769 | 0.93253027 | -0.1007775 | 1431.63552 | -0.1004784 | 0.02763745 | -3.6355888 | 0.00027735 | 0.00428183 |
| Arl8a | 863.348489 | 938.644843 | 0.91978185 | -0.1206364 | 900.996666 | -0.120121 | 0.03303898 | -3.6357376 | 0.00027719 | 0.00428183 |
| Asxl3 | 219.84917 | 256.669247 | 0.8565466 | -0.2233964 | 238.259209 | -0.224436 | 0.06178873 | -3.6323122 | 0.00028089 | 0.00433207 |
| Dnajb4 | 27.5877385 | 17.3607035 | 1.58909105 | 0.66820179 | 22.4742209 | 0.66893915 | 0.18422436 | 3.63111132 | 0.0002822 | 0.00434775 |
| Gsn | 546.512061 | 607.549266 | 0.89953538 | -0.1527481 | 577.030663 | -0.1553725 | 0.04280351 | -3.6299013 | 0.00028353 | 0.00436365 |
| Mfsd12 | 326.009721 | 366.999734 | 0.88831051 | -0.170864 | 346.504728 | -0.1710883 | 0.04716914 | -3.6271233 | 0.0002866 | 0.00440519 |
| Gpc4 | 4.23358829 | 10.2920198 | 0.41134669 | -1.2815733 | 7.26280395 | -1.3777923 | 0.37987953 | -3.6269191 | 0.00028682 | 0.00440519 |
| Hoxb5 | 638.958221 | 579.086876 | 1.10338923 | 0.1419418 | 609.022548 | 0.14347336 | 0.03957106 | 3.62571453 | 0.00028816 | 0.00441662 |
| Rps9 | 2137.34004 | 2001.30376 | 1.06797383 | 0.0948763 | 2069.3219 | 0.0970505 | 0.02676548 | 3.62595818 | 0.00028789 | 0.00441662 |
| Nceh1 | 565.027315 | 626.086979 | 0.90247415 | -0.1480425 | 595.557147 | -0.1481725 | 0.04087434 | -3.6250731 | 0.00028888 | 0.00442302 |
| Ppp6r3 | 1043.30503 | 951.081295 | 1.09696725 | 0.13352045 | 997.193163 | 0.13295624 | 0.03673798 | 3.61904086 | 0.0002957 | 0.00452273 |
| Poc1a | 131.931954 | 107.581517 | 1.22634406 | 0.29436379 | 119.756736 | 0.29240306 | 0.08082502 | 3.61772939 | 0.0002972 | 0.00454077 |
| Cnnm4 | 542.924703 | 597.564346 | 0.90856275 | -0.1383419 | 570.244524 | -0.1385541 | 0.03830133 | -3.6174765 | 0.00029749 | 0.00454077 |
| Tle3 | 998.530237 | 921.329029 | 1.08379331 | 0.11608965 | 959.929633 | 0.11698265 | 0.03237682 | 3.61316051 | 0.00030249 | 0.00461231 |
| Tubb5 | 14516.4827 | 15002.4643 | 0.96760655 | -0.0475076 | 14759.4735 | -0.0463513 | 0.01283087 | -3.612485 | 0.00030328 | 0.00461959 |
| Ganab | 1175.0725 | 1074.3769 | 1.09372465 | 0.12924957 | 1124.7247 | 0.12954949 | 0.0358751 | 3.6111258 | 0.00030487 | 0.00463911 |
| Trove2 | 4406.07601 | 4599.57071 | 0.95793201 | -0.0620048 | 4502.82336 | -0.0635161 | 0.01759503 | -3.6098896 | 0.00030633 | 0.00465649 |
| Lrrc56 | 108.305174 | 132.11304 | 0.8197917 | -0.2866707 | 120.209107 | -0.2867818 | 0.07946371 | -3.6089653 | 0.00030742 | 0.00466832 |
| Cbx1 | 478.13935 | 425.724178 | 1.12312003 | 0.16751212 | 451.931764 | 0.16752079 | 0.04642884 | 3.60811931 | 0.00030842 | 0.00467877 |
| Ptprn2 | 1372.86124 | 1476.19168 | 0.93000202 | -0.1046942 | 1424.52646 | -0.1005496 | 0.02787308 | -3.6074085 | 0.00030927 | 0.00468681 |
| Deup1 | 132.098524 | 160.31143 | 0.82401189 | -0.2792629 | 146.204976 | -0.2764223 | 0.07663658 | -3.6069241 | 0.00030985 | 0.00469077 |
| Dctn1 | 1730.26172 | 1832.90043 | 0.94400203 | -0.0831381 | 1781.58107 | -0.0829889 | 0.02302208 | -3.6047515 | 0.00031245 | 0.00472535 |
| Ap1b1 | 939.635203 | 1014.58189 | 0.92613047 | -0.1107126 | 977.108544 | -0.1108286 | 0.03076222 | -3.6027501 | 0.00031487 | 0.00475705 |
| Dcaf6 | 649.719639 | 726.888724 | 0.89383645 | -0.1619172 | 688.304182 | -0.1588325 | 0.04409182 | -3.602313 | 0.0003154 | 0.0047602 |
| Wee1 | 417.486025 | 369.728968 | 1.12916775 | 0.17525983 | 393.607496 | 0.17474472 | 0.04851977 | 3.60151561 | 0.00031637 | 0.00476997 |
| Snx10 | 19.3499148 | 10.8300224 | 1.78669204 | 0.83729099 | 15.0899685 | 0.86365751 | 0.24016299 | 3.59613067 | 0.00032299 | 0.00485987 |
| Gtpbp4 | 606.099466 | 551.703402 | 1.09859657 | 0.1356617 | 578.901434 | 0.13488508 | 0.03750714 | 3.59625052 | 0.00032284 | 0.00485987 |
| Dsel | 454.621108 | 511.059375 | 0.88956613 | -0.1688262 | 482.840241 | -0.1700226 | 0.04728679 | -3.5955628 | 0.00032369 | 0.00486554 |
| Cntnap4 | 271.325519 | 308.813377 | 0.87860676 | -0.1867105 | 290.069448 | -0.1869467 | 0.05201784 | -3.5938969 | 0.00032577 | 0.00489181 |
| Sh3bp4 | 832.58173 | 911.567262 | 0.91335194 | -0.1307572 | 872.074496 | -0.1284559 | 0.03574749 | -3.5934245 | 0.00032636 | 0.00489573 |
| Cdk2ap2 | 692.587648 | 755.642387 | 0.91655479 | -0.125707 | 724.115017 | -0.1290916 | 0.03593732 | -3.5921314 | 0.00032798 | 0.00491136 |
| Galnt12 | 283.228372 | 321.428836 | 0.88115421 | -0.1825336 | 302.328604 | -0.1831618 | 0.05099062 | -3.5920677 | 0.00032806 | 0.00491136 |
| A730017C20 | 27.7326143 | 43.1341859 | 0.64293816 | -0.6372481 | 35.4334 | -0.6262377 | 0.1743875 | -3.5910698 | 0.00032932 | 0.00492523 |
| Alpl | 7.03334536 | 2.44012296 | 2.88237334 | 1.52725721 | 4.73673406 | 1.5607193 | 0.43466243 | 3.59064685 | 0.00032986 | 0.00492579 |
| Retreg2 | 1636.98545 | 1735.38842 | 0.94329629 | -0.0842171 | 1686.18693 | -0.084265 | 0.02346879 | -3.5905144 | 0.00033003 | 0.00492579 |
| Podxl2 | 180.695618 | 212.216205 | 0.85146946 | -0.2319733 | 196.455912 | -0.2329163 | 0.06490098 | -3.5887944 | 0.00033221 | 0.00495341 |
| Gpr137b | 87.7535117 | 110.339766 | 0.79530268 | -0.3304241 | 99.0466386 | -0.3303428 | 0.09211895 | -3.5860464 | 0.00033573 | 0.00500085 |
| Osbpl2 | 829.018163 | 895.200017 | 0.92607032 | -0.1108064 | 862.10909 | -0.1109799 | 0.03095007 | -3.585771 | 0.00033608 | 0.0050011 |
| Onecut2 | 924.234199 | 834.807717 | 1.10712225 | 0.14681454 | 879.520958 | 0.1416397 | 0.03952704 | 3.58336183 | 0.0003392 | 0.00504241 |
| Fen1 | 322.010831 | 278.430662 | 1.15652072 | 0.20979112 | 300.220746 | 0.2131435 | 0.05949578 | 3.58249778 | 0.00034032 | 0.00505406 |
| Insig2 | 538.546712 | 596.891435 | 0.90225237 | -0.1483971 | 567.719073 | -0.1491558 | 0.04164744 | -3.5813918 | 0.00034177 | 0.00507042 |
| Map3k20 | 402.663634 | 357.765965 | 1.12549452 | 0.17055903 | 380.2148 | 0.17125097 | 0.04783145 | 3.5803005 | 0.0003432 | 0.00508656 |
| Bmp7 | 32.3215294 | 20.1907942 | 1.60080525 | 0.67879781 | 26.2561617 | 0.68974654 | 0.19283901 | 3.57679977 | 0.00034783 | 0.00514998 |
| Sec61b | 498.485688 | 448.428887 | 1.11162707 | 0.15267287 | 473.457287 | 0.15363943 | 0.04299593 | 3.57334793 | 0.00035245 | 0.00521318 |
| Mef2a | 1350.42437 | 1441.66803 | 0.93670966 | -0.0943261 | 1396.0462 | -0.0938402 | 0.0262673 | -3.5725117 | 0.00035357 | 0.00522464 |
| Psma4 | 918.230165 | 849.788845 | 1.08053921 | 0.11175142 | 884.009505 | 0.11127157 | 0.03117665 | 3.56906774 | 0.00035825 | 0.00528853 |

| | | | | | | | | | | |
|---|---|---|---|---|---|---|---|---|---|---|
| P4ha2 | 184.00834 | 215.767583 | 0.85280809 | -0.229707 | 199.887962 | -0.2271381 | 0.06367999 | -3.5668672 | 0.00036127 | 0.00532781 |
| Bcar1 | 365.343233 | 410.324983 | 0.89037531 | -0.1675145 | 387.834108 | -0.1676682 | 0.04702215 | -3.5657268 | 0.00036285 | 0.00534571 |
| Rasef | 334.442958 | 295.68424 | 1.13108145 | 0.17770282 | 315.063599 | 0.17788466 | 0.04991621 | 3.56366519 | 0.00036571 | 0.00538255 |
| Shc2 | 11.2506603 | 4.51494041 | 2.49187349 | 1.31723082 | 7.88280026 | 1.3176301 | 0.36990431 | 3.56208368 | 0.00036792 | 0.00540435 |
| Pkd2 | 832.874162 | 900.737328 | 0.92465821 | -0.1130079 | 866.805745 | -0.1128113 | 0.03166977 | -3.5621118 | 0.00036788 | 0.00540435 |
| Slc35f3 | 316.430198 | 357.15938 | 0.88596357 | -0.1746807 | 336.794788 | -0.1753437 | 0.04923324 | -3.561491 | 0.00036875 | 0.00540585 |
| Tgfb1 | 94.6312223 | 117.039012 | 0.80854427 | -0.3066013 | 105.835117 | -0.3074755 | 0.08632844 | -3.5616942 | 0.00036847 | 0.00540585 |
| Ntsr2 | 14.1977734 | 6.5333808 | 2.1731128 | 1.11976306 | 10.365577 | 1.07361499 | 0.30151764 | 3.56070379 | 0.00036986 | 0.00541673 |
| Tecr | 1156.43272 | 1241.8884 | 0.93118892 | -0.1028542 | 1199.16056 | -0.1023566 | 0.02875523 | -3.5595822 | 0.00037145 | 0.00543454 |
| B3gntl1 | 62.3309724 | 46.0723356 | 1.3528937 | 0.43604848 | 54.2016539 | 0.43653538 | 0.12268132 | 3.55828737 | 0.00037328 | 0.00545601 |
| Syt1 | 1576.79756 | 1695.76037 | 0.92984692 | -0.1049349 | 1636.27896 | -0.1016431 | 0.02857055 | -3.5576163 | 0.00037424 | 0.00546457 |
| Scap | 1095.94249 | 1179.04229 | 0.92951924 | -0.1054434 | 1137.49239 | -0.1053898 | 0.02963061 | -3.5567872 | 0.00037542 | 0.00547645 |
| Ahi1 | 666.141541 | 727.173375 | 0.91606976 | -0.1264706 | 696.657458 | -0.1259396 | 0.03543949 | -3.5536513 | 0.00037992 | 0.00553671 |
| Ptprs | 1594.21803 | 1494.46922 | 1.06674531 | 0.09321577 | 1544.34363 | 0.09247429 | 0.02603512 | 3.55190646 | 0.00038245 | 0.00556808 |
| Tes | 167.663629 | 140.724612 | 1.19143074 | 0.25269509 | 154.19412 | 0.25817786 | 0.07270553 | 3.55100731 | 0.00038376 | 0.00558165 |
| Vcan | 111.850539 | 89.8280352 | 1.24516293 | 0.31633453 | 100.839287 | 0.34066231 | 0.09596598 | 3.54982363 | 0.00038549 | 0.00560131 |
| Map1a | 387.88314 | 438.094264 | 0.8853874 | -0.1756193 | 412.988702 | -0.1824211 | 0.05142234 | -3.547506 | 0.0003889 | 0.00564529 |
| Ybx1 | 8442.4507 | 8138.02029 | 1.03740841 | 0.05298397 | 8290.23549 | 0.05279574 | 0.01488829 | 3.54612603 | 0.00039094 | 0.00566938 |
| Ccdc88c | 251.483517 | 214.102986 | 1.17459136 | 0.23215893 | 232.793251 | 0.24460418 | 0.06898721 | 3.54564509 | 0.00039165 | 0.0056706 |
| Brox | 1319.54328 | 1408.75122 | 0.93667587 | -0.0943782 | 1364.14725 | -0.0945357 | 0.02666317 | -3.5455544 | 0.00039179 | 0.0056706 |
| Timp2 | 1250.84701 | 1375.69846 | 0.90924504 | -0.1372589 | 1313.27273 | -0.1308653 | 0.03692245 | -3.5443296 | 0.00039361 | 0.0056859 |
| Csgalnact1 | 166.604901 | 196.601075 | 0.8474262 | -0.2388404 | 181.602988 | -0.2355571 | 0.0664566 | -3.5445247 | 0.00039332 | 0.0056859 |
| Rbm33 | 889.855599 | 818.091667 | 1.08772114 | 0.12130874 | 853.973633 | 0.12122662 | 0.03420925 | 3.54367962 | 0.00039458 | 0.00569439 |
| Sox12 | 860.644888 | 932.121629 | 0.92331822 | -0.1151001 | 896.383258 | -0.1150385 | 0.03246811 | -3.5431244 | 0.00039542 | 0.00570084 |
| P3h3 | 24.3140705 | 14.8344101 | 1.63903184 | 0.71284388 | 19.5742402 | 0.71353236 | 0.20142609 | 3.54240277 | 0.0003965 | 0.0057109 |
| Calm3 | 2767.17402 | 2913.84445 | 0.94966429 | -0.0745105 | 2840.50923 | -0.0743946 | 0.02100713 | -3.541399 | 0.00039801 | 0.00572711 |
| Vps8 | 276.390862 | 314.394456 | 0.8791213 | -0.1858659 | 295.392659 | -0.1888365 | 0.05335394 | -3.5393172 | 0.00040116 | 0.00576128 |
| Plod2 | 87.3170781 | 109.950271 | 0.79415064 | -0.3325154 | 98.6336744 | -0.3328666 | 0.09404259 | -3.5395308 | 0.00040084 | 0.00576128 |
| Ccdc86 | 71.8949678 | 54.958433 | 1.3081699 | 0.38754992 | 63.4267003 | 0.38491888 | 0.10877111 | 3.53879688 | 0.00040196 | 0.00576148 |
| Cpsf7 | 540.664564 | 489.589293 | 1.10432269 | 0.14316179 | 515.126928 | 0.14414375 | 0.04073124 | 3.53889905 | 0.0004018 | 0.00576148 |
| Ccnb1 | 2185.69422 | 2059.88535 | 1.06107567 | 0.08552754 | 2122.78979 | 0.0882488 | 0.02495421 | 3.53642894 | 0.00040558 | 0.00580776 |
| Sel1l3 | 1811.42539 | 1913.50491 | 0.94665312 | -0.0790922 | 1862.46515 | -0.0790089 | 0.02235464 | -3.5343395 | 0.0004088 | 0.00584822 |
| Anpep | 35.9994322 | 49.8225067 | 0.72255361 | -0.4688235 | 42.9109693 | -0.4793391 | 0.13575599 | -3.530887 | 0.00041417 | 0.00591938 |
| Ssbp2 | 571.941477 | 633.155275 | 0.90331945 | -0.1466918 | 602.548376 | -0.1462714 | 0.04142926 | -3.5306304 | 0.00041457 | 0.00591941 |
| Igf2os | 532.306398 | 588.717606 | 0.90417951 | -0.1453189 | 560.512001 | -0.1476262 | 0.04184656 | -3.5277978 | 0.00041903 | 0.00597736 |
| Sash1 | 416.820731 | 462.666723 | 0.90090925 | -0.1505463 | 439.743727 | -0.1522492 | 0.04320329 | -3.5240186 | 0.00042505 | 0.00605744 |
| Slc6a6 | 1988.77508 | 2114.50829 | 0.94053785 | -0.0884421 | 2051.64169 | -0.089233 | 0.02533356 | -3.5223226 | 0.00042778 | 0.00609047 |
| Hectd1 | 2878.42335 | 3010.87805 | 0.95600795 | -0.0649055 | 2944.6507 | -0.0659051 | 0.01871798 | -3.5209502 | 0.00043 | 0.0061162 |
| Ank | 2398.49771 | 2527.52348 | 0.9489517 | -0.0755934 | 2463.01059 | -0.0752315 | 0.02138152 | -3.5185299 | 0.00043394 | 0.00616634 |
| Naip5 | 7.19548078 | 14.1268077 | 0.50934938 | -0.9732725 | 10.6611441 | -0.9845478 | 0.27995309 | -3.5168313 | 0.00043673 | 0.00619999 |
| Mybbp1a | 1247.78687 | 1136.33789 | 1.09807733 | 0.13497965 | 1192.06238 | 0.13045678 | 0.03711006 | 3.51540185 | 0.00043909 | 0.00622155 |
| Osbpl5 | 141.694294 | 169.158015 | 0.83764458 | -0.2555899 | 155.426154 | -0.2555063 | 0.07268092 | -3.5154514 | 0.00043901 | 0.00622155 |
| Prpf38a | 386.976534 | 346.061861 | 1.11822936 | 0.16121613 | 366.519198 | 0.16230764 | 0.04619438 | 3.51357975 | 0.00044211 | 0.00625839 |
| Man2a2 | 591.345535 | 534.967947 | 1.10538498 | 0.14454892 | 563.156741 | 0.14638475 | 0.0416942 | 3.51091377 | 0.00044657 | 0.00631546 |
| Prdx3 | 202.414476 | 172.567056 | 1.17296129 | 0.2301554 | 187.490766 | 0.23034194 | 0.06562845 | 3.50978792 | 0.00044846 | 0.00633018 |
| Dnah8 | 249.006069 | 286.281819 | 0.86979351 | -0.2012551 | 267.643944 | -0.2081343 | 0.05929804 | -3.5099702 | 0.00044816 | 0.00633018 |
| Amer3 | 509.542437 | 562.212442 | 0.90631654 | -0.1419131 | 535.877439 | -0.1415862 | 0.04035497 | -3.5085186 | 0.00045061 | 0.00635441 |
| Lsm11 | 111.189665 | 88.8870257 | 1.25090995 | 0.32297794 | 100.038345 | 0.32321524 | 0.09220624 | 3.50535113 | 0.00045601 | 0.00642028 |
| Tln1 | 1268.50959 | 1374.75506 | 0.9227168 | -0.1160402 | 1321.63232 | -0.1228165 | 0.03503769 | -3.5052684 | 0.00045615 | 0.00642028 |
| Dnajc3 | 921.465975 | 853.448986 | 1.07969661 | 0.11062597 | 887.457481 | 0.11216614 | 0.03200415 | 3.50473709 | 0.00045706 | 0.006427 |
| Brcc3 | 498.671901 | 450.559642 | 1.10678333 | 0.14637282 | 474.615772 | 0.14659436 | 0.04184107 | 3.5036 | 0.00045901 | 0.00644839 |
| Stxbp1 | 2149.35096 | 2280.31389 | 0.94256802 | -0.0853314 | 2214.83243 | -0.0832963 | 0.02377681 | -3.5032568 | 0.00045961 | 0.00645059 |
| Sh3rf2 | 4.91992396 | 11.1110371 | 0.4427961 | -1.1752856 | 8.01548045 | -1.1579751 | 0.33061858 | -3.5024502 | 0.000461 | 0.00646403 |
| Msi2 | 1601.47677 | 1504.43923 | 1.06450081 | 0.09017704 | 1552.958 | 0.0890643 | 0.02543705 | 3.50136179 | 0.00046289 | 0.00648436 |
| G0s2 | 97.0637458 | 119.795619 | 0.81024454 | -0.3035707 | 108.429682 | -0.3039187 | 0.08685991 | -3.4989522 | 0.00046709 | 0.00653706 |
| Mdm2 | 697.360163 | 640.575439 | 1.08864643 | 0.12253547 | 668.967801 | 0.12270012 | 0.03509416 | 3.49631185 | 0.00047174 | 0.00659586 |
| Gpr149 | 1.78947464 | 0.0000001 | 17894746.4 | 24.0930328 | 0.89473727 | 3.4180872 | 0.97818999 | 3.49429787 | 0.00047531 | 0.00663956 |
| Ccar2 | 763.483189 | 830.402817 | 0.91941305 | -0.1212149 | 796.943003 | -0.120961 | 0.03464415 | -3.4915277 | 0.00048027 | 0.00669835 |
| Naip2 | 61.5336488 | 79.4811558 | 0.77419167 | -0.3692373 | 70.5074022 | -0.3667432 | 0.10504819 | -3.4911895 | 0.00048088 | 0.00669835 |
| Oplah | 69.363757 | 90.8550298 | 0.76345533 | -0.3893843 | 80.1093933 | -0.3673122 | 0.105205 | -3.4913951 | 0.00048051 | 0.00669835 |
| Zdhhc9 | 475.237711 | 425.73248 | 1.11628249 | 0.15870216 | 450.485095 | 0.15980736 | 0.04578809 | 3.49015152 | 0.00048275 | 0.00671811 |
| Agap1 | 1983.49462 | 2115.54162 | 0.93758241 | -0.0929826 | 2049.51812 | -0.0924179 | 0.02648376 | -3.4896074 | 0.00048373 | 0.00672549 |
| Pik3cd | 84.6430679 | 107.467553 | 0.7876151 | -0.3444373 | 96.0553105 | -0.3491101 | 0.1000626 | -3.4889171 | 0.00048498 | 0.00673656 |
| Ephb6 | 122.39424 | 147.73123 | 0.82849266 | -0.2714392 | 135.062735 | -0.2634954 | 0.07554739 | -3.4878154 | 0.00048698 | 0.00675804 |
| Ckap2l | 1058.41149 | 977.647414 | 1.08261064 | 0.11451447 | 1018.02945 | 0.11643141 | 0.03339573 | 3.48641633 | 0.00048954 | 0.00678714 |
| Ak2 | 608.788306 | 553.531568 | 1.09982581 | 0.13727505 | 581.159937 | 0.1385111 | 0.0397859 | 3.48141136 | 0.00049878 | 0.00690879 |
| Sqle | 1941.54885 | 2063.40608 | 0.94094365 | -0.0878198 | 2002.47746 | -0.0877435 | 0.02523652 | -3.4768478 | 0.00050735 | 0.00702089 |

| | | | | | | | | | | |
|---|---|---|---|---|---|---|---|---|---|---|
| Pde1c | 855.098025 | 790.922753 | 1.08113975 | 0.11255302 | 823.010389 | 0.11310516 | 0.03254119 | 3.47575367 | 0.00050942 | 0.00704302 |
| Mgst1 | 117.550077 | 142.025692 | 0.82766769 | -0.2728765 | 129.787885 | -0.2756526 | 0.07934813 | -3.473964 | 0.00051283 | 0.00708355 |
| Nr1d2 | 1389.27647 | 1301.77733 | 1.06721514 | 0.09385103 | 1345.5269 | 0.09385804 | 0.02702134 | 3.47347906 | 0.00051376 | 0.00708976 |
| 1110008L16I | 63.6453714 | 47.6855927 | 1.33468765 | 0.41650215 | 55.6654819 | 0.41608604 | 0.11983227 | 3.472237 | 0.00051614 | 0.00711602 |
| Cnksr2 | 2.97646768 | 0.35341233 | 8.42208204 | 3.07417693 | 1.66493991 | 2.97283635 | 0.85635873 | 3.47148483 | 0.00051759 | 0.00712935 |
| Pten | 811.648323 | 747.566553 | 1.08572049 | 0.11865274 | 779.607438 | 0.11793989 | 0.03398632 | 3.47021677 | 0.00052004 | 0.00714981 |
| Ptgfrn | 368.754223 | 414.395611 | 0.88986035 | -0.1683492 | 391.574917 | -0.1654554 | 0.04767745 | -3.4703074 | 0.00051986 | 0.00714981 |
| Pex19 | 889.578036 | 957.883579 | 0.92869118 | -0.1067292 | 923.730807 | -0.1068484 | 0.03080583 | -3.4684466 | 0.00052348 | 0.0071904 |
| Dnmt3a | 1835.21017 | 1699.56466 | 1.07981192 | 0.11078004 | 1767.38742 | 0.10557004 | 0.03046514 | 3.46527398 | 0.00052969 | 0.00726904 |
| Aff2 | 225.578954 | 259.362729 | 0.86974314 | -0.2013387 | 242.470842 | -0.2016744 | 0.05821773 | -3.4641413 | 0.00053193 | 0.00729296 |
| Abhd17b | 209.76312 | 178.36333 | 1.17604398 | 0.23394201 | 194.063225 | 0.23556591 | 0.06801046 | 3.46367182 | 0.00053286 | 0.00729895 |
| Nexmif | 639.49412 | 695.915324 | 0.91892519 | -0.1219807 | 667.704722 | -0.1228718 | 0.03549089 | -3.4620667 | 0.00053604 | 0.00733584 |
| Ctdsp1 | 1243.3488 | 1341.37397 | 0.92692182 | -0.1094804 | 1292.36139 | -0.1096737 | 0.0316843 | -3.4614543 | 0.00053727 | 0.007339 |
| Tmem134 | 884.956882 | 960.004235 | 0.92182602 | -0.1174336 | 922.480559 | -0.1169968 | 0.03379878 | -3.4615673 | 0.00053704 | 0.007339 |
| Ppp2r5a | 1980.11621 | 2089.39689 | 0.9476975 | -0.0775015 | 2034.75655 | -0.0770275 | 0.02225893 | -3.4605198 | 0.00053913 | 0.00735773 |
| Kansl1l | 335.715806 | 376.578879 | 0.89148867 | -0.1657116 | 356.147342 | -0.1639464 | 0.04738354 | -3.4599871 | 0.0005402 | 0.00736552 |
| Vopp1 | 889.551207 | 966.013581 | 0.92084752 | -0.1189658 | 927.782394 | -0.1187215 | 0.03432754 | -3.4584904 | 0.00054321 | 0.00739976 |
| Fam45a | 118.174563 | 95.6451079 | 1.23555262 | 0.30515645 | 106.909835 | 0.30573093 | 0.08843438 | 3.45715 | 0.00054592 | 0.00742925 |
| Ncapg | 847.85095 | 778.913487 | 1.08850465 | 0.12234757 | 813.382218 | 0.12364663 | 0.03576782 | 3.45692383 | 0.00054638 | 0.00742925 |
| Arf5 | 808.686812 | 875.192244 | 0.92401049 | -0.1140189 | 841.939528 | -0.1157763 | 0.03350578 | -3.4554116 | 0.00054945 | 0.0074642 |
| 2900052L18I | 72.20738 | 55.9033658 | 1.29164638 | 0.36921115 | 64.0553728 | 0.39874528 | 0.11541837 | 3.45478168 | 0.00055074 | 0.00747481 |
| Kcnk3 | 404.4877 | 450.849627 | 0.89716765 | -0.1565505 | 427.668663 | -0.1547777 | 0.04480774 | -3.4542618 | 0.0005518 | 0.00748045 |
| Tmem121 | 140.12515 | 168.129617 | 0.83343526 | -0.262858 | 154.127383 | -0.2626773 | 0.07604831 | -3.454085 | 0.00055216 | 0.00748045 |
| Fam20c | 442.034698 | 491.713219 | 0.89896851 | -0.1536575 | 466.873958 | -0.1555629 | 0.0450419 | -3.4537372 | 0.00055288 | 0.00748326 |
| Banf1 | 495.088878 | 440.899337 | 1.12290683 | 0.16723823 | 467.994108 | 0.17090221 | 0.04952937 | 3.45052251 | 0.0005595 | 0.00755915 |
| Rpl31 | 1741.37282 | 1635.22434 | 1.06491371 | 0.09073654 | 1688.29858 | 0.08971178 | 0.02599882 | 3.45060974 | 0.00055932 | 0.00755915 |
| Prickle2 | 401.92906 | 452.679624 | 0.88788856 | -0.1715495 | 427.304342 | -0.1642243 | 0.04763343 | -3.4476689 | 0.00056545 | 0.0076325 |
| Wdtc1 | 1217.62366 | 1300.92518 | 0.93596748 | -0.0954697 | 1259.27442 | -0.0952726 | 0.02763866 | -3.4470786 | 0.00056668 | 0.00764224 |
| Ppl | 603.709939 | 660.867271 | 0.91351163 | -0.130505 | 632.288605 | -0.1318399 | 0.03825495 | -3.4463485 | 0.00056822 | 0.00765595 |
| Arrdc3 | 855.753406 | 946.531421 | 0.90409403 | -0.1454553 | 901.142413 | -0.1501119 | 0.0435628 | -3.4458747 | 0.00056921 | 0.00766242 |
| Pmm2 | 335.718377 | 297.877997 | 1.12703315 | 0.17252995 | 316.798187 | 0.17363962 | 0.05040125 | 3.44514509 | 0.00057075 | 0.00767616 |
| Bend6 | 2.92550591 | 0.22670627 | 12.9043889 | 3.68978992 | 1.57610599 | 3.08561426 | 0.89587667 | 3.44424001 | 0.00057267 | 0.00769492 |
| Psrc1 | 189.138757 | 158.156383 | 1.19589708 | 0.25809324 | 173.64757 | 0.26089477 | 0.07578462 | 3.44258222 | 0.00057619 | 0.00772823 |
| Ttll6 | 6.65518269 | 13.2596607 | 0.50191199 | -0.9944937 | 9.9574216 | -1.0047648 | 0.29186232 | -3.4425987 | 0.00057615 | 0.00772823 |
| Dnpep | 1025.67491 | 1096.72598 | 0.93521529 | -0.0966296 | 1061.20044 | -0.0966235 | 0.02807455 | -3.4416772 | 0.00057812 | 0.00774712 |
| Peli2 | 3.31795796 | 0.34480499 | 9.62270861 | 3.26644304 | 1.83138137 | 2.86320822 | 0.83204137 | 3.44118492 | 0.00057917 | 0.00775422 |
| Eif1ad | 328.607495 | 286.064419 | 1.14871852 | 0.20002532 | 307.335957 | 0.20033093 | 0.0582547 | 3.43888009 | 0.00058413 | 0.00781347 |
| Bcl2 | 303.876548 | 266.786511 | 1.13902516 | 0.18779962 | 285.331529 | 0.18758364 | 0.05457861 | 3.43694403 | 0.00058832 | 0.00786244 |
| Lin28a | 356.759091 | 313.485924 | 1.13803863 | 0.18654953 | 335.122507 | 0.19480702 | 0.05669993 | 3.43575407 | 0.00059091 | 0.00787313 |
| 1700025G04 | 765.418387 | 826.242174 | 0.92638504 | -0.1103161 | 795.830281 | -0.1100494 | 0.03202975 | -3.4358493 | 0.0005907 | 0.00787313 |
| Prepl | 1114.12826 | 1206.0667 | 0.92377002 | -0.1143944 | 1160.09748 | -0.1110884 | 0.03233448 | -3.4356006 | 0.00059124 | 0.00787313 |
| Cd59a | 94.9800205 | 122.703302 | 0.77406247 | -0.3694781 | 108.841661 | -0.335052 | 0.09751805 | -3.4357949 | 0.00059082 | 0.00787313 |
| Pou4f1 | 310.999219 | 273.628954 | 1.13657278 | 0.18469006 | 292.314086 | 0.18449694 | 0.05370997 | 3.4350594 | 0.00059242 | 0.00788179 |
| Mta2 | 632.793323 | 579.489094 | 1.09198487 | 0.12695287 | 606.141208 | 0.12766721 | 0.03718137 | 3.43363416 | 0.00059555 | 0.00791624 |
| Cct5 | 2123.00011 | 2013.54539 | 1.0543592 | 0.07636645 | 2068.27275 | 0.07679234 | 0.02237234 | 3.43246733 | 0.00059812 | 0.00794326 |
| Fgf12 | 701.149582 | 640.103023 | 1.0953699 | 0.13141814 | 670.626302 | 0.13415245 | 0.03909369 | 3.43156299 | 0.00060011 | 0.00796266 |
| Map2k4 | 668.783543 | 614.684586 | 1.08801092 | 0.12169304 | 641.734065 | 0.12167968 | 0.0354709 | 3.43040885 | 0.00060267 | 0.00798945 |
| Spag5 | 1002.04165 | 884.058286 | 1.13345654 | 0.18072908 | 943.049966 | 0.19465926 | 0.05678973 | 3.42771912 | 0.00060867 | 0.00805462 |
| Nrtn | 72.0535403 | 91.4525763 | 0.78787874 | -0.3439545 | 81.7530582 | -0.3387887 | 0.09883237 | -3.4279128 | 0.00060824 | 0.00805462 |
| Tap2 | 259.770711 | 296.542294 | 0.87599886 | -0.1909991 | 278.156502 | -0.1909622 | 0.05575695 | -3.4249036 | 0.00061502 | 0.00813129 |
| Inpp4a | 810.579467 | 902.184556 | 0.89846303 | -0.154469 | 856.382011 | -0.1471227 | 0.04301711 | -3.4200971 | 0.00062599 | 0.00826895 |
| Chst1 | 104.63498 | 127.652271 | 0.81968757 | -0.286854 | 116.143625 | -0.2959654 | 0.08666286 | -3.4151356 | 0.0006375 | 0.00841356 |
| Krr1 | 445.143444 | 401.125641 | 1.1097357 | 0.15021612 | 423.134542 | 0.15062149 | 0.0441155 | 3.4142535 | 0.00063957 | 0.00843334 |
| Aurka | 1253.85079 | 1173.29457 | 1.06865814 | 0.09580041 | 1213.57268 | 0.09783311 | 0.02865977 | 3.4136037 | 0.0006411 | 0.00844596 |
| Mark2 | 574.527307 | 522.542621 | 1.09948411 | 0.13682675 | 548.534964 | 0.13619366 | 0.0399274 | 3.41103226 | 0.00064717 | 0.00851845 |
| Ppfia2 | 209.753439 | 240.654355 | 0.87159627 | -0.1982681 | 225.203897 | -0.1985412 | 0.05823377 | -3.4093821 | 0.0006511 | 0.00856254 |
| Prom2 | 495.967493 | 545.747643 | 0.90878541 | -0.1379884 | 520.857568 | -0.1375673 | 0.04035787 | -3.4086849 | 0.00065277 | 0.00857224 |
| Plppr3 | 115.698769 | 139.310817 | 0.83050815 | -0.2679338 | 127.504793 | -0.2677462 | 0.07855044 | -3.4085897 | 0.000653 | 0.00857224 |
| Atp2b2 | 816.25788 | 880.590758 | 0.9269435 | -0.1094467 | 848.424319 | -0.109193 | 0.0320438 | -3.4076162 | 0.00065533 | 0.00859526 |
| Dennd4a | 968.051099 | 1038.85273 | 0.93184633 | -0.101836 | 1003.45191 | -0.1006577 | 0.02954944 | -3.406417 | 0.00065822 | 0.00862548 |
| Klhl14 | 18.9364819 | 29.0901689 | 0.65095813 | -0.6193634 | 24.0133253 | -0.6197946 | 0.18202491 | -3.4049991 | 0.00066164 | 0.00866273 |
| Nxf1 | 781.638518 | 713.419465 | 1.09562264 | 0.13175098 | 747.528991 | 0.14240556 | 0.04185485 | 3.40236722 | 0.00066805 | 0.00873887 |
| Spats2 | 572.56415 | 625.455668 | 0.91543522 | -0.1274703 | 599.009909 | -0.1273018 | 0.03741838 | -3.4021208 | 0.00066865 | 0.00873904 |
| Pdcd11 | 268.457635 | 234.148387 | 1.1465278 | 0.19727133 | 251.303011 | 0.19445716 | 0.05717519 | 3.40107601 | 0.00067121 | 0.00876478 |
| Dusp3 | 254.36678 | 222.569764 | 1.14286314 | 0.19265265 | 238.468272 | 0.19238895 | 0.05657996 | 3.40030209 | 0.00067311 | 0.00878189 |
| Canx | 4017.90318 | 3820.11023 | 1.05177676 | 0.07282853 | 3919.0067 | 0.07310482 | 0.02153196 | 3.39517787 | 0.00068584 | 0.00894005 |
| Rps29 | 886.429676 | 821.333278 | 1.07925698 | 0.11003843 | 853.881477 | 0.11032798 | 0.03250881 | 3.39378686 | 0.00068933 | 0.00897612 |

| | | | | | | | | | | |
|---|---|---|---|---|---|---|---|---|---|---|
| Bmf | 239.099407 | 272.649609 | 0.87694755 | -0.1894375 | 255.874508 | -0.1862617 | 0.05488625 | -3.3935941 | 0.00068982 | 0.00897612 |
| Pabpc1 | 5234.18581 | 5042.15689 | 1.03808468 | 0.05392413 | 5138.17135 | 0.05392039 | 0.01589562 | 3.39215303 | 0.00069346 | 0.00900765 |
| Myo10 | 522.837211 | 573.725954 | 0.91130131 | -0.134 | 548.281583 | -0.1344801 | 0.03964384 | -3.3922063 | 0.00069332 | 0.00900765 |
| Dcaf8 | 1033.41061 | 1114.78199 | 0.92700691 | -0.109348 | 1074.0963 | -0.1076913 | 0.03175342 | -3.3914859 | 0.00069515 | 0.0090217 |
| Wasf2 | 1633.63605 | 1728.68315 | 0.94501763 | -0.0815869 | 1681.1596 | -0.0817116 | 0.02409857 | -3.3907253 | 0.00069708 | 0.00903886 |
| Psmb4 | 1157.5866 | 1083.78534 | 1.06809582 | 0.09504108 | 1120.68597 | 0.09617591 | 0.02838059 | 3.38879203 | 0.00070201 | 0.00909486 |
| Zfp36 | 70.3781362 | 88.6941064 | 0.79349282 | -0.3337109 | 79.5361212 | -0.3344501 | 0.09872473 | -3.3877035 | 0.0007048 | 0.00912306 |
| Tsr1 | 511.061131 | 461.951326 | 1.10630948 | 0.14575502 | 486.506228 | 0.14595299 | 0.04309037 | 3.38713733 | 0.00070626 | 0.00913393 |
| Adamts9 | 0.13075673 | 2.54152096 | 0.05144822 | -4.280735 | 1.33613875 | -3.0754282 | 0.9085268 | -3.3850715 | 0.0007116 | 0.00919493 |
| Pmf1 | 258.754254 | 226.154469 | 1.14414832 | 0.19427408 | 242.454361 | 0.20445394 | 0.06041394 | 3.384218 | 0.00071381 | 0.00921553 |
| Eif4a3 | 1161.24105 | 1084.72664 | 1.07053797 | 0.09833597 | 1122.98384 | 0.09892886 | 0.02924313 | 3.38297772 | 0.00071704 | 0.00924919 |
| Slc25a11 | 1273.01636 | 1188.8645 | 1.07078339 | 0.09866667 | 1230.94043 | 0.09861175 | 0.02915881 | 3.38188548 | 0.0007199 | 0.00927797 |
| Arhgef12 | 1063.95447 | 1140.32966 | 0.93302359 | -0.1000145 | 1102.14206 | -0.1029564 | 0.03044639 | -3.3815646 | 0.00072074 | 0.00928074 |
| Cdc37l1 | 349.336875 | 310.741999 | 1.12420232 | 0.16890169 | 330.039437 | 0.16800657 | 0.04976067 | 3.37629242 | 0.0007347 | 0.00945222 |
| Luzp1 | 555.023012 | 504.03286 | 1.10116434 | 0.1390298 | 529.527936 | 0.13645775 | 0.04046359 | 3.3723592 | 0.00074527 | 0.00957994 |
| Kdm5d | 203.634899 | 172.121884 | 1.18308546 | 0.24255429 | 187.878391 | 0.24736925 | 0.07338854 | 3.37067948 | 0.00074983 | 0.00962204 |
| Wipf1 | 326.082412 | 365.406118 | 0.89238356 | -0.1642642 | 345.744265 | -0.1665843 | 0.04942166 | -3.3706741 | 0.00074985 | 0.00962204 |
| Eif5a | 3312.29047 | 3171.42308 | 1.04441772 | 0.06269884 | 3241.85678 | 0.06295891 | 0.01868347 | 3.36976573 | 0.00075232 | 0.00964546 |
| Mcf2l | 1344.34017 | 1427.65094 | 0.94164486 | -0.086745 | 1385.99556 | -0.0868678 | 0.02578738 | -3.3686147 | 0.00075547 | 0.00967745 |
| Elavl3 | 2920.41895 | 3059.96361 | 0.95439663 | -0.0673391 | 2990.19128 | -0.0679872 | 0.02019879 | -3.3659047 | 0.00076293 | 0.00976459 |
| Asrgl1 | 82.8472077 | 64.8694371 | 1.27713776 | 0.35291415 | 73.8583223 | 0.34620716 | 0.10287905 | 3.36518628 | 0.00076492 | 0.0097816 |
| Sall2 | 309.05962 | 346.168872 | 0.89280015 | -0.1635908 | 327.614246 | -0.1635205 | 0.04862725 | -3.3627332 | 0.00077175 | 0.00986042 |
| Jun | 2106.34017 | 1975.26244 | 1.06635965 | 0.0926941 | 2040.8013 | 0.09392616 | 0.02794301 | 3.36134684 | 0.00077563 | 0.00990152 |
| Ears2 | 88.367006 | 69.5047947 | 1.27138 | 0.3463953 | 78.9359003 | 0.34503518 | 0.10267907 | 3.36032616 | 0.00077851 | 0.00992962 |
| Btf3l4 | 1048.1644 | 978.575059 | 1.07111293 | 0.0991106 | 1013.36973 | 0.09896403 | 0.02946771 | 3.35838912 | 0.00078398 | 0.00999088 |
| D16Ertd472e | 247.77641 | 215.943211 | 1.14741468 | 0.19838688 | 231.859811 | 0.19825802 | 0.05905361 | 3.35725481 | 0.00078721 | 0.01002334 |
| Arhgdia | 2690.90278 | 2555.75723 | 1.05287887 | 0.07433947 | 2623.33 | 0.07463383 | 0.02223265 | 3.35694769 | 0.00078808 | 0.01002587 |
| Gstm2 | 24.2362625 | 35.2208462 | 0.68812266 | -0.5392623 | 29.7285542 | -0.5390697 | 0.16062774 | -3.3560186 | 0.00079073 | 0.01005098 |
| Zcchc12 | 1413.52446 | 1508.7132 | 0.93690734 | -0.0940217 | 1461.11883 | -0.0989015 | 0.02947963 | -3.3549113 | 0.0007939 | 0.01008265 |
| Suz12 | 1778.89956 | 1668.58253 | 1.06611422 | 0.09236201 | 1723.74105 | 0.09187925 | 0.02739618 | 3.35372444 | 0.00079732 | 0.01010867 |
| Zcchc3 | 369.476823 | 410.027101 | 0.90110342 | -0.1502354 | 389.751962 | -0.1517491 | 0.04524684 | -3.3538056 | 0.00079708 | 0.01010867 |
| Pcdhb14 | 9.29320161 | 16.7702527 | 0.55414798 | -0.8516568 | 13.0317271 | -0.8514722 | 0.25397137 | -3.3526304 | 0.00080048 | 0.01014004 |
| Tkt | 1042.41068 | 1122.88427 | 0.92833314 | -0.1072855 | 1082.64747 | -0.1061982 | 0.03170585 | -3.3494832 | 0.00080962 | 0.01024718 |
| Dnajc2 | 480.397387 | 436.587023 | 1.10034738 | 0.13795906 | 458.492205 | 0.13829894 | 0.04130726 | 3.34805414 | 0.00081381 | 0.01029138 |
| Zc3h10 | 305.331456 | 343.943653 | 0.88773685 | -0.171796 | 324.637554 | -0.1703049 | 0.05088241 | -3.3470293 | 0.00081683 | 0.01032069 |
| Bicc1 | 575.427209 | 628.589529 | 0.91542602 | -0.1274848 | 602.008369 | -0.1256211 | 0.03754343 | -3.3460219 | 0.0008198 | 0.01034944 |
| Pak1 | 2332.07207 | 2444.18366 | 0.95413127 | -0.0677403 | 2388.12786 | -0.0677791 | 0.02027438 | -3.3430905 | 0.00082851 | 0.01044896 |
| Cdkl1 | 14.9086049 | 23.8562434 | 0.62493514 | -0.6782216 | 19.3824241 | -0.6773836 | 0.20263382 | -3.3428951 | 0.00082909 | 0.01044896 |
| Ppargc1a | 170.48979 | 199.740485 | 0.8535565 | -0.2284414 | 185.115138 | -0.2300515 | 0.06887534 | -3.3401141 | 0.00083744 | 0.0105452 |
| Abracl | 299.17913 | 263.31373 | 1.13620786 | 0.18422679 | 281.24643 | 0.18612449 | 0.05573262 | 3.33959707 | 0.000839 | 0.01055588 |
| Tmem94 | 480.848697 | 434.345026 | 1.1070662 | 0.14674149 | 457.596862 | 0.14641037 | 0.0438598 | 3.33814533 | 0.0008434 | 0.01060218 |
| Lpl | 16.5947052 | 9.42379742 | 1.76093611 | 0.81634257 | 13.0092512 | 0.8247224 | 0.24712814 | 3.33722583 | 0.00084619 | 0.01062734 |
| Orc2 | 1142.5914 | 1219.34026 | 0.93705706 | -0.0937912 | 1180.96583 | -0.0937027 | 0.02807979 | -3.3370158 | 0.00084683 | 0.01062734 |
| Npy1r | 1069.69611 | 1155.4348 | 0.9257953 | -0.1112349 | 1112.56545 | -0.1098407 | 0.03292039 | -3.3365558 | 0.00084823 | 0.01063594 |
| Abraxas2 | 317.518251 | 282.25687 | 1.12492656 | 0.16983082 | 299.887561 | 0.16979497 | 0.05089745 | 3.3360212 | 0.00084987 | 0.0106474 |
| Nol4 | 579.705042 | 632.06288 | 0.91716356 | -0.1247491 | 605.88396 | -0.1244155 | 0.03731213 | -3.3344532 | 0.00085467 | 0.01068954 |
| Cx3cl1 | 178.700204 | 238.767609 | 0.74842733 | -0.4180659 | 208.733906 | -0.3744943 | 0.11230547 | -3.3346043 | 0.00085421 | 0.01068954 |
| Zfp641 | 7.14032927 | 2.7862567 | 2.56269613 | 1.35766242 | 4.96329289 | 1.37863127 | 0.41358099 | 3.33340093 | 0.00085791 | 0.01070644 |
| Ankle1 | 265.717281 | 233.034943 | 1.14024651 | 0.18934576 | 249.376112 | 0.22824007 | 0.06846374 | 3.33373634 | 0.00085688 | 0.01070644 |
| Mak16 | 350.03507 | 311.216141 | 1.12473302 | 0.16958259 | 330.625605 | 0.17085491 | 0.05125683 | 3.33331038 | 0.00085819 | 0.01070644 |
| Rpain | 57.8873303 | 43.8961437 | 1.31873384 | 0.39915342 | 50.8917369 | 0.39930295 | 0.11981559 | 3.332646 | 0.00086024 | 0.0107186 |
| Tshz2 | 1168.54858 | 1258.83826 | 0.9282754 | -0.1073752 | 1213.69342 | -0.1051613 | 0.03155604 | -3.3325268 | 0.00086061 | 0.0107186 |
| Ybx2 | 84.3330661 | 65.8077153 | 1.28150728 | 0.35784167 | 75.0703906 | 0.35588385 | 0.10683148 | 3.33126378 | 0.00086453 | 0.01075831 |
| Ndufa4 | 697.073889 | 637.51973 | 1.0934154 | 0.1288416 | 667.296809 | 0.12745545 | 0.03829834 | 3.3279628 | 0.00087484 | 0.01087746 |
| Serp1 | 1448.99533 | 1362.67165 | 1.06334885 | 0.08861497 | 1405.83349 | 0.08969133 | 0.02695301 | 3.32769254 | 0.00087568 | 0.01087889 |
| Bex1 | 167.108615 | 141.494383 | 1.1810265 | 0.24004133 | 154.301499 | 0.23798325 | 0.07155982 | 3.32565486 | 0.00088211 | 0.01093699 |
| Rrs1 | 280.051057 | 246.856895 | 1.13446723 | 0.18201493 | 263.453976 | 0.18205775 | 0.05474584 | 3.32550836 | 0.00088257 | 0.01093699 |
| Ankrd10 | 464.720016 | 420.384338 | 1.10546463 | 0.14465286 | 442.552177 | 0.14629325 | 0.04398774 | 3.32577337 | 0.00088174 | 0.01093699 |
| Fkbp15 | 396.070476 | 438.226293 | 0.90380354 | -0.1459189 | 417.148384 | -0.1450749 | 0.043635 | -3.324737 | 0.00088502 | 0.01095813 |
| Chp1 | 854.95669 | 924.603825 | 0.92467354 | -0.112984 | 889.780258 | -0.1135601 | 0.03417854 | -3.3225567 | 0.00089197 | 0.01103491 |
| Commd7 | 346.134678 | 385.448403 | 0.89800522 | -0.1552043 | 365.79154 | -0.1556126 | 0.04685088 | -3.3214438 | 0.00089553 | 0.01106977 |
| Amt | 1.7582242 | 0.0000001 | 17582242 | 24.0676157 | 0.87911205 | 3.33740837 | 1.00525335 | 3.3199674 | 0.00090028 | 0.01111912 |
| Smpd3 | 1936.78356 | 2047.90753 | 0.9457378 | -0.0804878 | 1992.34554 | -0.0810083 | 0.02440203 | -3.3197371 | 0.00090102 | 0.01111912 |
| E2f3 | 334.973874 | 293.824392 | 1.14004788 | 0.18909441 | 314.399133 | 0.18978621 | 0.05722771 | 3.31633393 | 0.00091207 | 0.01124605 |
| Lrrc3b | 1.99842352 | 0.0000001 | 19984235.2 | 24.252359 | 0.99921171 | 3.2861362 | 0.99111112 | 3.31560822 | 0.00091444 | 0.01126592 |
| Ergic2 | 590.897337 | 540.710301 | 1.09281687 | 0.12805165 | 565.803819 | 0.12708969 | 0.03836317 | 3.3128046 | 0.00092365 | 0.01137001 |
| Larp7 | 498.445275 | 450.340105 | 1.10681964 | 0.14642015 | 474.39269 | 0.14605211 | 0.04409626 | 3.31212033 | 0.00092592 | 0.0113884 |

| | | | | | | | | | | |
|---|---|---|---|---|---|---|---|---|---|---|
| Srp68 | 915.501248 | 851.233886 | 1.07549906 | 0.10500627 | 883.367567 | 0.10501768 | 0.03170961 | 3.31185717 | 0.00092679 | 0.01138966 |
| Fgf2 | 5.63741732 | 1.87063783 | 3.01363376 | 1.5915041 | 3.75402748 | 1.66925607 | 0.50409679 | 3.31138008 | 0.00092837 | 0.01139021 |
| Npr2 | 0.81457817 | 4.12780897 | 0.19733912 | -2.3412511 | 2.47119347 | -2.3429518 | 0.70754143 | -3.3113988 | 0.00092831 | 0.01139021 |
| Cdh10 | 18.6145588 | 10.6691808 | 1.74470367 | 0.80298202 | 14.6418697 | 0.79887221 | 0.24138067 | 3.30959475 | 0.00093431 | 0.01145362 |
| Gabpb2 | 1233.36272 | 1157.25914 | 1.06576193 | 0.0918852 | 1195.31093 | 0.09260627 | 0.02799342 | 3.30814463 | 0.00093916 | 0.01150358 |
| Tbc1d8 | 566.90593 | 621.218839 | 0.91257041 | -0.1319922 | 594.062384 | -0.1324072 | 0.04005426 | -3.3056949 | 0.00094741 | 0.01159504 |
| Gpm6b | 14.6737417 | 7.42450066 | 1.97639443 | 0.98287089 | 11.0491211 | 0.96907882 | 0.29320427 | 3.30513201 | 0.00094932 | 0.01159919 |
| Zdhhc6 | 158.656071 | 134.536398 | 1.17927991 | 0.23790619 | 146.596234 | 0.23800683 | 0.07200785 | 3.30528987 | 0.00094878 | 0.01159919 |
| Lrp4 | 56.2503478 | 72.9740808 | 0.7708264 | -0.3755221 | 64.6122142 | -0.3763964 | 0.11389692 | -3.304711 | 0.00095074 | 0.01160706 |
| Ddx17 | 5219.87912 | 5023.05752 | 1.03918363 | 0.0554506 | 5121.46832 | 0.0565098 | 0.01710126 | 3.30442313 | 0.00095172 | 0.01160943 |
| Rptoros | 165.864905 | 140.597732 | 1.17971252 | 0.23843533 | 153.231318 | 0.23636301 | 0.07154326 | 3.30377768 | 0.00095391 | 0.01162662 |
| Cdk14 | 611.821977 | 668.001961 | 0.91589847 | -0.1267404 | 639.911969 | -0.1257148 | 0.03805465 | -3.3035327 | 0.00095475 | 0.01162722 |
| Naa15 | 1148.9189 | 1075.37635 | 1.06838773 | 0.09543532 | 1112.14762 | 0.0952176 | 0.02883674 | 3.30195439 | 0.00096014 | 0.01167621 |
| Sh3glb1 | 2634.81126 | 2516.05488 | 1.04719944 | 0.06653623 | 2575.43307 | 0.06669975 | 0.02020046 | 3.30189307 | 0.00096035 | 0.01167621 |
| Plcg2 | 42.3620148 | 30.5391827 | 1.38713649 | 0.47210975 | 36.4505987 | 0.46976593 | 0.14236101 | 3.29982164 | 0.00096746 | 0.01172518 |
| Pabpn1 | 1261.76864 | 1174.00092 | 1.0747595 | 0.10401386 | 1217.88478 | 0.10928825 | 0.03311341 | 3.3004224 | 0.00096539 | 0.01172518 |
| Clasp1 | 1662.3062 | 1765.23995 | 0.94168852 | -0.0866781 | 1713.77307 | -0.0867279 | 0.02628278 | -3.2997995 | 0.00096754 | 0.01172518 |
| Skor1 | 11.6160579 | 19.5388001 | 0.59451235 | -0.7502213 | 15.5774289 | -0.7571006 | 0.22943311 | -3.2998753 | 0.00096728 | 0.01172518 |
| Tfam | 593.213416 | 540.84368 | 1.09682971 | 0.13333956 | 567.028548 | 0.13226836 | 0.04010601 | 3.29796828 | 0.00097387 | 0.01179228 |
| H2afv | 901.480868 | 971.776407 | 0.92766285 | -0.1083275 | 936.628637 | -0.1096229 | 0.03326342 | -3.295599 | 0.00098212 | 0.01188246 |
| Cdc7 | 533.581055 | 485.518407 | 1.09899243 | 0.13618145 | 509.549731 | 0.13860853 | 0.04207795 | 3.2940892 | 0.00098741 | 0.01193673 |
| Uchl1 | 4544.23016 | 4728.1269 | 0.9611058 | -0.0572328 | 4636.17853 | -0.0564177 | 0.01713099 | -3.2933135 | 0.00099014 | 0.01195996 |
| Gpx4 | 896.748484 | 963.113265 | 0.93109348 | -0.1030021 | 929.930874 | -0.1030972 | 0.03130889 | -3.2929038 | 0.00099158 | 0.01196764 |
| Vangl2 | 301.670587 | 337.908351 | 0.8927586 | -0.163658 | 319.789469 | -0.1641401 | 0.04985143 | -3.2925855 | 0.00099271 | 0.01197145 |
| Bptf | 829.389817 | 743.761095 | 1.11512934 | 0.15721105 | 786.575456 | 0.1426576 | 0.04334704 | 3.29105763 | 0.00099811 | 0.01200735 |
| Pak7 | 118.545397 | 142.500118 | 0.83189684 | -0.2655235 | 130.522757 | -0.2597237 | 0.07891259 | -3.2912838 | 0.00099731 | 0.01200735 |
| Reck | 52.236588 | 68.4848325 | 0.76274682 | -0.3907238 | 60.3607102 | -0.3963239 | 0.12041074 | -3.2914329 | 0.00099678 | 0.01200735 |
| Mxi1 | 421.039598 | 379.383428 | 1.10979966 | 0.15029926 | 400.211513 | 0.14926198 | 0.04536474 | 3.29026417 | 0.00100093 | 0.0120315 |
| Ppp1r37 | 995.600077 | 1072.97859 | 0.92788438 | -0.107983 | 1034.28933 | -0.1067804 | 0.03246033 | -3.2895671 | 0.00100342 | 0.01205157 |
| Tnpo3 | 966.957824 | 899.196384 | 1.07535778 | 0.10481673 | 933.077104 | 0.10520231 | 0.03199294 | 3.28829766 | 0.00100795 | 0.01209624 |
| Set | 2937.67607 | 2812.43829 | 1.04452997 | 0.06285388 | 2875.05718 | 0.06286062 | 0.01912433 | 3.28694545 | 0.0010128 | 0.01213482 |
| Tab1 | 220.423369 | 251.565589 | 0.87620636 | -0.1906574 | 235.994479 | -0.1897998 | 0.05774094 | -3.2870917 | 0.00101228 | 0.01213482 |
| Romo1 | 446.82606 | 404.295176 | 1.10519761 | 0.14430434 | 425.560618 | 0.14525621 | 0.04421302 | 3.28537158 | 0.00101848 | 0.01219296 |
| Chrm2 | 153.650536 | 182.490578 | 0.84196422 | -0.2481692 | 168.070557 | -0.241704 | 0.07357926 | -3.284947 | 0.00102002 | 0.01220149 |
| Dhx33 | 274.204255 | 240.15815 | 1.14176536 | 0.19126619 | 257.181203 | 0.1912165 | 0.05823276 | 3.2836583 | 0.00102469 | 0.01224753 |
| Timeless | 323.106922 | 286.681561 | 1.12705861 | 0.17256254 | 304.894241 | 0.17470553 | 0.05321715 | 3.28288049 | 0.00102752 | 0.01227148 |
| Ppp1r14b | 247.24794 | 216.237417 | 1.14340961 | 0.19334232 | 231.742679 | 0.19678782 | 0.05995791 | 3.28209939 | 0.00103037 | 0.01229562 |
| Gm14326 | 86.0259205 | 68.4218712 | 1.25728687 | 0.33031386 | 77.2238958 | 0.33025538 | 0.1006802 | 3.28024174 | 0.00103718 | 0.01236691 |
| Kif20a | 1491.5606 | 1401.93681 | 1.06392855 | 0.08940127 | 1446.7487 | 0.08937615 | 0.02725212 | 3.27960405 | 0.00103953 | 0.01238493 |
| Gpr146 | 79.188039 | 98.068849 | 0.80747393 | -0.3085124 | 88.6284439 | -0.3088559 | 0.09422164 | -3.2779725 | 0.00104556 | 0.01244673 |
| Spag7 | 243.482844 | 213.704232 | 1.13934498 | 0.18820465 | 228.593538 | 0.1882812 | 0.05746062 | 3.27669991 | 0.00105028 | 0.01249293 |
| Sobp | 260.82071 | 298.801976 | 0.87288817 | -0.1961313 | 279.811343 | -0.1975258 | 0.06033267 | -3.2739434 | 0.00106058 | 0.01260532 |
| Mettl16 | 423.964013 | 383.975384 | 1.10414373 | 0.14292799 | 403.969698 | 0.14315148 | 0.0437427 | 3.27257987 | 0.00106571 | 0.01265613 |
| Smc2 | 2188.91868 | 2047.8214 | 1.06890117 | 0.09612846 | 2118.37004 | 0.10249234 | 0.03134284 | 3.27004049 | 0.00107532 | 0.01276007 |
| Sncb | 285.592585 | 322.313416 | 0.88607104 | -0.1745057 | 303.953001 | -0.1745466 | 0.05338182 | -3.2697769 | 0.00107632 | 0.01276105 |
| Gnpda2 | 247.931587 | 280.527868 | 0.88380377 | -0.178202 | 264.229728 | -0.1780237 | 0.05444873 | -3.2695662 | 0.00107713 | 0.01276105 |
| Ddx42 | 1141.26184 | 1070.99426 | 1.06560967 | 0.09167908 | 1106.12805 | 0.09180045 | 0.02810175 | 3.26671589 | 0.00108803 | 0.01287992 |
| Ankrd13d | 245.948255 | 280.244915 | 0.87761898 | -0.1883334 | 263.096585 | -0.1869093 | 0.05724526 | -3.2650614 | 0.0010944 | 0.01294505 |
| Ndrg3 | 1022.82478 | 1092.10093 | 0.93656617 | -0.0945472 | 1057.46286 | -0.0941343 | 0.02883545 | -3.2645328 | 0.00109645 | 0.01295889 |
| Fn1 | 46.1182606 | 75.1047862 | 0.61405222 | -0.7035667 | 60.6115233 | -0.6447541 | 0.19758837 | -3.2631179 | 0.00110194 | 0.01301339 |
| Hist1h2bc | 182.053415 | 155.803796 | 1.16847869 | 0.22463142 | 168.928605 | 0.22431243 | 0.06876577 | 3.26197779 | 0.00110638 | 0.01304505 |
| D630045J12I | 220.1208 | 190.695935 | 1.15430253 | 0.20702139 | 205.408367 | 0.2055222 | 0.06300263 | 3.2621208 | 0.00110582 | 0.01304505 |
| Hlf | 606.836449 | 550.334596 | 1.10266818 | 0.14099872 | 578.585523 | 0.13678589 | 0.04197978 | 3.2583754 | 0.00112052 | 0.0132013 |
| Slit1 | 103.750573 | 82.9251494 | 1.25113519 | 0.32323769 | 93.337861 | 0.3324727 | 0.10204855 | 3.25798557 | 0.00112206 | 0.01320477 |
| Edn3 | 48.1608085 | 63.3306884 | 0.76046558 | -0.3950452 | 55.7457484 | -0.392947 | 0.12061542 | -3.2578501 | 0.0011226 | 0.01320477 |
| Scfd2 | 175.169593 | 149.056624 | 1.17518825 | 0.23289187 | 162.113109 | 0.23250915 | 0.07142288 | 3.25538765 | 0.00113238 | 0.01330527 |
| Ide | 1167.86328 | 1079.93236 | 1.08142262 | 0.11293044 | 1123.89782 | 0.10924116 | 0.03355848 | 3.2552474 | 0.00113294 | 0.01330527 |
| Kcnq2 | 1640.89169 | 1732.4986 | 0.9471244 | -0.0783742 | 1686.69514 | -0.0784965 | 0.02413933 | -3.251811 | 0.00114672 | 0.01345651 |
| Camsap2 | 1593.5003 | 1685.62866 | 0.9453448 | -0.0810875 | 1639.56448 | -0.0809692 | 0.02491058 | -3.2503936 | 0.00115245 | 0.01351307 |
| Rnpepl1 | 1924.88938 | 2042.15296 | 0.94257845 | -0.0853154 | 1983.52117 | -0.0848628 | 0.02611495 | -3.2495855 | 0.00115573 | 0.01353011 |
| Rcor3 | 621.501364 | 673.869675 | 0.92228718 | -0.116712 | 647.685519 | -0.1165825 | 0.03587545 | -3.2496469 | 0.00115548 | 0.01353011 |
| Slc16a11 | 34.4469844 | 23.5582019 | 1.46220771 | 0.54814826 | 29.0025931 | 0.54781967 | 0.16866846 | 3.24790816 | 0.00116257 | 0.01359938 |
| Ndufs2 | 1397.42253 | 1480.17164 | 0.94409492 | -0.0829962 | 1438.79709 | -0.082326 | 0.02536726 | -3.2453623 | 0.00117301 | 0.01371074 |
| Rrbp1 | 914.334761 | 845.634506 | 1.08124107 | 0.11268822 | 879.984634 | 0.11209878 | 0.03456154 | 3.24345485 | 0.0011809 | 0.013792 |
| Cdkn2aipnl | 632.457758 | 581.488656 | 1.08765279 | 0.12121808 | 606.973207 | 0.1211126 | 0.03734727 | 3.24287693 | 0.00118329 | 0.01379826 |
| Tmsb10 | 1721.87566 | 1822.11828 | 0.94498567 | -0.0816356 | 1771.99697 | -0.079804 | 0.02460753 | -3.2430733 | 0.00118248 | 0.01379826 |
| Fbxo2 | 145.285623 | 170.464633 | 0.85229188 | -0.2305805 | 157.875128 | -0.2317743 | 0.07148134 | -3.242444 | 0.00118509 | 0.01380838 |

| | | | | | | | | | | |
|---|---|---|---|---|---|---|---|---|---|---|
| Gapdh | 4408.59774 | 4565.50653 | 0.96563168 | -0.0504551 | 4487.05214 | -0.0505745 | 0.01560713 | -3.2404759 | 0.0011933 | 0.01389313 |
| Kcnc2 | 1.88296289 | 0.0000001 | 18829628.9 | 24.1665012 | 0.9414814 | 3.41668768 | 1.05470379 | 3.23947605 | 0.0011975 | 0.01391629 |
| Zmym3 | 548.8061 | 597.066346 | 0.91917105 | -0.1215947 | 572.936223 | -0.1216335 | 0.03755053 | -3.2391968 | 0.00119867 | 0.01391629 |
| 4930539E08 | 134.356618 | 158.682236 | 0.84670232 | -0.2400732 | 146.519427 | -0.2391792 | 0.07384112 | -3.2391063 | 0.00119905 | 0.01391629 |
| Zfyve9 | 45.4277362 | 60.1713637 | 0.75497269 | -0.4055036 | 52.7995499 | -0.4084753 | 0.12609947 | -3.2393104 | 0.00119819 | 0.01391629 |
| Slc3a2 | 451.603026 | 407.571567 | 1.10803369 | 0.14800174 | 429.587297 | 0.15394506 | 0.04753773 | 3.23837653 | 0.00120212 | 0.01394102 |
| Ctif | 607.17121 | 659.80656 | 0.92022609 | -0.1199397 | 633.488885 | -0.1198792 | 0.03703529 | -3.2368922 | 0.00120839 | 0.01400278 |
| Slc2a3 | 4.05201917 | 0.90684897 | 4.46824036 | 2.1597068 | 2.47943397 | 2.12197176 | 0.65583107 | 3.23554624 | 0.0012141 | 0.01405798 |
| Ift57 | 360.812962 | 403.346537 | 0.89454831 | -0.1607687 | 382.079749 | -0.1603312 | 0.04956606 | -3.2346968 | 0.00121772 | 0.01408885 |
| Ly6h | 397.88898 | 441.557813 | 0.9011028 | -0.1502364 | 419.723396 | -0.1506583 | 0.04658041 | -3.2343698 | 0.00121911 | 0.01409399 |
| Ncoa3 | 881.721783 | 945.590904 | 0.93245586 | -0.1008927 | 913.656343 | -0.1015396 | 0.03140227 | -3.2335127 | 0.00122278 | 0.01412534 |
| Ogdhl | 607.844903 | 662.38257 | 0.9176644 | -0.1239615 | 635.113737 | -0.1228254 | 0.03801063 | -3.2313437 | 0.0012321 | 0.01422189 |
| Mis12 | 508.182916 | 462.449968 | 1.09889275 | 0.13605059 | 485.316442 | 0.13639532 | 0.04221549 | 3.23093031 | 0.00123388 | 0.01423139 |
| Ssh2 | 627.785949 | 566.580846 | 1.10802537 | 0.14799091 | 597.183397 | 0.14327664 | 0.04439101 | 3.22760516 | 0.00124831 | 0.01438301 |
| Fbxo31 | 296.798946 | 334.41363 | 0.88752048 | -0.1721477 | 315.606288 | -0.1719725 | 0.05328424 | -3.227455 | 0.00124897 | 0.01438301 |
| Lypd6 | 4.46048344 | 1.13218223 | 3.93972216 | 1.97809389 | 2.79633274 | 1.93928244 | 0.60099089 | 3.22680838 | 0.00125179 | 0.01440436 |
| Kcnip2 | 277.253193 | 239.510484 | 1.1575827 | 0.21111527 | 258.381838 | 0.20638924 | 0.06398616 | 3.22552917 | 0.0012574 | 0.01445766 |
| Prdx6 | 659.472459 | 718.018236 | 0.91846199 | -0.1227081 | 688.745348 | -0.122811 | 0.03809493 | -3.2238158 | 0.00126495 | 0.01453317 |
| Cct6a | 2247.23537 | 2145.26848 | 1.04753107 | 0.06699303 | 2196.25193 | 0.06786393 | 0.02105477 | 3.22320934 | 0.00126763 | 0.01455269 |
| Sall4 | 175.301275 | 203.147982 | 0.86292403 | -0.2126945 | 189.224629 | -0.21302 | 0.06611366 | -3.2220274 | 0.00127287 | 0.01460155 |
| Naa50 | 1638.53661 | 1543.82948 | 1.06134559 | 0.08589449 | 1591.18305 | 0.08745517 | 0.02714565 | 3.22170089 | 0.00127432 | 0.01460689 |
| Cpe | 2811.39447 | 3025.67262 | 0.92917999 | -0.10597 | 2918.53354 | -0.1042642 | 0.03237988 | -3.22003 | 0.00128177 | 0.01468095 |
| Cadm4 | 1235.45783 | 1314.69748 | 0.93972784 | -0.0896851 | 1275.07766 | -0.0909954 | 0.02826259 | -3.2196405 | 0.00128351 | 0.01468224 |
| Mxd1 | 283.763289 | 320.314139 | 0.88589061 | -0.1747995 | 302.038714 | -0.1744413 | 0.05418169 | -3.219562 | 0.00128387 | 0.01468224 |
| Hells | 473.625371 | 429.408155 | 1.10297246 | 0.14139678 | 451.516763 | 0.1415205 | 0.04397672 | 3.21807789 | 0.00129053 | 0.01474704 |
| Hif1an | 403.514808 | 362.259129 | 1.11388444 | 0.15559957 | 382.886968 | 0.15361269 | 0.04775132 | 3.21693078 | 0.0012957 | 0.01479472 |
| Polq | 226.618843 | 196.413361 | 1.15378527 | 0.20637475 | 211.516102 | 0.2102159 | 0.06537071 | 3.21575062 | 0.00130104 | 0.01484206 |
| Slc2a13 | 1939.79516 | 1842.24751 | 1.05295035 | 0.07443741 | 1891.02134 | 0.07360229 | 0.02288933 | 3.21557217 | 0.00130185 | 0.01484206 |
| Rnf180 | 118.426682 | 140.825291 | 0.84094754 | -0.2499123 | 129.625986 | -0.2494314 | 0.0775821 | -3.2150633 | 0.00130416 | 0.01485492 |
| Kif12 | 8.12138072 | 14.6332461 | 0.55499516 | -0.8494529 | 11.3773133 | -0.8472712 | 0.26354657 | -3.2148823 | 0.00130498 | 0.01485492 |
| Egr4 | 7.42260084 | 2.95392148 | 2.51279558 | 1.32929331 | 5.18826106 | 1.33142593 | 0.41424417 | 3.21410907 | 0.0013085 | 0.01488356 |
| Acadvl | 498.414803 | 452.437231 | 1.10162199 | 0.13962927 | 475.426017 | 0.14204299 | 0.04420831 | 3.2130379 | 0.00131339 | 0.01492133 |
| Cct3 | 1903.0319 | 1799.06444 | 1.05778974 | 0.08105288 | 1851.04817 | 0.08155678 | 0.02538384 | 3.21294066 | 0.00131383 | 0.01492133 |
| Cckbr | 3.28324462 | 8.08788971 | 0.40594577 | -1.3006411 | 5.68556707 | -1.2963686 | 0.40351777 | -3.2126679 | 0.00131508 | 0.01492407 |
| Mylk4 | 6.15418084 | 12.3541928 | 0.49814512 | -1.005362 | 9.25418671 | -1.0055746 | 0.31325043 | -3.2101301 | 0.00132675 | 0.01504495 |
| Eef1a2 | 2778.85228 | 2941.11303 | 0.94483015 | -0.0818731 | 2859.98265 | -0.0810692 | 0.02525767 | -3.2096849 | 0.00132881 | 0.01505675 |
| Runx1t1 | 2488.87805 | 2378.06918 | 1.04659615 | 0.06570486 | 2433.47362 | 0.06612618 | 0.02060621 | 3.20904081 | 0.00133179 | 0.01507899 |
| Cerk | 892.378874 | 959.901457 | 0.92965676 | -0.1052299 | 926.140165 | -0.1052967 | 0.03281861 | -3.2084438 | 0.00133455 | 0.0150988 |
| Fez1 | 132.392275 | 157.933507 | 0.83827858 | -0.2544983 | 145.162891 | -0.2554193 | 0.07965623 | -3.2065201 | 0.00134351 | 0.01518852 |
| Btaf1 | 486.536806 | 437.955641 | 1.11092714 | 0.1517642 | 462.246223 | 0.1514527 | 0.04727348 | 3.20375625 | 0.00135647 | 0.01532339 |
| Bik | 114.22971 | 136.251512 | 0.83837389 | -0.2543343 | 125.240611 | -0.2536471 | 0.07921106 | -3.2021672 | 0.00136398 | 0.01539644 |
| Fgl2 | 83.4142847 | 102.282923 | 0.81552504 | -0.2941989 | 92.848604 | -0.2957293 | 0.09236086 | -3.2018897 | 0.00136529 | 0.01539955 |
| Fam208a | 374.852945 | 336.34534 | 1.1144883 | 0.15638147 | 355.599142 | 0.1545491 | 0.04828299 | 3.20090185 | 0.00136998 | 0.01542895 |
| Psma6 | 919.911888 | 858.92275 | 1.07100655 | 0.0989673 | 889.417319 | 0.09879707 | 0.03086356 | 3.20109116 | 0.00136908 | 0.01542895 |
| Thsd4 | 1063.42035 | 995.647835 | 1.06806876 | 0.09500453 | 1029.53409 | 0.09506375 | 0.02970416 | 3.20035179 | 0.0013726 | 0.01543497 |
| Arfgef3 | 585.740703 | 639.221409 | 0.91633461 | -0.1260536 | 612.481055 | -0.1400132 | 0.0437476 | -3.2004769 | 0.001372 | 0.01543497 |
| Cd81 | 730.474718 | 787.639504 | 0.92742265 | -0.1087011 | 759.057111 | -0.1087817 | 0.03400193 | -3.1992803 | 0.00137771 | 0.01548072 |
| Sik1 | 818.118037 | 753.326599 | 1.0860071 | 0.11903354 | 785.722318 | 0.11729252 | 0.03667239 | 3.19838736 | 0.00138199 | 0.01551374 |
| Cblb | 107.673274 | 129.441092 | 0.83183225 | -0.2656355 | 118.557183 | -0.2654728 | 0.0830062 | -3.1982289 | 0.00138274 | 0.01551374 |
| Fbxo9 | 711.094647 | 652.604198 | 1.08962622 | 0.12383333 | 681.849423 | 0.12381009 | 0.03872429 | 3.19722058 | 0.00138759 | 0.01555631 |
| Rnaseh2c | 85.7452916 | 68.8775148 | 1.24489526 | 0.31602436 | 77.3114031 | 0.31870268 | 0.09970685 | 3.19639699 | 0.00139156 | 0.01558899 |
| Pls1 | 470.269301 | 518.020145 | 0.90782049 | -0.139521 | 494.144723 | -0.136909 | 0.04284551 | -3.19541 | 0.00139632 | 0.01563059 |
| Kcnip3 | 418.447761 | 459.281105 | 0.91109292 | -0.1343299 | 438.864433 | -0.1347411 | 0.04217109 | -3.195107 | 0.00139779 | 0.01563521 |
| Col11a1 | 17.9537674 | 10.2290257 | 1.75517863 | 0.81161786 | 14.0913965 | 0.80420053 | 0.25171748 | 3.19485367 | 0.00139902 | 0.01563713 |
| Hspg2 | 7.12871825 | 13.725695 | 0.51937029 | -0.9451646 | 10.4272065 | -0.9371459 | 0.29342927 | -3.1937711 | 0.00140427 | 0.01568406 |
| Uqcrq | 515.343549 | 467.182695 | 1.10308784 | 0.14154768 | 491.263122 | 0.14291317 | 0.04477015 | 3.19215307 | 0.00141216 | 0.01575733 |
| Kcnd2 | 347.101037 | 385.942824 | 0.8993587 | -0.1530315 | 366.52193 | -0.1530735 | 0.0479555 | -3.1919903 | 0.00141296 | 0.01575733 |
| Zbtb42 | 268.261448 | 301.963188 | 0.88839123 | -0.1707329 | 285.112318 | -0.171335 | 0.05368872 | -3.191266 | 0.00141651 | 0.01578501 |
| Pcsk9 | 127.024866 | 105.31586 | 1.20613235 | 0.27038823 | 116.170363 | 0.27738516 | 0.08701207 | 3.18789304 | 0.00143314 | 0.0159583 |
| Sufu | 160.909323 | 137.319727 | 1.17178592 | 0.22870902 | 149.114525 | 0.22860051 | 0.0717303 | 3.18694473 | 0.00143784 | 0.0159987 |
| Dpysl4 | 29.1080107 | 19.5650869 | 1.48775269 | 0.57313473 | 24.3365487 | 0.57524308 | 0.18068389 | 3.18369875 | 0.00145406 | 0.01616704 |
| Kbtbd11 | 684.619558 | 743.270884 | 0.92109024 | -0.1185856 | 713.945221 | -0.1179191 | 0.0371096 | -3.1775904 | 0.00148504 | 0.01649913 |
| Fcmr | 5.54115016 | 1.7975625 | 3.08259109 | 1.62414353 | 3.66935623 | 1.62300145 | 0.51094931 | 3.17644321 | 0.00149093 | 0.0165483 |
| Cxcr4 | 122.856864 | 145.269469 | 0.84571703 | -0.2417531 | 134.063167 | -0.246364 | 0.07756591 | -3.1761889 | 0.00149224 | 0.0165483 |
| Pcdhb22 | 61.4862224 | 79.082351 | 0.77749614 | -0.3630926 | 70.2842866 | -0.3597997 | 0.11328434 | -3.1760759 | 0.00149282 | 0.0165483 |
| Zfp971 | 129.69146 | 108.308313 | 1.19742849 | 0.2599395 | 118.999887 | 0.26019605 | 0.08193496 | 3.17564152 | 0.00149506 | 0.0165607 |
| Enpp3 | 42.1199121 | 55.4562897 | 0.75951551 | -0.3968487 | 48.7881008 | -0.3985887 | 0.12557199 | -3.174185 | 0.00150258 | 0.01663161 |

| | | | | | | | | | | |
|---|---|---|---|---|---|---|---|---|---|---|
| Fkbp8 | 1340.17872 | 1417.53266 | 0.94543057 | -0.0809566 | 1378.85569 | -0.081056 | 0.02553988 | -3.1737023 | 0.00150508 | 0.01664685 |
| Aldh1b1 | 133.684213 | 158.256712 | 0.84473013 | -0.2434376 | 145.970462 | -0.2403155 | 0.07576738 | -3.1717542 | 0.00151521 | 0.01674642 |
| Smg7 | 1716.92586 | 1808.84488 | 0.94918358 | -0.075241 | 1762.88537 | -0.074473 | 0.02348192 | -3.1715053 | 0.00151651 | 0.0167483 |
| Ndufaf4 | 280.192962 | 249.374647 | 1.12358239 | 0.16810592 | 264.783805 | 0.16809984 | 0.05303443 | 3.16963626 | 0.0015263 | 0.01684384 |
| Scnn1g | 49.4953734 | 36.6706967 | 1.34972547 | 0.432666 | 43.083035 | 0.43821157 | 0.13836647 | 3.16703575 | 0.00154001 | 0.01698255 |
| Grina | 732.871676 | 791.124397 | 0.92636718 | -0.110344 | 761.998037 | -0.110109 | 0.03477128 | -3.1666663 | 0.00154197 | 0.0169915 |
| Ngf | 197.158347 | 168.678022 | 1.16884431 | 0.22508278 | 182.918185 | 0.22479819 | 0.07100541 | 3.16593059 | 0.00154588 | 0.01702187 |
| Megf6 | 11.8490882 | 19.5338593 | 0.60659228 | -0.721201 | 15.6914737 | -0.714056 | 0.22558459 | -3.1653581 | 0.00154892 | 0.01704274 |
| Usp25 | 310.345515 | 347.853945 | 0.8921719 | -0.1646064 | 329.09973 | -0.1652097 | 0.05220249 | -3.164786 | 0.00155197 | 0.01706361 |
| Epha5 | 2.00920216 | 0.10853126 | 18.5126591 | 4.21044023 | 1.05886661 | 3.08883734 | 0.97619613 | 3.1641565 | 0.00155533 | 0.01707523 |
| Snx33 | 176.788536 | 205.175799 | 0.8616442 | -0.2148358 | 190.982168 | -0.2145218 | 0.06779499 | -3.1642719 | 0.00155471 | 0.01707523 |
| Psmb3 | 803.454014 | 745.759167 | 1.07736391 | 0.10750565 | 774.60659 | 0.1101788 | 0.03483066 | 3.16327062 | 0.00156007 | 0.01711461 |
| Mrm2 | 58.4116549 | 74.2868049 | 0.7862992 | -0.3468497 | 66.3492298 | -0.346684 | 0.10967336 | -3.1610589 | 0.00157197 | 0.01723234 |
| Slfn3 | 3.16545072 | 7.89160244 | 0.40111634 | -1.3179073 | 5.52852648 | -1.3043094 | 0.41275627 | -3.1599989 | 0.0015777 | 0.01728237 |
| Kcnh6 | 391.786477 | 432.135573 | 0.90662862 | -0.1414164 | 411.961025 | -0.1412752 | 0.04477284 | -3.1553781 | 0.0016029 | 0.01754551 |
| Rrm2 | 1828.59712 | 1705.33827 | 1.07227824 | 0.10067931 | 1766.9677 | 0.11630527 | 0.03690127 | 3.15179611 | 0.0016227 | 0.01774905 |
| Spc24 | 313.307771 | 278.770459 | 1.12389158 | 0.16850287 | 296.039115 | 0.17439847 | 0.05533907 | 3.15145297 | 0.0016246 | 0.0177494 |
| Snx27 | 447.279357 | 405.99916 | 1.10167557 | 0.13969943 | 426.639259 | 0.14054402 | 0.0445979 | 3.15135967 | 0.00162512 | 0.0177494 |
| Itgb8 | 49.364606 | 64.3163918 | 0.7675276 | -0.3817095 | 56.8404988 | -0.3728656 | 0.11833916 | -3.150822 | 0.00162812 | 0.01776901 |
| Nolc1 | 382.703562 | 343.320517 | 1.11471218 | 0.15667126 | 363.012039 | 0.15663138 | 0.04973656 | 3.14922026 | 0.00163707 | 0.01781423 |
| Pik3ca | 645.737365 | 596.552765 | 1.08244803 | 0.11429776 | 621.145065 | 0.1144987 | 0.03635501 | 3.14946129 | 0.00163572 | 0.01781423 |
| Lpin1 | 238.517578 | 271.732906 | 0.87776479 | -0.1880937 | 255.125242 | -0.1819001 | 0.05775971 | -3.1492563 | 0.00163687 | 0.01781423 |
| Pld1 | 248.141755 | 283.807095 | 0.87433246 | -0.1937461 | 265.974425 | -0.1905347 | 0.06049286 | -3.1497056 | 0.00163435 | 0.01781423 |
| Rnft1 | 159.142379 | 134.993991 | 1.17888491 | 0.23742287 | 147.068185 | 0.23749269 | 0.07551735 | 3.14487567 | 0.00166157 | 0.01796221 |
| Gm14325 | 181.794551 | 156.009137 | 1.16528143 | 0.22067843 | 168.901844 | 0.22162611 | 0.07043526 | 3.14652239 | 0.00165225 | 0.01796221 |
| Men1 | 337.615808 | 299.389446 | 1.12768106 | 0.17335909 | 318.502627 | 0.17165158 | 0.05457629 | 3.14516732 | 0.00165992 | 0.01796221 |
| Ggt7 | 784.300695 | 843.504813 | 0.92981176 | -0.1049894 | 813.902754 | -0.1043877 | 0.03317904 | -3.1461954 | 0.00165409 | 0.01796221 |
| Gfpt1 | 693.42918 | 749.985556 | 0.92459005 | -0.1131143 | 721.707368 | -0.1144839 | 0.03639796 | -3.145339 | 0.00165894 | 0.01796221 |
| Arfgap3 | 451.535405 | 494.065118 | 0.91391881 | -0.1298621 | 472.800261 | -0.1302133 | 0.04140411 | -3.1449365 | 0.00166123 | 0.01796221 |
| Chchd10 | 304.236945 | 341.507034 | 0.89086582 | -0.1667199 | 322.871989 | -0.1699003 | 0.05401752 | -3.1452813 | 0.00165927 | 0.01796221 |
| Emp2 | 723.218577 | 837.391352 | 0.86365661 | -0.2114703 | 780.304964 | -0.1936364 | 0.06154801 | -3.1461034 | 0.00165461 | 0.01796221 |
| Car11 | 39.7159069 | 53.0187449 | 0.7490918 | -0.4167856 | 46.3673258 | -0.4152145 | 0.13201764 | -3.1451439 | 0.00166005 | 0.01796221 |
| Rnpep | 760.769709 | 817.926597 | 0.93011978 | -0.1045116 | 789.348153 | -0.1046089 | 0.03328943 | -3.1424062 | 0.00167565 | 0.01810121 |
| Ina | 415.720003 | 375.345884 | 1.1075651 | 0.1473915 | 395.532943 | 0.14783913 | 0.04704976 | 3.14218696 | 0.00167691 | 0.01810158 |
| Phf24 | 20.7953398 | 12.8408327 | 1.61946973 | 0.6955215 | 16.8180862 | 0.68392692 | 0.21776104 | 3.1407221 | 0.00168532 | 0.0181681 |
| Mfge8 | 24.6169944 | 35.1279571 | 0.7007807 | -0.512965 | 29.8724757 | -0.5234713 | 0.16667415 | -3.1406866 | 0.00168552 | 0.0181681 |
| Cdh4 | 218.60729 | 248.869276 | 0.87840208 | -0.1870466 | 233.738283 | -0.1874018 | 0.05970579 | -3.1387539 | 0.00169668 | 0.01827506 |
| Klhl26 | 193.812615 | 222.169712 | 0.8723629 | -0.1969997 | 207.991163 | -0.195296 | 0.06229446 | -3.1350459 | 0.00171827 | 0.0184942 |
| Dync1i1 | 150.476845 | 174.716506 | 0.8612629 | -0.2154744 | 162.596675 | -0.2148997 | 0.06856411 | -3.1342885 | 0.00172271 | 0.01852855 |
| Limd1 | 764.963816 | 824.608771 | 0.92766878 | -0.1083183 | 794.786293 | -0.1075356 | 0.03433331 | -3.1321072 | 0.00173556 | 0.01865323 |
| Arsg | 114.082677 | 135.166585 | 0.84401538 | -0.2446588 | 124.624631 | -0.2457216 | 0.07851688 | -3.1295391 | 0.00175081 | 0.01880342 |
| Fchsd1 | 76.4396527 | 93.8169633 | 0.81477432 | -0.2955276 | 85.1283079 | -0.2966961 | 0.09481551 | -3.129194 | 0.00175287 | 0.01881189 |
| Hdlbp | 4350.24542 | 4509.83708 | 0.96461254 | -0.0519785 | 4430.04125 | -0.05199 | 0.01662364 | -3.1274723 | 0.00176316 | 0.01890872 |
| Cdkn1c | 150.365423 | 175.903759 | 0.85481643 | -0.2263135 | 163.134591 | -0.2215932 | 0.07087275 | -3.1266348 | 0.00176819 | 0.01894896 |
| Dhrs7 | 508.827979 | 461.057958 | 1.10360958 | 0.14222989 | 484.942968 | 0.14216392 | 0.04547709 | 3.12605547 | 0.00177168 | 0.01897261 |
| Calm2 | 6956.85385 | 7182.89603 | 0.9685305 | -0.0461306 | 7069.87494 | -0.0465194 | 0.01488897 | -3.1244184 | 0.00178157 | 0.01906472 |
| Shisa4 | 131.431794 | 155.312311 | 0.84624196 | -0.2408579 | 143.372052 | -0.2413228 | 0.07729408 | -3.1221387 | 0.00179542 | 0.01919911 |
| Acss3 | 90.899544 | 110.410916 | 0.82328403 | -0.2805378 | 100.65523 | -0.2765589 | 0.0886075 | -3.1211685 | 0.00180135 | 0.01924859 |
| Tap1 | 212.906513 | 241.712255 | 0.88082631 | -0.1830705 | 227.309384 | -0.1834571 | 0.05878583 | -3.1207715 | 0.00180378 | 0.01926068 |
| Galnt10 | 211.865288 | 241.230801 | 0.87826798 | -0.1872669 | 226.548045 | -0.1876674 | 0.06018044 | -3.1184126 | 0.00181828 | 0.01940154 |
| Gpc2 | 11.0138813 | 5.63782608 | 1.95356884 | 0.96611209 | 8.32585361 | 0.96516709 | 0.30955279 | 3.11794024 | 0.0018212 | 0.01941868 |
| Phf13 | 187.139581 | 214.632841 | 0.87190562 | -0.1977561 | 200.886211 | -0.1985473 | 0.06373438 | -3.11523 | 0.00183802 | 0.01958391 |
| Ddx19b | 438.617233 | 398.415048 | 1.10090529 | 0.13869036 | 418.51614 | 0.13860858 | 0.04451182 | 3.11397271 | 0.00184587 | 0.01965343 |
| Ddx11 | 280.879147 | 248.191703 | 1.1317024 | 0.17849463 | 264.535425 | 0.17978916 | 0.05778318 | 3.11144433 | 0.00186175 | 0.01980828 |
| Pus7 | 341.438017 | 307.316707 | 1.11102979 | 0.1518975 | 324.377362 | 0.15180428 | 0.04879277 | 3.11120451 | 0.00186326 | 0.01981015 |
| Fosl2 | 498.358322 | 544.416415 | 0.91539915 | -0.1275271 | 521.387369 | -0.1273844 | 0.04096556 | -3.1095486 | 0.00187373 | 0.01990726 |
| Tmem97 | 171.298373 | 147.952846 | 1.15779032 | 0.211374 | 159.625609 | 0.21124221 | 0.06793952 | 3.10926843 | 0.00187551 | 0.01991187 |
| Stra6 | 87.2107868 | 70.0822466 | 1.24440627 | 0.31545756 | 78.6465166 | 0.31805145 | 0.10235403 | 3.107366 | 0.00188763 | 0.0200118 |
| Thyn1 | 188.991658 | 163.265003 | 1.15757606 | 0.21110699 | 176.12833 | 0.2111549 | 0.06794963 | 3.10752084 | 0.00188664 | 0.0200118 |
| Las1l | 383.478419 | 347.150523 | 1.10464595 | 0.14358405 | 365.314471 | 0.14268395 | 0.04596622 | 3.10410471 | 0.00190856 | 0.02017757 |
| Spag9 | 1572.06303 | 1479.63754 | 1.06246495 | 0.08741525 | 1525.85029 | 0.08532078 | 0.02748603 | 3.10415009 | 0.00190827 | 0.02017757 |
| Ei24 | 1232.34564 | 1309.89106 | 0.9408001 | -0.0880399 | 1271.11835 | -0.0880437 | 0.02836387 | -3.1040814 | 0.00190871 | 0.02017757 |
| Phactr2 | 8.43411801 | 15.2792441 | 0.55199838 | -0.8572641 | 11.856681 | -0.8561867 | 0.27579539 | -3.1044272 | 0.00190648 | 0.02017757 |
| Mical2 | 498.374929 | 542.919169 | 0.9179542 | -0.1235059 | 520.647049 | -0.1249967 | 0.04027177 | -3.1038284 | 0.00191034 | 0.02018044 |
| Pigw | 134.026706 | 112.58598 | 1.19043869 | 0.25149332 | 123.306343 | 0.25204865 | 0.08123319 | 3.10277911 | 0.00191713 | 0.0202377 |
| Dnttip2 | 400.07462 | 360.342141 | 1.1102632 | 0.15090173 | 380.20838 | 0.1516566 | 0.0488855 | 3.10228208 | 0.00192035 | 0.02025729 |
| Adssl1 | 181.39333 | 210.888788 | 0.86013738 | -0.217361 | 196.141059 | -0.2069403 | 0.06675793 | -3.0998615 | 0.00193611 | 0.02040904 |

| | | | | | | | | | | |
|---|---|---|---|---|---|---|---|---|---|---|
| Zfp950 | 131.441626 | 110.637044 | 1.18804355 | 0.24858772 | 121.039335 | 0.2485091 | 0.08019906 | 3.0986534 | 0.00194402 | 0.02047787 |
| Tspo | 269.036372 | 304.105642 | 0.88468063 | -0.1767714 | 286.571006 | -0.1799823 | 0.05809229 | -3.098214 | 0.00194691 | 0.02049369 |
| 2310033P09 | 140.673165 | 119.078216 | 1.18135096 | 0.24043763 | 129.875691 | 0.23759698 | 0.07672571 | 3.09670616 | 0.00195684 | 0.0205836 |
| Pkdcc | 34.1607074 | 46.233327 | 0.73887625 | -0.4365953 | 40.1970171 | -0.4366479 | 0.14106429 | -3.0953823 | 0.00196559 | 0.02066104 |
| 4930402H24 | 706.946013 | 760.104426 | 0.93006433 | -0.1045976 | 733.52522 | -0.1059111 | 0.03422557 | -3.0945022 | 0.00197143 | 0.02070775 |
| Hook1 | 1327.44118 | 1249.91187 | 1.06202782 | 0.08682156 | 1288.67653 | 0.08702482 | 0.02813357 | 3.09327355 | 0.00197962 | 0.02077895 |
| Top2a | 8254.36079 | 7460.88112 | 1.10635201 | 0.14581048 | 7857.62095 | 0.14553322 | 0.04706111 | 3.09243098 | 0.00198524 | 0.0207927 |
| Ldb1 | 397.225762 | 360.450549 | 1.10202568 | 0.14015784 | 378.838155 | 0.14142619 | 0.04572685 | 3.09284786 | 0.00198246 | 0.0207927 |
| Ubn2 | 1363.63637 | 1258.90818 | 1.08318969 | 0.11528592 | 1311.27228 | 0.10733111 | 0.03470736 | 3.09245993 | 0.00198505 | 0.0207927 |
| Sorbs3 | 39.2868324 | 53.0661688 | 0.7403367 | -0.4337465 | 46.1765005 | -0.4191532 | 0.13555011 | -3.0922378 | 0.00198654 | 0.0207927 |
| Parp4 | 509.892853 | 556.227063 | 0.91669911 | -0.1254798 | 533.059958 | -0.1260699 | 0.04079954 | -3.0899835 | 0.00200168 | 0.02092759 |
| Bcl2l11 | 353.600303 | 390.28373 | 0.90600831 | -0.1424038 | 371.942016 | -0.1418068 | 0.04589367 | -3.0898986 | 0.00200225 | 0.02092759 |
| Top3a | 162.126613 | 138.63435 | 1.16945485 | 0.22583616 | 150.380481 | 0.22535255 | 0.07297519 | 3.08807056 | 0.00201461 | 0.02104192 |
| Plau | 31.0143482 | 42.1688423 | 0.73548019 | -0.4432416 | 36.5915951 | -0.4533538 | 0.14694639 | -3.0851648 | 0.00203439 | 0.02123363 |
| Srsf6 | 2524.03337 | 2413.09215 | 1.04597471 | 0.06484797 | 2468.56276 | 0.06688086 | 0.021684 | 3.08434216 | 0.00204003 | 0.02127745 |
| Dnajb2 | 568.081011 | 617.058267 | 0.92062783 | -0.11931 | 592.569639 | -0.1192677 | 0.03870496 | -3.0814562 | 0.00205991 | 0.02146969 |
| Mrpl21 | 122.27202 | 101.813174 | 1.20094497 | 0.26417004 | 112.042597 | 0.26634237 | 0.0864501 | 3.08087988 | 0.0020639 | 0.02149617 |
| Map3k5 | 354.351891 | 391.569232 | 0.90495336 | -0.1440847 | 372.960561 | -0.144372 | 0.04687221 | -3.08012 | 0.00206917 | 0.02153595 |
| Syndig1 | 12.3701465 | 6.72091415 | 1.84054524 | 0.88013321 | 9.54553024 | 0.89226355 | 0.28977999 | 3.07910688 | 0.00207622 | 0.02159416 |
| Zc2hc1a | 179.965849 | 206.383207 | 0.87199851 | -0.1976024 | 193.174528 | -0.1974162 | 0.06412363 | -3.0786806 | 0.00207919 | 0.02160992 |
| Kidins220 | 1817.07389 | 1914.20133 | 0.94925955 | -0.0751255 | 1865.63761 | -0.0763216 | 0.02480696 | -3.0766188 | 0.00209363 | 0.02174469 |
| Srsf1 | 4234.75005 | 4011.66613 | 1.0556088 | 0.07807528 | 4123.20809 | 0.08216753 | 0.02671614 | 3.07557622 | 0.00210096 | 0.02180557 |
| Shcbp1 | 345.168712 | 310.527754 | 1.11155511 | 0.15257948 | 327.848233 | 0.15951416 | 0.05187536 | 3.07495063 | 0.00210537 | 0.02183607 |
| Cebpa | 119.955879 | 99.4534538 | 1.20615096 | 0.27041048 | 109.704666 | 0.26589247 | 0.08649831 | 3.07396129 | 0.00211237 | 0.02187718 |
| Grik5 | 679.025603 | 730.888661 | 0.9290411 | -0.1061857 | 704.957132 | -0.1060595 | 0.03450281 | -3.0739392 | 0.00211252 | 0.02187718 |
| Arhgef16 | 225.172524 | 255.138682 | 0.88254953 | -0.1802508 | 240.155603 | -0.1802404 | 0.05863833 | -3.073764 | 0.00211377 | 0.02187718 |
| Cbx3 | 2424.26229 | 2322.97402 | 1.04360284 | 0.06157277 | 2373.61815 | 0.06197242 | 0.02018578 | 3.07010335 | 0.00213985 | 0.02213166 |
| Sox2 | 1068.31157 | 1135.69302 | 0.94066931 | -0.0882405 | 1102.00229 | -0.0879595 | 0.02865443 | -3.0696648 | 0.00214299 | 0.02214873 |
| Ilkap | 862.29067 | 921.858986 | 0.9353824 | -0.0963718 | 892.074828 | -0.0963092 | 0.03137933 | -3.0691928 | 0.00214638 | 0.02216829 |
| Lif | 5.97793783 | 2.21020689 | 2.70469604 | 1.43546647 | 4.09407226 | 1.44513173 | 0.47090934 | 3.06881096 | 0.00214913 | 0.02218119 |
| Cd209c | 46.9921425 | 35.1706146 | 1.33611946 | 0.418049 | 41.0813784 | 0.42736366 | 0.13941241 | 3.06546348 | 0.00217333 | 0.02240324 |
| Rab11fip5 | 45.3130232 | 59.5632733 | 0.76075442 | -0.3944973 | 52.4381481 | -0.3981206 | 0.12987484 | -3.0654176 | 0.00217366 | 0.02240324 |
| Glg1 | 1664.32677 | 1761.38184 | 0.94489833 | -0.081769 | 1712.85431 | -0.0852791 | 0.02782163 | -3.0652093 | 0.00217518 | 0.02240326 |
| Ect2 | 1713.88473 | 1592.58465 | 1.07616555 | 0.10590003 | 1653.23469 | 0.11139519 | 0.03635166 | 3.06437752 | 0.00218123 | 0.02245005 |
| Gtf2h3 | 301.788045 | 269.938909 | 1.11798646 | 0.16090271 | 285.863477 | 0.16071907 | 0.05246067 | 3.06361066 | 0.00218683 | 0.02249206 |
| Ampd2 | 637.812951 | 686.595229 | 0.92895046 | -0.1063264 | 662.20409 | -0.1062371 | 0.03468851 | -3.0626013 | 0.00219422 | 0.0225524 |
| Pgr15l | 4.41779758 | 1.15837514 | 3.81378834 | 1.93122478 | 2.78808626 | 1.9322923 | 0.63110443 | 3.06176317 | 0.00220037 | 0.02259997 |
| Hsd17b4 | 541.740972 | 495.580208 | 1.09314489 | 0.12848463 | 518.66059 | 0.12796602 | 0.04180421 | 3.06107976 | 0.0022054 | 0.02263593 |
| Nr2f1 | 2.10967873 | 6.38288953 | 0.33052095 | -1.5971864 | 4.24628403 | -1.5693069 | 0.51288792 | -3.0597462 | 0.00221525 | 0.02272122 |
| Jund | 1405.62303 | 1512.87888 | 0.9291048 | -0.1060868 | 1459.25095 | -0.1053432 | 0.03443176 | -3.0594788 | 0.00221722 | 0.02272579 |
| Dynll2 | 3947.11617 | 3792.14115 | 1.04086742 | 0.05778631 | 3869.62866 | 0.05825911 | 0.01904372 | 3.05922996 | 0.00221907 | 0.02272896 |
| Rps10 | 1436.17491 | 1354.47933 | 1.06031511 | 0.08449308 | 1395.32712 | 0.08373835 | 0.02738692 | 3.05760352 | 0.00223115 | 0.02283688 |
| Cdk4 | 2323.4919 | 2223.40086 | 1.04501709 | 0.06352654 | 2273.44638 | 0.06537435 | 0.02138889 | 3.05646319 | 0.00223965 | 0.02290811 |
| Npm3 | 93.1841313 | 75.4548261 | 1.23496582 | 0.30447111 | 84.3194786 | 0.30381689 | 0.09942666 | 3.05568833 | 0.00224545 | 0.02295155 |
| Akt2 | 1347.69564 | 1428.39917 | 0.94350072 | -0.0839045 | 1388.04741 | -0.0839477 | 0.02748023 | -3.0548382 | 0.00225182 | 0.02300084 |
| Tcirg1 | 723.164628 | 775.610882 | 0.93238071 | -0.1010089 | 749.387755 | -0.1008059 | 0.033003 | -3.0544472 | 0.00225476 | 0.02301498 |
| Hoxb7 | 24.1702038 | 15.9679273 | 1.51367196 | 0.59805258 | 20.0690654 | 0.60341556 | 0.19764148 | 3.05308163 | 0.00226504 | 0.02310406 |
| Hmgcr | 1469.39044 | 1550.38524 | 0.94775827 | -0.0774089 | 1509.88784 | -0.0807714 | 0.02646285 | -3.0522573 | 0.00227127 | 0.02314068 |
| Telo2 | 268.897302 | 300.746695 | 0.89409894 | -0.1614936 | 284.821998 | -0.162061 | 0.05309656 | -3.0521935 | 0.00227176 | 0.02314068 |
| 2900097C17 | 3778.66405 | 3635.02701 | 1.03951471 | 0.05591017 | 3706.84553 | 0.05588995 | 0.01831848 | 3.0510141 | 0.0022807 | 0.02320819 |
| Itgav | 533.912362 | 579.800922 | 0.92085463 | -0.1189547 | 556.856642 | -0.1186003 | 0.03887641 | -3.0507007 | 0.00228308 | 0.02320819 |
| Zfp688 | 43.0786923 | 56.9968461 | 0.75580835 | -0.4039076 | 50.0377691 | -0.4065731 | 0.1332643 | -3.0508779 | 0.00228173 | 0.02320819 |
| Trappc8 | 998.90487 | 1066.08973 | 0.93698011 | -0.0939097 | 1032.4973 | -0.0942292 | 0.0308957 | -3.0499139 | 0.00228907 | 0.02325312 |
| Ube2v2 | 593.895621 | 548.464701 | 1.0828329 | 0.11481062 | 571.180161 | 0.11429975 | 0.03747948 | 3.04966213 | 0.00229099 | 0.02325668 |
| Stim1 | 813.099367 | 873.966806 | 0.93035498 | -0.1041468 | 843.533086 | -0.103635 | 0.03399361 | -3.0486611 | 0.00229864 | 0.02331834 |
| Ndufb8 | 452.333204 | 410.134019 | 1.10289121 | 0.14129049 | 431.233612 | 0.14470604 | 0.047486 | 3.04734132 | 0.00230875 | 0.02340495 |
| Ero1lb | 706.125319 | 759.905012 | 0.9292284 | -0.1058948 | 733.015165 | -0.1056835 | 0.03468675 | -3.0467962 | 0.00231295 | 0.02342639 |
| Npy5r | 85.2137646 | 103.544891 | 0.82296446 | -0.281098 | 94.3793276 | -0.2821327 | 0.09260409 | -3.0466552 | 0.00231403 | 0.02342639 |
| Abcc4 | 13.0859455 | 7.25091148 | 1.8047311 | 0.8517839 | 10.1684284 | 0.90719696 | 0.29785697 | 3.04574697 | 0.00232103 | 0.02348123 |
| Scml4 | 855.193435 | 917.138636 | 0.93245819 | -0.1008891 | 886.166035 | -0.1008056 | 0.03311198 | -3.0443835 | 0.00233158 | 0.02357183 |
| Kcnk1 | 1358.16104 | 1449.96793 | 0.9366835 | -0.0943664 | 1404.06449 | -0.0935333 | 0.03073066 | -3.0436459 | 0.0023373 | 0.02361359 |
| Card6 | 36.8287369 | 48.9221752 | 0.75280252 | -0.4096566 | 42.875456 | -0.4148277 | 0.13630315 | -3.0434197 | 0.00233906 | 0.02361525 |
| Depdc1a | 522.5033 | 479.925742 | 1.08871697 | 0.12262895 | 501.214521 | 0.12274999 | 0.04034487 | 3.04251751 | 0.00234608 | 0.02366243 |
| Cox6c | 985.874615 | 924.992788 | 1.0658187 | 0.09196206 | 955.433701 | 0.09123055 | 0.02998628 | 3.04240933 | 0.00234693 | 0.02366243 |
| Hspe1 | 585.279467 | 541.147524 | 1.08155252 | 0.11310372 | 563.213496 | 0.11343955 | 0.03729792 | 3.04144448 | 0.00235446 | 0.02372226 |
| Mbnl3 | 278.487451 | 312.326721 | 0.89165426 | -0.1654437 | 295.407086 | -0.1641672 | 0.05399757 | -3.04027 | 0.00236366 | 0.02379879 |
| Fryl | 1518.46599 | 1613.87315 | 0.94088311 | -0.0879126 | 1566.16957 | -0.0959049 | 0.03156107 | -3.03871 | 0.00237593 | 0.02390613 |

| | | | | | | | | | | |
|---|---|---|---|---|---|---|---|---|---|---|
| C1qbp | 380.206931 | 342.772263 | 1.10921148 | 0.14953446 | 361.489597 | 0.15165833 | 0.04991304 | 3.03845139 | 0.00237797 | 0.02391042 |
| Kif23 | 1149.39794 | 1076.48172 | 1.06773568 | 0.09455455 | 1112.93983 | 0.09821664 | 0.03233227 | 3.03772787 | 0.00238369 | 0.02393542 |
| Sf3a3 | 1027.99312 | 964.50216 | 1.0658277 | 0.09197423 | 996.247638 | 0.09308235 | 0.03064132 | 3.03780521 | 0.00238308 | 0.02393542 |
| Hcrtr1 | 19.5009185 | 12.23676 | 1.59363413 | 0.67232045 | 15.8688391 | 0.67282852 | 0.22163197 | 3.03579186 | 0.00239905 | 0.02407333 |
| Ston2 | 943.224857 | 881.839962 | 1.06961002 | 0.09708488 | 912.532409 | 0.09658468 | 0.03183229 | 3.03417333 | 0.00241196 | 0.02418649 |
| Bend4 | 699.449683 | 646.684344 | 1.08159365 | 0.11315859 | 673.067014 | 0.11234714 | 0.03703809 | 3.03328657 | 0.00241906 | 0.02424128 |
| Dtx4 | 134.247018 | 112.597738 | 1.19227101 | 0.2537122 | 123.422378 | 0.25267767 | 0.08331077 | 3.03295339 | 0.00242173 | 0.02425166 |
| Nacad | 164.202736 | 189.321301 | 0.86732309 | -0.2053586 | 176.762018 | -0.2050781 | 0.06763562 | -3.0321026 | 0.00242857 | 0.0243037 |
| Xkr5 | 11.9088323 | 6.45512854 | 1.84486369 | 0.88351423 | 9.18198032 | 0.88311673 | 0.29138175 | 3.03078948 | 0.00243915 | 0.02439317 |
| Twnk | 124.710858 | 104.856556 | 1.18934727 | 0.25017002 | 114.783707 | 0.24784712 | 0.08179521 | 3.03009344 | 0.00244478 | 0.02443297 |
| Zfp692 | 152.324992 | 129.231899 | 1.17869499 | 0.23719044 | 140.778445 | 0.2421525 | 0.07992629 | 3.02969764 | 0.00244799 | 0.02444853 |
| Mrgpra6 | 1124.76597 | 1033.07344 | 1.08875703 | 0.12268204 | 1078.91971 | 0.11895377 | 0.03927088 | 3.02905823 | 0.00245317 | 0.02448383 |
| Atg9a | 570.649017 | 617.509348 | 0.92411397 | -0.1138573 | 594.079182 | -0.1142332 | 0.03774027 | -3.0268254 | 0.00247137 | 0.0246488 |
| Usp1 | 1292.58763 | 1219.19806 | 1.06019495 | 0.08432957 | 1255.89285 | 0.08570386 | 0.02832829 | 3.02538075 | 0.0024832 | 0.0247502 |
| Thrb | 78.2496713 | 62.6023192 | 1.24994844 | 0.32186859 | 70.4259952 | 0.32209334 | 0.106521 | 3.02375439 | 0.00249659 | 0.02486691 |
| Pde4d | 251.816264 | 283.532949 | 0.88813757 | -0.1711449 | 267.674606 | -0.1694985 | 0.05607437 | -3.0227453 | 0.00250493 | 0.02493322 |
| Alg13 | 220.714286 | 192.929579 | 1.14401476 | 0.19410567 | 206.821932 | 0.19467939 | 0.06441492 | 3.02227156 | 0.00250885 | 0.02493336 |
| Mtmr7 | 925.971999 | 987.12977 | 0.93804485 | -0.0922712 | 956.550885 | -0.0921387 | 0.03048583 | -3.0223445 | 0.00250825 | 0.02493336 |
| Gigyf2 | 929.702774 | 991.846931 | 0.93734501 | -0.0933479 | 960.774852 | -0.0932546 | 0.030858 | -3.0220576 | 0.00251063 | 0.02493336 |
| Tgfbr3 | 223.557162 | 252.860943 | 0.88411108 | -0.1777004 | 238.209052 | -0.1745828 | 0.05777424 | -3.0218103 | 0.00251268 | 0.02493336 |
| Vtn | 12.98442 | 20.5256161 | 0.63259587 | -0.660644 | 16.7550179 | -0.6674221 | 0.22087424 | -3.0217289 | 0.00251335 | 0.02493336 |
| Ctsz | 872.028558 | 812.42948 | 1.07335908 | 0.10213279 | 842.229019 | 0.10220432 | 0.03383388 | 3.02076824 | 0.00252134 | 0.02499587 |
| Samd1 | 979.055309 | 1042.60628 | 0.93904605 | -0.0907322 | 1010.8308 | -0.0909783 | 0.03012621 | -3.0199067 | 0.00252853 | 0.02505032 |
| Bahcc1 | 671.452333 | 620.524732 | 1.08207183 | 0.11379627 | 645.988532 | 0.11455793 | 0.03795325 | 3.01839548 | 0.00254117 | 0.02514258 |
| Rsl1d1 | 722.103478 | 669.801527 | 1.07808574 | 0.10847192 | 695.952502 | 0.10793247 | 0.03575831 | 3.01838823 | 0.00254123 | 0.02514258 |
| Iars | 1090.35205 | 1024.15175 | 1.06463916 | 0.09036454 | 1057.2519 | 0.09034676 | 0.02993825 | 3.01777053 | 0.00254642 | 0.02517708 |
| Ak4 | 138.942401 | 117.956418 | 1.17791303 | 0.23623302 | 128.449409 | 0.2340272 | 0.07758977 | 3.01621197 | 0.00255954 | 0.02528999 |
| Sestd1 | 342.989768 | 308.682249 | 1.11114186 | 0.15204302 | 325.836008 | 0.15219369 | 0.05049389 | 3.0141011 | 0.00257742 | 0.02544965 |
| Krt18 | 2355.90566 | 2469.44953 | 0.95402058 | -0.0679077 | 2412.6776 | -0.0657208 | 0.02181532 | -3.0125988 | 0.00259021 | 0.02555893 |
| Wfdc2 | 2561.72966 | 2394.87479 | 1.06967165 | 0.09716801 | 2478.30223 | 0.09578104 | 0.03181995 | 3.01009381 | 0.00261167 | 0.02575354 |
| Ccdc25 | 397.944852 | 361.992854 | 1.09931687 | 0.13660729 | 379.968853 | 0.13641031 | 0.04535076 | 3.00789455 | 0.00263064 | 0.02592339 |
| Rgs8 | 101.993251 | 123.423596 | 0.82636752 | -0.2751445 | 112.708423 | -0.2755457 | 0.09170142 | -3.0048143 | 0.00265743 | 0.02616995 |
| Stx5a | 225.643318 | 198.873119 | 1.13460944 | 0.18219577 | 212.258218 | 0.18264308 | 0.06080956 | 3.00352582 | 0.00266871 | 0.02624614 |
| Cers6 | 561.417143 | 518.42405 | 1.08293036 | 0.11494047 | 539.920597 | 0.11493051 | 0.03826275 | 3.00371774 | 0.00266703 | 0.02624614 |
| Cnih2 | 1.50409745 | 0.0000001 | 15040974.5 | 23.8423947 | 0.75204867 | 3.17293985 | 1.05668247 | 3.0027373 | 0.00267563 | 0.02629679 |
| Mtfr1 | 380.233749 | 345.63907 | 1.10008903 | 0.13762028 | 362.93641 | 0.1384053 | 0.04610016 | 3.00227371 | 0.00267971 | 0.02631942 |
| Nploc4 | 1127.89156 | 1063.08795 | 1.0609579 | 0.08536741 | 1095.48976 | 0.08552825 | 0.0284915 | 3.00188616 | 0.00268312 | 0.02633549 |
| Dnajc19 | 140.946787 | 119.72679 | 1.17723683 | 0.23540459 | 130.336789 | 0.23688457 | 0.07892527 | 3.00137816 | 0.00268761 | 0.02636201 |
| Tmx1 | 1575.81763 | 1495.85754 | 1.05345435 | 0.07512779 | 1535.83759 | 0.07543527 | 0.02513698 | 3.00096729 | 0.00269123 | 0.02638015 |
| Slc25a37 | 300.154518 | 266.606204 | 1.12583471 | 0.17099503 | 283.380361 | 0.17171511 | 0.05724219 | 2.99979994 | 0.00270157 | 0.02646396 |
| Abhd11 | 171.939585 | 149.449643 | 1.15048508 | 0.20224228 | 160.694614 | 0.20266628 | 0.06758753 | 2.99857521 | 0.00271245 | 0.02655301 |
| Fbln2 | 74.3217195 | 91.4848005 | 0.81239418 | -0.2997482 | 82.9032599 | -0.299816 | 0.10003269 | -2.9971806 | 0.00272489 | 0.02665718 |
| Spry2 | 223.856923 | 193.398314 | 1.15749159 | 0.21100171 | 208.627618 | 0.20967482 | 0.06998927 | 2.99581394 | 0.00273713 | 0.02675928 |
| Impad1 | 1505.66676 | 1428.57863 | 1.05396142 | 0.07582206 | 1467.1227 | 0.07586015 | 0.02533611 | 2.99415188 | 0.00275209 | 0.02688775 |
| Nrcam | 1404.82427 | 1478.53704 | 0.95014479 | -0.0737807 | 1441.68066 | -0.0737257 | 0.02462608 | -2.9938072 | 0.0027552 | 0.02690041 |
| Atl3 | 679.309839 | 630.86426 | 1.0767924 | 0.10674013 | 655.087049 | 0.1061564 | 0.03548826 | 2.99130998 | 0.00277783 | 0.02710354 |
| Mroh1 | 495.395585 | 541.715244 | 0.91449445 | -0.1289537 | 518.555414 | -0.1260752 | 0.04216607 | -2.9899687 | 0.00279006 | 0.02720492 |
| Zhx2 | 388.523144 | 426.429011 | 0.91110861 | -0.1343051 | 407.476077 | -0.134208 | 0.04497194 | -2.9842612 | 0.00284264 | 0.02769939 |
| Scaf1 | 1061.43551 | 1127.96896 | 0.94101482 | -0.0877107 | 1094.70223 | -0.0880631 | 0.02952331 | -2.9828316 | 0.00285595 | 0.02780999 |
| Add2 | 1014.80727 | 1083.8511 | 0.93629768 | -0.0949608 | 1049.32918 | -0.0944668 | 0.03167223 | -2.9826395 | 0.00285774 | 0.02780999 |
| Med21 | 203.867642 | 177.225284 | 1.15033045 | 0.20204836 | 190.546463 | 0.19853183 | 0.06659205 | 2.98131413 | 0.00287014 | 0.02791232 |
| Pdzd8 | 539.291354 | 495.443201 | 1.08850288 | 0.12234523 | 517.367277 | 0.12092837 | 0.04059896 | 2.97860733 | 0.00289562 | 0.02814159 |
| Scd2 | 3674.16797 | 3518.66937 | 1.04419244 | 0.06238762 | 3596.41867 | 0.05739493 | 0.01927228 | 2.97810865 | 0.00290033 | 0.0281505 |
| Nckap1 | 2616.96315 | 2730.56625 | 0.95839577 | -0.0613066 | 2673.7647 | -0.0606261 | 0.02035617 | -2.9782679 | 0.00289882 | 0.0281505 |
| Pgbd5 | 398.428141 | 439.388727 | 0.90677825 | -0.1411783 | 418.908434 | -0.1398104 | 0.04695748 | -2.9773822 | 0.00290721 | 0.02819883 |
| Cbwd1 | 92.7994393 | 76.4340626 | 1.21411104 | 0.27990037 | 84.6167508 | 0.27962758 | 0.09392871 | 2.97701935 | 0.00291066 | 0.02821363 |
| Med4 | 185.69963 | 161.750655 | 1.14806107 | 0.19919939 | 173.725142 | 0.20191383 | 0.06783782 | 2.9764198 | 0.00291635 | 0.02821363 |
| Dnajc16 | 284.178834 | 316.026337 | 0.89922516 | -0.1532457 | 300.102585 | -0.1531718 | 0.05146012 | -2.9765155 | 0.00291544 | 0.02821363 |
| B3gnt8 | 33.5261378 | 44.7163156 | 0.74975179 | -0.415515 | 39.1212266 | -0.4209932 | 0.14143125 | -2.9766631 | 0.00291404 | 0.02821363 |
| Cep295 | 599.422416 | 552.677212 | 1.08457958 | 0.11713591 | 576.049814 | 0.11870306 | 0.03989027 | 2.97573988 | 0.00292283 | 0.0282578 |
| Top1mt | 76.2436173 | 60.7656822 | 1.25471507 | 0.32735979 | 68.5046496 | 0.32652734 | 0.10975675 | 2.97500908 | 0.0029298 | 0.02830675 |
| Rhot1 | 671.745103 | 622.13544 | 1.07974094 | 0.11068521 | 646.940272 | 0.11066491 | 0.0372087 | 2.97416733 | 0.00293785 | 0.02836602 |
| Timm8b | 132.209153 | 110.847161 | 1.19271573 | 0.25425023 | 121.528157 | 0.25360409 | 0.08527794 | 2.97385339 | 0.00294086 | 0.02837658 |
| Exosc9 | 265.400057 | 237.165105 | 1.11905188 | 0.16227692 | 251.282581 | 0.16331477 | 0.05493889 | 2.97266257 | 0.00295229 | 0.02846836 |
| Zfyve28 | 646.254356 | 697.872648 | 0.9260348 | -0.1108617 | 672.063502 | -0.1098289 | 0.03695491 | -2.9719696 | 0.00295896 | 0.02851414 |
| Pigs | 800.383089 | 746.984237 | 1.07148592 | 0.09961289 | 773.683663 | 0.09952149 | 0.03350495 | 2.97035161 | 0.00297459 | 0.02864614 |
| Cdkn2c | 2316.46572 | 2217.93105 | 1.04442639 | 0.06271082 | 2267.19839 | 0.06351314 | 0.02138551 | 2.96991398 | 0.00297883 | 0.02866835 |

| | | | | | | | | | | |
|---|---|---|---|---|---|---|---|---|---|---|
| Sept10 | 565.222128 | 614.412522 | 0.91993914 | -0.1203897 | 589.817325 | -0.120128 | 0.04045257 | -2.9696013 | 0.00298187 | 0.02867893 |
| Ankrd29 | 140.101029 | 163.441311 | 0.85719472 | -0.2223051 | 151.771169 | -0.2231456 | 0.07520094 | -2.967324 | 0.00300404 | 0.02887348 |
| Rb1 | 2874.1072 | 2754.77342 | 1.04331891 | 0.06118021 | 2814.44031 | 0.06071243 | 0.02046742 | 2.96629678 | 0.00301409 | 0.02895133 |
| Rab15 | 453.662603 | 501.630433 | 0.90437616 | -0.1450051 | 477.646518 | -0.1419614 | 0.04790958 | -2.9631117 | 0.00304546 | 0.02923365 |
| Fat1 | 3900.301 | 4220.73439 | 0.92408113 | -0.1139086 | 4060.5177 | -0.1512503 | 0.05105177 | -2.962684 | 0.0030497 | 0.02925536 |
| Tmx4 | 1375.26912 | 1458.47929 | 0.9429473 | -0.0847509 | 1416.8742 | -0.0838761 | 0.02831347 | -2.9624098 | 0.00305241 | 0.02926249 |
| Smap2 | 1362.99235 | 1453.8545 | 0.93750258 | -0.0931054 | 1408.42343 | -0.0929744 | 0.03138974 | -2.9619375 | 0.0030571 | 0.02928846 |
| Prdx2 | 2972.73626 | 3092.65434 | 0.96122487 | -0.0570541 | 3032.6953 | -0.0573733 | 0.01937192 | -2.9616715 | 0.00305974 | 0.02929484 |
| Ppip5k1 | 817.566541 | 873.215982 | 0.9362707 | -0.0950024 | 845.391262 | -0.0945362 | 0.03192661 | -2.961046 | 0.00306596 | 0.02931652 |
| Sdc3 | 1377.18097 | 1491.20193 | 0.92353754 | -0.1147575 | 1434.19145 | -0.1130018 | 0.03816087 | -2.961196 | 0.00306447 | 0.02931652 |
| Atp13a2 | 491.939526 | 534.381168 | 0.92057796 | -0.1193882 | 513.160347 | -0.1216975 | 0.04110717 | -2.9604925 | 0.00307148 | 0.02935032 |
| Rasd1 | 99.2169443 | 118.165086 | 0.83964686 | -0.2521454 | 108.691015 | -0.2518156 | 0.08509904 | -2.959089 | 0.0030855 | 0.02946533 |
| Apc2 | 278.002481 | 309.502237 | 0.89822446 | -0.1548521 | 293.752359 | -0.1547958 | 0.05231768 | -2.9587672 | 0.00308872 | 0.02947712 |
| Esyt1 | 1725.88945 | 1819.26079 | 0.94867622 | -0.0760123 | 1772.57512 | -0.0759249 | 0.02566405 | -2.9584147 | 0.00309226 | 0.02949186 |
| Erdr1 | 356.74371 | 319.971856 | 1.11492215 | 0.15694298 | 338.357783 | 0.16630873 | 0.05622311 | 2.9580134 | 0.00309629 | 0.0295113 |
| Pde1b | 8.70223814 | 14.804743 | 0.58780069 | -0.766601 | 11.7534905 | -0.7692873 | 0.26030055 | -2.9553812 | 0.00312283 | 0.02974515 |
| Cntln | 626.147034 | 576.39066 | 1.08632404 | 0.11945452 | 601.268847 | 0.12130249 | 0.04107117 | 2.95347043 | 0.00314223 | 0.02989214 |
| Tshz3 | 117.152739 | 138.582677 | 0.84536352 | -0.2423562 | 127.867708 | -0.244298 | 0.08271577 | -2.9534638 | 0.00314229 | 0.02989214 |
| Wsb1 | 602.981004 | 553.691467 | 1.08901986 | 0.12303027 | 578.336235 | 0.13143017 | 0.044523 | 2.95196122 | 0.00315763 | 0.03001871 |
| Cat | 26.0781203 | 18.1739368 | 1.43491862 | 0.52096892 | 22.1260285 | 0.57449129 | 0.19479014 | 2.94928317 | 0.00318512 | 0.0301988 |
| Mrpl55 | 132.55378 | 113.013415 | 1.17290306 | 0.23008378 | 122.783597 | 0.23058343 | 0.07818142 | 2.94933794 | 0.00318456 | 0.0301988 |
| Nt5c3b | 445.600645 | 404.423315 | 1.1018174 | 0.13988515 | 425.01198 | 0.14093061 | 0.04778727 | 2.94912441 | 0.00318676 | 0.0301988 |
| Tmem25 | 197.373885 | 223.258427 | 0.88406018 | -0.1777835 | 210.316156 | -0.1791163 | 0.06072911 | -2.9494314 | 0.00318359 | 0.0301988 |
| Exo5 | 145.177922 | 167.822718 | 0.86506716 | -0.2091159 | 156.50032 | -0.20917 | 0.0709121 | -2.9497075 | 0.00318075 | 0.0301988 |
| Poldip2 | 317.942292 | 285.664097 | 1.11299353 | 0.15444521 | 301.803194 | 0.15582965 | 0.05291508 | 2.94490041 | 0.00323059 | 0.03059458 |
| Fam102a | 297.751203 | 330.802857 | 0.90008655 | -0.1518644 | 314.27703 | -0.1522274 | 0.05172167 | -2.9432031 | 0.00324835 | 0.03074318 |
| Ercc6 | 331.763683 | 297.321903 | 1.11584004 | 0.15813022 | 314.542793 | 0.15749664 | 0.05353304 | 2.9420453 | 0.00326052 | 0.03083867 |
| Dopey2 | 507.637778 | 549.89752 | 0.92314979 | -0.1153633 | 528.767649 | -0.1154283 | 0.03927009 | -2.9393429 | 0.00328909 | 0.03108903 |
| Ndufc2 | 656.862194 | 609.789913 | 1.07719426 | 0.10727845 | 633.326053 | 0.10754731 | 0.03659913 | 2.93852058 | 0.00329783 | 0.03115175 |
| Neurl1a | 470.222719 | 431.646647 | 1.08936956 | 0.12349346 | 450.934683 | 0.12315877 | 0.04192273 | 2.93775679 | 0.00330596 | 0.03120871 |
| Trim36 | 111.861686 | 131.421013 | 0.85117047 | -0.23248 | 121.64135 | -0.2318349 | 0.07897251 | -2.9356411 | 0.00332859 | 0.03140233 |
| Cnbp | 2997.73302 | 2879.37059 | 1.04110705 | 0.05811842 | 2938.55181 | 0.05818613 | 0.01982673 | 2.93473245 | 0.00333835 | 0.03147439 |
| Dpysl5 | 1640.32682 | 1723.10344 | 0.95196074 | -0.071026 | 1681.71513 | -0.0711638 | 0.02425599 | -2.9338654 | 0.00334769 | 0.03154238 |
| Fam78b | 186.154748 | 212.130073 | 0.87755001 | -0.1884467 | 199.14241 | -0.1874424 | 0.06389775 | -2.9334734 | 0.00335192 | 0.03156218 |
| Cap1 | 2507.11107 | 2617.58538 | 0.95779534 | -0.0622107 | 2562.34823 | -0.0621391 | 0.02119013 | -2.9324574 | 0.00336291 | 0.03164552 |
| Rce1 | 400.304797 | 438.035527 | 0.91386377 | -0.129949 | 419.170162 | -0.1308095 | 0.04461877 | -2.9317157 | 0.00337095 | 0.03170107 |
| Dhx15 | 2413.93487 | 2313.93908 | 1.04321453 | 0.06103587 | 2363.93698 | 0.06116899 | 0.02087333 | 2.93048577 | 0.00338432 | 0.03180665 |
| Proser2 | 11.3324638 | 5.96567583 | 1.89961106 | 0.92570406 | 8.64906971 | 0.92473442 | 0.31563592 | 2.92975024 | 0.00339235 | 0.03186182 |
| Gm11837 | 96.8685917 | 114.994471 | 0.84237608 | -0.2474636 | 105.931531 | -0.2473399 | 0.0845143 | -2.9266041 | 0.00342685 | 0.0321655 |
| Dsc2 | 909.085029 | 850.701483 | 1.06862989 | 0.09576227 | 879.893256 | 0.09581101 | 0.03274188 | 2.92625228 | 0.00343073 | 0.03218151 |
| Mphosph9 | 446.515346 | 405.261496 | 1.10179563 | 0.13985665 | 425.888421 | 0.14200657 | 0.04857708 | 2.92332474 | 0.00346315 | 0.03242028 |
| Gna11 | 1226.75863 | 1298.04069 | 0.94508487 | -0.0814842 | 1262.39966 | -0.0816193 | 0.02791967 | -2.9233614 | 0.00346274 | 0.03242028 |
| Arrb1 | 566.006297 | 612.674583 | 0.92382859 | -0.1143029 | 589.34044 | -0.1132521 | 0.03874296 | -2.9231649 | 0.00346493 | 0.03242028 |
| Chst5 | 1.09743026 | 4.15112576 | 0.26436931 | -1.9193734 | 2.62427791 | -1.9035894 | 0.65119656 | -2.9232178 | 0.00346434 | 0.03242028 |
| Caly | 9.62704099 | 16.407626 | 0.58673471 | -0.7692198 | 13.0174334 | -0.7626389 | 0.26095636 | -2.9224766 | 0.0034726 | 0.03247152 |
| Il13ra1 | 243.069088 | 215.006829 | 1.13051799 | 0.17698395 | 229.037959 | 0.17944423 | 0.061421 | 2.92154547 | 0.00348299 | 0.03253038 |
| Nlrc4 | 5.20736968 | 10.627005 | 0.49001291 | -1.0291083 | 7.91718723 | -1.0138249 | 0.34701967 | -2.9215198 | 0.00348328 | 0.03253038 |
| Srek1 | 513.372863 | 469.641151 | 1.09311729 | 0.12844821 | 491.507007 | 0.13046206 | 0.04470328 | 2.91840054 | 0.00351832 | 0.03281626 |
| Pacsin1 | 429.233367 | 469.114446 | 0.91498646 | -0.1281877 | 449.173906 | -0.1275891 | 0.04371753 | -2.9184874 | 0.00351734 | 0.03281626 |
| Snrnp70 | 1745.48202 | 1649.1393 | 1.05842 | 0.08191222 | 1697.31066 | 0.08727738 | 0.02992052 | 2.91697354 | 0.00353446 | 0.03292533 |
| Rdx | 3262.07089 | 3140.52627 | 1.03870199 | 0.0547818 | 3201.29858 | 0.05492829 | 0.01882984 | 2.91708801 | 0.00353316 | 0.03292533 |
| Synpr | 2.01787233 | 0.22737211 | 8.87475742 | 3.14970769 | 1.12262212 | 2.77905032 | 0.95304128 | 2.91598105 | 0.00354572 | 0.03296774 |
| Ppp2r3a | 39.0714491 | 28.4007318 | 1.3757198 | 0.46018666 | 33.7360903 | 0.46011675 | 0.15777472 | 2.91628937 | 0.00354222 | 0.03296774 |
| Grm5 | 185.779314 | 212.015113 | 0.87625505 | -0.1905772 | 198.897213 | -0.1886029 | 0.06468015 | -2.9159324 | 0.00354627 | 0.03296774 |
| Ccser1 | 153.867569 | 177.170779 | 0.86847035 | -0.2034515 | 165.519174 | -0.2031645 | 0.06967738 | -2.9157886 | 0.00354791 | 0.03296774 |
| Rras | 33.2805307 | 45.4328383 | 0.7325215 | -0.449057 | 39.3566844 | -0.4529583 | 0.15538789 | -2.9150166 | 0.0035567 | 0.03302869 |
| Setdb1 | 793.703555 | 738.170832 | 1.07523018 | 0.10464553 | 765.937194 | 0.10481432 | 0.03596171 | 2.91460916 | 0.00356134 | 0.03303045 |
| Wtip | 126.13704 | 146.775139 | 0.85938968 | -0.2186156 | 136.456089 | -0.217363 | 0.07457528 | -2.9146789 | 0.00356055 | 0.03303045 |
| Abca3 | 1149.56316 | 1221.38722 | 0.94119469 | -0.0874349 | 1185.47519 | -0.0863375 | 0.02962824 | -2.9140272 | 0.00356799 | 0.03307139 |
| Ccnj | 65.5186037 | 51.6547497 | 1.26839456 | 0.3430036 | 58.5866766 | 0.34318602 | 0.11782127 | 2.91276795 | 0.00358241 | 0.03318428 |
| Ackr3 | 6.62337997 | 2.69479127 | 2.45784527 | 1.29739409 | 4.65908552 | 1.25688484 | 0.43161661 | 2.91203999 | 0.00359077 | 0.03319947 |
| Eef1e1 | 188.264352 | 164.230846 | 1.14633978 | 0.19703473 | 176.247598 | 0.19978222 | 0.06860103 | 2.91223333 | 0.00358854 | 0.03319947 |
| Hcfc1r1 | 253.73937 | 283.668189 | 0.89449357 | -0.160857 | 268.703779 | -0.1614765 | 0.05544858 | -2.9121846 | 0.0035891 | 0.03319947 |
| Ormdl2 | 163.575461 | 141.909884 | 1.15267138 | 0.20498126 | 152.742672 | 0.20671007 | 0.07100783 | 2.91108862 | 0.00360172 | 0.03326402 |
| Nptn | 2378.1996 | 2487.34768 | 0.95611869 | -0.0647384 | 2432.77364 | -0.0660831 | 0.02270082 | -2.9110437 | 0.00360224 | 0.03326402 |
| Plk1 | 1018.63174 | 955.348679 | 1.0662408 | 0.0925333 | 986.990212 | 0.09569214 | 0.03288266 | 2.91010912 | 0.00361303 | 0.03334287 |
| Gm14403 | 7.32577826 | 3.2186284 | 2.27605593 | 1.18653601 | 5.27220323 | 1.19239646 | 0.40982738 | 2.90950904 | 0.00361997 | 0.03338616 |

| | | | | | | | | | | |
|---|---|---|---|---|---|---|---|---|---|---|
| Rad9a | 452.878451 | 492.247054 | 0.92002267 | -0.1202587 | 472.562753 | -0.120248 | 0.04133428 | -2.9091596 | 0.00362402 | 0.03340272 |
| Slc36a1 | 1328.21239 | 1256.44654 | 1.05711811 | 0.08013657 | 1292.32946 | 0.08016449 | 0.02755946 | 2.9087829 | 0.00362839 | 0.0334222 |
| Lrrc24 | 293.672019 | 328.400315 | 0.89425011 | -0.1612497 | 311.036167 | -0.1600226 | 0.05505242 | -2.9067317 | 0.00365226 | 0.03362123 |
| Acadl | 1046.74211 | 1112.0553 | 0.94126804 | -0.0873225 | 1079.39871 | -0.0894188 | 0.03076648 | -2.906371 | 0.00365648 | 0.03363913 |
| Snrpd3 | 550.418702 | 509.58315 | 1.08013521 | 0.11121192 | 530.000926 | 0.1125133 | 0.03874928 | 2.90362274 | 0.00368872 | 0.03389372 |
| Plxna2 | 2066.36278 | 2192.80069 | 0.94233953 | -0.0856811 | 2129.58174 | -0.091267 | 0.03143014 | -2.9038064 | 0.00368656 | 0.03389372 |
| Stip1 | 1394.51132 | 1312.85935 | 1.062194 | 0.08704728 | 1353.68533 | 0.08704258 | 0.02998288 | 2.90307574 | 0.00369517 | 0.03393194 |
| Nomo1 | 1533.52924 | 1614.64299 | 0.94976366 | -0.0743595 | 1574.08611 | -0.0731887 | 0.02521513 | -2.9025697 | 0.00370115 | 0.03396577 |
| Atp8a1 | 1617.84769 | 1699.25999 | 0.95208956 | -0.0708308 | 1658.55384 | -0.0719597 | 0.02479393 | -2.9023129 | 0.00370418 | 0.03397258 |
| Ndufb5 | 406.604952 | 371.795817 | 1.09362433 | 0.12911725 | 389.200384 | 0.12874593 | 0.0443856 | 2.9006239 | 0.00372421 | 0.0341351 |
| Egr1 | 539.271334 | 441.686508 | 1.22093685 | 0.28798858 | 490.478921 | 0.32650457 | 0.11260784 | 2.89948336 | 0.00373778 | 0.03423836 |
| Cldn7 | 1038.56718 | 977.248045 | 1.06274675 | 0.08779784 | 1007.90761 | 0.08798218 | 0.03034857 | 2.8990555 | 0.00374289 | 0.03426394 |
| Pianp | 709.203778 | 778.153894 | 0.9113927 | -0.1338553 | 743.678836 | -0.1283128 | 0.04427834 | -2.8978673 | 0.0037571 | 0.03437278 |
| Lig3 | 530.514192 | 484.626445 | 1.09468684 | 0.13051821 | 507.570318 | 0.12874035 | 0.04443462 | 2.89729855 | 0.00376391 | 0.03441392 |
| Tmem109 | 327.267057 | 295.305376 | 1.10823264 | 0.14826077 | 311.286216 | 0.14947005 | 0.05159986 | 2.89671429 | 0.00377093 | 0.03445682 |
| Scly | 607.093436 | 658.765506 | 0.92156227 | -0.1178464 | 632.929471 | -0.117618 | 0.0406615 | -2.8926129 | 0.00382052 | 0.03488841 |
| Mrps7 | 325.138806 | 293.990479 | 1.10595012 | 0.14528632 | 309.564642 | 0.14560617 | 0.05035047 | 2.89185327 | 0.00382977 | 0.03490836 |
| Sorbs2 | 734.33381 | 682.4768 | 1.07598355 | 0.10565602 | 708.405305 | 0.10617684 | 0.03671135 | 2.89220751 | 0.00382545 | 0.03490836 |
| 4933412E12I | 77.1494623 | 93.6023297 | 0.82422588 | -0.2788883 | 85.3758959 | -0.2759891 | 0.09543196 | -2.8919992 | 0.00382799 | 0.03490836 |
| Tprgl | 533.050879 | 575.5403 | 0.92617472 | -0.1106437 | 554.295589 | -0.1105281 | 0.03822955 | -2.8911697 | 0.00383811 | 0.03496288 |
| Ncor1 | 2879.82324 | 2733.52683 | 1.05351929 | 0.07521673 | 2806.67503 | 0.0713134 | 0.02467035 | 2.89065229 | 0.00384443 | 0.03498381 |
| Aak1 | 837.739478 | 892.299143 | 0.93885496 | -0.0910258 | 865.01931 | -0.0924339 | 0.03197748 | -2.8905955 | 0.00384513 | 0.03498381 |
| Ncor2 | 2030.42674 | 2142.50335 | 0.94768894 | -0.0775145 | 2086.46505 | -0.0848427 | 0.02936299 | -2.8894443 | 0.00385923 | 0.03504761 |
| Sdc2 | 306.365695 | 340.83222 | 0.89887539 | -0.153807 | 323.598958 | -0.151408 | 0.05239968 | -2.8894832 | 0.00385876 | 0.03504761 |
| Dkk3 | 1.34501378 | 4.75072848 | 0.28311738 | -1.8205278 | 3.04787103 | -1.8259966 | 0.63194171 | -2.8895016 | 0.00385853 | 0.03504761 |
| R3hdm1 | 1522.66022 | 1600.6673 | 0.9512659 | -0.0720794 | 1561.66376 | -0.071933 | 0.024909 | -2.8878325 | 0.00387906 | 0.03520613 |
| Pprc1 | 199.128742 | 173.17132 | 1.14989446 | 0.20150146 | 186.150031 | 0.20167678 | 0.06987389 | 2.88629652 | 0.00389805 | 0.03533514 |
| Ppp2ca | 2391.40953 | 2270.61776 | 1.05319776 | 0.07477635 | 2331.01365 | 0.07474234 | 0.02589479 | 2.88638499 | 0.00389695 | 0.03533514 |
| Elavl4 | 799.928608 | 747.522789 | 1.07010598 | 0.09775369 | 773.725698 | 0.09784278 | 0.03391419 | 2.88500982 | 0.00391401 | 0.03543654 |
| Heatr5b | 466.395758 | 506.217633 | 0.92133448 | -0.1182031 | 486.306695 | -0.1204168 | 0.04173727 | -2.8851153 | 0.0039127 | 0.03543654 |
| Usp53 | 459.602332 | 499.699756 | 0.91975697 | -0.1206754 | 479.651044 | -0.1187621 | 0.04116851 | -2.8847799 | 0.00391687 | 0.03544078 |
| Eif1ax | 1675.79909 | 1586.29438 | 1.05642377 | 0.07918867 | 1631.04673 | 0.08028217 | 0.02784107 | 2.88358775 | 0.00393173 | 0.0355535 |
| Iqgap3 | 658.402958 | 600.985768 | 1.09553835 | 0.13163999 | 629.694363 | 0.13671791 | 0.047427 | 2.88270213 | 0.0039428 | 0.03563186 |
| Pof1b | 237.402957 | 266.918242 | 0.889422 | -0.16906 | 252.160599 | -0.1677706 | 0.05820507 | -2.8824053 | 0.00394652 | 0.03564371 |
| Hmgb1 | 5816.16386 | 5644.22968 | 1.03046194 | 0.04329122 | 5730.19677 | 0.04411096 | 0.01531488 | 2.88026762 | 0.00397338 | 0.03586445 |
| Magee1 | 350.282423 | 385.597614 | 0.90841439 | -0.1385775 | 367.940018 | -0.137398 | 0.0477171 | -2.8794286 | 0.00398397 | 0.03593813 |
| Ppp2r2d | 525.077966 | 485.093288 | 1.08242678 | 0.11426944 | 505.085627 | 0.11420251 | 0.03968767 | 2.87753125 | 0.004008 | 0.03613297 |
| Adamts10 | 85.9801294 | 103.133064 | 0.83368152 | -0.2624317 | 94.5565964 | -0.2662936 | 0.09258716 | -2.8761393 | 0.00402572 | 0.03627064 |
| Mif | 916.221124 | 860.825234 | 1.06435207 | 0.08997546 | 888.523179 | 0.09061857 | 0.03151271 | 2.87561934 | 0.00403236 | 0.03630836 |
| Eif1 | 3271.81545 | 3143.37812 | 1.04085965 | 0.05777555 | 3207.59679 | 0.05789705 | 0.02013602 | 2.87529811 | 0.00403646 | 0.03632326 |
| Anp32e | 3705.87556 | 3568.53041 | 1.03848788 | 0.05448437 | 3637.20298 | 0.05644136 | 0.01965653 | 2.87137918 | 0.00408685 | 0.03675437 |
| Pnma1 | 26.2323432 | 35.8473642 | 0.73177886 | -0.4505204 | 31.0398536 | -0.450913 | 0.15706997 | -2.8707777 | 0.00409463 | 0.03680205 |
| Ss18 | 1097.12945 | 1037.19294 | 1.05778723 | 0.08104947 | 1067.1612 | 0.08139602 | 0.0283608 | 2.87001883 | 0.00410447 | 0.03686812 |
| Vps13a | 1032.14298 | 955.853973 | 1.07981241 | 0.1107807 | 993.998478 | 0.10169139 | 0.03543685 | 2.86965108 | 0.00410925 | 0.03688867 |
| Fkbp3 | 521.795172 | 482.566077 | 1.08129269 | 0.1127571 | 502.180624 | 0.11287306 | 0.03933925 | 2.86922262 | 0.00411482 | 0.03691632 |
| Ccna2 | 2135.84094 | 2038.48819 | 1.04775733 | 0.06730461 | 2087.16457 | 0.06812674 | 0.02374645 | 2.86892267 | 0.00411872 | 0.03692899 |
| Sprr2k | 4.66519033 | 1.55760442 | 2.995106 | 1.58260706 | 3.11139727 | 1.59766499 | 0.5570842 | 2.86790576 | 0.00413199 | 0.03702549 |
| Asf1b | 623.005333 | 670.059994 | 0.92977545 | -0.1050458 | 646.532664 | -0.1054132 | 0.03676542 | -2.8671831 | 0.00414143 | 0.03708772 |
| Uimc1 | 269.814638 | 241.660058 | 1.1165049 | 0.15898958 | 255.737348 | 0.1585471 | 0.05532474 | 2.86575413 | 0.00416017 | 0.03723304 |
| Cth | 4.36274111 | 1.47001039 | 2.96783012 | 1.56940851 | 2.91637565 | 1.56129101 | 0.54506045 | 2.86443645 | 0.00417752 | 0.0373432 |
| Adam28 | 65.5223901 | 52.4080971 | 1.2502341 | 0.32219826 | 58.9652435 | 0.32101106 | 0.11206418 | 2.86452862 | 0.0041763 | 0.0373432 |
| Nsd1 | 1721.21148 | 1813.30497 | 0.94921236 | -0.0751972 | 1767.25822 | -0.0788394 | 0.02753009 | -2.8637538 | 0.00418653 | 0.03735617 |
| Smyd3 | 204.73286 | 233.682501 | 0.87611549 | -0.190807 | 219.20768 | -0.179919 | 0.06282128 | -2.8639819 | 0.00418352 | 0.03735617 |
| Abat | 517.754403 | 612.13148 | 0.84582221 | -0.2415736 | 564.942941 | -0.2031853 | 0.07094654 | -2.8639204 | 0.00418433 | 0.03735617 |
| Gtf2ird1 | 508.141179 | 550.130652 | 0.92367363 | -0.1145449 | 529.135916 | -0.1144607 | 0.03997716 | -2.8631515 | 0.0041945 | 0.03740474 |
| Ccdc177 | 25.5034329 | 17.2656197 | 1.47712236 | 0.56278934 | 21.3845262 | 0.56335868 | 0.19682873 | 2.86217702 | 0.00420742 | 0.03749738 |
| Zswim1 | 147.614241 | 126.520043 | 1.16672613 | 0.22246596 | 137.067142 | 0.226802 | 0.07925333 | 2.86173457 | 0.0042133 | 0.0375272 |
| Zfp367 | 1017.33969 | 1076.9011 | 0.94469185 | -0.0820843 | 1047.12039 | -0.0822133 | 0.02874295 | -2.8602926 | 0.0042325 | 0.03767563 |
| Esyt2 | 1306.37776 | 1373.70899 | 0.95098581 | -0.0725043 | 1340.04337 | -0.0725348 | 0.02536799 | -2.8593031 | 0.00424573 | 0.03777068 |
| Psmb6 | 907.18882 | 849.746356 | 1.06759954 | 0.09437059 | 878.467588 | 0.09590695 | 0.03354638 | 2.85893556 | 0.00425065 | 0.03779178 |
| Crtap | 150.967229 | 175.374834 | 0.86082607 | -0.2162063 | 163.171032 | -0.2169162 | 0.07590863 | -2.8575959 | 0.00426864 | 0.03792893 |
| Napb | 345.187082 | 311.969251 | 1.1064779 | 0.14597464 | 328.578167 | 0.1460839 | 0.05113277 | 2.85695237 | 0.0042773 | 0.03798314 |
| Tox3 | 244.865598 | 273.04117 | 0.89680834 | -0.1571284 | 258.953384 | -0.1565872 | 0.05486475 | -2.8540577 | 0.00431647 | 0.03829627 |
| Efemp1 | 9.61256224 | 15.8337561 | 0.60709298 | -0.7200106 | 12.7231591 | -0.7202607 | 0.25237195 | -2.853965 | 0.00431773 | 0.03829627 |
| Melk | 538.763052 | 497.795839 | 1.08229722 | 0.11409674 | 518.279445 | 0.1170172 | 0.04101342 | 2.85314433 | 0.0043289 | 0.03837235 |
| Rhbdl3 | 8.66334783 | 4.23744518 | 2.04447431 | 1.03172994 | 6.45039641 | 1.04841131 | 0.36776921 | 2.85073158 | 0.00436188 | 0.03861197 |
| Gps1 | 934.922386 | 879.618615 | 1.06287244 | 0.08796847 | 907.270501 | 0.08832381 | 0.03098539 | 2.85049838 | 0.00436508 | 0.03861197 |

| | | | | | | | | | | |
|---|---|---|---|---|---|---|---|---|---|---|
| Uhrf1 | 1360.67235 | 1448.4516 | 0.93939787 | -0.0901918 | 1404.56198 | -0.0865394 | 0.03036091 | -2.8503565 | 0.00436703 | 0.03861197 |
| Maob | 103.783213 | 123.689947 | 0.8390594 | -0.2531551 | 113.73658 | -0.2604343 | 0.09137352 | -2.850216 | 0.00436895 | 0.03861197 |
| Lgr6 | 41.9003509 | 53.934022 | 0.77688163 | -0.3642333 | 47.9171863 | -0.3672765 | 0.12883669 | -2.8507139 | 0.00436212 | 0.03861197 |
| Bzw2 | 1012.11506 | 954.121754 | 1.06078187 | 0.08512802 | 983.118405 | 0.08483703 | 0.02977538 | 2.84923407 | 0.00438246 | 0.03870827 |
| Ran | 4207.34232 | 4070.07251 | 1.03372662 | 0.0478547 | 4138.70741 | 0.04911418 | 0.01724249 | 2.84843958 | 0.00439342 | 0.03878194 |
| Ptrh2 | 195.653069 | 172.13947 | 1.13659621 | 0.18471981 | 183.89627 | 0.18486336 | 0.06491306 | 2.84786106 | 0.00440141 | 0.03882938 |
| Nup88 | 930.673279 | 871.364992 | 1.06806366 | 0.09499763 | 901.019135 | 0.09504822 | 0.0333787 | 2.84757064 | 0.00440543 | 0.0388417 |
| Cntd1 | 127.081521 | 108.363268 | 1.17273613 | 0.22987844 | 117.722394 | 0.2300206 | 0.08080565 | 2.84659034 | 0.00441902 | 0.03893122 |
| 0610009B22 | 155.695306 | 134.781695 | 1.15516655 | 0.20810088 | 145.2385 | 0.20865274 | 0.07330556 | 2.8463427 | 0.00442246 | 0.03893122 |
| Csk | 594.288333 | 641.278434 | 0.92672434 | -0.1097878 | 617.783384 | -0.1100776 | 0.03867432 | -2.8462702 | 0.00442346 | 0.03893122 |
| D430041D05 | 246.643034 | 218.304933 | 1.12980971 | 0.17607981 | 232.473984 | 0.17197621 | 0.06045886 | 2.84451597 | 0.0044479 | 0.03911134 |
| Hspa9 | 2732.29204 | 2626.01278 | 1.04047172 | 0.05723776 | 2679.15241 | 0.05727448 | 0.02013572 | 2.84442193 | 0.00444921 | 0.03911134 |
| Snrpc | 528.478831 | 487.566909 | 1.08391038 | 0.11624547 | 508.02287 | 0.11787117 | 0.04145079 | 2.84364139 | 0.00446012 | 0.03916082 |
| Epn3 | 50.4933548 | 63.8619061 | 0.7906647 | -0.3388621 | 57.1776303 | -0.336702 | 0.11840084 | -2.8437466 | 0.00445865 | 0.03916082 |
| Cdk5rap3 | 284.857101 | 254.495469 | 1.11930127 | 0.1625984 | 269.676285 | 0.16282237 | 0.0572881 | 2.84216724 | 0.0044808 | 0.03931907 |
| Chn1 | 1079.26701 | 1149.40781 | 0.93897658 | -0.0908389 | 1114.33741 | -0.0922512 | 0.03247215 | -2.840933 | 0.00449818 | 0.0394482 |
| Iqsec1 | 966.0086 | 1033.07536 | 0.93508048 | -0.0968376 | 999.541979 | -0.0940962 | 0.03314837 | -2.8386381 | 0.00453065 | 0.0397095 |
| Myl12a | 856.78354 | 802.406308 | 1.0677677 | 0.09459782 | 829.594924 | 0.0952792 | 0.03357223 | 2.83803614 | 0.0045392 | 0.03973748 |
| Wipf3 | 487.392141 | 527.21561 | 0.92446455 | -0.1133101 | 507.303875 | -0.1145829 | 0.04037161 | -2.8382043 | 0.00453681 | 0.03973748 |
| Ankrd12 | 625.841673 | 580.044378 | 1.07895481 | 0.10963445 | 602.943026 | 0.10972074 | 0.03868411 | 2.83632558 | 0.00456359 | 0.03992737 |
| Micall2 | 79.4908575 | 96.099549 | 0.82717201 | -0.2737407 | 87.7952032 | -0.2765997 | 0.09754348 | -2.8356554 | 0.00457318 | 0.03998763 |
| Nhp2 | 191.167141 | 167.840668 | 1.13897987 | 0.18774225 | 179.503905 | 0.19056582 | 0.06721903 | 2.83499797 | 0.0045826 | 0.04004639 |
| Tead3 | 6.13224508 | 2.26453893 | 2.70794421 | 1.43719802 | 4.1983919 | 1.42644945 | 0.50352754 | 2.83291249 | 0.0046126 | 0.04028483 |
| Atl2 | 641.705116 | 594.269475 | 1.07982177 | 0.11079321 | 617.987295 | 0.11176785 | 0.03951908 | 2.82819972 | 0.00468106 | 0.04081058 |
| Ska2 | 560.085869 | 519.092399 | 1.07897143 | 0.10965667 | 539.589134 | 0.11095578 | 0.03922732 | 2.82853287 | 0.00467619 | 0.04081058 |
| Csmd1 | 2.55878685 | 6.23703919 | 0.41025666 | -1.2854014 | 4.39791292 | -1.3075764 | 0.46232839 | -2.8282416 | 0.00468045 | 0.04081058 |
| Vti1a | 165.367638 | 143.930165 | 1.14894357 | 0.20030795 | 154.648901 | 0.20004199 | 0.07074415 | 2.82768232 | 0.00468863 | 0.04085256 |
| Dlg2 | 15.7872246 | 23.4384684 | 0.67356042 | -0.5701207 | 19.6128464 | -0.5710609 | 0.2019884 | -2.8271964 | 0.00469575 | 0.04089056 |
| Exph5 | 154.737226 | 176.979445 | 0.87432315 | -0.1937615 | 165.858336 | -0.1945481 | 0.06886484 | -2.8250719 | 0.004727 | 0.04113851 |
| Mex3d | 436.057647 | 476.709554 | 0.91472395 | -0.1285917 | 456.3836 | -0.1294118 | 0.04582108 | -2.8242846 | 0.00473863 | 0.0412155 |
| Il22ra1 | 330.262268 | 365.042325 | 0.90472322 | -0.1444516 | 347.652296 | -0.143719 | 0.05094676 | -2.8209644 | 0.00478795 | 0.04162008 |
| Lingo3 | 11.761306 | 18.4133606 | 0.63873761 | -0.6467047 | 15.0873332 | -0.646311 | 0.22913235 | -2.8206885 | 0.00479207 | 0.04163147 |
| Hirip3 | 447.615703 | 405.80688 | 1.1030264 | 0.14146733 | 426.711291 | 0.15220109 | 0.05396828 | 2.82019514 | 0.00479945 | 0.04164671 |
| Lap3 | 385.440554 | 351.350086 | 1.09702706 | 0.13359912 | 368.39532 | 0.13470178 | 0.04776016 | 2.82037966 | 0.00479669 | 0.04164671 |
| Mapk9 | 633.925401 | 589.777949 | 1.07485436 | 0.10414119 | 611.851675 | 0.10492062 | 0.03720626 | 2.81997258 | 0.00480278 | 0.04165122 |
| Mycbp2 | 2266.07673 | 2363.25671 | 0.95887879 | -0.0605796 | 2314.66672 | -0.0680566 | 0.02415321 | -2.817706 | 0.00483681 | 0.04189732 |
| Itpr2 | 20.268805 | 28.5628694 | 0.70962076 | -0.4948799 | 24.4158371 | -0.4953376 | 0.17578658 | -2.8178351 | 0.00483486 | 0.04189732 |
| Apcdd1 | 140.353052 | 161.610642 | 0.86846417 | -0.2034618 | 150.981847 | -0.2037537 | 0.07232356 | -2.8172521 | 0.00484365 | 0.04193208 |
| Acvr1 | 434.876224 | 473.918807 | 0.91761757 | -0.1240351 | 454.397515 | -0.1242128 | 0.04410154 | -2.8165192 | 0.00485471 | 0.04200334 |
| Gpr83 | 4.77152184 | 9.89217803 | 0.48235301 | -1.0518387 | 7.33184983 | -1.0493598 | 0.37260935 | -2.8162465 | 0.00485884 | 0.04201449 |
| Rps20 | 902.581437 | 848.224972 | 1.0640826 | 0.08961015 | 875.403204 | 0.08973495 | 0.0318684 | 2.8157971 | 0.00486564 | 0.04202426 |
| Gpld1 | 774.290423 | 829.39305 | 0.93356271 | -0.0991812 | 801.841737 | -0.0974298 | 0.03460028 | -2.8158671 | 0.00486458 | 0.04202426 |
| Calu | 652.668567 | 608.612225 | 1.0723882 | 0.10082725 | 630.640396 | 0.1019695 | 0.03621696 | 2.81551772 | 0.00486987 | 0.04203632 |
| Myo6 | 334.039359 | 367.664325 | 0.90854439 | -0.1383711 | 350.851842 | -0.1361659 | 0.04837097 | -2.8150339 | 0.00487721 | 0.04207516 |
| Prkaca | 1508.2363 | 1580.42102 | 0.95432564 | -0.0674465 | 1544.32866 | -0.0674298 | 0.02395899 | -2.8143835 | 0.00488709 | 0.04211137 |
| Tbc1d32 | 400.756064 | 437.313515 | 0.91640448 | -0.1259436 | 419.034789 | -0.1254938 | 0.04458832 | -2.8144989 | 0.00488533 | 0.04211137 |
| Psg16 | 107.062984 | 89.62218 | 1.19460366 | 0.25653204 | 98.342582 | 0.25651879 | 0.09116109 | 2.81390657 | 0.00489435 | 0.04214938 |
| Bmp1 | 308.601332 | 274.466647 | 1.12436733 | 0.16911344 | 291.533989 | 0.19892152 | 0.07069983 | 2.81360671 | 0.00489891 | 0.04216422 |
| Zfp799 | 178.853191 | 202.845875 | 0.88171964 | -0.1816081 | 190.849533 | -0.1812549 | 0.06446225 | -2.8117993 | 0.00492652 | 0.04237724 |
| Srsf2 | 2714.11992 | 2585.25011 | 1.0498481 | 0.0701806 | 2649.68501 | 0.07264128 | 0.02584566 | 2.81057902 | 0.00494524 | 0.04251361 |
| Adamts1 | 1.60207285 | 0.0000001 | 16020728.5 | 23.9334364 | 0.80103638 | 3.03917567 | 1.08160676 | 2.80987119 | 0.00495613 | 0.04255785 |
| Prcc | 387.611917 | 422.236475 | 0.91799723 | -0.1234383 | 404.924196 | -0.1236813 | 0.04401635 | -2.809894 | 0.00495578 | 0.04255785 |
| Snx5 | 737.121212 | 689.58042 | 1.06894162 | 0.09618307 | 713.350816 | 0.09649488 | 0.03434368 | 2.80968366 | 0.00495902 | 0.042558 |
| Ubap2 | 1228.79411 | 1308.8642 | 0.93882475 | -0.0910722 | 1268.82915 | -0.0899435 | 0.03201425 | -2.8094847 | 0.00496209 | 0.04255967 |
| Cnst | 717.984441 | 769.081891 | 0.93356046 | -0.0991846 | 743.533165 | -0.0986326 | 0.03511854 | -2.8085625 | 0.00497632 | 0.04265707 |
| Csf1 | 49.3305683 | 62.0716658 | 0.79473569 | -0.331453 | 55.7011169 | -0.33239 | 0.11839546 | -2.8074553 | 0.00499346 | 0.04277924 |
| Adcy2 | 372.995547 | 407.683257 | 0.91491505 | -0.1282903 | 390.339402 | -0.1269887 | 0.04525041 | -2.8063544 | 0.00501056 | 0.04290089 |
| Kat7 | 1175.75495 | 1107.42691 | 1.06169982 | 0.08637592 | 1141.59093 | 0.08632296 | 0.03078809 | 2.80377786 | 0.00505077 | 0.04322024 |
| Frmd8 | 147.58911 | 126.829862 | 1.16367792 | 0.2186918 | 137.209486 | 0.21918152 | 0.07821003 | 2.80247342 | 0.00507124 | 0.04337036 |
| Cox20 | 118.145596 | 137.626235 | 0.85844858 | -0.2201964 | 127.886235 | -0.2220738 | 0.07925615 | -2.8019758 | 0.00507907 | 0.04341227 |
| Mapk4 | 251.260596 | 280.561136 | 0.89556451 | -0.1591307 | 265.910866 | -0.1575541 | 0.05624177 | -2.8013721 | 0.00508858 | 0.04344346 |
| Zmat4 | 35.3519371 | 46.0626189 | 0.76747562 | -0.3818072 | 40.7072779 | -0.3812513 | 0.13609307 | -2.8014009 | 0.00508813 | 0.04344346 |
| Fam118a | 202.73311 | 178.700789 | 1.13448358 | 0.18203573 | 190.716949 | 0.18393239 | 0.06568677 | 2.80014369 | 0.00510799 | 0.04357216 |
| Zfhx2 | 495.70819 | 454.176162 | 1.09144475 | 0.12623911 | 474.942176 | 0.12417966 | 0.04434915 | 2.80004587 | 0.00510953 | 0.04357216 |
| Cobl | 1100.66506 | 1163.68711 | 0.94584279 | -0.0803277 | 1132.17609 | -0.0800595 | 0.02859795 | -2.7994848 | 0.00511842 | 0.04362285 |
| Erap1 | 559.140768 | 603.448568 | 0.92657568 | -0.1100193 | 581.294668 | -0.1097353 | 0.03920263 | -2.7991836 | 0.0051232 | 0.04363848 |
| Sumo2 | 3010.619 | 2902.06678 | 1.03740514 | 0.05297942 | 2956.34289 | 0.05347307 | 0.01910476 | 2.79893947 | 0.00512707 | 0.0436464 |

| | | | | | | | | | | |
|---|---|---|---|---|---|---|---|---|---|---|
| Prpf38b | 612.971813 | 568.381477 | 1.07845142 | 0.1089612 | 590.676645 | 0.10926287 | 0.03904603 | 2.79830941 | 0.00513709 | 0.04368145 |
| Rpl37 | 809.334539 | 757.577137 | 1.06831965 | 0.09534337 | 783.455838 | 0.095076 | 0.03397529 | 2.79838696 | 0.00513585 | 0.04368145 |
| Fam122a | 133.500001 | 114.162693 | 1.16938377 | 0.22574848 | 123.831347 | 0.22507173 | 0.08044744 | 2.79774859 | 0.00514602 | 0.04370982 |
| C330027C09 | 942.428577 | 880.574202 | 1.07024323 | 0.09793871 | 911.501389 | 0.10543254 | 0.03768504 | 2.79772936 | 0.00514632 | 0.04370982 |
| Rpl36al | 617.752682 | 572.835046 | 1.07841286 | 0.10890961 | 595.293864 | 0.11203941 | 0.04007292 | 2.7958883 | 0.00517573 | 0.04393441 |
| Arf1 | 3825.86088 | 3683.5301 | 1.03863977 | 0.05469538 | 3754.69549 | 0.05574534 | 0.01994174 | 2.79541071 | 0.00518338 | 0.04394903 |
| Snta1 | 224.908967 | 251.306567 | 0.89495857 | -0.1601072 | 238.107767 | -0.1601 | 0.05726933 | -2.7955621 | 0.00518095 | 0.04394903 |
| Dgkd | 1151.64644 | 1217.24764 | 0.94610693 | -0.0799248 | 1184.44704 | -0.0801925 | 0.02869508 | -2.7946449 | 0.00519567 | 0.04402807 |
| Klhl11 | 205.133777 | 181.454479 | 1.13049718 | 0.17695739 | 193.294128 | 0.17692991 | 0.0633528 | 2.79277192 | 0.00522585 | 0.04425849 |
| Zfp930 | 219.65533 | 194.441809 | 1.1296713 | 0.17590305 | 207.048569 | 0.17550518 | 0.06288128 | 2.79105628 | 0.00525363 | 0.04444979 |
| Fam111a | 407.573796 | 371.921447 | 1.09585989 | 0.13206336 | 389.747622 | 0.13207451 | 0.04732459 | 2.79082197 | 0.00525744 | 0.04444979 |
| Chd3 | 1950.55486 | 1846.42822 | 1.05639355 | 0.07914739 | 1898.49154 | 0.07942224 | 0.0284566 | 2.79099499 | 0.00525463 | 0.04444979 |
| Zfp397 | 604.682198 | 561.506762 | 1.0768921 | 0.10687371 | 583.09448 | 0.1065859 | 0.03821196 | 2.78933309 | 0.00528167 | 0.04462922 |
| Nipal2 | 477.315755 | 438.420641 | 1.08871643 | 0.12262823 | 457.868198 | 0.1227198 | 0.04400159 | 2.78898566 | 0.00528734 | 0.04465167 |
| Ehf | 568.302272 | 527.73032 | 1.07688009 | 0.10685761 | 548.016296 | 0.10736799 | 0.03850798 | 2.78820114 | 0.00530016 | 0.04473446 |
| Reps1 | 801.560578 | 750.817881 | 1.06758323 | 0.09434855 | 776.189229 | 0.09416048 | 0.03377579 | 2.78780991 | 0.00530657 | 0.04476302 |
| Ptpra | 1010.1882 | 1067.99524 | 0.94587332 | -0.0802811 | 1039.09172 | -0.080169 | 0.02879005 | -2.784608 | 0.00535925 | 0.04518169 |
| Peli3 | 115.796269 | 137.159786 | 0.84424358 | -0.2442688 | 126.478027 | -0.2382018 | 0.08555025 | -2.784349 | 0.00536353 | 0.04519207 |
| Gcnt1 | 47.1933751 | 36.4555266 | 1.29454652 | 0.37244681 | 41.8244507 | 0.37730802 | 0.13561221 | 2.78225695 | 0.00539823 | 0.04540914 |
| Scin | 1224.43188 | 1157.16741 | 1.05812856 | 0.08151492 | 1190.79965 | 0.0813284 | 0.0292321 | 2.78216042 | 0.00539983 | 0.04540914 |
| Fam129b | 2559.97947 | 2660.79144 | 0.96211204 | -0.0557232 | 2610.38546 | -0.0553611 | 0.01989935 | -2.7820576 | 0.00540155 | 0.04540914 |
| Dcdc2a | 122.347228 | 142.754103 | 0.85704876 | -0.2225508 | 132.550665 | -0.2180386 | 0.07836485 | -2.7823523 | 0.00539664 | 0.04540914 |
| Lrrc42 | 568.074043 | 524.988922 | 1.08206863 | 0.113792 | 546.531482 | 0.11333125 | 0.0407409 | 2.78175624 | 0.00540656 | 0.04542555 |
| Tmem219 | 167.830657 | 193.079768 | 0.86922964 | -0.2021907 | 180.455212 | -0.2045845 | 0.07355957 | -2.7812091 | 0.00541568 | 0.04547641 |
| Prpf39 | 505.457745 | 467.07405 | 1.08217904 | 0.1139392 | 486.265898 | 0.11581644 | 0.04166516 | 2.77969481 | 0.005441 | 0.04566312 |
| LOC664787 | 2.71206783 | 6.30923519 | 0.42985683 | -1.2180719 | 4.51065141 | -1.2118985 | 0.43617669 | -2.7784577 | 0.00546176 | 0.04581142 |
| Nkain1 | 1051.90578 | 994.772554 | 1.05743346 | 0.08056688 | 1023.33917 | 0.08111778 | 0.02920669 | 2.77736945 | 0.00548008 | 0.04593911 |
| Wdfy1 | 298.342338 | 329.701249 | 0.90488689 | -0.1441906 | 314.021794 | -0.1440694 | 0.05188191 | -2.7768717 | 0.00548848 | 0.0459835 |
| Ubiad1 | 146.785058 | 168.669542 | 0.87025231 | -0.2004944 | 157.7273 | -0.2010379 | 0.07242851 | -2.7756741 | 0.00550874 | 0.04612714 |
| Elac2 | 271.462123 | 242.687096 | 1.11856843 | 0.16165352 | 257.074609 | 0.16160487 | 0.05827643 | 2.77307417 | 0.00555295 | 0.04647104 |
| Zfp970 | 157.799448 | 134.135083 | 1.17642189 | 0.23440553 | 145.967265 | 0.23483957 | 0.08472117 | 2.77191125 | 0.00557282 | 0.04657775 |
| Iars2 | 734.345753 | 782.727003 | 0.93818886 | -0.0920497 | 758.536377 | -0.0924899 | 0.03336811 | -2.7718042 | 0.00557465 | 0.04657775 |
| Syngr3 | 163.218407 | 185.698703 | 0.87894209 | -0.18616 | 174.458555 | -0.1865482 | 0.06730276 | -2.7717768 | 0.00557512 | 0.04657775 |
| Psmb9 | 161.332147 | 186.076188 | 0.86702199 | -0.2058595 | 173.704167 | -0.2058771 | 0.07429126 | -2.7712153 | 0.00558475 | 0.04663187 |
| Cdk13 | 658.2442 | 611.313472 | 1.07677032 | 0.10671054 | 634.778836 | 0.10608073 | 0.03830242 | 2.76955687 | 0.00561326 | 0.04684354 |
| Msl3l2 | 130.061023 | 150.406734 | 0.86472872 | -0.2096805 | 140.233878 | -0.210738 | 0.07610468 | -2.7690539 | 0.00562193 | 0.04688952 |
| Ptma | 34078.3579 | 34697.2909 | 0.98216244 | -0.0259664 | 34387.8334 | -0.0263803 | 0.00953158 | -2.7676765 | 0.00564575 | 0.04706164 |
| Xpr1 | 1725.45579 | 1801.84977 | 0.95760247 | -0.0625012 | 1763.65278 | -0.062468 | 0.02257682 | -2.7669101 | 0.00565904 | 0.04714591 |
| Hnrnpul1 | 2460.9908 | 2556.66041 | 0.96258024 | -0.0550213 | 2508.8256 | -0.0550074 | 0.01988425 | -2.7663799 | 0.00566825 | 0.04719611 |
| Ppa2 | 237.586421 | 211.854585 | 1.12145989 | 0.16537802 | 224.720503 | 0.16468425 | 0.05954062 | 2.76591415 | 0.00567635 | 0.04722944 |
| Rtraf | 941.12612 | 885.808181 | 1.06244912 | 0.08739375 | 913.46715 | 0.08606374 | 0.03111731 | 2.76578352 | 0.00567862 | 0.04722944 |
| Cbr1 | 217.897929 | 244.076579 | 0.89274411 | -0.1636814 | 230.987254 | -0.1635366 | 0.05914028 | -2.765232 | 0.00568823 | 0.04728284 |
| Arx | 6.53591915 | 2.86449408 | 2.28170105 | 1.19010978 | 4.70020652 | 1.27802959 | 0.46227383 | 2.7646592 | 0.00569823 | 0.04733938 |
| Ighmbp2 | 96.0491434 | 80.0763766 | 1.19946915 | 0.26239605 | 88.0627599 | 0.26297927 | 0.09517126 | 2.76322139 | 0.00572339 | 0.0475218 |
| Igf2 | 6777.71415 | 7060.45826 | 0.95995386 | -0.058963 | 6919.0862 | -0.063252 | 0.02289952 | -2.7621559 | 0.00574211 | 0.04765047 |
| Dopey1 | 452.415303 | 491.826751 | 0.91986721 | -0.1205025 | 472.121027 | -0.1220123 | 0.0441804 | -2.7616849 | 0.0057504 | 0.04769256 |
| Pir | 124.036277 | 106.03362 | 1.16978254 | 0.22624036 | 115.034949 | 0.22580057 | 0.0817675 | 2.76149533 | 0.00575373 | 0.04769356 |
| 5031425E22I | 113.566008 | 96.1481532 | 1.18115642 | 0.24020003 | 104.857081 | 0.24249737 | 0.08784729 | 2.76044228 | 0.00577232 | 0.04776744 |
| Ddx50 | 820.483523 | 770.848435 | 1.0643902 | 0.09002714 | 795.665979 | 0.09035917 | 0.03273209 | 2.76056849 | 0.00577009 | 0.04776744 |
| Ptprk | 695.641839 | 743.390804 | 0.93576869 | -0.0957761 | 719.516321 | -0.0947426 | 0.03431942 | -2.7606135 | 0.00576929 | 0.04776744 |
| Dennd3 | 296.36634 | 326.572247 | 0.9075062 | -0.1400206 | 311.469293 | -0.1399855 | 0.05072227 | -2.7598437 | 0.0057829 | 0.04782835 |
| Rpl18 | 1764.64082 | 1684.82162 | 1.04737546 | 0.06677871 | 1724.73122 | 0.0679522 | 0.02463485 | 2.75837708 | 0.00580891 | 0.04801669 |
| Grin3a | 182.13502 | 160.585041 | 1.13419668 | 0.18167083 | 171.360031 | 0.1813979 | 0.06578338 | 2.7575035 | 0.00582446 | 0.04811638 |
| Nectin1 | 606.893939 | 651.07749 | 0.9321378 | -0.1013848 | 628.985715 | -0.1010213 | 0.03663732 | -2.7573347 | 0.00582747 | 0.04811638 |
| Skp1a | 1998.99146 | 1911.30771 | 1.04587631 | 0.06471225 | 1955.14959 | 0.06410557 | 0.02327257 | 2.75455541 | 0.00587719 | 0.04847295 |
| Dtl | 793.439473 | 844.657583 | 0.93936228 | -0.0902464 | 819.048528 | -0.0899444 | 0.03265281 | -2.7545671 | 0.00587698 | 0.04847295 |
| C2cd4c | 39.3019917 | 50.6874202 | 0.7753796 | -0.3670253 | 44.9947059 | -0.3648403 | 0.13248213 | -2.7538831 | 0.00588928 | 0.04854561 |
| Slc38a5 | 200.848771 | 228.054119 | 0.88070662 | -0.1832666 | 214.451445 | -0.1839283 | 0.06681475 | -2.7528098 | 0.00590862 | 0.04867798 |
| Rbmxl1 | 813.069202 | 764.326785 | 1.0637717 | 0.08918856 | 788.697993 | 0.08922901 | 0.0324199 | 2.75229145 | 0.00591798 | 0.04872801 |
| Habp4 | 1146.74914 | 1215.86625 | 0.94315402 | -0.0844347 | 1181.30769 | -0.082808 | 0.03009075 | -2.7519427 | 0.00592429 | 0.04875161 |
| Degs1 | 627.552125 | 671.574786 | 0.93444861 | -0.0978128 | 649.563455 | -0.0979625 | 0.03559981 | -2.7517692 | 0.00592743 | 0.04875161 |
| Cd24a | 7998.5061 | 7794.78861 | 1.02613509 | 0.03722067 | 7896.64735 | 0.03731651 | 0.01356258 | 2.7514323 | 0.00593353 | 0.04877472 |
| Klhl22 | 360.142472 | 393.137888 | 0.91607165 | -0.1264677 | 376.64018 | -0.126349 | 0.04593162 | -2.750807 | 0.00594487 | 0.04881378 |
| Slc25a35 | 163.346022 | 189.459566 | 0.86216825 | -0.2139587 | 176.402794 | -0.2061632 | 0.07494366 | -2.7509096 | 0.005943 | 0.04881378 |
| Cmip | 1496.91718 | 1571.18149 | 0.95273346 | -0.0698554 | 1534.04933 | -0.0702098 | 0.0255307 | -2.7500161 | 0.00595923 | 0.04890465 |
| Ppm1g | 1304.09751 | 1368.97626 | 0.95260783 | -0.0700457 | 1336.53689 | -0.0698893 | 0.02542477 | -2.7488668 | 0.00598017 | 0.04904929 |
| Bckdhb | 96.7866843 | 81.1847548 | 1.19217807 | 0.25359974 | 88.9857194 | 0.25313746 | 0.0922073 | 2.7453081 | 0.00604541 | 0.0495322 |

| | | | | | | | | | | |
|---|---|---|---|---|---|---|---|---|---|---|
| D930020B18 | 113.74726 | 132.471167 | 0.85865673 | -0.2198466 | 123.109214 | -0.2204705 | 0.0803086 | -2.7452908 | 0.00604573 | 0.0495322 |
| Xlr3a | 4.39827468 | 1.55359396 | 2.8310323 | 1.50132821 | 2.97593422 | 1.55052905 | 0.56492571 | 2.74466008 | 0.00605736 | 0.04960008 |
| Ap2a1 | 1174.69949 | 1237.02389 | 0.94961746 | -0.0745816 | 1205.86169 | -0.0746239 | 0.02719084 | -2.7444497 | 0.00606124 | 0.04960447 |
| Cer1 | 677.038364 | 632.896489 | 1.06974581 | 0.09726802 | 654.967427 | 0.09810149 | 0.0357604 | 2.74329998 | 0.00608251 | 0.04972359 |
| Kras | 1411.36181 | 1342.50524 | 1.05128962 | 0.07216017 | 1376.93353 | 0.07187958 | 0.02620046 | 2.74344735 | 0.00607978 | 0.04972359 |
| Tmed9 | 1029.8031 | 972.253572 | 1.05919189 | 0.08296399 | 1001.02834 | 0.08431331 | 0.03073694 | 2.74306171 | 0.00608692 | 0.04973225 |
| Klf2 | 121.716896 | 104.173752 | 1.16840273 | 0.22453764 | 112.945324 | 0.22495681 | 0.08202908 | 2.74240311 | 0.00609914 | 0.04977721 |
| Ube2f | 222.435354 | 198.409662 | 1.12109134 | 0.16490382 | 210.422508 | 0.16470503 | 0.06005601 | 2.74252373 | 0.0060969 | 0.04977721 |
| Lrrc4c | 41.2528942 | 52.426059 | 0.78687765 | -0.3457888 | 46.8394765 | -0.3455203 | 0.12602056 | -2.7417772 | 0.00611078 | 0.04984257 |
| Fam135b | 26.7955587 | 36.2862958 | 0.73844844 | -0.4374309 | 31.5409271 | -0.4352282 | 0.15874911 | -2.7416105 | 0.00611388 | 0.04984257 |
| Pdcd5 | 267.910722 | 241.114589 | 1.11113443 | 0.15203338 | 254.512655 | 0.15220945 | 0.05554071 | 2.7405024 | 0.00613453 | 0.04992887 |
| Gprin1 | 1293.01455 | 1359.14819 | 0.95134184 | -0.0719643 | 1326.08137 | -0.0717363 | 0.0261779 | -2.7403383 | 0.0061376 | 0.04992887 |
| Rab3d | 879.968154 | 954.353718 | 0.92205661 | -0.1170728 | 917.160936 | -0.1131148 | 0.04127794 | -2.74032 | 0.00613794 | 0.04992887 |
| Arc | 211.995652 | 237.590708 | 0.89227249 | -0.1644437 | 224.79318 | -0.1627583 | 0.05938805 | -2.7405901 | 0.0061329 | 0.04992887 |
| Kansl3 | 1284.0213 | 1353.60479 | 0.94859394 | -0.0761374 | 1318.81304 | -0.0765378 | 0.02793398 | -2.7399537 | 0.00614478 | 0.04995713 |
