## Supplementary Data 2 for "FGFR1 is critical for *Rbl2* loss-driven tumor development and requires PLCG1 activation for continued growth of small cell lung cancer": Supplementaty Data 2.pdf

| Term | Count | % | PValue | Genes | List Total | Pop Hits | Pop Total | Fold Enrichm | Bonferroni | Benjamini | FDR |
|---|---|---|---|---|---|---|---|---|---|---|---|
| GO:0006412~translation | 88 | 0.4197 | 1.40E-20 | RPL18, RPL1 | 1286 | 319 | 13588 | 2.91478522 | 4.48E-17 | 4.48E-17 | 2.54E-17 |
| GO:0030182~neuron differentiation | 77 | 0.3672 | 1.84E-09 | GPRIN1, EDN | 1286 | 399 | 13588 | 2.03907124 | 5.87E-06 | 2.94E-06 | 3.33E-06 |
| GO:0007155~cell adhesion | 96 | 0.4578 | 1.03E-08 | CLDN7, TLN1 | 1286 | 561 | 13588 | 1.80810206 | 3.30E-05 | 1.10E-05 | 1.87E-05 |
| GO:0022610~biological adhesion | 96 | 0.4578 | 1.11E-08 | CLDN7, TLN1 | 1286 | 562 | 13588 | 1.8048848 | 3.56E-05 | 8.89E-06 | 2.02E-05 |
| GO:0031175~neuron projection development | 49 | 0.2337 | 1.88E-08 | GPRIN1, NRT | 1286 | 218 | 13588 | 2.37494828 | 6.00E-05 | 1.20E-05 | 3.40E-05 |
| GO:0006928~cell motion | 69 | 0.3291 | 4.15E-08 | EDN3, NRTN, | 1286 | 367 | 13588 | 1.98654129 | 1.33E-04 | 2.21E-05 | 7.52E-05 |
| GO:0007049~cell cycle | 100 | 0.4769 | 4.97E-08 | SEPT5, SEPT4 | 1286 | 611 | 13588 | 1.72931202 | 1.59E-04 | 2.27E-05 | 9.00E-05 |
| GO:0000904~cell morphogenesis involved in differentiation | 47 | 0.2242 | 5.92E-08 | DCC, ABLIM1 | 1286 | 212 | 13588 | 2.34248364 | 1.89E-04 | 2.36E-05 | 1.07E-04 |
| GO:0030030~cell projection organization | 62 | 0.2957 | 6.46E-08 | GPRIN1, NRT | 1286 | 319 | 13588 | 2.05359868 | 2.06E-04 | 2.29E-05 | 1.17E-04 |
| GO:0048666~neuron development | 58 | 0.2766 | 8.39E-08 | GPRIN1, NRT | 1286 | 292 | 13588 | 2.09874518 | 2.68E-04 | 2.68E-05 | 1.52E-04 |
| GO:0000902~cell morphogenesis | 60 | 0.2862 | 1.12E-07 | NRP1, UCHL1 | 1286 | 309 | 13588 | 2.05166921 | 3.58E-04 | 3.25E-05 | 2.03E-04 |
| GO:0048667~cell morphogenesis involved in neuron differentiatior | 41 | 0.1955 | 2.99E-07 | ABLIM1, DCC | 1286 | 182 | 13588 | 2.38027447 | 9.55E-04 | 7.97E-05 | 5.42E-04 |
| GO:0051301~cell division | 55 | 0.2623 | 3.11E-07 | SEPT5, SEPT4 | 1286 | 281 | 13588 | 2.06809716 | 9.94E-04 | 7.65E-05 | 5.64E-04 |
| GO:0007409~axonogenesis | 38 | 0.1812 | 3.54E-07 | ABLIM1, DCC | 1286 | 163 | 13588 | 2.46326174 | 0.00113001 | 8.08E-05 | 6.42E-04 |
| GO:0032989~cellular component morphogenesis | 64 | 0.3052 | 4.24E-07 | NRP1, UCHL1 | 1286 | 351 | 13588 | 1.92658168 | 0.00135534 | 9.04E-05 | 7.70E-04 |
| GO:0032990~cell part morphogenesis | 45 | 0.2146 | 4.32E-07 | ABLIM1, DCC | 1286 | 212 | 13588 | 2.24280349 | 0.00137824 | 8.62E-05 | 7.83E-04 |
| GO:0060429~epithelium development | 53 | 0.2528 | 5.39E-07 | NRP1, NPNT, | 1286 | 271 | 13588 | 2.06643214 | 0.00172207 | 1.01E-04 | 9.78E-04 |
| GO:0048858~cell projection morphogenesis | 43 | 0.2051 | 7.47E-07 | ABLIM1, DCC | 1286 | 202 | 13588 | 2.24921855 | 0.00238407 | 1.33E-04 | 0.00135429 |
| GO:0048812~neuron projection morphogenesis | 39 | 0.186 | 9.40E-07 | ABLIM1, DCC | 1286 | 176 | 13588 | 2.34135091 | 0.00300074 | 1.58E-04 | 0.00170512 |
| GO:0016477~cell migration | 48 | 0.2289 | 1.03E-06 | CER1, DCC, E | 1286 | 240 | 13588 | 2.11321928 | 0.00329321 | 1.65E-04 | 0.00187159 |
| GO:0006796~phosphate metabolic process | 125 | 0.5961 | 1.23E-06 | ILKAP, RNASI | 1286 | 866 | 13588 | 1.52512939 | 0.00392922 | 1.87E-04 | 0.00223375 |
| GO:0006793~phosphorus metabolic process | 125 | 0.5961 | 1.23E-06 | ILKAP, RNASI | 1286 | 866 | 13588 | 1.52512939 | 0.00392922 | 1.87E-04 | 0.00223375 |
| GO:0043549~regulation of kinase activity | 41 | 0.1955 | 1.29E-06 | HMGCR, CSF | 1286 | 192 | 13588 | 2.25630184 | 0.00412127 | 1.88E-04 | 0.00234315 |
| GO:0035295~tube development | 51 | 0.2432 | 1.33E-06 | GNA13, NRP | 1286 | 264 | 13588 | 2.04117772 | 0.00423183 | 1.84E-04 | 0.00240614 |
| GO:0042127~regulation of cell proliferation | 86 | 0.4101 | 1.33E-06 | EDN3, E2F3, | 1286 | 538 | 13588 | 1.68900426 | 0.00423833 | 1.77E-04 | 0.00240985 |
| GO:0045859~regulation of protein kinase activity | 40 | 0.1908 | 1.48E-06 | HMGCR, CSF | 1286 | 186 | 13588 | 2.2722788 | 0.00470968 | 1.89E-04 | 0.00267848 |
| GO:0022402~cell cycle process | 67 | 0.3195 | 2.58E-06 | AURKB, PTTG | 1286 | 393 | 13588 | 1.80134468 | 0.00820007 | 3.17E-04 | 0.00467167 |
| GO:0007411~axon guidance | 26 | 0.124 | 3.02E-06 | ABLIM1, DCC | 1286 | 98 | 13588 | 2.80325007 | 0.00959922 | 3.57E-04 | 0.00547262 |
| GO:0030879~mammary gland development | 24 | 0.1145 | 3.12E-06 | WNT5A, IRS2 | 1286 | 86 | 13588 | 2.94867807 | 0.0099324 | 3.56E-04 | 0.00566351 |
| GO:0051338~regulation of transferase activity | 41 | 0.1955 | 3.32E-06 | HMGCR, CSF | 1286 | 199 | 13588 | 2.17693444 | 0.01054288 | 3.65E-04 | 0.00601345 |
| GO:0000278~mitotic cell cycle | 47 | 0.2242 | 3.97E-06 | PPP2R3A, CE | 1286 | 244 | 13588 | 2.03527267 | 0.01259796 | 4.23E-04 | 0.00719304 |
| GO:0006470~protein amino acid dephosphorylation | 28 | 0.1335 | 5.68E-06 | ILKAP, PPP2F | 1286 | 114 | 13588 | 2.59518158 | 0.01798533 | 5.85E-04 | 0.01029697 |
| GO:0032535~regulation of cellular component size | 35 | 0.1669 | 5.80E-06 | NRP1, CAPZA | 1286 | 161 | 13588 | 2.29697748 | 0.01836341 | 5.79E-04 | 0.01051543 |
| GO:0022403~cell cycle phase | 57 | 0.2718 | 8.84E-06 | PTTG1, AURK | 1286 | 328 | 13588 | 1.83618139 | 0.02786448 | 8.56E-04 | 0.01603306 |
| GO:0048070~regulation of pigmentation during development | 8 | 0.0382 | 1.56E-05 | EDN3, ADAM | 1286 | 11 | 13588 | 7.68443376 | 0.04866107 | 0.00146613 | 0.02829995 |
| GO:0042325~regulation of phosphorylation | 51 | 0.2432 | 2.03E-05 | HMGCR, TGF | 1286 | 290 | 13588 | 1.85817558 | 0.06296532 | 0.00185642 | 0.03689309 |
| GO:0045664~regulation of neuron differentiation | 25 | 0.1192 | 2.08E-05 | NRP1, SOX2, | 1286 | 102 | 13588 | 2.58972952 | 0.06429402 | 0.00184424 | 0.03769791 |
| GO:0048870~cell motility | 50 | 0.2385 | 2.41E-05 | DCC, CER1, E | 1286 | 284 | 13588 | 1.86022824 | 0.07407769 | 0.00207797 | 0.04365925 |
| GO:0051674~localization of cell | 50 | 0.2385 | 2.41E-05 | DCC, CER1, E | 1286 | 284 | 13588 | 1.86022824 | 0.07407769 | 0.00207797 | 0.04365925 |
| GO:0048729~tissue morphogenesis | 44 | 0.2098 | 2.41E-05 | WNT5A, COB | 1286 | 238 | 13588 | 1.95339598 | 0.07421344 | 0.00202719 | 0.0437424 |
| GO:0008285~negative regulation of cell proliferation | 42 | 0.2003 | 2.71E-05 | CER1, WNT5 | 1286 | 224 | 13588 | 1.98114308 | 0.08285166 | 0.00221513 | 0.04905888 |
| GO:0002009~morphogenesis of an epithelium | 35 | 0.1669 | 2.89E-05 | WNT5A, COB | 1286 | 173 | 13588 | 2.13764957 | 0.08812292 | 0.00230359 | 0.05232765 |
| GO:0060284~regulation of cell development | 33 | 0.1574 | 3.01E-05 | NRP1, SOX2, | 1286 | 159 | 13588 | 2.19296341 | 0.09160959 | 0.0023407 | 0.05450011 |
| GO:0050767~regulation of neurogenesis | 29 | 0.1383 | 3.46E-05 | NRP1, SOX2, | 1286 | 132 | 13588 | 2.32133937 | 0.10466291 | 0.0026288 | 0.06270762 |
| GO:0048732~gland development | 38 | 0.1812 | 3.70E-05 | WNT5A, FGF | 1286 | 197 | 13588 | 2.03813027 | 0.11157055 | 0.00274738 | 0.06709919 |
| GO:0048754~branching morphogenesis of a tube | 23 | 0.1097 | 4.15E-05 | GNA13, WNT | 1286 | 93 | 13588 | 2.61312062 | 0.12434393 | 0.00301322 | 0.07531013 |
| GO:0016337~cell-cell adhesion | 43 | 0.2051 | 4.31E-05 | CLDN7, THRA | 1286 | 236 | 13588 | 1.92517859 | 0.12865434 | 0.00305569 | 0.07810782 |
| GO:0008104~protein localization | 105 | 0.5008 | 4.31E-05 | SLC9A2, GRIK | 1286 | 753 | 13588 | 1.47336006 | 0.12880226 | 0.00299304 | 0.07820408 |
| GO:0016311~dephosphorylation | 30 | 0.1431 | 4.61E-05 | ILKAP, PPP2F | 1286 | 141 | 13588 | 2.24810562 | 0.13695989 | 0.00312901 | 0.08353761 |
| GO:0040007~growth | 37 | 0.1765 | 5.45E-05 | WNT5A, FKB | 1286 | 193 | 13588 | 2.0256247 | 0.15986489 | 0.00362243 | 0.09878548 |
| GO:0051174~regulation of phosphorus metabolic process | 51 | 0.2432 | 5.55E-05 | HMGCR, TGF | 1286 | 301 | 13588 | 1.79026883 | 0.16245387 | 0.0036114 | 0.1005349 |
| GO:0019220~regulation of phosphate metabolic process | 51 | 0.2432 | 5.55E-05 | HMGCR, TGF | 1286 | 301 | 13588 | 1.79026883 | 0.16245387 | 0.0036114 | 0.1005349 |
| GO:0008361~regulation of cell size | 25 | 0.1192 | 5.63E-05 | NRP1, TGFB3 | 1286 | 108 | 13588 | 2.44585565 | 0.16454908 | 0.00358922 | 0.10195462 |
| GO:0000087~M phase of mitotic cell cycle | 37 | 0.1765 | 6.10E-05 | CETN3, CEP5 | 1286 | 194 | 13588 | 2.01518334 | 0.17703579 | 0.00381315 | 0.11048974 |
| GO:0044093~positive regulation of molecular function | 51 | 0.2432 | 8.37E-05 | CASR, ADCY2 | 1286 | 306 | 13588 | 1.76101607 | 0.2347308 | 0.00513155 | 0.15167611 |
| GO:0007067~mitosis | 36 | 0.1717 | 8.95E-05 | CETN3, CEP5 | 1286 | 190 | 13588 | 2.00199722 | 0.24866914 | 0.00537999 | 0.16208917 |
| GO:0000280~nuclear division | 36 | 0.1717 | 8.95E-05 | CETN3, CEP5 | 1286 | 190 | 13588 | 2.00199722 | 0.24866914 | 0.00537999 | 0.16208917 |
| GO:0051960~regulation of nervous system development | 30 | 0.1431 | 1.15E-04 | NRP1, SOX2, | 1286 | 148 | 13588 | 2.1417763 | 0.30727373 | 0.00677547 | 0.20808193 |
| GO:0007010~cytoskeleton organization | 53 | 0.2528 | 1.19E-04 | TLN1, BCAR1 | 1286 | 326 | 13588 | 1.71780095 | 0.31577075 | 0.00687557 | 0.21506973 |
| GO:0035239~tube morphogenesis | 33 | 0.1574 | 1.30E-04 | GNA13, WNT | 1286 | 171 | 13588 | 2.03907124 | 0.33993632 | 0.00739075 | 0.23542509 |
| GO:0009967~positive regulation of signal transduction | 33 | 0.1574 | 1.46E-04 | FGFR1, FKBP | 1286 | 172 | 13588 | 2.02721617 | 0.37193031 | 0.00812652 | 0.2635454 |

| | | | | | | | | | | | |
|---|---|---|---|---|---|---|---|---|---|---|---|
| GO:0007242~intracellular signaling cascade | 120 | 0.5723 | 1.70E-04 | GNA13, CASF | 1286 | 915 | 13588 | 1.38571756 | 0.41965799 | 0.00933781 | 0.30825967 |
| GO:0010647~positive regulation of cell communication | 35 | 0.1669 | 1.80E-04 | FGFR1, FKBP | 1286 | 189 | 13588 | 1.95668452 | 0.43801507 | 0.00971992 | 0.32643901 |
| GO:0000279~M phase | 47 | 0.2242 | 1.83E-04 | CETN3, CEP5 | 1286 | 283 | 13588 | 1.7547934 | 0.44248289 | 0.00969044 | 0.3309529 |
| GO:0048285~organelle fission | 36 | 0.1717 | 1.88E-04 | CETN3, CEP5 | 1286 | 197 | 13588 | 1.93086026 | 0.45120281 | 0.00978828 | 0.3398673 |
| GO:0007156~homophilic cell adhesion | 25 | 0.1192 | 2.11E-04 | PTPRF, CLSTN | 1286 | 117 | 13588 | 2.25771291 | 0.49014648 | 0.01080622 | 0.38147935 |
| GO:0033043~regulation of organelle organization | 30 | 0.1431 | 2.36E-04 | RAB3A, CAPZ | 1286 | 154 | 13588 | 2.05833047 | 0.52940938 | 0.01189326 | 0.426763 |
| GO:0001763~morphogenesis of a branching structure | 26 | 0.124 | 2.38E-04 | WNT5A, GNA | 1286 | 125 | 13588 | 2.19774806 | 0.53262028 | 0.01181424 | 0.43063095 |
| GO:0007267~cell-cell signaling | 47 | 0.2242 | 3.23E-04 | WNT5A, SYT1 | 1286 | 290 | 13588 | 1.71243632 | 0.64353442 | 0.01574424 | 0.58355864 |
| GO:0007017~microtubule-based process | 37 | 0.1765 | 3.47E-04 | KIF23, UCHL1 | 1286 | 211 | 13588 | 1.8528226 | 0.66973801 | 0.0166458 | 0.62661721 |
| GO:0006897~endocytosis | 34 | 0.1622 | 3.55E-04 | STON2, ABCA | 1286 | 188 | 13588 | 1.91088978 | 0.67870287 | 0.01680333 | 0.64213255 |
| GO:0010324~membrane invagination | 34 | 0.1622 | 3.55E-04 | STON2, ABCA | 1286 | 188 | 13588 | 1.91088978 | 0.67870287 | 0.01680333 | 0.64213255 |
| GO:0051270~regulation of cell motion | 23 | 0.1097 | 3.68E-04 | COL18A1, GN | 1286 | 107 | 13588 | 2.27121699 | 0.69123763 | 0.01713363 | 0.66456377 |
| GO:0010959~regulation of metal ion transport | 14 | 0.0668 | 3.75E-04 | ATP2B2, CAC | 1286 | 48 | 13588 | 3.08177812 | 0.69882815 | 0.01724201 | 0.67859167 |
| GO:0043523~regulation of neuron apoptosis | 19 | 0.0906 | 3.92E-04 | SNCB, SNCA, | 1286 | 80 | 13588 | 2.5094479 | 0.71479189 | 0.0177623 | 0.70927813 |
| GO:0033674~positive regulation of kinase activity | 26 | 0.124 | 4.45E-04 | CSF1, TGFB3, | 1286 | 130 | 13588 | 2.11321928 | 0.75900244 | 0.01984231 | 0.80412102 |
| GO:0043067~regulation of programmed cell death | 78 | 0.372 | 4.74E-04 | SNCB, SNCA, | 1286 | 560 | 13588 | 1.47170629 | 0.7802867 | 0.02082771 | 0.85614783 |
| GO:0030029~actin filament-based process | 32 | 0.1526 | 4.97E-04 | TLN1, CNN3, | 1286 | 176 | 13588 | 1.92110844 | 0.79602941 | 0.02154238 | 0.89796166 |
| GO:0045860~positive regulation of protein kinase activity | 25 | 0.1192 | 5.21E-04 | CSF1, MET, T | 1286 | 124 | 13588 | 2.13026138 | 0.81072132 | 0.02224262 | 0.93998698 |
| GO:0060541~respiratory system development | 25 | 0.1192 | 5.21E-04 | WNT5A, RAB | 1286 | 124 | 13588 | 2.13026138 | 0.81072132 | 0.02224262 | 0.93998698 |
| GO:0010941~regulation of cell death | 78 | 0.372 | 5.64E-04 | SNCB, SNCA, | 1286 | 563 | 13588 | 1.46386416 | 0.83544985 | 0.02377338 | 1.01864586 |
| GO:0007268~synaptic transmission | 32 | 0.1526 | 6.07E-04 | SYT1, RAB3A | 1286 | 178 | 13588 | 1.89952295 | 0.85620927 | 0.02519553 | 1.094353 |
| GO:0016310~phosphorylation | 95 | 0.4531 | 6.51E-04 | RNASEL, SNC | 1286 | 718 | 13588 | 1.39802111 | 0.87534956 | 0.02667974 | 1.17448213 |
| GO:0007215~glutamate signaling pathway | 7 | 0.0334 | 6.76E-04 | APP, GNAQ, | 1286 | 13 | 13588 | 5.68943654 | 0.88485082 | 0.02733143 | 1.21892882 |
| GO:0043405~regulation of MAP kinase activity | 19 | 0.0906 | 7.31E-04 | HMGCR, MET | 1286 | 84 | 13588 | 2.38995038 | 0.90332683 | 0.02914187 | 1.31690214 |
| GO:0030036~actin cytoskeleton organization | 30 | 0.1431 | 7.70E-04 | TLN1, CNN3, | 1286 | 165 | 13588 | 1.92110844 | 0.91482995 | 0.03031967 | 1.3878099 |
| GO:0051347~positive regulation of transferase activity | 26 | 0.124 | 7.96E-04 | CSF1, TGFB3, | 1286 | 135 | 13588 | 2.0349519 | 0.92158066 | 0.03093947 | 1.43400313 |
| GO:0030334~regulation of cell migration | 20 | 0.0954 | 8.51E-04 | COL18A1, GN | 1286 | 92 | 13588 | 2.29697748 | 0.93408868 | 0.03262007 | 1.53113054 |
| GO:0010810~regulation of cell-substrate adhesion | 12 | 0.0572 | 9.09E-04 | DDR1, FBLN2 | 1286 | 40 | 13588 | 3.16982893 | 0.94537078 | 0.03442002 | 1.63596538 |
| GO:0045597~positive regulation of cell differentiation | 31 | 0.1478 | 9.61E-04 | HMGB1, THR | 1286 | 175 | 13588 | 1.87170851 | 0.95366999 | 0.03591038 | 1.72788612 |
| GO:0051129~negative regulation of cellular component organizatio | 20 | 0.0954 | 9.76E-04 | NRP1, CAPZA | 1286 | 93 | 13588 | 2.2722788 | 0.95589012 | 0.03605254 | 1.75526297 |
| GO:0051130~positive regulation of cellular component organizatio | 24 | 0.1145 | 9.84E-04 | CALY, ACTN4 | 1286 | 122 | 13588 | 2.07857635 | 0.9569975 | 0.03592595 | 1.76943475 |
| GO:0045634~regulation of melanocyte differentiation | 5 | 0.0238 | 0.00102148 | ADAMTS9, G | 1286 | 6 | 13588 | 8.80508035 | 0.96185363 | 0.03684792 | 1.83619749 |
| GO:0050932~regulation of pigment cell differentiation | 5 | 0.0238 | 0.00102148 | ADAMTS9, G | 1286 | 6 | 13588 | 8.80508035 | 0.96185363 | 0.03684792 | 1.83619749 |
| GO:0010627~regulation of protein kinase cascade | 28 | 0.1335 | 0.00131969 | FGFR1, LITAF | 1286 | 155 | 13588 | 1.90871419 | 0.98530898 | 0.04682851 | 2.36621624 |
| GO:0040012~regulation of locomotion | 22 | 0.1049 | 0.00135828 | COL18A1, GN | 1286 | 110 | 13588 | 2.11321928 | 0.98701563 | 0.04763707 | 2.43460068 |
| GO:0043069~negative regulation of programmed cell death | 39 | 0.186 | 0.00145092 | FGFR1, SNCB | 1286 | 244 | 13588 | 1.68884328 | 0.9903472 | 0.05025447 | 2.59859613 |
| GO:0043085~positive regulation of catalytic activity | 41 | 0.1955 | 0.00150537 | CASR, ADCY2 | 1286 | 261 | 13588 | 1.65980825 | 0.99189102 | 0.05153435 | 2.6948604 |
| GO:0030324~lung development | 22 | 0.1049 | 0.00152933 | CEBPA, WNT | 1286 | 111 | 13588 | 2.09418127 | 0.99248975 | 0.05177962 | 2.7371988 |
| GO:0060562~epithelial tube morphogenesis | 22 | 0.1049 | 0.00152933 | WNT5A, KAT | 1286 | 111 | 13588 | 2.09418127 | 0.99248975 | 0.05177962 | 2.7371988 |
| GO:0060688~regulation of morphogenesis of a branching structure | 9 | 0.0429 | 0.0015605 | WNT5A, FGF | 1286 | 25 | 13588 | 3.80379471 | 0.99320289 | 0.05225465 | 2.79224225 |
| GO:0060548~negative regulation of cell death | 39 | 0.186 | 0.00156497 | FGFR1, SNCB | 1286 | 245 | 13588 | 1.68195004 | 0.99329945 | 0.05185772 | 2.80013363 |
| GO:0045184~establishment of protein localization | 86 | 0.4101 | 0.00164664 | COPA, AP1B1 | 1286 | 656 | 13588 | 1.38518947 | 0.99484085 | 0.05393304 | 2.94419789 |
| GO:0006468~protein amino acid phosphorylation | 84 | 0.4006 | 0.00173132 | RNASEL, TGF | 1286 | 640 | 13588 | 1.38680016 | 0.99606595 | 0.05605596 | 3.09337388 |
| GO:0051726~regulation of cell cycle | 35 | 0.1669 | 0.00178951 | HMGB1, ILKA | 1286 | 214 | 13588 | 1.72809988 | 0.99673471 | 0.0573069 | 3.19576553 |
| GO:0000122~negative regulation of transcription from RNA polyme | 37 | 0.1765 | 0.00188404 | GLIS3, THRB, | 1286 | 231 | 13588 | 1.69240505 | 0.99758747 | 0.05964781 | 3.36186637 |
| GO:0015031~protein transport | 85 | 0.4054 | 0.00189297 | COPA, AP1B1 | 1286 | 651 | 13588 | 1.37959784 | 0.99765546 | 0.05933511 | 3.37753858 |
| GO:0030323~respiratory tube development | 22 | 0.1049 | 0.001927 | CEBPA, WNT | 1286 | 113 | 13588 | 2.05711612 | 0.99789754 | 0.05978476 | 3.43726604 |
| GO:0001764~neuron migration | 16 | 0.0763 | 0.00196519 | DCC, SATB2, | 1286 | 70 | 13588 | 2.41510775 | 0.99813954 | 0.06034912 | 3.50424275 |
| GO:0042981~regulation of apoptosis | 74 | 0.3529 | 0.0020355 | SNCB, SNCA, | 1286 | 553 | 13588 | 1.41390802 | 0.99851463 | 0.06184869 | 3.62743839 |
| GO:0051493~regulation of cytoskeleton organization | 20 | 0.0954 | 0.00211208 | PSRC1, CAPZ | 1286 | 99 | 13588 | 2.13456493 | 0.9988377 | 0.06349963 | 3.76145461 |
| GO:0021952~central nervous system projection neuron axonogene | 6 | 0.0286 | 0.00213085 | DCC, EPHB3, | 1286 | 11 | 13588 | 5.76332532 | 0.99890549 | 0.06345004 | 3.79426567 |
| GO:0008284~positive regulation of cell proliferation | 43 | 0.2051 | 0.00233015 | EDN3, HMGB | 1286 | 284 | 13588 | 1.59979629 | 0.99942194 | 0.06854554 | 4.14210321 |
| GO:0019226~transmission of nerve impulse | 36 | 0.1717 | 0.00240624 | SYT1, RAB3A | 1286 | 226 | 13588 | 1.68309501 | 0.99954699 | 0.07006235 | 4.27459009 |
| GO:0001655~urogenital system development | 26 | 0.124 | 0.00248751 | WNT5A, FGF | 1286 | 146 | 13588 | 1.88163361 | 0.99965084 | 0.07169257 | 4.41590518 |
| GO:0001667~ameboidal cell migration | 12 | 0.0572 | 0.00257365 | CER1, EDN3, | 1286 | 45 | 13588 | 2.81762571 | 0.99973505 | 0.07342384 | 4.56546999 |
| GO:0021537~telencephalon development | 16 | 0.0763 | 0.00263175 | KAT2A, SOX2 | 1286 | 72 | 13588 | 2.34802143 | 0.99978006 | 0.07435776 | 4.66622892 |
| GO:0000910~cytokinesis | 9 | 0.0429 | 0.00270056 | FMN2, PRC1, | 1286 | 27 | 13588 | 3.52203214 | 0.99982358 | 0.07556236 | 4.78543058 |
| GO:0016358~dendrite development | 11 | 0.0525 | 0.00273212 | BBS1, BDNF, | 1286 | 39 | 13588 | 2.98018104 | 0.99984056 | 0.07575041 | 4.8400604 |
| GO:0035264~multicellular organism growth | 11 | 0.0525 | 0.00273212 | BBS1, KAT2A | 1286 | 39 | 13588 | 2.98018104 | 0.99984056 | 0.07575041 | 4.8400604 |
| GO:0030705~cytoskeleton-dependent intracellular transport | 10 | 0.0477 | 0.00279655 | DYNC1I1, AP | 1286 | 33 | 13588 | 3.2018474 | 0.9998703 | 0.07680355 | 4.95147065 |
| GO:0051247~positive regulation of protein metabolic process | 21 | 0.1002 | 0.00284941 | HMGB1, TGF | 1286 | 109 | 13588 | 2.03566995 | 0.99989052 | 0.07753472 | 5.04280661 |
| GO:0034599~cellular response to oxidative stress | 8 | 0.0382 | 0.00317565 | GPX1, EPAS1 | 1286 | 22 | 13588 | 3.84221688 | 0.99996153 | 0.08531114 | 5.60458971 |

| | | | | | | | | | | | |
|---|---|---|---|---|---|---|---|---|---|---|---|
| GO:0051094~positive regulation of developmental process | 34 | 0.1622 | 0.003373 | FGFR1, HMG | 1286 | 214 | 13588 | 1.6787256 | 0.99997957 | 0.08962476 | 5.94289358 |
| GO:0043066~negative regulation of apoptosis | 37 | 0.1765 | 0.00338637 | FGFR1, SNCB | 1286 | 239 | 13588 | 1.63575551 | 0.99998042 | 0.08922428 | 5.96577454 |
| GO:0051924~regulation of calcium ion transport | 10 | 0.0477 | 0.00348964 | ATP2B2, CAC | 1286 | 34 | 13588 | 3.10767542 | 0.99998594 | 0.09107293 | 6.14230638 |
| GO:0010720~positive regulation of cell development | 12 | 0.0572 | 0.00370709 | ASCL1, PLXN | 1286 | 47 | 13588 | 2.69772675 | 0.999993 | 0.09569829 | 6.51301668 |
| GO:0051146~striated muscle cell differentiation | 18 | 0.0858 | 0.00371892 | RXRA, MET, | 1286 | 89 | 13588 | 2.13696332 | 0.99999326 | 0.09522211 | 6.53314308 |
| GO:0051272~positive regulation of cell motion | 11 | 0.0525 | 0.00405002 | COL18A1, HM | 1286 | 41 | 13588 | 2.83480636 | 0.99999767 | 0.10244821 | 7.0948078 |
| GO:0031346~positive regulation of cell projection organization | 8 | 0.0382 | 0.00419772 | PLXNB2, TGF | 1286 | 23 | 13588 | 3.67516397 | 0.99999855 | 0.10515844 | 7.34433504 |
| GO:0010604~positive regulation of macromolecule metabolic proc | 81 | 0.3863 | 0.00420312 | E2F3, MEF2A | 1286 | 633 | 13588 | 1.35205973 | 0.99999858 | 0.10447035 | 7.35344649 |
| GO:0043269~regulation of ion transport | 14 | 0.0668 | 0.00466196 | ATP2B2, CAC | 1286 | 62 | 13588 | 2.38589274 | 0.99999967 | 0.11433675 | 8.12452128 |
| GO:0010970~microtubule-based transport | 7 | 0.0334 | 0.00483788 | DYNC1I1, AP | 1286 | 18 | 13588 | 4.1090375 | 0.99999981 | 0.11749877 | 8.41853518 |
| GO:0021953~central nervous system neuron differentiation | 11 | 0.0525 | 0.00487439 | DCC, ARX, AS | 1286 | 42 | 13588 | 2.76731097 | 0.99999983 | 0.11744403 | 8.47943583 |
| GO:0048762~mesenchymal cell differentiation | 12 | 0.0572 | 0.00520765 | ALDH1A2, FG | 1286 | 49 | 13588 | 2.58761545 | 0.99999994 | 0.12404245 | 9.03363935 |
| GO:0001558~regulation of cell growth | 18 | 0.0858 | 0.00528338 | NRP1, PSRC1 | 1286 | 92 | 13588 | 2.06727973 | 0.99999996 | 0.12480679 | 9.15912916 |
| GO:0008203~cholesterol metabolic process | 15 | 0.0715 | 0.00529572 | SREBF1, CYP | 1286 | 70 | 13588 | 2.26416352 | 0.99999996 | 0.12416616 | 9.17956336 |
| GO:0032270~positive regulation of cellular protein metabolic proc | 19 | 0.0906 | 0.00555428 | HMGB1, TGF | 1286 | 100 | 13588 | 2.00755832 | 0.99999998 | 0.1288942 | 9.60669451 |
| GO:0016044~membrane organization | 40 | 0.1908 | 0.00557278 | STON2, ABC | 1286 | 272 | 13588 | 1.55383771 | 0.99999998 | 0.12836778 | 9.63718223 |
| GO:0046907~intracellular transport | 58 | 0.2766 | 0.00578391 | COPA, AP1B1 | 1286 | 431 | 13588 | 1.42188769 | 0.99999999 | 0.13196162 | 9.98445165 |
| GO:0043244~regulation of protein complex disassembly | 11 | 0.0525 | 0.00582483 | GSN, CFL1, S | 1286 | 43 | 13588 | 2.7029549 | 0.99999999 | 0.1318961 | 10.0516174 |
| GO:0044092~negative regulation of molecular function | 23 | 0.1097 | 0.00607858 | GTPBP4, THR | 1286 | 132 | 13588 | 1.84106226 | 1 | 0.13628681 | 10.4670224 |
| GO:0060485~mesenchyme development | 12 | 0.0572 | 0.00611908 | ALDH1A2, FG | 1286 | 50 | 13588 | 2.53586314 | 1 | 0.13618176 | 10.5331479 |
| GO:0010769~regulation of cell morphogenesis involved in differen | 12 | 0.0572 | 0.00611908 | NRP1, ULK1, | 1286 | 50 | 13588 | 2.53586314 | 1 | 0.13618176 | 10.5331479 |
| GO:0043524~negative regulation of neuron apoptosis | 12 | 0.0572 | 0.00611908 | NRAS, BDNF, | 1286 | 50 | 13588 | 2.53586314 | 1 | 0.13618176 | 10.5331479 |
| GO:0007018~microtubule-based movement | 19 | 0.0906 | 0.00617766 | KIF23, DYNC | 1286 | 101 | 13588 | 1.98768151 | 1 | 0.13645048 | 10.6287309 |
| GO:0046578~regulation of Ras protein signal transduction | 29 | 0.1383 | 0.00629154 | ARFGAP3, TB | 1286 | 181 | 13588 | 1.69291048 | 1 | 0.13784338 | 10.8142551 |
| GO:0016049~cell growth | 10 | 0.0477 | 0.00639757 | NOTCH2, API | 1286 | 37 | 13588 | 2.85570174 | 1 | 0.13905534 | 10.9866629 |
| GO:0030335~positive regulation of cell migration | 10 | 0.0477 | 0.00639757 | COL18A1, HM | 1286 | 37 | 13588 | 2.85570174 | 1 | 0.13905534 | 10.9866629 |
| GO:0060444~branching involved in mammary gland duct morphog | 7 | 0.0334 | 0.0065133 | WNT5A, DDR | 1286 | 19 | 13588 | 3.89277237 | 1 | 0.14044256 | 11.174474 |
| GO:0045793~positive regulation of cell size | 7 | 0.0334 | 0.0065133 | BCL2, CD81, | 1286 | 19 | 13588 | 3.89277237 | 1 | 0.14044256 | 11.174474 |
| GO:0051253~negative regulation of RNA metabolic process | 44 | 0.2098 | 0.0067685 | GLIS3, THRB, | 1286 | 310 | 13588 | 1.49970401 | 1 | 0.14457413 | 11.5873171 |
| GO:0051325~interphase | 12 | 0.0572 | 0.00715115 | CCND1, APP, | 1286 | 51 | 13588 | 2.48614033 | 1 | 0.15111958 | 12.2029393 |
| GO:0001952~regulation of cell-matrix adhesion | 6 | 0.0286 | 0.00725707 | DDR1, PIK3C | 1286 | 14 | 13588 | 4.52832704 | 1 | 0.1521855 | 12.3726428 |
| GO:0021535~cell migration in hindbrain | 4 | 0.0191 | 0.00729005 | PLXNA2, ULK | 1286 | 5 | 13588 | 8.45287714 | 1 | 0.15183358 | 12.4254099 |
| GO:0046668~regulation of retinal cell programmed cell death | 4 | 0.0191 | 0.00729005 | ZFP91, BDNF | 1286 | 5 | 13588 | 8.45287714 | 1 | 0.15183358 | 12.4254099 |
| GO:0060762~regulation of branching involved in mammary gland c | 4 | 0.0191 | 0.00729005 | WNT5A, ETV | 1286 | 5 | 13588 | 8.45287714 | 1 | 0.15183358 | 12.4254099 |
| GO:0007169~transmembrane receptor protein tyrosine kinase sigr | 30 | 0.1431 | 0.00767544 | FGFR1, NRTN | 1286 | 192 | 13588 | 1.65095257 | 1 | 0.15819356 | 13.0398811 |
| GO:0007346~regulation of mitotic cell cycle | 18 | 0.0858 | 0.00816358 | HMGB1, ILKA | 1286 | 96 | 13588 | 1.98114308 | 1 | 0.16634004 | 13.8123173 |
| GO:0032868~response to insulin stimulus | 13 | 0.062 | 0.00826243 | IRS2, WDTC1 | 1286 | 59 | 13588 | 2.32812294 | 1 | 0.1671252 | 13.967957 |
| GO:0006813~potassium ion transport | 26 | 0.124 | 0.00835973 | HCN1, KCNC | 1286 | 160 | 13588 | 1.71699067 | 1 | 0.16787052 | 14.1208904 |
| GO:0048741~skeletal muscle fiber development | 9 | 0.0429 | 0.00836999 | GPX1, APP, D | 1286 | 32 | 13588 | 2.97171462 | 1 | 0.16701699 | 14.137 |
| GO:0007243~protein kinase cascade | 35 | 0.1669 | 0.00857945 | GNA13, WNT | 1286 | 236 | 13588 | 1.56700583 | 1 | 0.16978369 | 14.4652782 |
| GO:0044087~regulation of cellular component biogenesis | 17 | 0.0811 | 0.00874153 | PSRC1, CAPZ | 1286 | 89 | 13588 | 2.01824314 | 1 | 0.17165644 | 14.7185038 |
| GO:0006357~regulation of transcription from RNA polymerase II pı | 77 | 0.3672 | 0.0091501 | THRA, THRB, | 1286 | 616 | 13588 | 1.32076205 | 1 | 0.17786906 | 15.353668 |
| GO:0032956~regulation of actin cytoskeleton organization | 13 | 0.062 | 0.00946305 | CAPZA1, RDX | 1286 | 60 | 13588 | 2.28932089 | 1 | 0.18228757 | 15.8371575 |
| GO:0019725~cellular homeostasis | 47 | 0.2242 | 0.00953525 | TMX1, GNA1 | 1286 | 343 | 13588 | 1.44783245 | 1 | 0.18245808 | 15.9483206 |
| GO:0045792~negative regulation of cell size | 12 | 0.0572 | 0.00961847 | EI24, NRP1, | 1286 | 53 | 13588 | 2.39232372 | 1 | 0.1828165 | 16.0762914 |
| GO:0045892~negative regulation of transcription, DNA-dependent | 43 | 0.2051 | 0.00984878 | GLIS3, THRB, | 1286 | 308 | 13588 | 1.47513684 | 1 | 0.18568441 | 16.4294867 |
| GO:0008283~cell proliferation | 36 | 0.1717 | 0.00994345 | WNT5A, ELF | 1286 | 247 | 13588 | 1.53999786 | 1 | 0.18621064 | 16.5742685 |
| GO:0050772~positive regulation of axonogenesis | 6 | 0.0286 | 0.01005033 | PLXNB2, ROE | 1286 | 15 | 13588 | 4.22643857 | 1 | 0.18693535 | 16.737432 |
| GO:0021955~central nervous system neuron axonogenesis | 6 | 0.0286 | 0.01005033 | DCC, EPHB3, | 1286 | 15 | 13588 | 4.22643857 | 1 | 0.18693535 | 16.737432 |
| GO:0021954~central nervous system neuron development | 9 | 0.0429 | 0.0101535 | DCC, ARX, AS | 1286 | 33 | 13588 | 2.88166266 | 1 | 0.1875883 | 16.8946354 |
| GO:0045446~endothelial cell differentiation | 5 | 0.0238 | 0.01049044 | COL18A1, GF | 1286 | 10 | 13588 | 5.28304821 | 1 | 0.19210239 | 17.4061286 |
| GO:0032970~regulation of actin filament-based process | 13 | 0.062 | 0.01079486 | CAPZA1, RDX | 1286 | 61 | 13588 | 2.25179104 | 1 | 0.19600569 | 17.8656818 |
| GO:0010721~negative regulation of cell development | 10 | 0.0477 | 0.01088097 | BDNF, NRP1, | 1286 | 40 | 13588 | 2.64152411 | 1 | 0.1963074 | 17.9952336 |
| GO:0051640~organelle localization | 12 | 0.0572 | 0.01107483 | MFN2, FMN2 | 1286 | 54 | 13588 | 2.34802143 | 1 | 0.19834114 | 18.2861988 |
| GO:0014031~mesenchymal cell development | 11 | 0.0525 | 0.01113912 | ALDH1A2, ED | 1286 | 47 | 13588 | 2.47291618 | 1 | 0.19827531 | 18.3824709 |
| GO:0042592~homeostatic process | 73 | 0.3481 | 0.01127594 | LDLR, GNA11 | 1286 | 584 | 13588 | 1.32076205 | 1 | 0.19936281 | 18.5870101 |
| GO:0050804~regulation of synaptic transmission | 18 | 0.0858 | 0.01218537 | RAB3A, MYO | 1286 | 100 | 13588 | 1.90189736 | 1 | 0.2125259 | 19.9342807 |
| GO:0051656~establishment of organelle localization | 9 | 0.0429 | 0.01220392 | FMN2, KIF1B | 1286 | 34 | 13588 | 2.79690788 | 1 | 0.21167151 | 19.9615335 |
| GO:0010639~negative regulation of organelle organization | 13 | 0.062 | 0.01226675 | CAPZA1, ESR | 1286 | 62 | 13588 | 2.21547183 | 1 | 0.21150754 | 20.0538114 |
| GO:0045596~negative regulation of cell differentiation | 28 | 0.1335 | 0.01237622 | HMGB3, NRF | 1286 | 182 | 13588 | 1.6255533 | 1 | 0.21205785 | 20.2143284 |
| GO:0016125~sterol metabolic process | 15 | 0.0715 | 0.01239458 | SREBF1, CYP | 1286 | 77 | 13588 | 2.05833047 | 1 | 0.2112183 | 20.2412239 |
| GO:0050769~positive regulation of neurogenesis | 10 | 0.0477 | 0.01280217 | ASCL1, PLXN | 1286 | 41 | 13588 | 2.57709669 | 1 | 0.21625225 | 20.8360159 |

| | | | | | | | | | | | |
|---|---|---|---|---|---|---|---|---|---|---|---|
| GO:0048167~regulation of synaptic plasticity | 11 | 0.0525 | 0.01291076 | SYP, GRM5, / | 1286 | 48 | 13588 | 2.4213971 | 1 | 0.21674951 | 20.9937682 |
| GO:0043408~regulation of MAPKKK cascade | 17 | 0.0811 | 0.01321147 | FGFR1, PSAP | 1286 | 93 | 13588 | 1.93143698 | 1 | 0.22008377 | 21.4290869 |
| GO:0001525~angiogenesis | 22 | 0.1049 | 0.01321505 | COL18A1, GN | 1286 | 133 | 13588 | 1.74777535 | 1 | 0.21900852 | 21.4342589 |
| GO:0050810~regulation of steroid biosynthetic process | 6 | 0.0286 | 0.01350012 | SREBF1, INSI | 1286 | 16 | 13588 | 3.96228616 | 1 | 0.22205528 | 21.8448101 |
| GO:0048598~embryonic morphogenesis | 48 | 0.2289 | 0.0135323 | CER1, WNT5, | 1286 | 359 | 13588 | 1.41273713 | 1 | 0.2213985 | 21.8910305 |
| GO:0032411~positive regulation of transporter activity | 4 | 0.0191 | 0.01355861 | PLCG2, STIM | 1286 | 6 | 13588 | 7.04406428 | 1 | 0.22066404 | 21.9288054 |
| GO:0046890~regulation of lipid biosynthetic process | 7 | 0.0334 | 0.01401642 | SREBF1, WD | 1286 | 22 | 13588 | 3.36193977 | 1 | 0.22611046 | 22.5832352 |
| GO:0030900~forebrain development | 26 | 0.124 | 0.01407791 | WNT5A, SOX | 1286 | 167 | 13588 | 1.645021 | 1 | 0.22586115 | 22.670735 |
| GO:0010628~positive regulation of gene expression | 62 | 0.2957 | 0.01409124 | E2F3, MEF2A | 1286 | 488 | 13588 | 1.34241389 | 1 | 0.22493505 | 22.6896827 |
| GO:0044057~regulation of system process | 30 | 0.1431 | 0.01412387 | RAB3A, EDN: | 1286 | 201 | 13588 | 1.57702932 | 1 | 0.22428953 | 22.7360727 |
| GO:0051329~interphase of mitotic cell cycle | 11 | 0.0525 | 0.01488591 | CCND1, APP, | 1286 | 49 | 13588 | 2.37198083 | 1 | 0.23378666 | 23.8119093 |
| GO:0001657~ureteric bud development | 10 | 0.0477 | 0.01496429 | FGFR1, BDNF | 1286 | 42 | 13588 | 2.51573724 | 1 | 0.23373588 | 23.921759 |
| GO:0009890~negative regulation of biosynthetic process | 56 | 0.2671 | 0.01498353 | THRA, THRB, | 1286 | 434 | 13588 | 1.36336728 | 1 | 0.23287724 | 23.9486901 |
| GO:0051056~regulation of small GTPase mediated signal transduct | 33 | 0.1574 | 0.01511819 | ARFGAP3, RA | 1286 | 228 | 13588 | 1.52930343 | 1 | 0.23359736 | 24.1370141 |
| GO:0007059~chromosome segregation | 13 | 0.062 | 0.01566672 | NUSAP1, CEN | 1286 | 64 | 13588 | 2.14623834 | 1 | 0.23987948 | 24.899542 |
| GO:0006979~response to oxidative stress | 16 | 0.0763 | 0.01580073 | TXNIP, EPAS: | 1286 | 87 | 13588 | 1.94319015 | 1 | 0.24054016 | 25.0847338 |
| GO:0000226~microtubule cytoskeleton organization | 19 | 0.0906 | 0.01608527 | KIF11, PSRC1 | 1286 | 111 | 13588 | 1.8086111 | 1 | 0.24318543 | 25.4765163 |
| GO:0007167~enzyme linked receptor protein signaling pathway | 38 | 0.1812 | 0.0160984 | FGFR1, NRTN | 1286 | 273 | 13588 | 1.4707387 | 1 | 0.24222969 | 25.4945409 |
| GO:0016481~negative regulation of transcription | 49 | 0.2337 | 0.01627316 | GLIS3, THRA, | 1286 | 372 | 13588 | 1.39177077 | 1 | 0.24339906 | 25.7341474 |
| GO:0031344~regulation of cell projection organization | 12 | 0.0572 | 0.01646339 | NRP1, ULK1, | 1286 | 57 | 13588 | 2.22444135 | 1 | 0.24475568 | 25.9941461 |
| GO:0032271~regulation of protein polymerization | 12 | 0.0572 | 0.01646339 | ACTR3, GSN, | 1286 | 57 | 13588 | 2.22444135 | 1 | 0.24475568 | 25.9941461 |
| GO:0031589~cell-substrate adhesion | 12 | 0.0572 | 0.01646339 | VWF, SORBS: | 1286 | 57 | 13588 | 2.22444135 | 1 | 0.24475568 | 25.9941461 |
| GO:0007519~skeletal muscle tissue development | 14 | 0.0668 | 0.01651437 | MET, CACNB | 1286 | 72 | 13588 | 2.05451875 | 1 | 0.24429815 | 26.0636675 |
| GO:0010558~negative regulation of macromolecule biosynthetic p | 54 | 0.2575 | 0.01653013 | THRA, THRB, | 1286 | 418 | 13588 | 1.3649981 | 1 | 0.24339201 | 26.0851533 |
| GO:0009891~positive regulation of biosynthetic process | 69 | 0.3291 | 0.01659197 | E2F3, MEF2A | 1286 | 557 | 13588 | 1.30890602 | 1 | 0.24308487 | 26.1693936 |
| GO:0040008~regulation of growth | 36 | 0.1717 | 0.01668974 | FGFR1, NRP1 | 1286 | 256 | 13588 | 1.48585731 | 1 | 0.24323874 | 26.3023779 |
| GO:0008064~regulation of actin polymerization or depolymerizatic | 11 | 0.0525 | 0.01707762 | ACTR3, GSN, | 1286 | 50 | 13588 | 2.32454121 | 1 | 0.24706068 | 26.8277685 |
| GO:0000302~response to reactive oxygen species | 9 | 0.0429 | 0.01719063 | GPX1, ERCC6 | 1286 | 36 | 13588 | 2.64152411 | 1 | 0.24738385 | 26.9801774 |
| GO:0010557~positive regulation of macromolecule biosynthetic pr | 66 | 0.3148 | 0.01719405 | E2F3, MEF2A | 1286 | 530 | 13588 | 1.31577805 | 1 | 0.24633441 | 26.9847721 |
| GO:0010629~negative regulation of gene expression | 53 | 0.2528 | 0.01737031 | THRA, THRB, | 1286 | 410 | 13588 | 1.36586124 | 1 | 0.2474447 | 27.2218657 |
| GO:0042493~response to drug | 16 | 0.0763 | 0.01743679 | SNX27, SNCA | 1286 | 88 | 13588 | 1.92110844 | 1 | 0.24718603 | 27.3111017 |
| GO:0001755~neural crest cell migration | 7 | 0.0334 | 0.01748185 | EDN3, NRTN, | 1286 | 23 | 13588 | 3.21576848 | 1 | 0.24666623 | 27.3715301 |
| GO:0043254~regulation of protein complex assembly | 13 | 0.062 | 0.01761275 | PSRC1, CAPZ | 1286 | 65 | 13588 | 2.11321928 | 1 | 0.24720305 | 27.5467896 |
| GO:0045941~positive regulation of transcription | 60 | 0.2862 | 0.01786046 | E2F3, MEF2A | 1286 | 475 | 13588 | 1.33466481 | 1 | 0.24915567 | 27.8773694 |
| GO:0031327~negative regulation of cellular biosynthetic process | 55 | 0.2623 | 0.01857802 | THRA, THRB, | 1286 | 430 | 13588 | 1.35147745 | 1 | 0.25673464 | 28.82694 |
| GO:0019216~regulation of lipid metabolic process | 12 | 0.0572 | 0.01863512 | SREBF1, WD | 1286 | 58 | 13588 | 2.18608892 | 1 | 0.25632894 | 28.9019831 |
| GO:0031644~regulation of neurological system process | 19 | 0.0906 | 0.01906946 | RAB3A, MYO | 1286 | 113 | 13588 | 1.77660028 | 1 | 0.26039583 | 29.4704394 |
| GO:0031328~positive regulation of cellular biosynthetic process | 68 | 0.3243 | 0.01939806 | E2F3, MEF2A | 1286 | 552 | 13588 | 1.30162057 | 1 | 0.26316568 | 29.8976525 |
| GO:0006469~negative regulation of protein kinase activity | 11 | 0.0525 | 0.01949871 | SPRY2, CDKN | 1286 | 51 | 13588 | 2.27896197 | 1 | 0.26324676 | 30.0280178 |
| GO:0032147~activation of protein kinase activity | 11 | 0.0525 | 0.01949871 | DGKB, ERCC6 | 1286 | 51 | 13588 | 2.27896197 | 1 | 0.26324676 | 30.0280178 |
| GO:0033673~negative regulation of kinase activity | 11 | 0.0525 | 0.01949871 | SPRY2, CDKN | 1286 | 51 | 13588 | 2.27896197 | 1 | 0.26324676 | 30.0280178 |
| GO:0030832~regulation of actin filament length | 11 | 0.0525 | 0.01949871 | ACTR3, GSN, | 1286 | 51 | 13588 | 2.27896197 | 1 | 0.26324676 | 30.0280178 |
| GO:0051276~chromosome organization | 52 | 0.248 | 0.01987634 | HP1BP3, UTY | 1286 | 404 | 13588 | 1.35999261 | 1 | 0.26653385 | 30.5150815 |
| GO:0040017~positive regulation of locomotion | 10 | 0.0477 | 0.02007581 | COL18A1, HN | 1286 | 44 | 13588 | 2.40138555 | 1 | 0.2677337 | 30.7710717 |
| GO:0060538~skeletal muscle organ development | 14 | 0.0668 | 0.02046406 | MET, CACNB | 1286 | 74 | 13588 | 1.99899122 | 1 | 0.27107176 | 31.2667726 |
| GO:0000165~MAPKKK cascade | 19 | 0.0906 | 0.02071304 | WNT5A, FGF | 1286 | 114 | 13588 | 1.76101607 | 1 | 0.27279232 | 31.5828838 |
| GO:0051173~positive regulation of nitrogen compound metabolic | 65 | 0.31 | 0.02075815 | E2F3, MEF2A | 1286 | 526 | 13588 | 1.30569633 | 1 | 0.2722017 | 31.6400187 |
| GO:0043406~positive regulation of MAP kinase activity | 12 | 0.0572 | 0.02101219 | SPAG9, MAP | 1286 | 59 | 13588 | 2.14903656 | 1 | 0.27395557 | 31.9608922 |
| GO:0042744~hydrogen peroxide catabolic process | 5 | 0.0238 | 0.02121589 | GPX1, GPX4, | 1286 | 12 | 13588 | 4.40254018 | 1 | 0.27513031 | 32.2171504 |
| GO:0070301~cellular response to hydrogen peroxide | 5 | 0.0238 | 0.02121589 | GPX1, GPX4, | 1286 | 12 | 13588 | 4.40254018 | 1 | 0.27513031 | 32.2171504 |
| GO:0030834~regulation of actin filament depolymerization | 7 | 0.0334 | 0.02149239 | GSN, CFL1, S | 1286 | 24 | 13588 | 3.08177812 | 1 | 0.27709634 | 32.563537 |
| GO:0045934~negative regulation of nucleobase, nucleoside, nucle | 51 | 0.2432 | 0.02178867 | GLIS3, THRA, | 1286 | 397 | 13588 | 1.35735747 | 1 | 0.27925639 | 32.932851 |
| GO:0030855~epithelial cell differentiation | 20 | 0.0954 | 0.02203055 | COL18A1, PS | 1286 | 123 | 13588 | 1.71806446 | 1 | 0.28079928 | 33.232931 |
| GO:0006265~DNA topological change | 4 | 0.0191 | 0.02207393 | TOP1, TOP3A | 1286 | 7 | 13588 | 6.03776938 | 1 | 0.28017649 | 33.2866214 |
| GO:0002011~morphogenesis of an epithelial sheet | 4 | 0.0191 | 0.02207393 | NOTCH2, DA | 1286 | 7 | 13588 | 6.03776938 | 1 | 0.28017649 | 33.2866214 |
| GO:0010638~positive regulation of organelle organization | 11 | 0.0525 | 0.02216171 | MEN1, CFL1, | 1286 | 52 | 13588 | 2.23513578 | 1 | 0.28003826 | 33.3951279 |
| GO:0034614~cellular response to reactive oxygen species | 6 | 0.0286 | 0.02259993 | GPX1, GPX4, | 1286 | 18 | 13588 | 3.52203214 | 1 | 0.28365863 | 33.9343183 |
| GO:0007274~neuromuscular synaptic transmission | 6 | 0.0286 | 0.02259993 | RAB3A, KIF1E | 1286 | 18 | 13588 | 3.52203214 | 1 | 0.28365863 | 33.9343183 |
| GO:0051969~regulation of transmission of nerve impulse | 18 | 0.0858 | 0.0228085 | RAB3A, MYO | 1286 | 107 | 13588 | 1.77747416 | 1 | 0.28479246 | 34.1895039 |
| GO:0030833~regulation of actin filament polymerization | 10 | 0.0477 | 0.02305633 | ACTR3, GSN, | 1286 | 45 | 13588 | 2.34802143 | 1 | 0.28632952 | 34.4915038 |
| GO:0007595~lactation | 8 | 0.0382 | 0.02314966 | ATP2B2, CCN | 1286 | 31 | 13588 | 2.72673456 | 1 | 0.28622667 | 34.6048976 |
| GO:0045935~positive regulation of nucleobase, nucleoside, nucleo | 63 | 0.3005 | 0.02316003 | E2F3, MEF2A | 1286 | 510 | 13588 | 1.30522368 | 1 | 0.28525531 | 34.6174818 |

| | | | | | | | | | | | |
|---|---|---|---|---|---|---|---|---|---|---|---|
| GO:0048747~muscle fiber development | 9 | 0.0429 | 0.0234941 | GPX1, APP, D | 1286 | 38 | 13588 | 2.50249652 | 1 | 0.2876679 | 35.0217756 |
| GO:0050768~negative regulation of neurogenesis | 9 | 0.0429 | 0.0234941 | BDNF, NRP1, | 1286 | 38 | 13588 | 2.50249652 | 1 | 0.2876679 | 35.0217756 |
| GO:0050905~neuromuscular process | 12 | 0.0572 | 0.02360485 | ATP2B2, APP | 1286 | 60 | 13588 | 2.11321928 | 1 | 0.28774161 | 35.1552823 |
| GO:0051348~negative regulation of transferase activity | 11 | 0.0525 | 0.02507875 | SPRY2, CDKN | 1286 | 53 | 13588 | 2.19296341 | 1 | 0.30174879 | 36.9075189 |
| GO:0021543~pallium development | 11 | 0.0525 | 0.02507875 | ARX, BBS1, A | 1286 | 53 | 13588 | 2.19296341 | 1 | 0.30174879 | 36.9075189 |
| GO:0048589~developmental growth | 17 | 0.0811 | 0.0251084 | WNT5A, SETI | 1286 | 100 | 13588 | 1.79623639 | 1 | 0.30094256 | 36.9423127 |
| GO:0050957~equilibrioception | 3 | 0.0143 | 0.0251229 | SOX2, POU4F | 1286 | 3 | 13588 | 10.5660964 | 1 | 0.29998994 | 36.9593172 |
| GO:0007412~axon target recognition | 3 | 0.0143 | 0.0251229 | BDNF, UCHL1 | 1286 | 3 | 13588 | 10.5660964 | 1 | 0.29998994 | 36.9593172 |
| GO:0051386~regulation of nerve growth factor receptor signaling p | 3 | 0.0143 | 0.0251229 | SPRY2, SPRY1 | 1286 | 3 | 13588 | 10.5660964 | 1 | 0.29998994 | 36.9593172 |
| GO:0010035~response to inorganic substance | 16 | 0.0763 | 0.02530308 | CASR, ROMO | 1286 | 92 | 13588 | 1.83758199 | 1 | 0.30070508 | 37.1702578 |
| GO:0022612~gland morphogenesis | 15 | 0.0715 | 0.02531199 | WNT5A, FGF | 1286 | 84 | 13588 | 1.88680293 | 1 | 0.29970576 | 37.1806799 |
| GO:0003001~generation of a signal involved in cell-cell signaling | 15 | 0.0715 | 0.02531199 | SYT1, RAB3A | 1286 | 84 | 13588 | 1.88680293 | 1 | 0.29970576 | 37.1806799 |
| GO:0003006~reproductive developmental process | 36 | 0.1717 | 0.02540316 | WNT5A, SEPT | 1286 | 264 | 13588 | 1.44083133 | 1 | 0.29953196 | 37.2871376 |
| GO:0051172~negative regulation of nitrogen compound metabolic | 51 | 0.2432 | 0.02551434 | GLIS3, THRA, | 1286 | 401 | 13588 | 1.34381775 | 1 | 0.299558 | 37.416747 |
| GO:0030837~negative regulation of actin filament polymerization | 7 | 0.0334 | 0.02608064 | GSN, SCIN, C | 1286 | 25 | 13588 | 2.958507 | 1 | 0.30405815 | 38.0729778 |
| GO:0031647~regulation of protein stability | 7 | 0.0334 | 0.02608064 | GTPBP4, CCD | 1286 | 25 | 13588 | 2.958507 | 1 | 0.30405815 | 38.0729778 |
| GO:0042692~muscle cell differentiation | 19 | 0.0906 | 0.02629836 | RXRA, MET, I | 1286 | 117 | 13588 | 1.71586181 | 1 | 0.30510442 | 38.3235365 |
| GO:0050770~regulation of axonogenesis | 9 | 0.0429 | 0.02718698 | NRP1, ULK1, | 1286 | 39 | 13588 | 2.43832994 | 1 | 0.31261592 | 39.3362863 |
| GO:0043086~negative regulation of catalytic activity | 17 | 0.0811 | 0.02731211 | HMGCR, INTS | 1286 | 101 | 13588 | 1.77845187 | 1 | 0.31272147 | 39.4776223 |
| GO:0051693~actin filament capping | 6 | 0.0286 | 0.02834515 | GSN, SCIN, C | 1286 | 19 | 13588 | 3.33666203 | 1 | 0.32142676 | 40.6326642 |
| GO:0048169~regulation of long-term neuronal synaptic plasticity | 6 | 0.0286 | 0.02834515 | SYP, GRM5, I | 1286 | 19 | 13588 | 3.33666203 | 1 | 0.32142676 | 40.6326642 |
| GO:0032392~DNA geometric change | 5 | 0.0238 | 0.02839888 | TOP1, HMGB | 1286 | 13 | 13588 | 4.06388324 | 1 | 0.32082483 | 40.692168 |
| GO:0021544~subpallium development | 5 | 0.0238 | 0.02839888 | BBS1, ASCL1, | 1286 | 13 | 13588 | 4.06388324 | 1 | 0.32082483 | 40.692168 |
| GO:0033554~cellular response to stress | 51 | 0.2432 | 0.02893105 | WNT5A, HM | 1286 | 404 | 13588 | 1.3338389 | 1 | 0.32469028 | 41.2784696 |
| GO:0048562~embryonic organ morphogenesis | 24 | 0.1145 | 0.02952649 | FGFR1, SATB | 1286 | 161 | 13588 | 1.57507027 | 1 | 0.32908719 | 41.9279856 |
| GO:0015672~monovalent inorganic cation transport | 40 | 0.1908 | 0.02958594 | KCNC2, SLC3 | 1286 | 303 | 13588 | 1.39486421 | 1 | 0.32852094 | 41.9924646 |
| GO:0014706~striated muscle tissue development | 20 | 0.0954 | 0.02972999 | RXRA, MET, I | 1286 | 127 | 13588 | 1.66395219 | 1 | 0.32873231 | 42.1484123 |
| GO:0030001~metal ion transport | 55 | 0.2623 | 0.02975806 | KCNC2, KCNI | 1286 | 442 | 13588 | 1.31478575 | 1 | 0.32788606 | 42.1787459 |
| GO:0010975~regulation of neuron projection development | 10 | 0.0477 | 0.02993989 | NRP1, ULK1, | 1286 | 47 | 13588 | 2.24810562 | 1 | 0.32844145 | 42.3749372 |
| GO:0030308~negative regulation of cell growth | 10 | 0.0477 | 0.02993989 | EI24, NRP1, U | 1286 | 47 | 13588 | 2.24810562 | 1 | 0.32844145 | 42.3749372 |
| GO:0030155~regulation of cell adhesion | 16 | 0.0763 | 0.03010816 | GTPBP4, EGF | 1286 | 94 | 13588 | 1.7984845 | 1 | 0.32886968 | 42.5559276 |
| GO:0048568~embryonic organ development | 33 | 0.1574 | 0.03088706 | FGFR1, SOX2 | 1286 | 241 | 13588 | 1.44680988 | 1 | 0.3347607 | 43.3867339 |
| GO:0048015~phosphoinositide-mediated signaling | 9 | 0.0429 | 0.03126349 | S1PR1, GNAC | 1286 | 40 | 13588 | 2.3773717 | 1 | 0.33700324 | 43.7841729 |
| GO:0060603~mammary gland duct morphogenesis | 7 | 0.0334 | 0.03127512 | WNT5A, DDR | 1286 | 26 | 13588 | 2.84471827 | 1 | 0.33600637 | 43.796412 |
| GO:0009260~ribonucleotide biosynthetic process | 18 | 0.0858 | 0.03142181 | TCIRG1, ATP1 | 1286 | 111 | 13588 | 1.71342104 | 1 | 0.33620506 | 43.950544 |
| GO:0014033~neural crest cell differentiation | 8 | 0.0382 | 0.03190109 | ALDH1A2, ED | 1286 | 33 | 13588 | 2.56147792 | 1 | 0.33930987 | 44.4513525 |
| GO:0014032~neural crest cell development | 8 | 0.0382 | 0.03190109 | ALDH1A2, ED | 1286 | 33 | 13588 | 2.56147792 | 1 | 0.33930987 | 44.4513525 |
| GO:0043491~protein kinase B signaling cascade | 4 | 0.0191 | 0.03287003 | RPS6KB2, PIK | 1286 | 8 | 13588 | 5.28304821 | 1 | 0.34660248 | 45.4509295 |
| GO:0045176~apical protein localization | 4 | 0.0191 | 0.03287003 | HCN1, VANG | 1286 | 8 | 13588 | 5.28304821 | 1 | 0.34660248 | 45.4509295 |
| GO:0045540~regulation of cholesterol biosynthetic process | 4 | 0.0191 | 0.03287003 | SREBF1, PEX5 | 1286 | 8 | 13588 | 5.28304821 | 1 | 0.34660248 | 45.4509295 |
| GO:0008593~regulation of Notch signaling pathway | 4 | 0.0191 | 0.03287003 | ASCL1, SOX2, | 1286 | 8 | 13588 | 5.28304821 | 1 | 0.34660248 | 45.4509295 |
| GO:0007406~negative regulation of neuroblast proliferation | 4 | 0.0191 | 0.03287003 | BDNF, CTNN | 1286 | 8 | 13588 | 5.28304821 | 1 | 0.34660248 | 45.4509295 |
| GO:0006164~purine nucleotide biosynthetic process | 21 | 0.1002 | 0.03308039 | TCIRG1, ADS | 1286 | 137 | 13588 | 1.61962062 | 1 | 0.34730125 | 45.6656766 |
| GO:0051494~negative regulation of cytoskeleton organization | 10 | 0.0477 | 0.03386969 | S1PR1, GSN, | 1286 | 48 | 13588 | 2.20127009 | 1 | 0.35290972 | 46.4643709 |
| GO:0042254~ribosome biogenesis | 18 | 0.0858 | 0.03390803 | KRR1, GTPBP | 1286 | 112 | 13588 | 1.69812264 | 1 | 0.35212345 | 46.5028807 |
| GO:0046486~glycerolipid metabolic process | 20 | 0.0954 | 0.03426226 | LPL, PLD1, PI | 1286 | 129 | 13588 | 1.63815448 | 1 | 0.35399582 | 46.857466 |
| GO:0060079~regulation of excitatory postsynaptic membrane pote | 6 | 0.0286 | 0.03493595 | SNCA, GRIK5 | 1286 | 20 | 13588 | 3.16982893 | 1 | 0.35850534 | 47.5257026 |
| GO:0043270~positive regulation of ion transport | 6 | 0.0286 | 0.03493595 | ATP2B2, PLC | 1286 | 20 | 13588 | 3.16982893 | 1 | 0.35850534 | 47.5257026 |
| GO:0043407~negative regulation of MAP kinase activity | 6 | 0.0286 | 0.03493595 | SPRY2, SPRY1 | 1286 | 20 | 13588 | 3.16982893 | 1 | 0.35850534 | 47.5257026 |
| GO:0007212~dopamine receptor signaling pathway | 6 | 0.0286 | 0.03493595 | CALY, GNAQ, | 1286 | 20 | 13588 | 3.16982893 | 1 | 0.35850534 | 47.5257026 |
| GO:0051254~positive regulation of RNA metabolic process | 52 | 0.248 | 0.03630219 | E2F3, THRA, | 1286 | 419 | 13588 | 1.31130552 | 1 | 0.3686184 | 48.8566012 |
| GO:0032409~regulation of transporter activity | 5 | 0.0238 | 0.0368556 | PLCG2, PKD2 | 1286 | 14 | 13588 | 3.77360587 | 1 | 0.37197684 | 49.3865664 |
| GO:0046579~positive regulation of Ras protein signal transduction | 5 | 0.0238 | 0.0368556 | NRAS, NOTCI | 1286 | 14 | 13588 | 3.77360587 | 1 | 0.37197684 | 49.3865664 |
| GO:0045665~negative regulation of neuron differentiation | 8 | 0.0382 | 0.03700777 | ZFP91, SOX2, | 1286 | 34 | 13588 | 2.48614033 | 1 | 0.37207332 | 49.5313718 |
| GO:0031333~negative regulation of protein complex assembly | 7 | 0.0334 | 0.03709986 | GSN, SCIN, C | 1286 | 27 | 13588 | 2.73935833 | 1 | 0.37168754 | 49.6188183 |
| GO:0032272~negative regulation of protein polymerization | 7 | 0.0334 | 0.03709986 | GSN, SCIN, C | 1286 | 27 | 13588 | 2.73935833 | 1 | 0.37168754 | 49.6188183 |
| GO:0010811~positive regulation of cell-substrate adhesion | 7 | 0.0334 | 0.03709986 | FBLN2, EGFL | 1286 | 27 | 13588 | 2.73935833 | 1 | 0.37168754 | 49.6188183 |
| GO:0050808~synapse organization | 10 | 0.0477 | 0.03814105 | ATP2B2, APP | 1286 | 49 | 13588 | 2.15634621 | 1 | 0.3788515 | 50.5975916 |
| GO:0022604~regulation of cell morphogenesis | 16 | 0.0763 | 0.0385261 | GNA13, NRP1 | 1286 | 97 | 13588 | 1.74286127 | 1 | 0.38075356 | 50.9549952 |
| GO:0010608~posttranscriptional regulation of gene expression | 22 | 0.1049 | 0.03872089 | ZFP36, GTPB | 1286 | 148 | 13588 | 1.57063595 | 1 | 0.38114972 | 51.1348639 |
| GO:0014020~primary neural tube formation | 9 | 0.0429 | 0.04062712 | KAT2A, COBL | 1286 | 42 | 13588 | 2.26416352 | 1 | 0.39474435 | 52.8624758 |
| GO:0016071~mRNA metabolic process | 39 | 0.186 | 0.04171256 | SCAF1, RNAS | 1286 | 302 | 13588 | 1.3644959 | 1 | 0.40181942 | 53.8202607 |

| | | | | | | | | | | | |
|---|---|---|---|---|---|---|---|---|---|---|---|
| GO:0030835~negative regulation of actin filament depolymerizatio | 6 | 0.0286 | 0.04239723 | GSN, SCIN, C | 1286 | 21 | 13588 | 3.01888469 | 1 | 0.40578747 | 54.4149142 |
| GO:0043242~negative regulation of protein complex disassembly | 8 | 0.0382 | 0.04262405 | GSN, SCIN, C | 1286 | 35 | 13588 | 2.41510775 | 1 | 0.40631375 | 54.6103122 |
| GO:0045185~maintenance of protein location | 7 | 0.0334 | 0.04357419 | TLN1, FGFR1 | 1286 | 28 | 13588 | 2.64152411 | 1 | 0.41215968 | 55.4202675 |
| GO:0009152~purine ribonucleotide biosynthetic process | 17 | 0.0811 | 0.04365287 | TCIRG1, ATP | 1286 | 107 | 13588 | 1.6787256 | 1 | 0.41157292 | 55.4867227 |
| GO:0001822~kidney development | 17 | 0.0811 | 0.04365287 | FGFR1, NPNT | 1286 | 107 | 13588 | 1.6787256 | 1 | 0.41157292 | 55.4867227 |
| GO:0006886~intracellular protein transport | 36 | 0.1717 | 0.0446994 | STON2, PACS | 1286 | 276 | 13588 | 1.37818649 | 1 | 0.41800795 | 56.3617928 |
| GO:0006605~protein targeting | 20 | 0.0954 | 0.04482718 | PACS1, TXNI | 1286 | 133 | 13588 | 1.58888668 | 1 | 0.41776356 | 56.4675181 |
| GO:0045893~positive regulation of transcription, DNA-dependent | 51 | 0.2432 | 0.04527209 | E2F3, THRA, | 1286 | 416 | 13588 | 1.29536278 | 1 | 0.41978956 | 56.8337481 |
| GO:0010648~negative regulation of cell communication | 26 | 0.124 | 0.04572733 | CER1, ONECU | 1286 | 186 | 13588 | 1.47698122 | 1 | 0.42186854 | 57.205473 |
| GO:0046543~development of secondary female sexual characteris | 4 | 0.0191 | 0.04590592 | WNT5A, NCC | 1286 | 9 | 13588 | 4.69604285 | 1 | 0.42197447 | 57.3504667 |
| GO:0042304~regulation of fatty acid biosynthetic process | 4 | 0.0191 | 0.04590592 | SREBF1, WD | 1286 | 9 | 13588 | 4.69604285 | 1 | 0.42197447 | 57.3504667 |
| GO:0050910~detection of mechanical stimulus involved in sensory | 4 | 0.0191 | 0.04590592 | ATP2B2, SLC | 1286 | 9 | 13588 | 4.69604285 | 1 | 0.42197447 | 57.3504667 |
| GO:0033598~mammary gland epithelial cell proliferation | 4 | 0.0191 | 0.04590592 | WNT5A, CCN | 1286 | 9 | 13588 | 4.69604285 | 1 | 0.42197447 | 57.3504667 |
| GO:0032869~cellular response to insulin stimulus | 9 | 0.0429 | 0.04593993 | WDTC1, IRS2 | 1286 | 43 | 13588 | 2.21150855 | 1 | 0.42106109 | 57.3780279 |
| GO:0045727~positive regulation of translation | 5 | 0.0238 | 0.04660298 | EIF5A, PTMS, | 1286 | 15 | 13588 | 3.52203214 | 1 | 0.42456468 | 57.9119828 |
| GO:0031076~embryonic camera-type eye development | 5 | 0.0238 | 0.04660298 | ALDH1A1, AL | 1286 | 15 | 13588 | 3.52203214 | 1 | 0.42456468 | 57.9119828 |
| GO:0051057~positive regulation of small GTPase mediated signal t | 5 | 0.0238 | 0.04660298 | NRAS, NOTC | 1286 | 15 | 13588 | 3.52203214 | 1 | 0.42456468 | 57.9119828 |
| GO:0045747~positive regulation of Notch signaling pathway | 3 | 0.0143 | 0.04710888 | ASCL1, SOX2, | 1286 | 4 | 13588 | 7.92457232 | 1 | 0.42693574 | 58.3151314 |
| GO:0032234~regulation of calcium ion transport via store-operate | 3 | 0.0143 | 0.04710888 | PLCG2, STIM | 1286 | 4 | 13588 | 7.92457232 | 1 | 0.42693574 | 58.3151314 |
| GO:0050942~positive regulation of pigment cell differentiation | 3 | 0.0143 | 0.04710888 | ADAMTS9, B | 1286 | 4 | 13588 | 7.92457232 | 1 | 0.42693574 | 58.3151314 |
| GO:0045651~positive regulation of macrophage differentiation | 3 | 0.0143 | 0.04710888 | LIF, CSF1, RB | 1286 | 4 | 13588 | 7.92457232 | 1 | 0.42693574 | 58.3151314 |
| GO:0032236~positive regulation of calcium ion transport via store- | 3 | 0.0143 | 0.04710888 | PLCG2, STIM | 1286 | 4 | 13588 | 7.92457232 | 1 | 0.42693574 | 58.3151314 |
| GO:0045636~positive regulation of melanocyte differentiation | 3 | 0.0143 | 0.04710888 | ADAMTS9, B | 1286 | 4 | 13588 | 7.92457232 | 1 | 0.42693574 | 58.3151314 |
| GO:0006163~purine nucleotide metabolic process | 23 | 0.1097 | 0.04718803 | TCIRG1, ADS | 1286 | 160 | 13588 | 1.51887636 | 1 | 0.42633498 | 58.3778724 |
| GO:0080135~regulation of cellular response to stress | 13 | 0.062 | 0.04811446 | BRCC3, BECN | 1286 | 75 | 13588 | 1.83145671 | 1 | 0.43156102 | 59.1056552 |
| GO:0016192~vesicle-mediated transport | 56 | 0.2671 | 0.04972896 | COPA, LDLR, | 1286 | 466 | 13588 | 1.26974549 | 1 | 0.44134335 | 60.3453143 |
| GO:0009628~response to abiotic stimulus | 33 | 0.1574 | 0.04990895 | RAB3A, ARPP | 1286 | 251 | 13588 | 1.38916806 | 1 | 0.44138943 | 60.4812924 |
| GO:0008202~steroid metabolic process | 23 | 0.1097 | 0.04998338 | SREBF1, CYP | 1286 | 161 | 13588 | 1.50944235 | 1 | 0.44073158 | 60.5373981 |
| GO:0008299~isoprenoid biosynthetic process | 6 | 0.0286 | 0.05074452 | ALDH1A1, AL | 1286 | 22 | 13588 | 2.88166266 | 1 | 0.44463181 | 61.1068109 |
| GO:0019218~regulation of steroid metabolic process | 6 | 0.0286 | 0.05074452 | SREBF1, INSI | 1286 | 22 | 13588 | 2.88166266 | 1 | 0.44463181 | 61.1068109 |
| GO:0006812~cation transport | 61 | 0.2909 | 0.05228043 | KCNC2, SLC3 | 1286 | 515 | 13588 | 1.25151822 | 1 | 0.45353015 | 62.2322704 |
| GO:0010605~negative regulation of macromolecule metabolic pro | 60 | 0.2862 | 0.05264016 | THRA, THRB, | 1286 | 506 | 13588 | 1.25289681 | 1 | 0.45469677 | 62.491385 |
| GO:0032268~regulation of cellular protein metabolic process | 36 | 0.1717 | 0.05310082 | HMGB1, TGF | 1286 | 280 | 13588 | 1.35849811 | 1 | 0.45650139 | 62.8207496 |
| GO:0009968~negative regulation of signal transduction | 24 | 0.1145 | 0.05324082 | CER1, ONECU | 1286 | 171 | 13588 | 1.4829609 | 1 | 0.45624156 | 62.9203012 |
| GO:0009201~ribonucleoside triphosphate biosynthetic process | 15 | 0.0715 | 0.05395772 | TCIRG1, ATP | 1286 | 93 | 13588 | 1.7042091 | 1 | 0.45965147 | 63.4261578 |
| GO:0009206~purine ribonucleoside triphosphate biosynthetic proc | 15 | 0.0715 | 0.05395772 | TCIRG1, ATP | 1286 | 93 | 13588 | 1.7042091 | 1 | 0.45965147 | 63.4261578 |
| GO:0006397~mRNA processing | 34 | 0.1622 | 0.05411093 | SCAF1, RNAS | 1286 | 262 | 13588 | 1.37117282 | 1 | 0.45946838 | 63.5334161 |
| GO:0060537~muscle tissue development | 20 | 0.0954 | 0.05416036 | RXRA, MET, | 1286 | 136 | 13588 | 1.55383771 | 1 | 0.45863237 | 63.5679582 |
| GO:0043009~chordate embryonic development | 51 | 0.2432 | 0.05458082 | GNA13, COB | 1286 | 421 | 13588 | 1.27997843 | 1 | 0.46013232 | 63.8605207 |
| GO:0048878~chemical homeostasis | 45 | 0.2146 | 0.05525629 | LDLR, GNA11 | 1286 | 365 | 13588 | 1.30266942 | 1 | 0.4632071 | 64.3258844 |
| GO:0034622~cellular macromolecular complex assembly | 29 | 0.1383 | 0.05597844 | CALY, HP1BP | 1286 | 217 | 13588 | 1.41205897 | 1 | 0.46653436 | 64.8171454 |
| GO:0022613~ribonucleoprotein complex biogenesis | 20 | 0.0954 | 0.0575546 | KRR1, GTPBP | 1286 | 137 | 13588 | 1.54249583 | 1 | 0.47501652 | 65.8672485 |
| GO:0030307~positive regulation of cell growth | 5 | 0.0238 | 0.05763726 | BCL2, CD81, | 1286 | 16 | 13588 | 3.30190513 | 1 | 0.47436827 | 65.9214992 |
| GO:0009145~purine nucleoside triphosphate biosynthetic process | 15 | 0.0715 | 0.05813139 | TCIRG1, ATP | 1286 | 94 | 13588 | 1.68607922 | 1 | 0.47619967 | 66.2440806 |
| GO:0006695~cholesterol biosynthetic process | 6 | 0.0286 | 0.05998416 | CYP51, INSIG | 1286 | 23 | 13588 | 2.75637298 | 1 | 0.48606388 | 67.4281067 |
| GO:0035023~regulation of Rho protein signal transduction | 14 | 0.0668 | 0.06006799 | ARHGEF4, AF | 1286 | 86 | 13588 | 1.72006221 | 1 | 0.48540704 | 67.480742 |
| GO:0016486~peptide hormone processing | 4 | 0.0191 | 0.06108398 | PCSK2, ECE1, | 1286 | 10 | 13588 | 4.22643857 | 1 | 0.49019001 | 68.1122821 |
| GO:0008105~asymmetric protein localization | 4 | 0.0191 | 0.06108398 | HCN1, VANG | 1286 | 10 | 13588 | 4.22643857 | 1 | 0.49019001 | 68.1122821 |
| GO:0006884~cell volume homeostasis | 4 | 0.0191 | 0.06108398 | SLC12A7, SL | 1286 | 10 | 13588 | 4.22643857 | 1 | 0.49019001 | 68.1122821 |
| GO:0021756~striatum development | 4 | 0.0191 | 0.06108398 | BBS1, ALDH1 | 1286 | 10 | 13588 | 4.22643857 | 1 | 0.49019001 | 68.1122821 |
| GO:0031575~G1/S transition checkpoint | 4 | 0.0191 | 0.06108398 | CCND1, FBX | 1286 | 10 | 13588 | 4.22643857 | 1 | 0.49019001 | 68.1122821 |
| GO:0009792~embryonic development ending in birth or egg hatch | 51 | 0.2432 | 0.061581 | GNA13, COB | 1286 | 425 | 13588 | 1.26793157 | 1 | 0.49191798 | 68.416994 |
| GO:0009100~glycoprotein metabolic process | 21 | 0.1002 | 0.06169407 | ST6GAL1, IM | 1286 | 147 | 13588 | 1.50944235 | 1 | 0.49142482 | 68.4859275 |
| GO:0031401~positive regulation of protein modification process | 13 | 0.062 | 0.06184059 | MEN1, LIF, Z | 1286 | 78 | 13588 | 1.76101607 | 1 | 0.49112663 | 68.5750477 |
| GO:0016331~morphogenesis of embryonic epithelium | 13 | 0.062 | 0.06184059 | KAT2A, COBL | 1286 | 78 | 13588 | 1.76101607 | 1 | 0.49112663 | 68.5750477 |
| GO:0009142~nucleoside triphosphate biosynthetic process | 15 | 0.0715 | 0.0625187 | TCIRG1, ATP | 1286 | 95 | 13588 | 1.66833101 | 1 | 0.49386578 | 68.9843997 |
| GO:0055074~calcium ion homeostasis | 15 | 0.0715 | 0.0625187 | CCKBR, PIK3 | 1286 | 95 | 13588 | 1.66833101 | 1 | 0.49386578 | 68.9843997 |
| GO:0043434~response to peptide hormone stimulus | 15 | 0.0715 | 0.0625187 | WDTC1, IRS2 | 1286 | 95 | 13588 | 1.66833101 | 1 | 0.49386578 | 68.9843997 |
| GO:0043583~ear development | 16 | 0.0763 | 0.06446203 | FGFR1, MYO | 1286 | 104 | 13588 | 1.6255533 | 1 | 0.50367733 | 70.1297918 |
| GO:0007423~sensory organ development | 33 | 0.1574 | 0.06482579 | FGFR1, HMG | 1286 | 257 | 13588 | 1.35673612 | 1 | 0.50455916 | 70.3396981 |
| GO:0055002~striated muscle cell development | 10 | 0.0477 | 0.06495439 | ACTG1, GPX1 | 1286 | 54 | 13588 | 1.95668452 | 1 | 0.50413354 | 70.4135766 |
| GO:0007219~Notch signaling pathway | 10 | 0.0477 | 0.06495439 | DTX4, NCSTN | 1286 | 54 | 13588 | 1.95668452 | 1 | 0.50413354 | 70.4135766 |

| | | | | | | | | | | | |
|---|---|---|---|---|---|---|---|---|---|---|---|
| GO:0046903~secretion | 29 | 0.1383 | 0.06735796 | SYT1, RAB3A | 1286 | 221 | 13588 | 1.38650134 | 1 | 0.51613949 | 71.76274 |
| GO:0043062~extracellular structure organization | 21 | 0.1002 | 0.06901426 | COL18A1, RE | 1286 | 149 | 13588 | 1.48918138 | 1 | 0.52386127 | 72.6583845 |
| GO:0007517~muscle organ development | 24 | 0.1145 | 0.0691986 | RXRA, MET, I | 1286 | 176 | 13588 | 1.44083133 | 1 | 0.52369303 | 72.7563888 |
| GO:0006650~glycerophospholipid metabolic process | 14 | 0.0668 | 0.06975484 | PLD1, PIK3CE | 1286 | 88 | 13588 | 1.68096989 | 1 | 0.52548546 | 73.0501095 |
| GO:0006916~anti-apoptosis | 14 | 0.0668 | 0.06975484 | EEF1A2, CLU, | 1286 | 88 | 13588 | 1.68096989 | 1 | 0.52548546 | 73.0501095 |
| GO:0010740~positive regulation of protein kinase cascade | 14 | 0.0668 | 0.06975484 | FGFR1, PDCD | 1286 | 88 | 13588 | 1.68096989 | 1 | 0.52548546 | 73.0501095 |
| GO:0006325~chromatin organization | 39 | 0.186 | 0.06990604 | HP1BP3, UTY | 1286 | 315 | 13588 | 1.30818337 | 1 | 0.52514057 | 73.1294295 |
| GO:0050982~detection of mechanical stimulus | 5 | 0.0238 | 0.06993611 | ATP2B2, SLC | 1286 | 17 | 13588 | 3.10767542 | 1 | 0.52416335 | 73.1451775 |
| GO:0007093~mitotic cell cycle checkpoint | 6 | 0.0286 | 0.07011357 | CCND1, MAD | 1286 | 24 | 13588 | 2.64152411 | 1 | 0.52396141 | 73.2379441 |
| GO:0001569~patterning of blood vessels | 6 | 0.0286 | 0.07011357 | SEMA5A, GN | 1286 | 24 | 13588 | 2.64152411 | 1 | 0.52396141 | 73.2379441 |
| GO:0042542~response to hydrogen peroxide | 6 | 0.0286 | 0.07011357 | GPX1, GPX4, | 1286 | 24 | 13588 | 2.64152411 | 1 | 0.52396141 | 73.2379441 |
| GO:0001843~neural tube closure | 8 | 0.0382 | 0.07040495 | KAT2A, COBL | 1286 | 39 | 13588 | 2.16740439 | 1 | 0.52435446 | 73.3896034 |
| GO:0060606~tube closure | 8 | 0.0382 | 0.07040495 | KAT2A, COBL | 1286 | 39 | 13588 | 2.16740439 | 1 | 0.52435446 | 73.3896034 |
| GO:0008584~male gonad development | 8 | 0.0382 | 0.07040495 | WNT5A, HM | 1286 | 39 | 13588 | 2.16740439 | 1 | 0.52435446 | 73.3896034 |
| GO:0030384~phosphoinositide metabolic process | 11 | 0.0525 | 0.07043779 | PIK3CB, PIGV | 1286 | 63 | 13588 | 1.84487398 | 1 | 0.52340196 | 73.4066457 |
| GO:0007005~mitochondrion organization | 15 | 0.0715 | 0.07194631 | CEBPA, SEPT | 1286 | 97 | 13588 | 1.63393244 | 1 | 0.53006604 | 74.1784425 |
| GO:0035058~sensory cilium assembly | 3 | 0.0143 | 0.07365477 | BBS1, VANGL | 1286 | 5 | 13588 | 6.33965785 | 1 | 0.53761531 | 75.0269673 |
| GO:0007216~metabotropic glutamate receptor signaling pathway | 3 | 0.0143 | 0.07365477 | HOMER3, HC | 1286 | 5 | 13588 | 6.33965785 | 1 | 0.53761531 | 75.0269673 |
| GO:0032414~positive regulation of ion transmembrane transporte | 3 | 0.0143 | 0.07365477 | PLCG2, STIM | 1286 | 5 | 13588 | 6.33965785 | 1 | 0.53761531 | 75.0269673 |
| GO:0060751~mammary gland duct branch elongation | 3 | 0.0143 | 0.07365477 | WNT5A, ESR | 1286 | 5 | 13588 | 6.33965785 | 1 | 0.53761531 | 75.0269673 |
| GO:0045649~regulation of macrophage differentiation | 3 | 0.0143 | 0.07365477 | LIF, CSF1, RB | 1286 | 5 | 13588 | 6.33965785 | 1 | 0.53761531 | 75.0269673 |
| GO:0048087~positive regulation of pigmentation during developm | 3 | 0.0143 | 0.07365477 | ADAMTS9, B | 1286 | 5 | 13588 | 6.33965785 | 1 | 0.53761531 | 75.0269673 |
| GO:0051271~negative regulation of cell motion | 7 | 0.0334 | 0.07618206 | GTPBP4, ACT | 1286 | 32 | 13588 | 2.31133359 | 1 | 0.54904582 | 76.2339978 |
| GO:0017157~regulation of exocytosis | 7 | 0.0334 | 0.07618206 | SEPT5, RAB3 | 1286 | 32 | 13588 | 2.31133359 | 1 | 0.54904582 | 76.2339978 |
| GO:0001666~response to hypoxia | 11 | 0.0525 | 0.07675506 | ECE1, EPAS1, | 1286 | 64 | 13588 | 1.81604782 | 1 | 0.55072006 | 76.4998877 |
| GO:0009416~response to light stimulus | 15 | 0.0715 | 0.07699158 | HMGCR, USP | 1286 | 98 | 13588 | 1.61725966 | 1 | 0.55074642 | 76.6088213 |
| GO:0034613~cellular protein localization | 37 | 0.1765 | 0.0776833 | STON2, PACS | 1286 | 299 | 13588 | 1.30751026 | 1 | 0.55297434 | 76.9246644 |
| GO:0008088~axon cargo transport | 4 | 0.0191 | 0.07826521 | APP, KIF1B, U | 1286 | 11 | 13588 | 3.84221688 | 1 | 0.55465369 | 77.1872473 |
| GO:0045136~development of secondary sexual characteristics | 4 | 0.0191 | 0.07826521 | WNT5A, NCC | 1286 | 11 | 13588 | 3.84221688 | 1 | 0.55465369 | 77.1872473 |
| GO:0008354~germ cell migration | 4 | 0.0191 | 0.07826521 | CXCR4, KIT, K | 1286 | 11 | 13588 | 3.84221688 | 1 | 0.55465369 | 77.1872473 |
| GO:0006268~DNA unwinding during replication | 4 | 0.0191 | 0.07826521 | TOP1, TOP3A | 1286 | 11 | 13588 | 3.84221688 | 1 | 0.55465369 | 77.1872473 |
| GO:0001934~positive regulation of protein amino acid phosphoryla | 10 | 0.0477 | 0.07834813 | LIF, ZFP91, H | 1286 | 56 | 13588 | 1.88680293 | 1 | 0.55393423 | 77.2244324 |
| GO:0007015~actin filament organization | 10 | 0.0477 | 0.07834813 | ACTN4, GSN, | 1286 | 56 | 13588 | 1.88680293 | 1 | 0.55393423 | 77.2244324 |
| GO:0009259~ribonucleotide metabolic process | 18 | 0.0858 | 0.08036209 | TCIRG1, ATP | 1286 | 125 | 13588 | 1.52151788 | 1 | 0.56236751 | 78.1102186 |
| GO:0002064~epithelial cell development | 6 | 0.0286 | 0.0811218 | COL18A1, GF | 1286 | 25 | 13588 | 2.53586314 | 1 | 0.56480458 | 78.435818 |
| GO:0060078~regulation of postsynaptic membrane potential | 6 | 0.0286 | 0.0811218 | SNCA, GRIK5 | 1286 | 25 | 13588 | 2.53586314 | 1 | 0.56480458 | 78.435818 |
| GO:0045639~positive regulation of myeloid cell differentiation | 6 | 0.0286 | 0.0811218 | LIF, HMGB1, | 1286 | 25 | 13588 | 2.53586314 | 1 | 0.56480458 | 78.435818 |
| GO:0048514~blood vessel morphogenesis | 26 | 0.124 | 0.08278187 | GNA13, FGFF | 1286 | 198 | 13588 | 1.38746721 | 1 | 0.57135905 | 79.1314462 |
| GO:0070482~response to oxygen levels | 11 | 0.0525 | 0.08341303 | ECE1, EPAS1, | 1286 | 65 | 13588 | 1.78810863 | 1 | 0.57312879 | 79.3903212 |
| GO:0060445~branching involved in salivary gland morphogenesis | 5 | 0.0238 | 0.08346073 | FGFR1, NRP1 | 1286 | 18 | 13588 | 2.93502678 | 1 | 0.57223642 | 79.4097602 |
| GO:0006471~protein amino acid ADP-ribosylation | 5 | 0.0238 | 0.08346073 | GNA14, GNA | 1286 | 18 | 13588 | 2.93502678 | 1 | 0.57223642 | 79.4097602 |
| GO:0070727~cellular macromolecule localization | 37 | 0.1765 | 0.08360409 | STON2, PACS | 1286 | 301 | 13588 | 1.29882248 | 1 | 0.57178211 | 79.4680843 |
| GO:0045944~positive regulation of transcription from RNA polyme | 43 | 0.2051 | 0.08364595 | ABLIM1, HM | 1286 | 358 | 13588 | 1.26911214 | 1 | 0.57087003 | 79.4850846 |
| GO:0006754~ATP biosynthetic process | 13 | 0.062 | 0.08383606 | TCIRG1, ATP | 1286 | 82 | 13588 | 1.67511285 | 1 | 0.57063289 | 79.5621277 |
| GO:0051235~maintenance of location | 7 | 0.0334 | 0.08602227 | TLN1, FGFR1 | 1286 | 33 | 13588 | 2.24129318 | 1 | 0.57932526 | 80.4286594 |
| GO:0060443~mammary gland morphogenesis | 7 | 0.0334 | 0.08602227 | WNT5A, DDF | 1286 | 33 | 13588 | 2.24129318 | 1 | 0.57932526 | 80.4286594 |
| GO:0006874~cellular calcium ion homeostasis | 14 | 0.0668 | 0.08610539 | CCKBR, PIK3 | 1286 | 91 | 13588 | 1.6255533 | 1 | 0.57859793 | 80.4609073 |
| GO:0006644~phospholipid metabolic process | 22 | 0.1049 | 0.0894064 | PLD1, PIK3CE | 1286 | 163 | 13588 | 1.4260989 | 1 | 0.5918849 | 81.701858 |
| GO:0006396~RNA processing | 51 | 0.2432 | 0.08948891 | SCAF1, RNAS | 1286 | 437 | 13588 | 1.23311423 | 1 | 0.59114525 | 81.7319013 |
| GO:0060627~regulation of vesicle-mediated transport | 13 | 0.062 | 0.09001115 | STON2, SEPT | 1286 | 83 | 13588 | 1.65493077 | 1 | 0.59228657 | 81.9209753 |
| GO:0006457~protein folding | 18 | 0.0858 | 0.09005881 | FKBP8, TTC9 | 1286 | 127 | 13588 | 1.49755697 | 1 | 0.59140269 | 81.9381385 |
| GO:0050673~epithelial cell proliferation | 6 | 0.0286 | 0.09299006 | WNT5A, CCN | 1286 | 26 | 13588 | 2.43832994 | 1 | 0.60262953 | 82.9646019 |
| GO:0051495~positive regulation of cytoskeleton organization | 6 | 0.0286 | 0.09299006 | CFL1, PSRC1, | 1286 | 26 | 13588 | 2.43832994 | 1 | 0.60262953 | 82.9646019 |
| GO:0048610~reproductive cellular process | 23 | 0.1097 | 0.09315483 | SEPT4, DNM | 1286 | 173 | 13588 | 1.40474114 | 1 | 0.60222817 | 83.0206331 |
| GO:0001656~metanephros development | 10 | 0.0477 | 0.09330215 | FGFR1, BDNF | 1286 | 58 | 13588 | 1.82174076 | 1 | 0.60175685 | 83.0705836 |
| GO:0032870~cellular response to hormone stimulus | 10 | 0.0477 | 0.09330215 | WDTC1, IRS2 | 1286 | 58 | 13588 | 1.82174076 | 1 | 0.60175685 | 83.0705836 |
| GO:0007281~germ cell development | 15 | 0.0715 | 0.0934739 | SEPT4, HMG | 1286 | 101 | 13588 | 1.56922224 | 1 | 0.60138852 | 83.1286401 |
| GO:0006323~DNA packaging | 15 | 0.0715 | 0.0934739 | HIST1H2BC, | 1286 | 101 | 13588 | 1.56922224 | 1 | 0.60138852 | 83.1286401 |
| GO:0009205~purine ribonucleoside triphosphate metabolic proces | 15 | 0.0715 | 0.0934739 | TCIRG1, ATP | 1286 | 101 | 13588 | 1.56922224 | 1 | 0.60138852 | 83.1286401 |
| GO:0009150~purine ribonucleotide metabolic process | 17 | 0.0811 | 0.09495427 | TCIRG1, ATP | 1286 | 119 | 13588 | 1.50944235 | 1 | 0.60637311 | 83.6213166 |
| GO:0042391~regulation of membrane potential | 17 | 0.0811 | 0.09495427 | SCD2, GNA1 | 1286 | 119 | 13588 | 1.50944235 | 1 | 0.60637311 | 83.6213166 |
| GO:0050885~neuromuscular process controlling balance | 7 | 0.0334 | 0.09652538 | ATP2B2, APP | 1286 | 34 | 13588 | 2.17537279 | 1 | 0.61164009 | 84.1293045 |

| | | | | | | | | | | | |
|---|---|---|---|---|---|---|---|---|---|---|---|
| GO:0014902~myotube differentiation | 4 | 0.0191 | 0.09728165 | GPX1, MET, I | 1286 | 12 | 13588 | 3.52203214 | 1 | 0.61358891 | 84.368487 |
| GO:0051928~positive regulation of calcium ion transport | 4 | 0.0191 | 0.09728165 | ATP2B2, PLC | 1286 | 12 | 13588 | 3.52203214 | 1 | 0.61358891 | 84.368487 |
| GO:0048588~developmental cell growth | 4 | 0.0191 | 0.09728165 | APP, ULK1, D | 1286 | 12 | 13588 | 3.52203214 | 1 | 0.61358891 | 84.368487 |
| GO:0032508~DNA duplex unwinding | 4 | 0.0191 | 0.09728165 | TOP1, TOP3A | 1286 | 12 | 13588 | 3.52203214 | 1 | 0.61358891 | 84.368487 |
| GO:0048488~synaptic vesicle endocytosis | 4 | 0.0191 | 0.09728165 | RAB3A, SNCA | 1286 | 12 | 13588 | 3.52203214 | 1 | 0.61358891 | 84.368487 |
| GO:0051647~nucleus localization | 4 | 0.0191 | 0.09728165 | CDC42BPA, S | 1286 | 12 | 13588 | 3.52203214 | 1 | 0.61358891 | 84.368487 |
| GO:0009165~nucleotide biosynthetic process | 23 | 0.1097 | 0.0975971 | TCIRG1, ADS | 1286 | 174 | 13588 | 1.39666792 | 1 | 0.61377497 | 84.467243 |
| GO:0042472~inner ear morphogenesis | 11 | 0.0525 | 0.09775277 | FGFR1, SPRY | 1286 | 67 | 13588 | 1.73473225 | 1 | 0.61332827 | 84.5157611 |
| GO:0030902~hindbrain development | 11 | 0.0525 | 0.09775277 | KAT2A, ALDH | 1286 | 67 | 13588 | 1.73473225 | 1 | 0.61332827 | 84.5157611 |
| GO:0019217~regulation of fatty acid metabolic process | 5 | 0.0238 | 0.09815822 | SREBF1, WD | 1286 | 19 | 13588 | 2.78055169 | 1 | 0.61386982 | 84.6414532 |
| GO:0031399~regulation of protein modification process | 22 | 0.1049 | 0.09845468 | HMGB1, GTP | 1286 | 165 | 13588 | 1.40881286 | 1 | 0.61397991 | 84.7327504 |
| GO:0009725~response to hormone stimulus | 22 | 0.1049 | 0.09845468 | GPR83, HMG | 1286 | 165 | 13588 | 1.40881286 | 1 | 0.61397991 | 84.7327504 |
| GO:0009199~ribonucleoside triphosphate metabolic process | 15 | 0.0715 | 0.09941987 | TCIRG1, ATP | 1286 | 102 | 13588 | 1.55383771 | 1 | 0.61670401 | 85.0264425 |
| GO:0055082~cellular chemical homeostasis | 33 | 0.1574 | 0.0998837 | GNA11, CLST | 1286 | 268 | 13588 | 1.30104919 | 1 | 0.61745622 | 85.165673 |
