## Supplementary Data 3 for "FGFR1 is critical for *Rbl2* loss-driven tumor development and requires PLCG1 activation for continued growth of small cell lung cancer": Supplementaty Data 3 .pdf

| Term | Count | % | PValue | Genes | List Total | Pop Hits | Pop Total | Fold Enrichm | Bonferroni | Benjamini | FDR |
|---|---|---|---|---|---|---|---|---|---|---|---|
| GO:0007409~axonogenesis | 12 | 0.6006 | 3.31E-06 | DCC, ARX, BD | 160 | 163 | 13588 | 6.25214724 | 0.00390677 | 0.00390677 | 0.0053358 |
| GO:0048729~tissue morphogenesis | 14 | 0.7007 | 4.42E-06 | WNT5A, ESR | 160 | 238 | 13588 | 4.99558824 | 0.00521568 | 0.00261125 | 0.0071281 |
| GO:0048812~neuron projection morphogenesis | 12 | 0.6006 | 6.93E-06 | DCC, ARX, BD | 160 | 176 | 13588 | 5.79034091 | 0.0081699 | 0.00273075 | 0.01118191 |
| GO:0048666~neuron development | 15 | 0.75075 | 8.45E-06 | DCC, FGFR1, | 160 | 292 | 13588 | 4.36258562 | 0.00995969 | 0.00249928 | 0.01364367 |
| GO:0060688~regulation of morphogenesis of a branching structure | 6 | 0.3003 | 9.06E-06 | WNT5A, FGF | 160 | 25 | 13588 | 20.382 | 0.01066674 | 0.00214251 | 0.01461738 |
| GO:0048667~cell morphogenesis involved in neuron differentiation | 12 | 0.6006 | 9.54E-06 | DCC, ARX, BD | 160 | 182 | 13588 | 5.59945055 | 0.01123268 | 0.00188094 | 0.01539726 |
| GO:0031175~neuron projection development | 13 | 0.65065 | 9.65E-06 | DCC, FGFR1, | 160 | 218 | 13588 | 5.06433486 | 0.01136333 | 0.00163129 | 0.01557738 |
| GO:0060429~epithelium development | 14 | 0.7007 | 1.79E-05 | WNT5A, ESR | 160 | 271 | 13588 | 4.38726937 | 0.02093136 | 0.0026407 | 0.02883124 |
| GO:0030182~neuron differentiation | 17 | 0.85085 | 1.79E-05 | DCC, FGFR1, | 160 | 399 | 13588 | 3.6183584 | 0.02095144 | 0.0023499 | 0.02885918 |
| GO:0048858~cell projection morphogenesis | 12 | 0.6006 | 2.54E-05 | DCC, ARX, BD | 160 | 202 | 13588 | 5.0450495 | 0.02957111 | 0.00299721 | 0.04090946 |
| GO:0000904~cell morphogenesis involved in differentiation | 12 | 0.6006 | 3.95E-05 | DCC, ARX, BD | 160 | 212 | 13588 | 4.80707547 | 0.04573999 | 0.00424724 | 0.06380101 |
| GO:0032990~cell part morphogenesis | 12 | 0.6006 | 3.95E-05 | DCC, ARX, BD | 160 | 212 | 13588 | 4.80707547 | 0.04573999 | 0.00424724 | 0.06380101 |
| GO:0006928~cell motion | 15 | 0.75075 | 1.04E-04 | DCC, SATB2, | 160 | 367 | 13588 | 3.47104905 | 0.11604928 | 0.01022684 | 0.16800846 |
| GO:0048732~gland development | 11 | 0.55055 | 1.07E-04 | WNT5A, FGF | 160 | 197 | 13588 | 4.74200508 | 0.11923787 | 0.00971921 | 0.17292611 |
| GO:0001763~morphogenesis of a branching structure | 9 | 0.45045 | 1.09E-04 | WNT5A, FGF | 160 | 125 | 13588 | 6.1146 | 0.12093707 | 0.0091648 | 0.1755539 |
| GO:0021953~central nervous system neuron differentiation | 6 | 0.3003 | 1.24E-04 | DCC, ARX, SC | 160 | 42 | 13588 | 12.1321429 | 0.13607301 | 0.00970375 | 0.19918501 |
| GO:0001501~skeletal system development | 13 | 0.65065 | 1.32E-04 | WNT5A, FGF | 160 | 285 | 13588 | 3.87377193 | 0.14489522 | 0.0097355 | 0.21314786 |
| GO:0001764~neuron migration | 7 | 0.35035 | 1.65E-04 | DCC, ARX, SA | 160 | 70 | 13588 | 8.4925 | 0.17756008 | 0.01143296 | 0.2661133 |
| GO:0021952~central nervous system projection neuron axonogenes | 4 | 0.2002 | 2.42E-04 | DCC, DCX, NI | 160 | 11 | 13588 | 30.8818182 | 0.2492862 | 0.01580329 | 0.39009375 |
| GO:0000902~cell morphogenesis | 13 | 0.65065 | 2.79E-04 | DCC, NR2E1, | 160 | 309 | 13588 | 3.57289644 | 0.28162632 | 0.01725805 | 0.44986736 |
| GO:0051216~cartilage development | 7 | 0.35035 | 2.99E-04 | WNT5A, FGF | 160 | 78 | 13588 | 7.62147436 | 0.29858059 | 0.01757617 | 0.48227287 |
| GO:0007155~cell adhesion | 18 | 0.9009 | 3.08E-04 | PCDHA2, NEI | 160 | 561 | 13588 | 2.72486631 | 0.30529543 | 0.01719655 | 0.49532136 |
| GO:0022610~biological adhesion | 18 | 0.9009 | 3.14E-04 | PCDHA2, NEI | 160 | 562 | 13588 | 2.72001779 | 0.31055406 | 0.01676099 | 0.50562724 |
| GO:0051960~regulation of nervous system development | 9 | 0.45045 | 3.48E-04 | BDNF, SOX5, | 160 | 148 | 13588 | 5.16435811 | 0.3381347 | 0.01778316 | 0.56098265 |
| GO:0030030~cell projection organization | 13 | 0.65065 | 3.73E-04 | DCC, FGFR1, | 160 | 319 | 13588 | 3.46089342 | 0.35696597 | 0.01823002 | 0.60010072 |
| GO:0007156~homophilic cell adhesion | 8 | 0.4004 | 4.45E-04 | PCDHA2, FRE | 160 | 117 | 13588 | 5.80683761 | 0.40948237 | 0.02084981 | 0.71547449 |
| GO:0022612~gland morphogenesis | 7 | 0.35035 | 4.47E-04 | WNT5A, FGF | 160 | 84 | 13588 | 7.07708333 | 0.41134136 | 0.02017482 | 0.7197417 |
| GO:0016477~cell migration | 11 | 0.55055 | 5.25E-04 | DCC, ARX, GI | 160 | 240 | 13588 | 3.89239583 | 0.462908 | 0.02275872 | 0.84373417 |
| GO:0060284~regulation of cell development | 9 | 0.45045 | 5.61E-04 | BDNF, SOX5, | 160 | 159 | 13588 | 4.80707547 | 0.48572436 | 0.02347005 | 0.90239182 |
| GO:0021954~central nervous system neuron development | 5 | 0.25025 | 5.67E-04 | DCC, ARX, DC | 160 | 33 | 13588 | 12.8674242 | 0.48910084 | 0.02289195 | 0.91128965 |
| GO:0007267~cell-cell signaling | 12 | 0.6006 | 6.11E-04 | WNT5A, FGF | 160 | 290 | 13588 | 3.51413793 | 0.51515512 | 0.02384205 | 0.98196582 |
| GO:0021955~central nervous system neuron axonogenesis | 4 | 0.2002 | 6.45E-04 | DCC, DCX, NI | 160 | 15 | 13588 | 22.6466667 | 0.53423757 | 0.02434645 | 1.03614841 |
| GO:0048754~branching morphogenesis of a tube | 7 | 0.35035 | 7.70E-04 | WNT5A, ESR | 160 | 93 | 13588 | 6.3922043 | 0.5981935 | 0.02809117 | 1.23520564 |
| GO:0030900~forebrain development | 9 | 0.45045 | 7.75E-04 | ARX, WNT5A | 160 | 167 | 13588 | 4.57679641 | 0.600745 | 0.02743939 | 1.24378171 |
| GO:0032989~cellular component morphogenesis | 13 | 0.65065 | 8.69E-04 | DCC, NR2E1, | 160 | 351 | 13588 | 3.14537037 | 0.64270741 | 0.02981703 | 1.39315707 |
| GO:0050767~regulation of neurogenesis | 8 | 0.4004 | 9.14E-04 | BDNF, SOX5, | 160 | 132 | 13588 | 5.1469697 | 0.66117498 | 0.03044885 | 1.46446737 |
| GO:0051094~positive regulation of developmental process | 10 | 0.5005 | 9.33E-04 | LIF, FGFR1, L | 160 | 214 | 13588 | 3.96845794 | 0.66888976 | 0.03023634 | 1.49539935 |
| GO:0002009~morphogenesis of an epithelium | 9 | 0.45045 | 9.75E-04 | WNT5A, FREI | 160 | 173 | 13588 | 4.41806358 | 0.68500025 | 0.03073883 | 1.56235299 |
| GO:0007411~axon guidance | 7 | 0.35035 | 0.0010134 | DCC, ARX, BD | 160 | 98 | 13588 | 6.06607143 | 0.69894759 | 0.03109755 | 1.62310329 |
| GO:0045597~positive regulation of cell differentiation | 9 | 0.45045 | 0.00105106 | LIF, LPL, ADA | 160 | 175 | 13588 | 4.36757143 | 0.71209224 | 0.03142177 | 1.682954 |
| GO:0035295~tube development | 11 | 0.55055 | 0.00109138 | WNT5A, FGF | 160 | 264 | 13588 | 3.53854167 | 0.72552763 | 0.03180582 | 1.74698104 |
| GO:0060445~branching involved in salivary gland morphogenesis | 4 | 0.2002 | 0.00112754 | FGFR1, DAG1 | 160 | 18 | 13588 | 18.8722222 | 0.73704325 | 0.03205465 | 1.80436953 |
| GO:0045664~regulation of neuron differentiation | 7 | 0.35035 | 0.00124775 | BDNF, SOX5, | 160 | 102 | 13588 | 5.82818627 | 0.7719666 | 0.03458454 | 1.99492469 |
| GO:0060762~regulation of branching involved in mammary gland du | 3 | 0.15015 | 0.00132956 | WNT5A, ETV | 160 | 5 | 13588 | 50.955 | 0.79304356 | 0.03597076 | 2.12440499 |
| GO:0016337~cell-cell adhesion | 10 | 0.5005 | 0.0018413 | PCDHA2, FRE | 160 | 236 | 13588 | 3.59851695 | 0.88719634 | 0.04838366 | 2.93073093 |
| GO:0048870~cell motility | 11 | 0.55055 | 0.00188045 | DCC, ARX, GI | 160 | 284 | 13588 | 3.28934859 | 0.89231599 | 0.04831714 | 2.99217098 |
| GO:0051674~localization of cell | 11 | 0.55055 | 0.00188045 | DCC, ARX, GI | 160 | 284 | 13588 | 3.28934859 | 0.89231599 | 0.04831714 | 2.99217098 |
| GO:0048568~embryonic organ development | 10 | 0.5005 | 0.0021236 | LIF, FGFR1, S | 160 | 241 | 13588 | 3.52385892 | 0.91930035 | 0.05324776 | 3.37287916 |
| GO:0010627~regulation of protein kinase cascade | 8 | 0.4004 | 0.00230718 | LIF, FGFR1, G | 160 | 155 | 13588 | 4.38322581 | 0.93509684 | 0.0565279 | 3.65938332 |
| GO:0043523~regulation of neuron apoptosis | 6 | 0.3003 | 0.00244491 | GPX1, BDNF, | 160 | 80 | 13588 | 6.369375 | 0.944884 | 0.05859474 | 3.87380585 |
| GO:0048562~embryonic organ morphogenesis | 8 | 0.4004 | 0.00285084 | FGFR1, SATB | 160 | 161 | 13588 | 4.21987578 | 0.96595992 | 0.0666584 | 4.50319213 |
| GO:0030879~mammary gland development | 6 | 0.3003 | 0.00334926 | WNT5A, CCN | 160 | 86 | 13588 | 5.925 | 0.9811676 | 0.07636983 | 5.27068333 |
| GO:0021543~pallium development | 5 | 0.25025 | 0.00338603 | ARX, ZEB2, TI | 160 | 53 | 13588 | 8.01179245 | 0.98197269 | 0.07572211 | 5.32708466 |
| GO:0007435~salivary gland morphogenesis | 4 | 0.2002 | 0.003742 | FGFR1, DAG1 | 160 | 27 | 13588 | 12.5814815 | 0.98819036 | 0.08182042 | 5.87137218 |
| GO:0035239~tube morphogenesis | 8 | 0.4004 | 0.00398705 | WNT5A, ESR | 160 | 171 | 13588 | 3.97309942 | 0.99117439 | 0.08538048 | 6.24434929 |
| GO:0042127~regulation of cell proliferation | 15 | 0.75075 | 0.00428571 | WNT5A, FGF | 160 | 538 | 13588 | 2.3677974 | 0.99381227 | 0.08987209 | 6.69706675 |
| GO:0033598~mammary gland epithelial cell proliferation | 3 | 0.15015 | 0.00464121 | WNT5A, CCN | 160 | 9 | 13588 | 28.3083333 | 0.99594573 | 0.09529394 | 7.23325931 |
| GO:0043408~regulation of MAPKKK cascade | 6 | 0.3003 | 0.00468295 | LIF, FGFR1, E | 160 | 93 | 13588 | 5.47903226 | 0.99614212 | 0.09447784 | 7.29602571 |
| GO:0007431~salivary gland development | 4 | 0.2002 | 0.00555761 | FGFR1, DAG1 | 160 | 31 | 13588 | 10.9580645 | 0.99863767 | 0.10931479 | 8.60214362 |
| GO:0048589~developmental growth | 6 | 0.3003 | 0.00635694 | WNT5A, GPX | 160 | 100 | 13588 | 5.0955 | 0.99947423 | 0.12206569 | 9.7806689 |
| GO:0048704~embryonic skeletal system morphogenesis | 5 | 0.25025 | 0.0066547 | SATB2, HOXE | 160 | 64 | 13588 | 6.63476562 | 0.99963129 | 0.12540246 | 10.2160212 |

| | | | | | | | | | | | |
|---|---|---|---|---|---|---|---|---|---|---|---|
| GO:0055123~digestive system development | 4 | 0.2002 | 0.00721202 | WNT5A, TRP | 160 | 34 | 13588 | 9.99117647 | 0.99981028 | 0.13310104 | 11.0255924 |
| GO:0001655~urogenital system development | 7 | 0.35035 | 0.00739475 | WNT5A, FGF | 160 | 146 | 13588 | 4.07174658 | 0.99984743 | 0.13416776 | 11.2895294 |
| GO:0040007~growth | 8 | 0.4004 | 0.00762435 | WNT5A, GPX | 160 | 193 | 13588 | 3.52020725 | 0.99988399 | 0.13597909 | 11.6201318 |
| GO:0006468~protein amino acid phosphorylation | 16 | 0.8008 | 0.00809552 | WNT5A, FGF | 160 | 640 | 13588 | 2.123125 | 0.99993388 | 0.1416674 | 12.2949631 |
| GO:0048598~embryonic morphogenesis | 11 | 0.55055 | 0.00965116 | WNT5A, FGF | 160 | 359 | 13588 | 2.60215877 | 0.99998969 | 0.16423988 | 14.4888268 |
| GO:0030324~lung development | 6 | 0.3003 | 0.00977332 | WNT5A, FGF | 160 | 111 | 13588 | 4.59054054 | 0.99999109 | 0.16381088 | 14.6589083 |
| GO:0060562~epithelial tube morphogenesis | 6 | 0.3003 | 0.00977332 | WNT5A, ESR | 160 | 111 | 13588 | 4.59054054 | 0.99999109 | 0.16381088 | 14.6589083 |
| GO:0021537~telencephalon development | 5 | 0.25025 | 0.01002942 | ARX, ZEB2, T | 160 | 72 | 13588 | 5.89756944 | 0.99999344 | 0.16542289 | 15.0144439 |
| GO:0030323~respiratory tube development | 6 | 0.3003 | 0.01050752 | WNT5A, FGF | 160 | 113 | 13588 | 4.50929204 | 0.9999963 | 0.17028051 | 15.6744484 |
| GO:0007242~intracellular signaling cascade | 20 | 1.001 | 0.01053601 | WNT5A, CNK | 160 | 915 | 13588 | 1.85628415 | 0.99999642 | 0.16841669 | 15.7136246 |
| GO:0043009~chordate embryonic development | 12 | 0.6006 | 0.01064411 | LIF, FGFR1, S | 160 | 421 | 13588 | 2.42066508 | 0.99999686 | 0.16775274 | 15.8621152 |
| GO:0009792~embryonic development ending in birth or egg hatchin | 12 | 0.6006 | 0.01136629 | LIF, FGFR1, S | 160 | 425 | 13588 | 2.39788235 | 0.99999868 | 0.17580958 | 16.8479084 |
| GO:0035272~exocrine system development | 4 | 0.2002 | 0.01210635 | FGFR1, DAG1 | 160 | 41 | 13588 | 8.28536585 | 0.99999945 | 0.18381816 | 17.8468399 |
| GO:0060348~bone development | 6 | 0.3003 | 0.01250644 | IGSF10, SATE | 160 | 118 | 13588 | 4.31822034 | 0.99999966 | 0.1869464 | 18.3821866 |
| GO:0030855~epithelial cell differentiation | 6 | 0.3003 | 0.01474878 | GPX1, SPRR1 | 160 | 123 | 13588 | 4.14268293 | 0.99999998 | 0.21415424 | 21.3225384 |
| GO:0060541~respiratory system development | 6 | 0.3003 | 0.01522752 | WNT5A, FGF | 160 | 124 | 13588 | 4.10927419 | 0.99999999 | 0.21769791 | 21.9372809 |
| GO:0048706~embryonic skeletal system development | 5 | 0.25025 | 0.01623816 | SATB2, HOXE | 160 | 83 | 13588 | 5.11596386 | 1 | 0.2277528 | 23.2202956 |
| GO:0008285~negative regulation of cell proliferation | 8 | 0.4004 | 0.01631841 | WNT5A, BDN | 160 | 224 | 13588 | 3.03303571 | 1 | 0.22610628 | 23.3213149 |
| GO:0010740~positive regulation of protein kinase cascade | 5 | 0.25025 | 0.01970867 | LIF, FGFR1, G | 160 | 88 | 13588 | 4.82528409 | 1 | 0.2636716 | 27.4771138 |
| GO:0045596~negative regulation of cell differentiation | 7 | 0.35035 | 0.02018479 | LIF, BDNF, CC | 160 | 182 | 13588 | 3.26634615 | 1 | 0.26620775 | 28.0435293 |
| GO:0060444~branching involved in mammary gland duct morphoger | 3 | 0.15015 | 0.02042189 | WNT5A, ESR | 160 | 19 | 13588 | 13.4092105 | 1 | 0.26599421 | 28.324043 |
| GO:0043524~negative regulation of neuron apoptosis | 4 | 0.2002 | 0.02062923 | BDNF, RHOA | 160 | 50 | 13588 | 6.794 | 1 | 0.26545625 | 28.5685003 |
| GO:0007160~cell-matrix adhesion | 4 | 0.2002 | 0.02062923 | FREM1, RHO | 160 | 50 | 13588 | 6.794 | 1 | 0.26545625 | 28.5685003 |
| GO:0042325~regulation of phosphorylation | 9 | 0.45045 | 0.02081659 | LIF, DGKB, ZE | 160 | 290 | 13588 | 2.63560345 | 1 | 0.26471254 | 28.7887428 |
| GO:0016310~phosphorylation | 16 | 0.8008 | 0.02116005 | WNT5A, FGF | 160 | 718 | 13588 | 1.89247911 | 1 | 0.26567921 | 29.1908076 |
| GO:0032147~activation of protein kinase activity | 4 | 0.2002 | 0.02173466 | DGKB, GRM1 | 160 | 51 | 13588 | 6.66078431 | 1 | 0.26909022 | 29.8587194 |
| GO:0045859~regulation of protein kinase activity | 7 | 0.35035 | 0.02216157 | DGKB, ZEB2, | 160 | 186 | 13588 | 3.19610215 | 1 | 0.27085736 | 30.3511114 |
| GO:0043066~negative regulation of apoptosis | 8 | 0.4004 | 0.02239398 | FGFR1, GPX1 | 160 | 239 | 13588 | 2.84267782 | 1 | 0.2705619 | 30.6178131 |
| GO:0002062~chondrocyte differentiation | 3 | 0.15015 | 0.02470082 | FGFR1, HSPG | 160 | 21 | 13588 | 12.1321429 | 1 | 0.29131032 | 33.2135177 |
| GO:0043069~negative regulation of programmed cell death | 8 | 0.4004 | 0.02472731 | FGFR1, GPX1 | 160 | 244 | 13588 | 2.78442623 | 1 | 0.28876275 | 33.2427848 |
| GO:0006793~phosphorus metabolic process | 18 | 0.9009 | 0.02494366 | WNT5A, FGF | 160 | 866 | 13588 | 1.76518476 | 1 | 0.2881316 | 33.4814083 |
| GO:0006796~phosphate metabolic process | 18 | 0.9009 | 0.02494366 | WNT5A, FGF | 160 | 866 | 13588 | 1.76518476 | 1 | 0.2881316 | 33.4814083 |
| GO:0060548~negative regulation of cell death | 8 | 0.4004 | 0.02521329 | FGFR1, GPX1 | 160 | 245 | 13588 | 2.77306122 | 1 | 0.28803226 | 33.7776547 |
| GO:0019220~regulation of phosphate metabolic process | 9 | 0.45045 | 0.0253063 | LIF, DGKB, ZE | 160 | 301 | 13588 | 2.53928571 | 1 | 0.28623628 | 33.8795627 |
| GO:0051174~regulation of phosphorus metabolic process | 9 | 0.45045 | 0.0253063 | LIF, DGKB, ZE | 160 | 301 | 13588 | 2.53928571 | 1 | 0.28623628 | 33.8795627 |
| GO:0043549~regulation of kinase activity | 7 | 0.35035 | 0.02536634 | DGKB, ZEB2, | 160 | 192 | 13588 | 3.09622396 | 1 | 0.28416053 | 33.945275 |
| GO:0007169~transmembrane receptor protein tyrosine kinase signa | 7 | 0.35035 | 0.02536634 | IGSF10, EPHA | 160 | 192 | 13588 | 3.09622396 | 1 | 0.28416053 | 33.945275 |
| GO:0009069~serine family amino acid metabolic process | 3 | 0.15015 | 0.02696528 | SHMT1, CTH, | 160 | 22 | 13588 | 11.5806818 | 1 | 0.29657651 | 35.6727161 |
| GO:0010628~positive regulation of gene expression | 12 | 0.6006 | 0.02838019 | LIF, SATB2, S | 160 | 488 | 13588 | 2.08831967 | 1 | 0.30687126 | 37.1659081 |
| GO:0048514~blood vessel morphogenesis | 7 | 0.35035 | 0.02897413 | FGFR1, GPX1 | 160 | 198 | 13588 | 3.00239899 | 1 | 0.30950196 | 37.7829679 |
| GO:0031589~cell-substrate adhesion | 4 | 0.2002 | 0.02903583 | FREM1, RHO | 160 | 57 | 13588 | 5.95964912 | 1 | 0.3073537 | 37.8467466 |
| GO:0009100~glycoprotein metabolic process | 6 | 0.3003 | 0.02925987 | ST6GALNAC5 | 160 | 147 | 13588 | 3.46632653 | 1 | 0.30667512 | 38.0778087 |
| GO:0031099~regeneration | 3 | 0.15015 | 0.02931012 | LIF, GPX1, NE | 160 | 23 | 13588 | 11.0771739 | 1 | 0.30449183 | 38.1295152 |
| GO:0051338~regulation of transferase activity | 7 | 0.35035 | 0.02978509 | DGKB, ZEB2, | 160 | 199 | 13588 | 2.98731156 | 1 | 0.3060258 | 38.6163277 |
| GO:0007423~sensory organ development | 8 | 0.4004 | 0.03156405 | FGFR1, BDNF | 160 | 257 | 13588 | 2.64357977 | 1 | 0.31858197 | 40.4078741 |
| GO:0060686~negative regulation of prostatic bud formation | 2 | 0.1001 | 0.03469785 | WNT5A, BMI | 160 | 3 | 13588 | 56.6166667 | 1 | 0.34171625 | 43.4451092 |
| GO:0001503~ossification | 5 | 0.25025 | 0.0357967 | IGSF10, SATE | 160 | 106 | 13588 | 4.00589623 | 1 | 0.34775273 | 44.4752569 |
| GO:0060603~mammary gland duct morphogenesis | 3 | 0.15015 | 0.03680542 | WNT5A, ESR | 160 | 26 | 13588 | 9.79903846 | 1 | 0.35292443 | 45.405398 |
| GO:0050673~epithelial cell proliferation | 3 | 0.15015 | 0.03680542 | WNT5A, CCN | 160 | 26 | 13588 | 9.79903846 | 1 | 0.35292443 | 45.405398 |
| GO:0001822~kidney development | 5 | 0.25025 | 0.03686017 | FGFR1, BDNF | 160 | 107 | 13588 | 3.96845794 | 1 | 0.35060845 | 45.455459 |
| GO:0008361~regulation of cell size | 5 | 0.25025 | 0.03794174 | TRO, ESR1, T | 160 | 108 | 13588 | 3.93171296 | 1 | 0.35619534 | 46.4356758 |
| GO:0032535~regulation of cellular component size | 6 | 0.3003 | 0.04074339 | TRO, ESR1, T | 160 | 161 | 13588 | 3.16490683 | 1 | 0.37440486 | 48.8985148 |
| GO:0007167~enzyme linked receptor protein signaling pathway | 8 | 0.4004 | 0.04159103 | IGSF10, EPHA | 160 | 273 | 13588 | 2.48864469 | 1 | 0.37780475 | 49.6224538 |
| GO:0051173~positive regulation of nitrogen compound metabolic pr | 12 | 0.6006 | 0.04492762 | LIF, SATB2, U | 160 | 526 | 13588 | 1.93745247 | 1 | 0.39869736 | 52.3796999 |
| GO:0046903~secretion | 7 | 0.35035 | 0.0455646 | BDNF, NLRC4 | 160 | 221 | 13588 | 2.68993213 | 1 | 0.40026133 | 52.8897189 |
| GO:0060750~epithelial cell proliferation involved in mammary gland | 2 | 0.1001 | 0.04599583 | WNT5A, ESR | 160 | 4 | 13588 | 42.4625 | 1 | 0.40039234 | 53.2320853 |
| GO:0007507~heart development | 7 | 0.35035 | 0.04723816 | HSPG2, SEMA | 160 | 223 | 13588 | 2.66580717 | 1 | 0.40598798 | 54.2053988 |
| GO:0009967~positive regulation of signal transduction | 6 | 0.3003 | 0.05156997 | LIF, FGFR1, G | 160 | 172 | 13588 | 2.9625 | 1 | 0.43150955 | 57.4526081 |
| GO:0045941~positive regulation of transcription | 11 | 0.55055 | 0.0525144 | LIF, SATB2, S | 160 | 475 | 13588 | 1.96668421 | 1 | 0.43462172 | 58.1312815 |
| GO:0007517~muscle organ development | 6 | 0.3003 | 0.05587917 | LIF, GPX1, RH | 160 | 176 | 13588 | 2.89517045 | 1 | 0.45255301 | 60.4675961 |
| GO:0006355~regulation of transcription, DNA-dependent | 25 | 1.25125 | 0.05593209 | BACH2, SOX2 | 160 | 1465 | 13588 | 1.44923208 | 1 | 0.44997243 | 60.5033479 |
| GO:0060443~mammary gland morphogenesis | 3 | 0.15015 | 0.05672162 | WNT5A, ESR | 160 | 33 | 13588 | 7.72045455 | 1 | 0.45184795 | 61.0330953 |

| | | | | | | | | | | | |
|---|---|---|---|---|---|---|---|---|---|---|---|
| GO:0048546~digestive tract morphogenesis | 3 | 0.15015 | 0.05672162 | WNT5A, TRP | 160 | 33 | 13588 | 7.72045455 | 1 | 0.45184795 | 61.0330953 |
| GO:0046668~regulation of retinal cell programmed cell death | 2 | 0.1001 | 0.0571624 | BDNF, FGF2 | 160 | 5 | 13588 | 33.97 | 1 | 0.45162201 | 61.3259431 |
| GO:0060751~mammary gland duct branch elongation | 2 | 0.1001 | 0.0571624 | WNT5A, ESR | 160 | 5 | 13588 | 33.97 | 1 | 0.45162201 | 61.3259431 |
| GO:0045860~positive regulation of protein kinase activity | 5 | 0.25025 | 0.05770693 | DGKB, ZEB2, | 160 | 124 | 13588 | 3.42439516 | 1 | 0.45201193 | 61.6848669 |
| GO:0007268~synaptic transmission | 6 | 0.3003 | 0.0587503 | SLC17A6, LYN | 160 | 178 | 13588 | 2.86264045 | 1 | 0.45530031 | 62.3638859 |
| GO:0031328~positive regulation of cellular biosynthetic process | 12 | 0.6006 | 0.05949384 | LIF, SATB2, U | 160 | 552 | 13588 | 1.84619565 | 1 | 0.45680027 | 62.8408686 |
| GO:0014706~striated muscle tissue development | 5 | 0.25025 | 0.06192449 | GPX1, RHOA, | 160 | 127 | 13588 | 3.34350394 | 1 | 0.46779422 | 64.3608762 |
| GO:0009891~positive regulation of biosynthetic process | 12 | 0.6006 | 0.06261683 | LIF, SATB2, U | 160 | 557 | 13588 | 1.82962298 | 1 | 0.46886388 | 64.7830371 |
| GO:0043067~regulation of programmed cell death | 12 | 0.6006 | 0.06454159 | FGFR1, GPX1 | 160 | 560 | 13588 | 1.81982143 | 1 | 0.47664669 | 65.9321651 |
| GO:0051252~regulation of RNA metabolic process | 25 | 1.25125 | 0.06460249 | BACH2, SOX2 | 160 | 1488 | 13588 | 1.42683132 | 1 | 0.47421397 | 65.9679412 |
| GO:0043010~camera-type eye development | 5 | 0.25025 | 0.06630091 | BMP7, MEIS1 | 160 | 130 | 13588 | 3.26634615 | 1 | 0.48057329 | 66.9516663 |
| GO:0048705~skeletal system morphogenesis | 5 | 0.25025 | 0.06630091 | SATB2, HOXE | 160 | 130 | 13588 | 3.26634615 | 1 | 0.48057329 | 66.9516663 |
| GO:0033674~positive regulation of kinase activity | 5 | 0.25025 | 0.06630091 | DGKB, ZEB2, | 160 | 130 | 13588 | 3.26634615 | 1 | 0.48057329 | 66.9516663 |
| GO:0010941~regulation of cell death | 12 | 0.6006 | 0.0665048 | FGFR1, GPX1 | 160 | 563 | 13588 | 1.81012433 | 1 | 0.47892324 | 67.0679485 |
| GO:0001568~blood vessel development | 7 | 0.35035 | 0.06711241 | FGFR1, GPX1 | 160 | 244 | 13588 | 2.43637295 | 1 | 0.47941534 | 67.4122058 |
| GO:0060685~regulation of prostatic bud formation | 2 | 0.1001 | 0.06819908 | WNT5A, BMI | 160 | 6 | 13588 | 28.3083333 | 1 | 0.4823876 | 68.0194928 |
| GO:0007442~hindgut morphogenesis | 2 | 0.1001 | 0.06819908 | WNT5A, GLI3 | 160 | 6 | 13588 | 28.3083333 | 1 | 0.4823876 | 68.0194928 |
| GO:0044093~positive regulation of molecular function | 8 | 0.4004 | 0.06828512 | NLRC4, DGKE | 160 | 306 | 13588 | 2.22026144 | 1 | 0.48016199 | 68.0671219 |
| GO:0016049~cell growth | 3 | 0.15015 | 0.06943315 | ESR1, DCX, D | 160 | 37 | 13588 | 6.88581081 | 1 | 0.483398 | 68.6962723 |
| GO:0001525~angiogenesis | 5 | 0.25025 | 0.07083454 | FGFR1, GPX1 | 160 | 133 | 13588 | 3.19266917 | 1 | 0.48784502 | 69.4485166 |
| GO:0010647~positive regulation of cell communication | 6 | 0.3003 | 0.07116724 | LIF, FGFR1, G | 160 | 189 | 13588 | 2.69603175 | 1 | 0.48688587 | 69.6245967 |
| GO:0021536~diencephalon development | 3 | 0.15015 | 0.07274457 | ARX, WNT5A | 160 | 38 | 13588 | 6.70460526 | 1 | 0.49208807 | 70.4465164 |
| GO:0001944~vasculature development | 7 | 0.35035 | 0.07356749 | FGFR1, GPX1 | 160 | 250 | 13588 | 2.3779 | 1 | 0.49351354 | 70.8669974 |
| GO:0051347~positive regulation of transferase activity | 5 | 0.25025 | 0.07394337 | DGKB, ZEB2, | 160 | 135 | 13588 | 3.14537037 | 1 | 0.49275784 | 71.0571819 |
| GO:0060537~muscle tissue development | 5 | 0.25025 | 0.07552348 | GPX1, RHOA, | 160 | 136 | 13588 | 3.12224265 | 1 | 0.49777967 | 71.8440199 |
| GO:0043405~regulation of MAP kinase activity | 4 | 0.2002 | 0.07552636 | ZEB2, TRP73, | 160 | 84 | 13588 | 4.04404762 | 1 | 0.49524363 | 71.845437 |
| GO:0030111~regulation of Wnt receptor signaling pathway | 3 | 0.15015 | 0.07610609 | DKK3, CCND1 | 160 | 39 | 13588 | 6.53269231 | 1 | 0.49546108 | 72.1290432 |
| GO:0016358~dendrite development | 3 | 0.15015 | 0.07610609 | BDNF, DCX, D | 160 | 39 | 13588 | 6.53269231 | 1 | 0.49546108 | 72.1290432 |
| GO:0045935~positive regulation of nucleobase, nucleoside, nucleoti | 11 | 0.55055 | 0.07659981 | LIF, SATB2, S | 160 | 510 | 13588 | 1.83171569 | 1 | 0.49527374 | 72.3684537 |
| GO:0060687~regulation of branching involved in prostate gland mor | 2 | 0.1001 | 0.07910736 | ESR1, BMP7 | 160 | 7 | 13588 | 24.2642857 | 1 | 0.50439831 | 73.5549222 |
| GO:0021772~olfactory bulb development | 2 | 0.1001 | 0.07910736 | ARX, NR2E1 | 160 | 7 | 13588 | 24.2642857 | 1 | 0.50439831 | 73.5549222 |
| GO:0043085~positive regulation of catalytic activity | 7 | 0.35035 | 0.08629486 | NLRC4, DGKE | 160 | 261 | 13588 | 2.27768199 | 1 | 0.53384398 | 76.6964226 |
| GO:0001657~ureteric bud development | 3 | 0.15015 | 0.08647528 | FGFR1, BDNF | 160 | 42 | 13588 | 6.06607143 | 1 | 0.53209035 | 76.770575 |
| GO:0021988~olfactory lobe development | 2 | 0.1001 | 0.08988875 | ARX, NR2E1 | 160 | 8 | 13588 | 21.23125 | 1 | 0.54403798 | 78.1325452 |
| GO:0010557~positive regulation of macromolecule biosynthetic pro | 11 | 0.55055 | 0.09304238 | LIF, SATB2, S | 160 | 530 | 13588 | 1.76259434 | 1 | 0.55451593 | 79.3239379 |
| GO:0045165~cell fate commitment | 5 | 0.25025 | 0.09407774 | FGFR1, SOX5 | 160 | 147 | 13588 | 2.88860544 | 1 | 0.55619511 | 79.7016145 |
| GO:0046328~regulation of JNK cascade | 3 | 0.15015 | 0.09723919 | EDA2R, ZEB2 | 160 | 45 | 13588 | 5.66166667 | 1 | 0.56626136 | 80.8151788 |
| GO:0042476~odontogenesis | 3 | 0.15015 | 0.09723919 | BMP7, GLI3, | 160 | 45 | 13588 | 5.66166667 | 1 | 0.56626136 | 80.8151788 |
